# Automated 3D multi-color single-molecule localization microscopy

**DOI:** 10.1101/2023.10.23.563122

**Authors:** Rory M. Power, Aline Tschanz, Timo Zimmermann, Jonas Ries

## Abstract

Since its inception, single molecule localization microscopy (SMLM) has enabled imaging scientists to visualize biological structures with unprecedented resolution. Particularly powerful implementations capable of 3D, multi-color and high-throughput imaging have yielded key biological insights although widespread access to such technologies has been limited. The purpose of this protocol is to provide a guide for interested researchers to establish high-end SMLM in their laboratories. We detail the initial configuration and subsequent assembly of the SMLM, including instructions for alignment of all optical pathways, software/hardware integration and operation of the instrument. We describe validation steps including the preparation and imaging of test- and biological samples with structures of well-defined geometry and assist the user in troubleshooting and benchmarking performance. Additionally, we provide a walkthrough of the reconstruction of a super-resolved dataset from acquired raw images using the Super-resolution Microscopy Analysis Platform (SMAP). Depending on the instrument configuration, the cost of components is in the range $80,000 – 160,000, a fraction of the cost of a commercial instrument. A builder with some experience of optical systems is expected to require 3 - 6 months from the start of system construction to attain high-quality 3D and multi-color biological images.

## Introduction

Single-molecule localization microscopy (SMLM) is a super-resolution fluorescence microscopy technique based on the precise 2D/3D localization of individual fluorophores. Unambiguous localization relies on spatial sparsity of the single molecule emitters. Following many rounds of imaging, each comprising a small fraction of all fluorophores, their positions are determined with a precision in the single nanometer range (under ideal conditions) and used to reconstruct a super-resolution image. The required sparsity of fluorophores may be regulated through photochemical on/off switching of the fluorophores, for example, in the cases of (fluorescence) photoactivation localization microscopy, ((f)PALM) [1,2] or (direct) stochastic optical reconstruction microscopy ((d)STORM) [3,4]. On-off binding of fluorophores provides an alternative route, for example, in DNA-based point-accumulation in nanoscale topology microscopy (DNA-PAINT) [5] target structures are tagged with short single-stranded DNA, and fluorescently-labelled complementary strands stochastically bind and detach to elicit a blinking effect.

Engineering labs operating highly-optimized custom microscopes have driven the development of SMLM with their creations typically outperforming commercially available variants in terms of image quality, stability and robustness. Furthermore, the custom-build approach provides agency in the design process, freedom from reliance on service technicians to maintain advanced imaging devices, and ability to upgrade hardware freely. The efforts of many to democratize SMLM technology have centered largely on developing particularly low cost systems [6–10]. While this is a worthwhile pursuit, such systems often omit powerful features, lack the convenience and feature set of turnkey systems, and generally do not reflect the current state-of-the-art. For more advanced implementations, the technical challenge associated with replicating a system with limited documentation and instruction limits the widespread application of the associated advances. Nevertheless, the benefits of the custom-build approach make dissemination of these devices of crucial importance for biological imaging.

### Development of the protocol

In this protocol, we seek to bridge the gap between entry-level open-source projects and costly, though not necessarily more capable, commercial systems by providing a detailed and complete guide to allow non-specialist labs to establish a single molecule localization microscope with state-of-the art performance and advanced features and apply it to their biological studies. To encourage builders to configure the instrument as they wish, the microscope presented is modular and configurable in design. The scheme outlined includes the necessary resources to construct and use an advanced SMLM system. All necessary validation steps, processes for acquiring benchmark images and their subsequent analysis are similarly included.

More than a decade at the forefront of SMLM development has provided a unique view on the desirable features of the associated microscope. While many technical developments from our lab and others have expanded the capabilities of SMLM, there is clearly a judgement to be made regarding cost/complexity vs. performance/usability. In this regard, we assert that the following features are suitable to cover most applications and user needs spanning (d)STORM, DNA-PAINT and PALM workflows and are well-suited to a multi-functional system covering many such applications.

The first feature concerns an inverted microscope body. In order to deliver a spatial resolution at least one order of magnitude better than in standard widefield epi-fluorescence microscopy, mechanical disturbances such as drift and vibration have to be minimized. A suitably custom machined body provides a substantial improvement over a commercial inverted microscope stand in this regard. We included long range (several mm) refocusing to accommodate different sample types, hot-swappable optics (e.g., dichroic mirrors), transmission illumination, and the controlled heating of the microscope for live-cell imaging under physiological conditions. These features are typically missing in custom microscopes, but have proven useful across the developmental lifetime of the microscope reported herein. We also include a focus lock system to minimize axial drift. While real-time lateral drift correction schemes have been reported [11], we correct it during post-processing.

From an illumination perspective, homogeneity is a primary concern but so too is the possibility for high quality TIRF illumination. Although schemes effectively demonstrating both exist [12–15], they have not been transferred to open-source systems with multiple laser lines. Achieving suitable irradiance of the sample in often power-hungry applications is of similar concern. Even when sufficient laser power and minimally-lossy schemes are available, statically sized illumination fields do not allow for optimization of the intensity^1^ with respect to the background generated nor avoid damage to expensive objective lenses where high power lasers are used to illuminate an unnecessarily large field of view. Particularly in multi-user settings, such field adjustable illumination, which maintains homogeneity and TIRF-compatibility are invaluable.

On the emission side, multi-color imaging is, of course, desirable and can be achieved serially. However, many powerful methods such as (d)STORM and smFRET (single-molecule Förster resonance energy transfer) require synchronous acquisition of multiple color channels to discriminate as many as 4 spectrally-overlapping far-red fluorophores [16] and elucidate close contacts between FRET-pairs, respectively [17,18]. Given the resolution of modern sCMOS cameras, image splitters forming a pair of images on a single camera provide such capabilities without sacrificing field of view and offer a doubling of imaging speed even for applications where serial acquisition is tolerable. Given the inherently 3D nature of biological samples, the ability to discriminate emitters based on depth information is critical to many SMLM applications and the related PSF-engineering approaches to do so have been among the most impactful extensions of the technology [19,20]. To date, however, open-source SMLM systems have not provided an option for 3D imaging which we view to be one of the key areas of innovation in SMLM. One exception in this regard, utilized a specialist sample preparation whereby the known geometry of a labelled microbead provides a fixed reference to the z-position of an emitter [7,21]. However, more optimized 3D imaging strategies, which offer a greater degree of control of the PSF shape have not been employed [20,22].

Advanced microscopes such as SMLM systems require suitable control systems in the form of software with an intuitive interface and powerful acquisition engine, electronics and hardware synchronization. Moreover, SMLM acquisitions are typically long, requiring tens of thousands of images to reconstruct a single super-resolution view of the sample. As such, and given the need for statistical significance and perturbations in biological studies, automation of SMLM plays a key role in its success. The supplied control systems and associated hardware must therefore be capable of unsupervised imaging, not only moving from point to point in a predetermined manner, but also performing quality checks and where appropriate modulating photoactivation parameters in line with on-the-fly detected emitter density. Naturally, the data and metadata structures should streamline subsequent analysis by being fully integrated with freely-available modern analysis software.

The capabilities of various open-source SMLM projects have been well-summarized as they relate to a recent implementation, the NanoPro 1.0 [23]. We do not replicate this discussion but extend the analysis of these systems in Supplementary Table 1: Features Comparison of Open-Source SMLM Systems. As can be seen from the feature list, the 3D-SMLM system reported herein delivers all desired features highlighted in Danial et al. [23], including several missing in the NanoPro 1.0 such as fully-integrated 3D capability. In addition, Supplementary Table 1 expands this wish list to consider many additional features. Ultimately, our 3D-SMLM has been designed to optimize performance and functionality while maintaining sensitivity to pricing such that the instrument represents a substantial saving compared to similar commercial systems. Labs wishing for an entry-level SMLM system without many features described are advised to seek out many of the excellent open-source SMLM projects, some of which are optimized for low cost [7–9] and ease of construction [6].

The instrument detailed herein, represents the 6^th^ generation refinement of our SMLM developments across more than a decade of development and features improvements to the optical, mechanical, electronic and computational designs reported previously [24–30]. These microscopes were instrumental for key biological discoveries [31–34]. Microscopes constructed from this scheme are currently in use in research and service settings at the European Molecular Biology Laboratory (EMBL), while implementation of previous generations has been carried out in several labs worldwide. We note that the most recent generation of the microscope has been developed at the EMBL Imaging Centre, as part of efforts to develop custom imaging technologies in partnership with EMBL developer labs. The particular challenges inherent in deployment of such technologies in a multi-user environment for advanced service provision require a greater flexibility of approach than would be the case for a single research lab with a narrower research focus. The aspects developed to serve the range of needs encountered are fully represented in the instrument described herein accordingly.

Successful implementation of the protocol is contingent on some familiarity with the construction of optical systems and a working understanding of SMLM. We expect a full-time builder would require 6 months to successfully implement the protocol described, culminating in the reconstruction of exemplary super-resolution images, while an expert builder may require as little as 3 months. A full list of materials including parts list (see Supplementary Table 2. Parts List and Supplementary Note 1: Using the parts list and understanding part names), CAD models and associated toleranced drawings, software, and electronic schematics can be found on GitHub (https://github.com/ries-lab/3DSMLM). CAD models are included in SolidWorks (Dassault Systèmes) assembly and part file format for those wishing to modify the design. For those wishing to follow the protocol as written, the CAD assemblies are also provided in eDrawing format, which can be viewed using a freely available eDrawing viewer (https://www.edrawingsviewer.com/download-edrawings). The viewer also allows suppression of components, configuration of the assemblies, and cut away views, all of which are helpful to follow the protocol. Toleranced drawings for the custom mechanical parts are provided as SolidWorks drawings and as .pdf for manufacture by a workshop with precision computer numerical control (CNC) milling capabilities. We advise that labs wishing to replicate the system first familiarize themselves with the assemblies and contact the EMBL mechanical and electronic workshops to discuss appropriate sourcing of parts and to seek direct correspondence with the authors before embarking on the protocol to clarify uncertainties and check for forthcoming updates.

We note that the parts provided at the time of publication are by no means final as the system undergoes continuous improvements and the online resources noted will be maintained accordingly. Furthermore, we encourage builders to engage in this effort by modifying or otherwise suggesting modifications of part files to further improve their functionality or suggest desirable features that could be developed.

### Overview of the automated 3D multi-color single molecule localization microscope

A CAD rendering of the microscope is shown in Figure 1. In essence, the SMLM comprises an inverted epi-fluorescence microscope constructed around a robust, yet flexible, custom microscope body. The body incorporates xy stages, a z-focusing piezo and long-range travel platform for the objective lens/piezo as well as switchable kinematic mounts for dichroics, housing for a tube lens and optional filters and, if specified, the capacity to operate at elevated temperatures suitable to mammalian systems. Samples mounted on coverslips are subject to widefield laser illumination via an objective lens, which collects fluorescence and separates the emitted light from the excitation source on a dichroic mirror. The emission path further comprises a tube lens and additional optics for spectral discrimination and image relaying to produce two synchronously captured color channels on a scientific complementary metal-oxide semiconductor (sCMOS) camera. For example, ratiometric imaging with two spectrally distinct channels may be employed to separate multiple spectrally overlapping far-red dyes suitable to dSTORM imaging. For 3D imaging, a distortion-free and tunable astigmatism module allows fine control over the extension of the PSF in two orthogonal axes either side of the native focus. As such, one may attain localization precisions as high as 2 nm in xy and 8 nm in z under idealized conditions [27] using a calibrated spline model of the PSF [25]. Excitation can be provided by single-mode or multi-mode laser sources, or a combination of the two, depending on specific requirements to yield a uniform illumination field using various homogenization schemes. The laser excitation can be centered/offset at the back focal plane of the imaging objective lens via a motorized mirror to provide epi-/HILO/TIRF illumination. A focus lock system is included, comprising a near infrared (NIR) laser at 808 nm under total internal reflection from the coverslip and subsequent height sensitive detection using a position-sensitive detector with feedback to the piezo-coupled objective lens. Sample throughput presents a challenge to SMLM since large numbers of camera frames are required to reconstruct a single image and constant user input may be required during the experiments. Thus, the microscope provides automated acquisition capabilities comprising multiple field of views (FOVs) and featuring real-time feedback to activation lasers to regulate the number of emitters during a single exposure for unsupervised acquisition of data.

**Figure 1.**
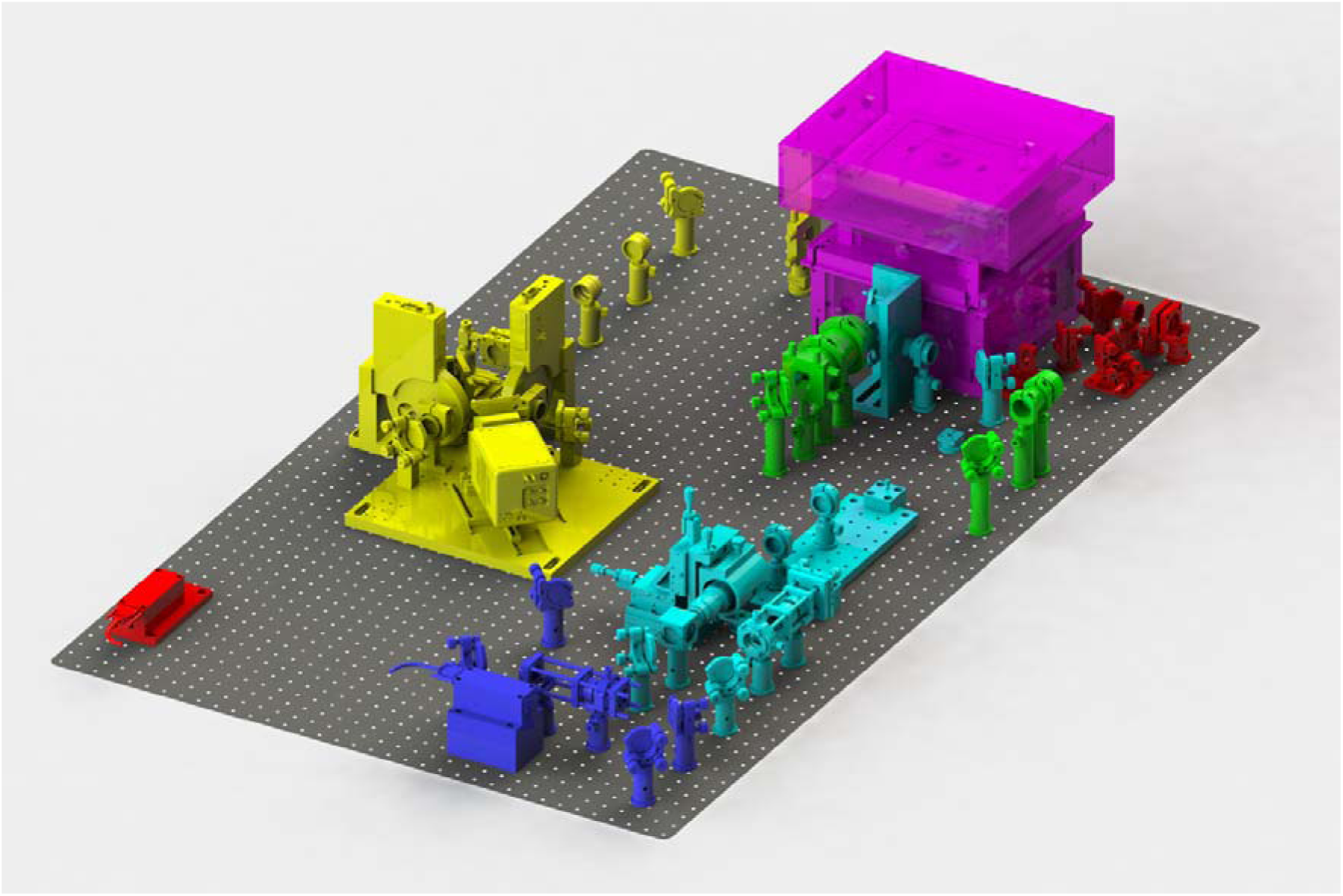
A full rendering of the 3D-SMLM. The system features a custom microscope body (magenta), emission path with 3D and synchronous multi-channel/color imaging capabilities (yellow) and focus lock path (red). Multiple illumination sources are available: single-mode illumination path with fiber-coupled laser engine source (cyan) ii) single-mode booster laser (blue), multi-mode illumination path (green). The configuration shown includes all optional modules. For scale, the underlying optical breadboard measures 1.5 × 0.9 m.

The microscope comprises four optomechanical subsystems: the body, and the excitation, emission and focus lock paths, described below (see Figure 1). The beam paths are shown in Figure 2. There is some flexibility in the system to tune the configuration to the end-user’s needs, budget and proficiencies in optical system construction. The optomechanical configuration must be determined before sourcing commercial and custom parts. For selection of costly hardware elements such as stages, cameras and lasers, please refer to the respective section under Materials. The CAD assembly files provided are configurable allowing the builder to construct the appropriate system *in silico* before proceeding. The parts list is similarly structured to allow easy identification of components required for different configurations. Supplementary Table 3 is provided to estimate the final cost for the system.

**Figure 2.**
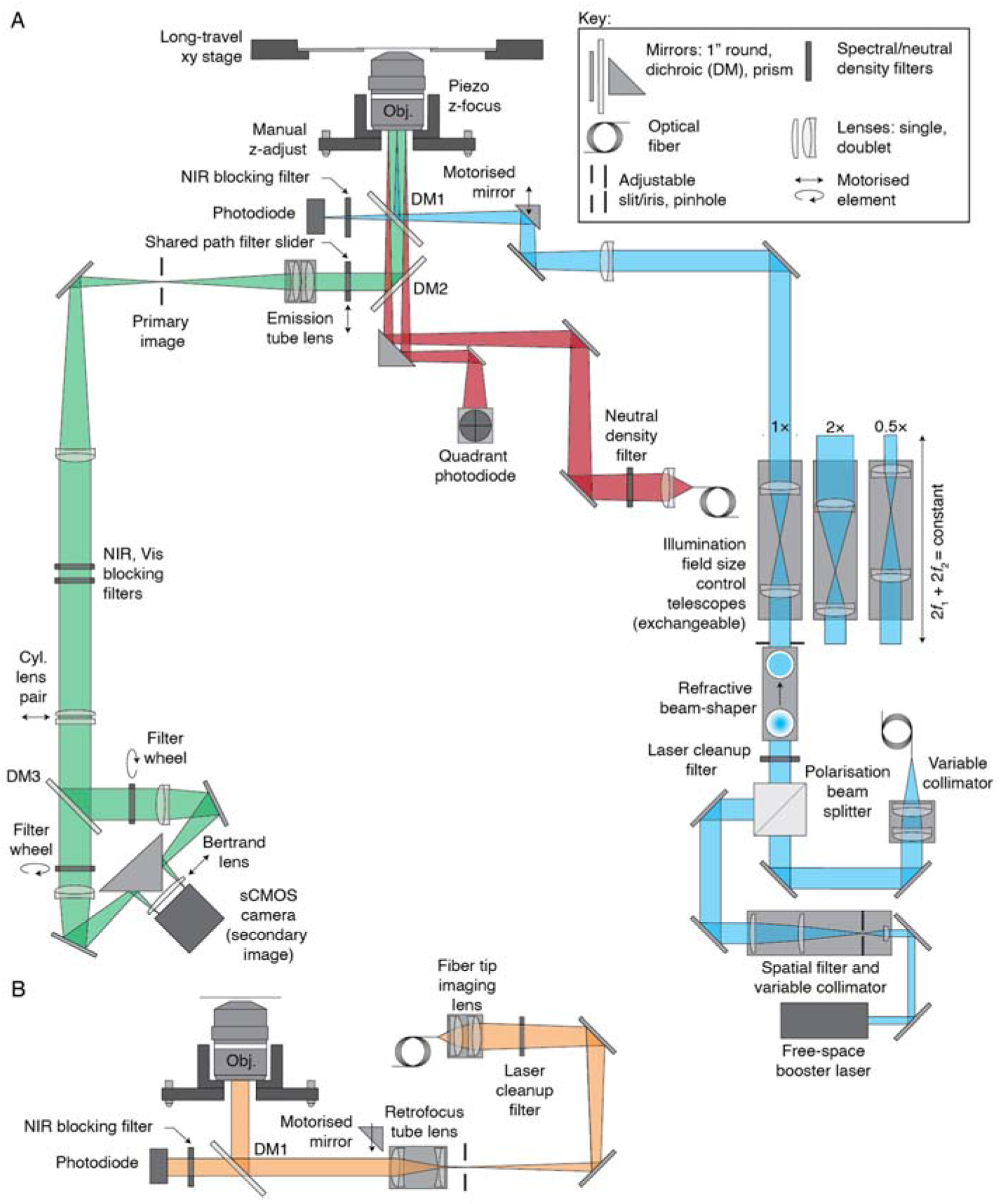
Schematic of the optical and motion control aspects of the 3D-SMLM. A) Single-mode illumination path(s) (blue): The single-mode laser(s) are spatially filtered, expanded and homogenized. The resulting flat-top intensity profile is further expanded by exchangeable telescopes to control the size of the illumination field. Setting the angle for TIRF/HILO illumination is accomplished by positioning of a motorized mirror. The laser power is monitored in real time by measuring the <1% transmission of the laser lines at the illumination dichroic mirror. Emission path (green): A primary image is produced by a tube lens, restricted in spatial extent by a slit and relayed in a 4f configuration via a spectrally discriminative image splitter to form two secondary images on an sCMOS camera. A variable astigmatic lens (cylindrical lens pair) can be removed/inserted for 2D/3D imaging, while a Bertrand lens allows viewing of the objective back focal plane. Focus lock path (red): The focus lock laser is separated from emitted fluorescence by the emission dichroic mirror, focused to the objective back focal plane and launched obliquely to return under the TIRF condition allowing axial drift of the coverslip relative to the objective lens to be measured via a quadrant photodiode and corrected via the control loop of the piezo flexure stage mounting the objective. B) For clarity the multi-mode laser illumination path (orange) has been shown separately but all modules are co-compatible. A magnified image of the tip of a square core multimode fiber (coupled to a multi-mode laser engine) is produced and re-imaged onto the sample via a tube lens in a retrofocus arrangement and the objective lens. The multi-mode laser is coupled into the microscope by translation of the motorized mirror (HILO/TIRF) fully out of the beam path

#### The body

The body comprises an inverted microscope frame constructed from aluminum, stainless steel, brass and plastic (detailed in the technical drawings). It has been designed with several criteria in mind. First, it should provide an extremely stable platform suited to SMLM (see Protocol: Performing drift and vibration checks). Second, the body should retain much of the flexibility of a commercial inverted microscope stand and be able to accommodate multiple sample preparations/formats. Thirdly, the body should be easily assembled from appropriately toleranced custom parts using dowel pins to constrain the placement of elements.

Considering the specifics of the body, a highly rigid square base and frame provides up to four optical ports for the emission, excitation and focus lock optics. A removable cover of the body mounts a white LED ring suited to brightfield imaging. Below lies an xy microscope stage. In the base configuration, a Smaract SOM-12090 is specified but mounting points are included for legacy Smaract stages, which are in use on all previous generations of the microscope. With some adaptation of part files and assuming existence of a device adapter for µManager, other stages can also be used. Below the stages sits an objective lens long-range travel stage held in place under spring tension. The stage can be mechanically-driven over several mm in z by a system of belt-coupled fine-pitch actuators. A piezo flexure stage (Physik Instrumente PIFOC P-726.1CD) sits atop the stage and mounts the objective. Below the objective stage is an optional recess for a heating foil and temperature probe. The lower half of the body is thermally isolated from the top to obviate thermal drift. An optical mounting pillar is situated at the core of the body in the lower section and provides a single interface to the dichroic and broadband mirrors routing laser/emitted light into or out of the optical ports on the side of the body. The pillar and associated optical mounts are available in right- and left-handed versions to allow the microscope to be constructed in the desired orientation on the optical table by switching the excitation optical port between two opposing sides of the body. A primary dichroic mirror (DM1, TIRF quality) is situated at the top of the pillar below the isolating layer, which couples in the excitation light while passing the longer wave emission and focus locking laser. Below this sits a secondary dichroic mirror (DM2, super-resolution imaging quality), which reflects the emitted light towards the tube lens (via an optional piezo resonant slider, COTS-ELL9, in which additional filters may be placed as desired) and again passes the longwave focus lock laser. A subsequent broadband (NIR coated) prism mirror below DM2 reflects the focus lock laser on its way in and out of the body. See Supplementary Note 2: Selecting spectrally discriminative optics for a list of filters currently used on our most recent generations of the microscope.

#### Illumination path

The illumination path may be configured by the builder to include suitably homogenized single-mode and/or multi-mode laser sources. These sources can be combined and together comprise the illumination path. The rationale for offering a range of options in this regard relates to the properties of single- and multi-mode lasers as well as the desire to be able to configure highly capable systems to suit a range of budgetary constraints. Single-mode lasers require homogenization to achieve a uniform illumination field. Often this is simply achieved using a field aperture to apodize the Gaussian profile and pass only the approximately uniform core of the beam. While this is a workable solution, it results in substantial loss of power from the Gaussian input. For example, achieving a uniformity of ± 10%, results in a loss of ca. 90% of the beam power. Conversely, the coupling of light into a multi-mode fiber, combined with appropriate mode scrambling, produces a highly homogeneous illumination spot with only fiber insertion and transmission losses to consider. Nevertheless, single-mode laser systems allow for high-fidelity TIRF illumination since they can be focused to a very small spot on the back focal plane of the imaging objective lens, while multi-mode schemes are generally incompatible with high-quality TIRF illumination. The scheme outlined permits the combination of up to two TIRF compatible single-mode sources and one multi-mode source. The single-mode sources are homogenized by reshaping rather than apodizing the Gaussian beam with only negligible loss of power. The following description considers multiple laser sources:

##### Single-mode laser engine

the fiber-coupled output of the single-mode laser engine is expanded, and passed by an optional polarizing beam splitter that is included for booster laser integration (see below). The Gaussian beam is subsequently homogenized by a refractive beam shaper. An iris, placed at the exit aperture of the beam shaper, is conjugated to the object via two 4f systems. The first comprises one of four exchangeable telescopic systems (4f Keplerian arrangement), which allow the homogenized illumination field area to be varied by a factor of ∼32.5× (0.5 to 2.85× magnification to achieve Ill13.5 - 77 µm homogenized illumination fields). A piezo resonant stage actuated mirror situated intermediate within the second 4f system (tube lens/objective lens) allows tuning of the illumination angle for epi-/HILO/TIRF illumination. The lasers are coupled into the objective lens via the primary dichroic mirror (DM1) to reconstitute a variable-sized homogenized illumination field. A laser clean-up filter installed in the shared path with the optional booster laser removes any contribution from wavelengths outside a narrow window defined by the laser line that otherwise would contribute to background.

##### Single-mode booster laser

The optional free-space booster laser, typically at 640 nm, is spatially filtered and expanded before being steered to and combined with the common optical path at the polarizing beam splitter.

##### Multi-mode laser engine

Additionally/alternatively, multimode excitation can be coupled into the microscope from a custom low-cost laser engine [35]. The multimode path consists of a compound achromatic lens that projects a magnified image of the square multimode fiber tip onto an adjustable slit. This image is subsequently contracted and re-imaged onto the object via a tube lens and the objective lens. The tube lens is of a retrofocus type using only commercial parts. The retrofocus design allows the lens focal length to remain relatively short (f = 125 mm), with a much longer working distance on the infinite conjugate side (350 mm) such that it can be placed outside the body while maintaining strong coupling of the multi-mode laser into the objective lens. When using the multi-mode illumination, the piezo resonant stage actuated mirror used to position the single-mode beam(s) at the back focal plane for epi-/HILO/TIRF illumination is moved fully out of the beam path, thus passing the multi-mode lasers.

#### Emission path

The emission path configuration is largely fixed, although the selection of dichroic mirrors and filters is at the discretion of the user to suit the application in question (See Supplementary Note 2: Selecting spectrally discriminative optics). The optical scheme has been developed to work with 45 mm parfocal length (mounting flange to object side focus) objective lenses, specifically those that do not rely on a paired tube lens to correct residual chromatic aberrations (Olympus and Nikon are major manufacturers conforming to this design principle, whereas Leica and Zeiss opt to use the tube lens for chromatic correction). Moreover, to achieve a suitable pixel size (when demagnified into the object space), we specify the use of one of three objective lenses (all Olympus, RMS-threaded): 100x/1.5 oil (UPLAPO100XOHR), 100x/1.35 silicone oil, (UPLSAPO100XS) 100x/1.45 oil (UPLXAPO100XO). Note, that the latter is untested but provides a comparatively inexpensive option with regard to the 100x/1.5 oil lens and is expected to provide good performance. With limited modification of the part files, it is possible to use objective lenses from other manufacturers (See Supplementary Note 2: Using non-specified objective lenses). Emitted fluorescence is collected by the objective lens, passes the primary dichroic mirror (DM1) and is reflected by the secondary dichroic mirror (DM2). An optional motorized filter slider can be incorporated into the infinity space before a tube lens produces a primary image. This image is limited in spatial extent by an adjustable slit to ensure that the multiple color channels do not spatially overlap on the camera. A first relay lens restores the image to an infinite conjugate and produces an accessible conjugate to the objective back focal plane (BFP) where a variable astigmatism module comprising a pair of rotationally offset cylindrical lenses can be situated for 3D imaging. Placement of the astigmatism module at the BFP, affords minimized distortion and provides extensibility by maintaining compatibility with advanced PSF-engineering approaches using deformable mirrors (e.g., for aberration correction [36,37]) or phase masks (e.g., for double-helix PSFs [20]). A subsequent tertiary dichroic mirror (DM3) in the reconstituted infinity space, splits the emitted light between short and longwave channels, each of which includes a motorized filter wheel. Note, that these channels will often be referred to as reflected or transmitted based on their reflection/transmission at DM3. This should not be confused with reflective or transmissive imaging that generate contrast based on scattering and absorption of light by the sample. A second relay lens produces secondary images in the two pathways, which are recombined by a knife-edge mirror to coincide with the camera chip. Note, it is possible to perform 3D imaging via the biplane method whereby two images with a small defocus offset are used to attribute axial position to an emitter. Biplane imaging is beyond the scope of this article but it is worthy of note that with small modifications it is can be realized with the 3D-SMLM. In this case, one would omit the astigmatic lens and associated components, typically exchange DM3 for a 50:50 beam splitter (although biplane/ratiometric imaging has been reported using a dichroic mirror with the two channels encoding both depth and emitter information [38]) and appropriately defocus one of the second relay lenses.

A Bertrand lens may be introduced in front of the camera via a motorized stage, thus producing an image of the back focal plane on the camera, which aids in observing unwanted bubbles in immersion oils and alignment of the lasers more generally. The emission path has been designed using ray-tracing software (Zemax, OpticStudio) to achieve diffraction limited performance over a large field of view and for control of the 3D point spread function (see Supplementary Figures 1 - 3).

#### Focus lock path

The focus lock path comprises a NIR laser, typically 785 or 808 nm, which is steered to pass through the secondary and primary dichroic mirrors (DM2, DM1) with its focus laterally offset at the back focal plane to incline the collimated beam at the coverslip/sample boundary. In the case of oil objectives where the refractive index contrast between the immersion media (n = 1.51) and sample (n = 1.33 - 1.4) is substantial, the inclination is set to fulfil the condition of total internal reflection resulting in a strong focus lock signal. In the case of silicone objective lenses for which the refractive index contrast between the immersion media (n = 1.406) and sample is smaller, the inclination is set to maximize Fresnel reflections, which increase with incident angle. Such reflections are weaker but still easy to detect. In either case, the reflected light passes out of the objective on the opposing side of the back focal plane and is then spatially separated from the incoming beam to arrive at a position sensitive detector. The detector captures any change in height between the objective and coverslip as a positional offset and provides a corrective signal to the piezo flexure stage driving the objective lens.

#### Control and synchronization

Control of the microscope including previewing and acquisition functions is carried out using the open-source µManager software [39] and a custom htSMLM [26] (high-throughput single molecule localization microscopy) graphical user interface (GUI). µManager allows common hardware functions to be mapped to specific core functionalities providing great flexibility to specific hardware choices. As long as the hardware chosen permits the required functionality, and a device adapter for the hardware is available, the user may deviate from the specific hardware choices outlined in the protocol (see Supplementary Notes 3 - 6, Using non-specified objective lenses/single-mode lasers and laser-engines/XY stages/cameras). Pre-compiled versions of the htSMLM GUI and device adapters, which are not available in the core µManager build, are provided as part of the protocol. A guide to configuration of µManager and the htSMLM GUI is provided in the protocol. The microscope makes use of the open-source microFPGA project for synchronization of hardware components and for monitoring of signals related to the microscope focus lock system. Here, we provide microscope-specific configuration files for the FPGA, board files for reproduction and a detailed build up yielding the required functionality. A more generally applicable description of the highly-configurable microFPGA [40] has been provided elsewhere and may prove useful for particularly adventurous builders who want to further expand the microscope functionality or use microFPGA [40] for other projects. A full list of software controllable hardware elements used in the microscope is provided in Supplementary Table 4: Micro-Manager Hardware Configuration.

### Applications

The relative benefits of SMLM with regard to other super-resolution microscopies have been covered in detail elsewhere [41–44]. In general, applications requiring spatial resolution beyond that which can typically be achieved by stimulated emission depletion (STED) [45], structured illumination (SIM) [46] or image-scanning microscopy (ISM) [47] may be ideally suited to the use of 3D-SMLM.

The microscope presented herein has been used extensively across multiple generations of development for 3D super-resolution imaging of biological samples with an associated localization precision, σ typically down to 2/8 nm laterally/axially under ideal conditions (the associated resolution is typically 2.3σ [48]). Using oil or silicone oil objectives as well as TIRF/HILO/epi-illumination one may image structures close to the coverslip boundary or as far as 50 µm beyond. The microscope acquisition workflow is flexible enough to allow for multiple SMLM imaging modalities including (f)PALM [1,2], (d)STORM [3,4] and DNA-PAINT [5]. For example, in our group we used the automated high-throughput mode to determine the distribution of the majority of proteins during clathrin-mediated endocytosis [32] and the high 3D resolution allowed us to investigate the precise dynamics of membrane curvature generation [33]. Furthermore, we extended 3D imaging to cells and investigated the structure and stoichiometry of nuclear pore complexes [49] and the kinetochore [34].

The microscope has the flexibility to provide a field of view as large as 77 µm in diameter, or to optimize for increased laser intensities over a smaller field of view for increased speed, blinking contrast, and lower auto-fluorescence background [27]. Combined with the automated imaging capabilities, the instrument is thus equipped for high-throughput applications, akin to previous studies which resolved the structural organization of large numbers of endocytic proteins through such workflows [32].

### Comparisons with other methods and limitations

Regarding the specific 3D-SMLM implementation presented here, one must consider the associated limitations relative to other SMLM setups. As discussed previously, the microscope is functionally more capable than previous open-source SMLMs as defined by our experiences with SMLM. Others may have other preferences, for example, the maximum illumination field is a relatively modest 77 µm in line with comparable commercial systems (e.g., Nikon N-STORM, ONI NanoImager, Bruker Vutara, Zeiss Elyra), whereas several reports have described SMLMs, with larger fields [12,13,50]. We have constrained the system as such for several reasons. First, the illumination field stated approximately corresponds to the maximum practical field of view offered by high-NA oil and silicone oil lenses for which field aberrations are low. Since a spatially invariant point spread function (PSF) is critical for precise fitting of single molecules, this is a crucial factor in the use of SMLM for structural biology [16]. Secondly, a larger illumination field requires more laser power, which increases the background caused by autofluorescence in the objective and optics, and which can lead to damage of the expensive objective lenses, especially in TIRF illumination where the laser is focused onto a tiny spot at the periphery of the back focal plane and misalignments may result in damage to the objective back aperture. Thirdly, a smaller illumination field allows one to image two channels onto a single camera while maintaining spatial sampling requirements. For users wishing to optimize throughput, we again note that the most effective strategy is automation. In pursuit of performance, we have selected high-quality components that make the 3D-SMLM a relatively expensive build at the upper end of the budget ($160k). Furthermore, we have optimized the system for stability and ease of construction/use using a large number of custom parts, which have to be sourced, rather than relying on more generic commercial options. There are certain sub-systems, for example, the dual-channel image splitter, that can be assembled largely from commercial components if preferred, although the assembly and alignment process is less-straightforward. With regard to similar commercial systems, (e.g., Nikon N-STORM, ONI NanoImager, Bruker Vutara, Zeiss Elyra) control of the microscope may seem cumbersome and will require a greater degree of expertise to achieve the highest performance. However, this is a general point shared by most custom microscopes and the fine control of the device afforded by the control scheme presented will ultimately deliver superior results when employed correctly.

In addition to the benefits of the technique discussed, 3D-SMLM features unique limitations. Notably, the low temporal resolution arising from the need to acquire tens to hundreds of thousands of frames to restore single super-resolved images renders live-cell time-lapse imaging challenging [51], which may be better suited for e.g., SIM/ISM [46,47]. Nevertheless, in the implementation presented, automation of the microscope ensures that long acquisitions may proceed unsupervised to provide high-throughputs. An additional bottleneck to throughput can be the image reconstruction. However, with proper management and appropriate IT infrastructure the data volume/processing is manageable. We recommend on the fly analysis during data acquisition to help to identify problems before a more complete reconstruction is carried out in post-processing.

Although the resolution achievable with 3D-SMLM is far superior to diffraction-limited techniques as well as STED/SIM/ISM [45–47], more specialized fluorescence microscopies such as interferometric SMLM analogues employing dual opposed imaging pathways (e.g., 4pi-SMS) [52] or coordinate-targeted approaches such as MINFLUX/MINSTED [53,54] typically achieve superior resolution in one or all dimensions. However, a recent study based on Exchange-PAINT and 3D-SMLM achieved Ångström level resolution in fluorescence microscopy for the first time [55]. Setting aside this impressive demonstration, 3D-SMLM offers a simpler, less expensive and more robust route to obtaining super-resolution images than offered by MINFLUX/MINSTED/4pi-SMS.

## Overview of the Protocol

Full setup and validation of the microscope performance is an involved procedure encompassing multiple disciplines. The protocol is broken down into a number of discrete tasks, which should be carried out in the order specified and care must be taken to properly benchmark performance as indicated before moving on.

Prior to the protocol we include an Experimental Design section detailing first steps such as hardware selection and provision of an appropriate environment in which the system may be constructed as well as an Equipment Set Up section, which describes best practices for optical alignment amongst other useful guidelines.

The protocol, which details the construction of the system and its subsequent validation, is subdivided as follows.

First, the microscope body, comprising more than half of the custom parts of the microscope is built up (steps 1 - 67). Before any optical alignment can be performed, it is advisable that the various hardware elements can be controlled and configured from a single interface. To this end, the microFPGA, which allows for monitoring and analog/digital control of hardware, is constructed (unless specifically sourced in an assembled state, see Electronics rationale and sourcing) (steps 68 - 103) and configured (steps 104 - 107). Next, all software for acquisition is set up and configured, including µManager and the EMU htSMLM plugin, which together provide the user interfaces for the microscope (steps 108 - 130). Having installed µManager, it is possible to fully test the microFPGA functionality (for self-assembled microFPGAs only) (steps 131 - 137). Electrical connections between many of the hardware elements and the microFPGA are then made (steps 138 - 142). Subsequently, a camera and transmission target are set up to provide an infinite conjugate reference, the use of which allows us to correctly focus the emission optics (steps 143 - 152). The microscope alignment then commences with routing of an alignment laser through the emission path (steps 153 - 174) and aligning the various lenses to produce a pair of secondary images on the camera corresponding to the two pathways of an image splitter (steps 175 - **Error! Reference source not found.**). Following from the emission path, the illumination path is then set up for simple epi-fluorescence imaging with a spatially inhomogeneous Gaussian illumination field (steps 230 - 258). The microscope is then focused and initial checks of PSF quality are performed (steps 260 - 272). The illumination optics are subsequently augmented by a field variable, beam homogenization unit (steps 273 - 313) as well as optional steps of setting up of a booster laser (steps 314 - 335), configuring for TIRF/HILO illumination (steps 336 - 344), installation of a multi-mode laser engine (steps 345 - 373), and calibration of the photodiode used to monitor the laser power (steps 374 - 379). Subsequently, a series of checks and calibrations relating to the 2D imaging performance are carried out to prevent crosstalk between the image splitter channels, determine the object space pixel size and assess the field-dependence of the PSF under homogeneous illumination (steps 380 - 392).

Having achieved a final illumination configuration, the emission path is extended for 3D functionality via the installation and adjustment of a variable astigmatic lens (steps 393 - 414). The following tasks relate to the focus lock system, including the alignment of the focus lock laser into and out of the body via (total internal) reflection at the coverslip the body (steps 415 - 433), setup of the focus lock electronics (steps 434 - 445), focus locking the microscope (steps 446 - 449) and establishing the stable operating regime of the focus lock laser (steps 450 - 458). The microscope is fully functional at this stage and further steps require the installation of the SMLM data analysis software SMAP (steps 460 - 465) for analysis of the data. As a first step towards 3D imaging a PSF calibration is performed using a fluorescent bead sample and the 3D imaging performance of the system is adjusted (steps 466- 473). Having completed the PSF calibration, the next steps detail the use of SMAP for 2D and 3D calibration based on a Gaussian or experimentally-derived model of the PSF respectively (steps 474 - 487). The following section considers the rendering of the localized data including the merging, filtering and drift-correction of the individual localizations (steps 488 - 494). At this juncture the necessary tools and knowledge have been established to make additional quality checks including checks of the focus lock systems’ ability to correct axial drift, as well as assessment of lateral drift and vibrations (steps 495 - 517). Subsequent performance benchmarking of the microscope for biological imaging includes 2D (steps 518 - 550) and 3D single-color and 3D multi-color imaging (steps 551 - 562) of nuclear pore complexes.

## Experimental Design

### Personnel

For successful completion of the protocol including planning, procurement of commercial and custom parts, construction/alignment and validation, one full-time scientist or engineer is required. Preferably this person would have some familiarity with optical systems although such experience is not strictly required. Alternatively, it may be desirable to split certain tasks over more specialized or suitable individuals. For example, samples may be prepared by life-science technicians. From the point where all required hardware is available, we assert that the protocol may be completed in 3 - 6 months.

### Laboratory environment

For best results and ease of implementation, the selection of a microscopy-dedicated laboratory environment is preferable. We note, however, that much of the development of this microscope has taken place in general laboratory space with excellent results. This distinguishes the 3D-SMLM system reported from other super-resolution microscopes, which may suffer more from thermal drift or vibrations such as 4pi SMLM [52] analogues, which require perfect co-alignment of two opposed imaging paths and STED, which requires perfect co-alignment of a toroidal depletion beam with the excitation beam [45]. Nevertheless, the ideal environment is temperature controlled (to within ± 1 °C) to obviate thermal drift, which under hysteresis results in the slow misalignment of optical components. Since the microscope may exhibit substantial thermal inertia, the desired operating temperature should be maintained over evenings, weekends and other periods of closure. Moreover, the laboratory itself should not have too strong vibrations (better than 10 µm/s RMS velocity, vibration criteria category: VC-C/VC-D). Ideally the lab would be located on the ground or basement level of buildings and to avoid coupling of vibrations through plastic-lined laboratory floors, the optical table would be supported directly on the concrete floor below with cut-outs in the plastic lining (if present) made accordingly. For more remote locations, higher building levels may also be acceptable. Similarly, sources of vibration native to the building itself should be avoided and the laboratory should be situated away from compressors or circulation systems (for example, those relating to cooling/cryogenics or ventilation) as well as lifts/elevators and centrifuges. Nevertheless, even in relatively poorly suited surroundings, the optical table does efficiently isolate the microscope. Sources of dust should be avoided as much as possible. For example, air conditioning systems should provide filtered air and, if possible, a slightly higher pressure than surrounding rooms to prevent ingress of dust. Dust generation in the laboratory can be further minimized by regular cleaning and using replaceable adhesive floor mats at the entrances to remove soiling from footwear. Ultimately some dust is unavoidable but its impact can be minimized by enclosing the optical table and microscope itself.

The lab space itself should be large enough to accommodate a 1.5 × 0.9 m optical table, fitted with an appropriate enclosure for laser safety and stability, as well as a desk for the instrument control PC while maintaining access to three or more sides of the optical table, ideally including the two long sides. We note a suitable optical table in the parts list. To achieve optimal results similar to those presented in Figure 8 and Figure 9, one should aim to enclose the optical table itself and provide additional sub-divided enclosures on the table itself around the various optical modules to obviate drift and prevent specular/diffuse reflections of the lasers escaping the optical table or being captured by the camera. The design of each enclosure will depend on the optical table, placement of electronics and other factors that cannot be predicted *a priori* and so is not included in the protocol.

For builders wishing to use a pre-existing optical table, it should be at least 1.5 × 0.9 m with a 25 mm pitch, M6 tapped hole pattern and not so much larger that access for alignment of the optical pathways is impaired. Imperial configured tables with a 1” pitch, ¼”-20 tapped hole pattern cannot be used without redesign of many components.

We note that in tight spaces with only weak air conditioning systems, the additional thermal burden of lasers, controllers, and PCs may result in a greater fluctuation of temperature during operation. In any case, we would recommend powering the instrument several hours before its intended usage to allow rooms to equilibrate as best as possible. The optical table specified in the parts list uses an integrated design whereby the four legs are attached as a single frame. Alternatively, one may use individual floated legs and an appropriate table top at least 100 mm thick if such a setup is available. Two optimal layouts of the microscope in typical/small rooms or space of 2.25 × 3.45 and 2.35 × 2.90 m are shown in Supplementary Figures 4 and 5. Ideally the desk for the PC(s) should be positioned such that one can pivot on a lab chair from the desk to the body of the microscope where the sample is placed.

Ideally, all sources of heat and vibration should be located off the optical table. In practice this may not always be possible but one should seek to locate all electronics off the optical table. Necessary exceptions are free-space booster lasers and the camera. The specified sCMOS camera is fan-cooled as standard. However, operation of the fan can result in noticeable vibrations in images. As such, the fan-cooling should be switched off and water cooling should be provided. The cooling unit should not be located on the optical table. Instructions for doing so are included in the protocol. This process may differ for other cameras. Regarding the lasers, fiber-coupled options can be readily accommodated off the optical table and builders should seek this approach where possible. The small fiber-coupled focus lock laser can be located on the optical table if desired and a part file is provided to mount it as such (FAB-EMBL-000088). The multi-mode laser engine, particularly the agitation unit used for mode-scrambling in the fiber, should not be located on the optical table.

All local guidelines regarding laser safety should be followed. Note, during system alignment where the path of the beam may be ill-defined additional care is warranted. Laser-safety glasses should be used and minimal laser power/appropriate beam-blocking as needed. A particular hazard is presented by collimated lasers emerging from high NA objective lenses. A high NA objective can launch a laser over a solid angle approaching 2π, with only minor steering of the beam on the infinity-side resulting in large swings in angle. Although the beam waist will be small and thus will expand noticeably away from the objective lens due to diffraction, the laser(s) can still cause permanent damage to eyes. Take particular care whenever aligning the laser to the objective lens or when installing the objective lens to project a realigned beam. Block the beam passage as a first step when unsure of the beam pointing out of the objective and follow the beam as it propagates. The instructions provided are designed to ensure that highly inclined beams are not launched directly off the near side of the optical table but nevertheless, the responsibility for laser safety ultimately resides with the builder. We have operated the microscope with a security relay to close the laser interlocks only when the microscope cover is in place. The system features an override mechanism to allow uninterrupted use only while aligning the system. Those wishing to implement a similar approach are encouraged to contact the EMBL electronic workshop for further information.

### Mechanical parts rationale and sourcing

The microscope detailed requires many tens of custom mechanical parts. Where possible we have used commercially available off-the-shelf (COTS) components given that requirements for i) stability, ii) ease of assembly and iii) complexity are met. For example, while it is possible to construct an inverted body fully from COTS parts, stability and functionality may be lacking and the assembly may be needlessly complex. We provide toleranced drawings for the custom parts for fabrication at a suitable facility. Note, where tolerances are not specifically defined by the ISO standard, we assume a relatively coarse tolerance of ± 0.05 mm (or ± 2 thousands of an inch for workshops more familiar with such conventions) on mechanical dimensions. Facilities with well-operated, maintained and calibrated computer-numerically controlled (CNC) mills are generally required for timely production of many of the parts and should be capable of holding tolerances at or approaching 5× better than this coarse measure. Particularly for dowel-pinned components, holding such tight tolerances is required to minimize the effects of tolerance stacking. We use dowel pins throughout for alignment as well as pocketed features to define the position of several components. The need to remove parts and install temporary alignment fixtures means that holes for dowel pins should be machined for a slip/transition fit. We also note that tightly toleranced dowel pins of the recommended specification should be used throughout. When a press/interference fit is required, this is indicated from the toleranced drawing. In such cases, for example for the idler or bearings for the belt-drive objective lens translation mechanism, ideally the associated commercial part should be provided for the workshop to judge tolerances. We urge caution in installing these parts as incorrect placement may be difficult to correct once dowel pins or bearings have been pressed in place. Many workshops will work with an external facility to anodize machined components, which is strongly recommended owing to laser safety and general back reflection concerns. Moreover, dowel-pin holes have been specified assuming subsequent anodizing to avoid under-sized holes. Nevertheless, a supply of minimally under-sized dowel-pins is a wise precaution. Many parts require tapping to produce an internal thread for the mount of optical components (e.g., RMS thread for the objective lens), the thread specification for these threads should be available from the manufacturer, the threads are called-out in the corresponding feature in the CAD file and bore sizes are sized at 83.3% thread.

Since it is undesirable to leave expensive optical components with mechanical workshops, commonly available optical component thread adapters provide a budget-friendly option to allow the facility to test fits. Additionally, very fine-pitch threads (< 0.25 mm), required in one case, may prove difficult for some workshops to machine successfully. For this reason, we provide an alternative option based on a commercial fine pitch actuator and an additional, more-easily fabricated component with a machined hex-key interface. In addition to the machined components, two of the part files are supplied for 3D printing (in .sldprt and .stl format). These parts are included for completeness to mount certain boards and do not warrant high-precision CNC machining. One of these parts (FAB- EMBL-000095) includes bores for heat-installed threaded inserts, which can be installed using a soldering iron on a low setting.

### Electronics rationale and sourcing

The microscope makes use of the microFPGA control platform to trigger lasers and the camera, modulate laser diodes associated with a low-cost laser engine, and monitor signals from power and position monitoring photodiodes. The generic/multi-purpose microFPGA design, functionality and assembly has been described by Deschamps et al. [40]. Here and in the protocol, we refer to a specific implementation of the microFPGA that is suited to the needs of the microscope while providing some additional functionality. For example, additional transistor-transistor logic (TTL) or pulse-width modulation (PWM) signals can be provided to drive e.g., flipper mirrors and servo motors. This added functionality is included in the base configuration described, as, in our collective experience, it is typical that builders will want to control additional hardware later and the various signal conversion boards included as standard, already provide the auxiliary functionality.

At the core of the microFPGA is a low-cost hobby-grade FPGA (Alchitry Au+). In addition, various commercially available and custom boards are connected to the FPGA. These additional boards act to protect the FPGA, ensure proper connection with labelled SMB connectors, scale I/O voltages appropriately, and low-pass filter PWM signals to produce analog outputs. The data required for reproduction of all custom boards is included (see: https://github.com/ries-lab/3DSMLM), with original board files (in Altium), the Gerber files, bill of materials, schematics, and board files as .pdf. A list of all boards and parts required for the microFPGA is given in the parts list. Depending on the builder’s expertise in electronics we suggest the following options. Firstly, a pre-assembled microFPGA can be sourced directly. Those wishing to make use of this option should contact the technology transfer department of the European Molecular Biology Laboratory (EMBL/EMBLEM). For those with access to a capable electronics workshop it may be preferable to purchase all commercial and custom boards and self-assemble and test the microFPGA. There are few specific requirements on PCB manufacturers used and the builder is advised to select on the basis of professional recommendation, price and general rapport. A full guide to assemble the microFPGA from the complete electronics boards is included in the protocol. We note that there is no need for the builder to interact with or compile the microFPGA code, which is simply flashed to the device via the manufacturer supplied software.

In addition to the microFPGA the microscope employs a custom white-light ring LED with associated controller as well as an amplifier for the QPD used in the focus lock system. The former is not strictly required and can be omitted (if transmission imaging functionality is not required) or replaced by a suitable triggerable/directly modulatable (not pulse-width modulated) ring LED source. Doing so will require some modification of the CAD files to mount the device. Again, full data required for reproduction of the custom boards including the board-mounted LED ring is included (see https://github.com/ries-lab/3DSMLM). Conversely, the focus lock electronics are required and comprise a commercially available amplifier board (see parts list), which simply requires appropriate packaging including provision of appropriate power and making connections to various input/outputs, which can be handled by the builder or electronics workshop as necessary.

### General components sourcing

Under typical procurement conditions, 4 months is sufficient to acquire the necessary materials. Where certain components are not immediately available or facing undue delays, it is possible to reorganize some of the protocol or otherwise to begin sections before completing earlier ones. Achieving the highest level of performance is contingent on the correct function of various components, none more so than the objective lens. We note that all new/pristine standard oil objectives listed that we have received to us delivered the expected performance. Testing lenses from demo stocks may be less informative than one would expect as the use and misuse of these examples often renders their performance sub-par. Compared with the standard oil lenses, the analogous silicone oil objective lens, however, exhibit far stronger field-dependent aberrations (primarily coma), which make precise fitting of experimentally-determined average PSFs difficult. The builder is urged in this case to directly purchase the objective lens tested if the performance is determined to be sufficient over the required field of view and limit the field of view of the microscope accordingly. For example, it is not recommended to use illumination fields larger than ca. Ill30 µm with the silicone objective lens (limiting the recommended telescopic magnification to 1×, see iii Emission path). Moreover, for this reason, it is recommended to purchase an oil objective alongside the silicone objective to make setting up the microscope easier.

### Computational infrastructure

Handling the data stream from the sCMOS camera is the primary consideration for the computer. The typical maximum data rate of the instrument is > 200 MB/s (See Supplementary Note 7: Camera data rate) and more commonly it is substantially less than this. Additionally, one must consider storage capacity. For the largest imaging field and a particularly long acquisition of 500,000 camera frames, a single field of view may occupy up to 1 TB. For a standard field of view, and a more modest acquisition of 200,000 frames, ca. 100 GB is required. In either case, one may quickly exceed the limits of local storage and should consider networked solutions. To achieve the required data rate and storage volume, we stream the data directly to networked storage preferably via a 10 Gbps (1.25 GB/s) fiber connection. Other local solutions include single solid-state drives (SSDs). To extend the capacity of a number of inexpensive SSDs one may use them in a RAID0 configuration. Similarly, one may achieve the required data rate with several mechanical hard disk drives (HDDs) in RAID0 and in this way provide storage capacity in the tens of TB. The builder is advised to contact local IT infrastructure management to better understand the options available locally. During initial setup of the instrument, it is typically sufficient to buffer images in RAM before they are more slowly written to disk or to use a single modest-capacity SSD for the minimal test data required.

Producing super-resolved images from the 3D-SMLM data is contingent on fitting of the single emitters and rendering the molecule positions *in silico*. Without appropriate consideration of this aspect and its hardware underpinnings, fitting may ultimately limit throughput more than the microscope itself. Considering the analysis machine, here one may specify according to the available budget. However, since the fitting of emitters is performed using CUDA, a recent Nvidia GPU provides a minimum requirement. Excellent results (several thousand localizations per second) can be achieved with a relatively inexpensive consumer grade GPU. See the Equipment section for details. Note that a separate computer for analysis is not a requirement but can provide benefit depending on the setting. In research-based settings where the microscope and acquisition PC is hosted and used by a single lab, we have predominantly performed single-molecule fitting on the microscope acquisition PC. Whereas in a service setting we have preferred to separate the task between two machines to allow users to perform data analysis at a later date without interrupting other users acquiring data.

### Data analysis and rendering

During data acquisition the raw camera frames and metadata describing the state of the microscope are saved by µManager. This data requires extensive analysis steps to yield biological insights: fitting of single molecule positions, post-processing such as filtering and drift correction, rendering as a super-resolution image and quantitative evaluation. We developed the super-resolution microscopy software platform SMAP [28] and here detail the analysis steps using this solution, but there are numerous other software, notably Picasso [56] and ThunderStorm [57] that can be equally used. For an overview of the performance of different localization algorithms, we recommend users study the results of the SMLM software challenge [58].

To analyze 3D or multi-color data in SMAP, first the PSF of the microscope and/or channel transformation are calibrated from image stacks of fluorescent beads. Then SMAP identifies possible single-molecule events in the raw camera frames and determines their position with sub-pixel precision by maximum likelihood estimation. Localizations with low accuracy can be filtered out, before residual drift is corrected and a super-resolution image is rendered. These steps can be performed on the fly during data acquisition. SMAP provides numerous plugins to segment and quantitatively analyze cellular structures based on the coordinates of the detected fluorophores.

## Materials

### Reagents

CAUTION: All reagents present potential hazards and should be handled correctly by personnel trained in general laboratory safety and materials handling. Specific hazards are noted below with the respective reagent.

- β−mercaptoethanol (Sigma-Aldrich, cat. no. M6250). CAUTION: Handle with care. β - mercaptoethanol is a toxic, corrosive, and strong irritant that poses risks to human health upon contact, inhalation, or ingestion, potentially causing skin, eye, and respiratory irritation, as well as sensitization and allergic reactions.
- SNAP-Surface BG-AF647 (New England Biolabs, cat. no. S9136S)
- Bovine Serum Albumin (BSA, Sigma-Aldrich, cat. No. A7030)
- Catalase (Sigma-Aldrich, cat. No. C3155)
- Isopropanol (Merck, cat. no. 67-63-0). CAUTION: Handle with care. Isopropanol is highly flammable liquid and vapor, causes serious eye irritation, and may cause drowsiness or dizziness.
- Petrol ether (Carl Roth GmbH, cat. no. 3295). CAUTION: Handle with care. Highly flammable liquid and vapor, may be fatal if swallowed and enters airways, causes skin irritation, may cause drowsiness or dizziness and is toxic to aquatic life with long lasting effects.
- Coverslips 12 × 12 mm (Marienfeld Superior, cat. no. 0101000)
- Coverslips No. 1.5H Ill24mm (Marienfeld Superior, cat. no. 0117640)
- DMEM (Gibco, cat. no. 11880-02)
- Dithiothreitol (DTT, Biomol, cat. no. 04010.10). CAUTION: DTT is a reducing agent that can cause eye and skin irritation upon contact, and may lead to respiratory irritation or allergic reactions when inhaled or ingested in significant amounts.
- Fetal Bovine Serum (FBS, Gibco, cat. no. 10270-106)
- 2-Deoxy-D-glucose (Sigma-Aldrich, cat. no. D3179)
- Glucose Oxidase (Sigma-Aldrich, cat. no. 49180)
- GlutaMAX (Gibco, cat. no. 35050-038)
- Glycerol (Merck, cat. no. 1.04094)
- Hydrochloric acid, HCl (Sigma-Aldrich, cat. no. 7647-01-0)
- Immersion oil (Olympus, cat. No. IMMOIL-F30CC)
- Silicone immersion oil (Olympus, cat. No. SIL300CS-30CC)
- MEM NEAA (Gibco, cat. no. 11140-035)
- Methanol (Sigma-Aldrich, cat. no. 67-56-1)
- Millex-GP filters, 0.22 um (Merck, cat. no. SLGP033RS)
- Sodium chloride (NaCl, Merck, cat. no. 7647-14-5)
- Ammonium chloride (NH_4_Cl, Merck, cat. no. 12125-02-9). CAUTION: Handle with care. Ammonium chloride may cause skin, eye, and respiratory irritation upon contact, inhalation, or ingestion, with potential for more severe effects following prolonged exposure.
- Parafilm (Thermo Fisher, cat. no. HS234526B)
- PBS, 1 (waiting for specifications from the media kitchen)
- PBS, 2x (waiting for specifications from the media kitchen)
- Paraformaldehyde, 16% (PFA, Electron Microscopy Sciences, cat. no. 15710). CAUTION: Handle with care. PFA is flammable, harmful if swallowed or inhaled, causes skin irritation, may cause an allergic skin reaction, causes serious eye damage, may cause respiratory irritation, is suspected of causing genetic defects if exposed, and may cause cancer if exposed.
- TetraSpeck fluorescent beads (100 nm) (Thermo Fisher, cat. no. T7279)
- Tris-HCl, pH8, 1M (waiting for specifications from the media kitchen)
- Triton X-100 (Sigma Aldrich, cat. no. X100). CAUTION: Handle with care. Triton X-100 is harmful if swallowed, can cause skin irritation and serious eye damage. Very toxic to aquatic life with long lasting effects.
- TrypLe (Gibco, cat. no. 12604013)
- Anhydrous, deoxygenated DMSO (Sigma Aldrich, cat. no. 900645-4X2ML)
- WGA-CF680 (Biotium, cat. no. 29029-1)

### Equipment

A full list of parts shared between all configurations is provided in Supplementary Table 2.

#### Optical and mechanical components

The microscope uses many commercially available off-the-shelf (COTS) optical components including spherical, cylindrical and widefield tube lenses, dielectric mirrors, dichroic and polarizing beam splitters, spectral filters and optical fibers. In addition, all configurations require an objective lens either of oil- or silicone oil-immersion type and a refractive beam homogenizer is required for certain configurations. Independent of the final configuration, the microscope is composed of > 90 custom mechanical parts, which need to be sourced from an appropriate workshop (see: Mechanical parts rationale and sourcing). In addition, > 350 COTS optical and optomechanical components are required as well as additional electronic and mechanical components. The parts list is correct at the time of publication, however, improvements and additions to the design are ongoing. An up-to-date parts list can be downloaded from: https://github.com/ries-lab/3DSMLM, where the associated CAD files are also hosted.

#### Electronic components

The microscope uses many custom and commercially available electronic components. The custom electronics can be sourced as discussed in Electronics rationale and sourcing. Transmission illumination is provided by an LED ring source, which directly mounts to the microscope cover. A QPD and associated amplifier are used for focus locking and an optional single-element photodiode (with electronics integrated into the FPGA and signal conversion unit) can be used for recording of experimental laser power. At the heart of the microscope control scheme lies a low-cost hobby-grade field-programmable gate array (FPGA, Alchitry Au) and a number of signal conversion boards (see https://github.com/mufpga/MicroFPGA [40]) to provide synchronization of hardware, readout of the QPD, analog/digital control of the single/multi-mode laser engine(s), and more generally to provide the builder with the flexibility to add and control additional hardware.

#### Motion control/Mechatronics

Several motion control/mechatronic systems are used in the microscope and together provide a substantial portion of the budget. This includes an objective lens piezo flexure stage to focus and focus lock the objective lens as well as the long travel XY stage for sample positioning. Additional piezo resonant stages are required for positioning the laser incidence at the objective back focal plane to incline the illumination suitably for epi-/HILO/TIRF illumination modes and to select between 2D/3D modes by positioning an astigmatic lens in the emission path. Furthermore, several motorized filter wheels and piezo-sliders are required to select emission filters and introduce a Bertrand lens to the emission path to view the objective back focal plane on a camera.

#### Laser sources

The microscope uses lasers for excitation of various fluorophores as well as for on-switching in (d)STORM/PALM and to provide a positional reference for the focus lock system. The microscope can be configured for different requirements and budgets through the provision of various laser sources. For lower budgets and where TIRF illumination is not required, a low cost multi-mode laser engine can be constructed according to a previously published scheme, which provides a well-homogenized illumination source [35]. This is a particularly budget-friendly option where only diode lines (typically 405, 488, 640 nm) are required as the range 520 - 620 nm (typically 561 nm) requires an additional commercially packaged laser head. Where TIRF illumination is desired, the base-configuration includes a single mode fiber-coupled laser engine. Note, it is possible to combine several illumination options as desired or add additional ones as the need arises.

Typically, the single-mode laser engine is the largest budgetary portion of the microscope, if included. While the standard configuration includes excitation/activation sources at 405, 488, 561, 640 nm, the microscope can in principle be adapted to other applications by specifying alternative laser lines that are similarly commercially available at time of purchase. However, the stated wavelengths are the most commonly used in SMLM and biological imaging more broadly. For power hungry applications an additional free-space single-mode laser can be provided at a single wavelength, most commonly at 640 nm owing to the tendency to perform dSTORM imaging (with far-red dyes) at high laser power. As default configuration from the manufacturer, the recommended booster lasers do not allow for digital modulation. Ensure to specify this option when sourcing these components.

The single-mode lasers are homogenized and expanded to produce a uniform and variably sized illumination field at the object. It is possible to adapt the scheme provided to omit the variable magnification or homogenization units (or both), opting instead for a fixed size illumination field and lossy homogenization scheme based on use of a field aperture to heavily apodize the Gaussian beam. This configuration is not specifically supported but it is comparatively simple to implement with some considerations of the necessary beam expansion. Those wishing to develop such a scheme are advised to contact us to discuss any possible issues that may arise. Additionally, builders may use other single-mode lasers or engines than those specified, for example, where hardware already exists in the lab (see Supplementary Note 4: Using non-specified single-mode lasers and laser-engines). However, we cannot comment on the availability of µManager device adapters for unsupported hardware.

The focus locking system uses a near infrared 808 nm fiber-coupled single-mode diode laser to be spectrally separable from the excitation/activation sources and emitted fluorescence, which even for far-red dyes is negligible beyond ca. 750 nm.

#### Imaging camera

A single camera is used to capture fluorescence images, even when two spectral channels are employed. We have used several cameras successfully across the many generations of the microscope. The base configuration specifies a Hamamatsu Fusion BT sCMOS camera, which represents a substantial fraction of the total budget. See Supplementary Note 7: Camera data rate for a discussion of camera data rate to determine whether it is necessary to purchase the CoaXPress frame grabber card for the camera. Typically for the FOVs defined by the illumination system and typical exposure times, this is not necessary but for those wishing to substantially adapt the design for larger FOV imaging, the card may be necessary. The Hamamatsu Fusion BT sCMOS camera has three imaging modes, defined here as slow, normal and fast mode. Each has their benefits for imaging, with the slow mode having just 0.7 electron read noise being particularly well-suited to slowSTORM [27]. Unless otherwise noted, use the normal mode for general imaging e.g., of fluorescent bead and dye samples and for various calibration/alignment procedures. For the dSTORM imaging presented as a route to validation of the microscope, the slow mode is optimal, but the normal mode can be used e.g., for larger field of views where the limited frame rate of the slow mode makes imaging less practical.

Lower-cost generation II sCMOS cameras such as the Hamamatsu Orca Flash 4.0 v2 have also been used and are also well-suited as the imaging camera, albeit with higher noise and lower quantum efficiency. Similar cameras from other manufacturers for example Andor Zyla 4.2, PCO Edge 4.2 may be equally performant but prospective builders are recommended to thoroughly test cameras and their µManager device adapters before committing to purchases. Although we have operated 3D- SMLM systems with electron-multiplying charge-coupled device (EMCCD) cameras (Photometrics Evolve 512, Andor iXon), they require substantial adaptation of the emission path to increase the magnification to accommodate the larger pixel size.

Owing to their readout architecture, sCMOS cameras are characterized by having pixel to pixel variability in noise. For older generation II sCMOS chips and particularly for uncooled industrial CMOS cameras, the variation can be large enough to require calibration of the noise characteristics such that it may be accounted for in localizing fluorophores. Builders wishing to use generation II sCMOS cameras should follow published protocols for generation of the dark offset and variance maps required [30,59] and the SMAP user guide detailing how to include these in final localization analyses. For the generation III Hamamatsu Fusion BT, the read noise is low enough and design of the camera is such that the fraction of pixels with abnormal noise characteristics is vanishingly small and therefore calibration of the noise characteristics is not required.

#### Microscope control computer

The microscope control computer can be configured as desired with a few points worth noting. If the same machine is to be used for analysis tasks, one should ensure that both sets of requirements are met. Smooth performance can be achieved by choosing a central processing unit (CPU) with at least 4 cores, for 8 threads in the tested 64-bit Windows 10/11 environment. Lesser configurations may result in µManager 2.0 struggling to achieve a suitable image refresh rate (>10 Hz) during acquisitions, which is crucial to be able to assess conditions for blinking. We strongly recommend installing the operating system on an SSD and to reserve this disk for the operating system and required software and drivers.

Other than these concerns, refer to the Computational infrastructure section for a discussion of data transfer and storage. Depending on the specific configuration chosen, 1 - 4 PCI express (PCIe) ports are required. For example, high-speed SSDs, camera frame grabbers, network cards and GPUs may all occupy this interface. Consult the manufacturer of these components for specification of the port. For example, frame-grabber cards often specify PCI express x8 or x16. One PCIe port is also recommended for a modest graphics processing unit (GPU). Depending on the model chosen, the GPU may occupy more back panel space and block other PCIe ports. There are many USB and RS232 serial devices to connect. As such, it is recommended to use high-quality mountable USB and serial hubs which may be situated closer to the hardware elements and allow a greater number of communications channels than the back/front panel connectors of the acquisition computer. At the time of publication, the following computer was used for all microscope control and data acquisition tasks: Dell Precision Tower 5820, Windows 10 Professional 64-bit, Intel Xeon W-2102 Processor, 32 GB random-access memory (RAM), 512 GB SSD, 2 GB HDD, Nvidia Quadro P1000. The following guidelines provide a suggested configuration:

- Operating system: Windows 10 64-bit
- CPU: 4-core CPU
- GPU: 4GB VRAM
- RAM: ≥16 GB
- Operating system SSD: ≥ 256 GB*
- Storage disk(s): either SSD (or multiple SSDs in RAID0): ≥ 1 TB
- Storage disk(s): or 1-10 Gbps fiber network, corresponding network card and network data storage with sufficient write-speed
- sCMOS camera interface: 1x USB 3.0 port

*Note a single SSD may be used for the operating system and data storage as desired.

#### Image analysis computer

The image analysis computer may be used for analysis as well, assuming both sets of requirements are met. For on-the-fly data analysis, which is helpful for quality checking and optimization of experimental parameters, this is often the preferred route as the acquired data are inherently available. We have often adopted this approach. We have also split the acquisition and analysis tasks over two machines, each connected to a file server where data are stored via a 10 Gbps network connection. SMAP uses GPU acceleration for parallelized fitting of experimentally-derived PSFs based on CUDA (Nvidia), as such a CUDA-compatible GPU is required. Naturally, performance will scale with hardware availability. Note, if using a single machine for analysis and acquisition, avoid connecting the monitor to the GPU used for analysis as this may interfere with the monitor refresh. In this case, favor the motherboard-integrated GPU for connection of the monitor. At the time of publication, the following computer was used for image analysis: Dell Precision Tower 3650, Windows 10 Professional 64-bit, Intel Core i7-11700 Processor, 32 GB random-access memory (RAM), 1 TB SSD, Nvidia GeForce GTX 1650 GPU. The following guidelines provide a suggested configuration.

- Operating system: Windows 10 64-bit
- CPU: 4-core CPU
- GPU: Nvidia CUDA-compatible GPU, minimum Compute Capability 6.1, ≥ 4 GB VRAM, e.g., GTX 3070
- RAM: ≥ 16 GB
- Operating system SSD: ≥ 256 GB
- Storage disk(s): either HDD or SSD: ≥ 1 TB
- Storage disk(s): 1 - 10 Gbps fiber network, corresponding network card and access to network data storage (Only required when using separate PCs for acquisition/analysis)

#### Software for hardware setup

In order to test and configure the various hardware elements prior to full integration with µManager, it is necessary to install the following software that is either provided with the hardware or freely available from the manufacturer.

- Hamamatsu DCAM API
- Hamamatsu HCImage Live
- Toptica, TOPAS for iBeam smart laser(s)
- Toptica, TOPAS for iChrome MLE laser engine
- Thorlabs, Elliptec, ELLO software
- Thorlabs, FWxC filter wheel software
- Physik Instrumente, PIMikroMove
- Smaract, SCUConfigure, SCUFirmwareUploader
- Beam profiling camera software (e.g., Coherent BeamView)
- Infinite conjugate viewing camera software (e.g., FLIR Spinnaker SDK/SpinView)
- Alchitry, AlchitryLabs

#### Software for microscope control

The following software is required to control the microscope. Additional device adapters and a graphical user interface will be added to the base µManager installation during the protocol.

- µManager 2.0 latest official release
- ImageJ (installed as part of µManager)

#### Software for image analysis

The Super-resolution microscopy analysis platform (SMAP) [28] provides the basis for super-resolution image reconstruction in the protocol. SMAP may be installed with Matlab offering greater extensibility or as a standalone version requiring only the Matlab runtime engine, which is freely available. Please consult the SMAP documentation for installation notes and for tutorials to get accustomed with the data analysis workflow:

- SMAP latest official release for Matlab and Matlab 2022a or newer with toolboxes: Optimization, Image processing, Curve fitting, Statistics and Machine Learning

Or

- SMAP standalone version, latest release (installs the Matlab runtime).
- µManager 1.4 latest official release. This is a legacy version µManager required to load images into SMAP.
- ImageJ (installed as part of µManager)
- Nvidia graphics drivers
- Nvidia CUDA toolkit and graphics driver, version 10.1 or newer (for GeForce products 471.41 or later)

### Tools

Completion of the protocol requires a number of tools including screwdrivers, balldrivers/hex keys (L-shaped). These are assumed to be available and not included in the parts list. In addition, several tools that are less commonplace are included in the parts list. Furthermore, there are several assembled tools including alignment jigs/fixtures/optical references that are included in the parts list and as individual CAD assemblies.

#### Oscilloscope/lab power supply

An oscilloscope is helpful for checking triggering signals from the microFPGA and camera and is critical to correct setup of the focus lock electronics. Many options are available in this regard and even an inexpensive entry level model is sufficient to probe the signals required. If the microFPGA has been directly sourced in an assembled state, then the required testing has been performed. However, if the builder is assembling boards that have been printed/assembled at a third-party manufacturing facility, then a variable lab power supply is also needed. Many options are available for oscilloscopes and lab power supplies with one option for each suggested in the parts list (Supplementary Table 2).

#### Laser power meter

A laser power meter covering the spectral region 405 - 808 nm is necessary to check the correct functioning of the lasers, to determine the intensity at the sample, to aid alignment of spatial filters and fibers, and to calibrate the photodiode used for experimental laser power measurement. A digital interface is preferable for easy logging of laser power. Many options are available in this regard, with one option suggested for ease of coupling to the microscope body in the parts list (Supplementary Table 2).

#### Laser beam profiling camera

A laser beam profiling camera covering the spectral region 405 - 808 nm is necessary to aid the alignment of the refractive beam shaper if applicable and to check for mode hopping of the NIR diode laser used for the focus lock system. For the former, camera software with on-the-fly calculation of various beam dimensions is preferred since the refractive beam shaper requires a tightly specified input beam diameter for optimum performance. An open source project using µManager/ImageJ is available in this regard, requiring only a compatible camera [60]. However, for truly quantitative results one must consider variations in pixel response if, for example, a low-cost industrial CMOS camera is used (with the addition of a high neutral density blocking filter to obviate overexposure of the sensor). Commercially available laser profiling cameras include the necessary corrections to meet the relevant industrial ISO standards for definition of various beam parameters and software to allow readout thereof. Many options are available in this regard, with one option suggested in the parts list (Supplementary Table 2).

#### Alignment laser

An alignment laser and associated optomechanics are needed for alignment of the emission path. The emission path must be aligned with dichroic beamsplitters in place since they impart a shift to the system optical axis upon transmission. For a 3 mm dichroic in air, the shift is approximately 1 mm for 45-degree incidence, assuming a refractive index of 1.5. Although even µW power levels are visible to the naked eye depending on wavelength, adjusting lens tilt by viewing the inherently dim back reflection requires a moderate power alignment laser (>1 mW) which is passed efficiently by the primary dichroic mirror. Since the primary dichroic will typically be selected for reflection at typical 3D-SMLM laser wavelengths: 405, 488, 561, 640 nm, it is advisable to avoid these wavelengths and a narrow window around them of ±10 nm. The most suitable commonly available wavelength, which features the additional benefit of being ideally suited to visual inspection, is 532 nm. We note a 532 nm laser system in the parts list. Note that a 532 nm laser will be reflected by the image splitter dichroic (assuming a 665 nm long-pass dichroic, see Supplementary Note 2). To aid alignment of the image splitter system the tilt of the lenses is fixed by pins and the bleed-through from the dichroic is sufficient to align the lens to the beam-axis. The alignment laser itself is mounted to an optomechanical assembly that allows alignment of the laser (x/y, pitch/yaw) to the optical axis of the objective lens port (which provides a sufficiently accurate proxy for the objective lens optical axis) via two adjustable irises. The parts required are included in the parts list (Supplementary Table 2) and with all CAD files.

#### Infinite conjugate transmission target and infinite conjugate viewing camera

Assembly of the emission path requires an infinite conjugate transmission source to provide a reference for focusing of an object at infinity (as provided by an infinity corrected objective imaging at its native/design focal plane). Since incorrect spacing of the emission path lenses may produce a focused image featuring strong spherical and axial chromatic aberrations, this reference provides a route to check that the emission path does indeed focus to the correct plane for which the objective lens is designed. Setting up such a source requires an infinite conjugate viewing camera. The infinite conjugate transmission target assembly includes a transmission target (a coverslip with flecks of ink from a permanent marker is sufficient) and a focusable lens (f = 200 mm). This may be backlit by a suitable white light LED, LED array or flashlight provided by a smartphone as desired. Such a backlight is required for several other steps in the protocol. The infinite conjugate viewing camera comprises a camera and focusable lens (f = 100 mm). Any c-mount camera can be used in principle. Although the camera may also be used for beam profiling (as desired), using the imaging sCMOS camera is not advised as it is useful to have continued access to the infinite referencing assemblies even after the microscope is constructed. To aid focusing of the two assemblies, an adjustable optomechanical scheme is provided as a CAD assembly using COTS components. The parts required are also included in the parts list (Supplementary Table 2).

#### Shear plate interferometer

Assembly of both the excitation and emission paths requires a shear plate interferometer (referred to as shear plate throughout the protocol) to collimate single-mode fiber sources and judge distance between lenses. The shear plate provides a readout of the collimation state of the laser via the presence of interference fringes and their orientation with regard to a witness line. The parts required and the CAD assembly are included in the parts list (Supplementary Table 2). Refer to the section: Use of the shear plate interferometer for correct operation of the shear plate.

#### Alignment targets

Several targets are used throughout to aid with construction and alignment. For example, owing to the position of the emission path tube lens within the microscope body, it is difficult to use the back reflection method (see Orienting and aligning optical elements) to align the lens to the alignment laser beam installed in the objective port. Instead, an alignment target is used to position the tube lens mount appropriately such that the alignment laser passes through two obstructions in series. In some cases, it is necessary to precisely position and configure COTS components to position a target (often an iris) at a specific beam height. In these cases, use calipers to set the height of the assembly. For example, in the typical case of a ½” optical post and optical post holder, the height can be adjusted by sliding the post into the holder with the callipers and tightening the screw to secure it in place. Omit the components mounted to the post and use the calipers to push the post slowly into the holder with the holder screw just tight enough to provide some resistance. Determine the correct spacing of the holder base and top of the post and secure in position at said spacing. When using the target, ensure that the axis of the iris is coaxially aligned to the beam (i.e., not tilted). Moreover, for iris targets mounted on posts, consider that rotating about the post axis results in both translation and rotation of the iris since the iris itself is not aligned to the rotational axis of the post.

### Reagent preparation

#### Enzyme stock solution

Prepare 5 mL of a 20x concentrated enzyme stock solution by mixing 60 mg Glucose Oxidase (168 U/mg), 3 mL of 85% Glycerol, 250 µL of 1 M Tris-HCl pH 8, 330 µL Catalase (44560 U/mg) and 1.42 mL dH_2_O. Let the solution mix on a rotating wheel for 1-2 h. Store at -20°C.

#### dSTORM imaging base buffer

Prepare 100 mL of blinking buffer by mixing 50 mL of 10% 2- Deoxy-D-glucose, 5 mL of 1 M Tris-HCl pH 8, 1 mL of 1 M NaCl. Fill the solution up to 100 mL by adding 44 mL dH_2_O. Mix the solution by inverting the measuring cylinder 3-5 times. Aliquot the buffer into 950 µL volumes and store at -20°C.

#### dSTORM imaging buffer

Prepare the final dSTORM imaging buffer by adding 50 µL of the 20x concentrated enzyme stock solution and 10 µL of β-mercaptoethanol (14.3 M) to the 950 µL buffer stock solution.

CRITICAL: Prepare this solution freshly each time. Oxygen depletion is achieved enzymatically via Glucose Oxidase and Catalase. Over time this leads to the acidification of the buffer, which is detrimental to imaging as it can alter the photophysical properties of the dyes.

#### Cell fixation solution

Prepare 6.5 mL of 2.4% PFA cell fixation solution by combining 975 µL of 16% PFA, 2275 µL of dH_2_O and 3250 µL of 2x PBS.

CRITICAL: Prepare this solution freshly each time.

#### Cell permeabilization stock solution

Prepare 500 mL cell permeabilization stock solution by adding 2 mL of Triton X-100 to 250 mL 2x PBS and 248 mL dH_2_O. Store at room temperature.

#### Quenching stock solution

Prepare 1 L of 1M NH_4_Cl stock solution by dissolving 26.745 g NH_4_Cl in 1 L 1x PBS. Mix using a magnetic stirrer and then autoclave the stock solution.

For 1L of the final quenching solution at 100 mM NH_4_Cl, add 100 mL to 900 mL PBS. Store at room temperature.

#### SNAP-Surface BG-AF647 aliquots

Prepare 1 mM SNAP-Surface BG-AF647 stock solutions by dissolving 50 nmol BG-AF647 in 50 µL anhydrous DMSO. Vortex the solution, aliquot and store at - 80°C.

#### WGA-CF680 aliquots

Prepare a 1 mg/mL stock solution by dissolving 1 mg of WGA-CF680 in 1 mL dH2O. Aliquot and store at -80°C.

#### DTT aliquots

Prepare 1 M DTT solution by dissolving 1.55 g DTT in 10 mL dH2O. Aliquot and store at -20°C.

#### BSA stock solution

Prepare a 2% w/v BSA stock solution by dissolving 1g BSA in 50 mL PBS. Filter the solution through 0.22 um Millex-GP filters. Store at 4°C.

#### Labelling solution

Prepare 1 mL of labelling solution by adding 1 µL of 1 mM SNAP-Surface BG- AF647, 1 µL of 1 M DTT and 250 µL of 2% BSA to 748 µL of PBS.

CRITICAL: Prepare this solution freshly each time.

#### Culture medium

Prepare culture medium by adding 50 mL FBS, 5 mL MEM NEAA and 5 mL GlutaMAX to 500 mL DMEM. Store at 4°C, pre-warm to room temperature before using.

#### Objective lens cleaning solution

Prepare a mixture of 85% Petrol ether and 15% Isopropanol vol/vol in a dropper bottle.

### Sample preparation

#### Magnetic sample holder

All samples described in the following sections use the same sample holder set up (see Supplementary Figure 6). In short, a magnetic metal plate with a hole in its center serves as a base for the sample Ill24 mm coverslip. By placing a ring magnet on top, the coverslip is fixed in place, and a chamber is formed that can be filled with the required imaging solution. The magnet ring is encased in a hard PVC casing. To ensure thorough sealing, an O-ring is installed in a corresponding groove on the underside of the magnet casing. Since the ring magnet is in direct contact with the sample and the sample buffer, it must be wrapped in fresh parafilm to avoid carryover of fluorescent material between experiments. To wrap the magnet, a square piece of parafilm is placed on the underside of the ring magnet (clean side facing outward). The edges of the square piece of parafilm are then folded over the outside of the ring and fixed on top. Using a 1 mL pipette tip, the parafilm is pushed through the center of the ring from below (see Supplementary Figure 6). The edges of the resulting tube can then be pulled over the top using the same pipette tip. The magnet should now be completely wrapped by parafilm and can be placed on top of the coverslip and filled with imaging solution. For prolonged imaging of biological samples, acidification of the imaging buffer can be slowed down by additionally sealing the chamber with parafilm on top.

#### Fluorescent bead samples

Fluorescent bead samples are required for instrument alignment and PSF assessment/fitting. For the former, a denser bead sample (ideally 50 - 100 beads per Ill35 µm FOV) is preferable to better indicate variations in intensity across the FOV while for the latter, a sparser bead sample (ideally 10 - 20 beads per Ill35 µm FOV) is preferable such that the PSFs derived from the bead containing sub-volumes do not overlap with each other. To prepare a bead sample, add 40 µL of 1 M MgCl_2_ at the center of a Ill24 mm coverslip, already placed on the sample holder. Dilute 0.8 µL of the bead stock solution in 360 µL dH_2_O and vortex the solution for 20 s. Then add the bead solution to the MgCl_2_ solution on the coverslip and pipette up and down until mixed. Incubate the sample for 3 min. Then replace the mix with 400 µL dH_2_O. Such a bead sample is generally stable for an hour, after which the beads will start to detach. To keep the beads attached to the coverslip for longer one can replace the dH_2_O with 400 µL of a 0.1 M MgCl_2_ solution.

Alternatively, one can also immobilize beads in the long term on a coverslip using poly-L-lysine. To this end, add 100 µL of poly-L-lysine to the center of a Ill24 mm coverslip. After 10 minutes gently wash the solution away using dH_2_O. Dry the coverslip using clean compressed air. The presence of the poly-L-lysine layer can be seen as a surface film by tilting the coverslip to the light. Mix the undiluted bead stock by sonicating/vortexing for 1 minute to break up aggregated beads. Prepare a bead stock as a 1:50 or 1:250 vol/vol dilution with distilled water for the dense and sparse sample respectively. Mix the diluted stock as previously described and pipette 100 µL onto the coverslip to cover the poly-L-lysine coating. Leave for 10 minutes. Gently wash away the bead stock with distilled water and dry with clean compressed air. Bead samples can be stored dry at room temperature for several weeks if protected from light.

When imaging fluorescent bead samples, ensure that they are immersed in water or buffer solution. This is particularly important when assessing the PSF quality or 3D imaging performance since spherical aberrations in aqueous media (n = 1.33 - 1.34) will be comparable to those encountered in tissues (n = 1.36 - 1.4), whereas air (n = 1.00) provides a much larger refractive index mismatch between the coverslip and sample and stronger spherical aberration accordingly. In addition, when assessing conditions for (total internal) reflection from the coverslip, aqueous media provides a reasonable approximation of tissue and allows the required obliquity of the incoming beam to be determined more accurately, with only small readjustments required for tissue.

#### Fluorescent dye sandwich

A dye sandwich is used to assess and optimize the uniformity of the illumination field. To this end, place 1 µL Alexa Fluor 647 dye on a Ill24 mm round glass coverslip. Place a smaller, 12 × 12 mm glass coverslip on top of the droplet and gently press down until all the volume is spread evenly. Seal the dye sandwich using clear nail polish. Mount the sample and illuminate at low laser power.

#### Coverslip cleaning

Before any biological sample can be prepared, high-precision Ill24 mm round glass coverslips have to be cleaned to remove any contamination. First separate them into a coverslip mini-rack that can withstand strong acids. Then place the entire rack into a glass container and submerge it in a methanol:hydrochloric acid (50:50) mixture. Incubate the coverslips overnight (16 –24 h) while stirring using a magnetic stirrer. Following that, rinse the coverslips repeatedly with dH_2_O until a neutral pH is reached. Place them into a laminar flow cell culture hood to let them air-dry without risking new contamination. Finalize the cleaning by irradiating the coverslips inside the laminar flow cell culture hood with ultraviolet light for 30 min.

#### Biological sample preparation

Nuclear pore complexes (NPC) are used as reference standards to test the performance of the microscope [49]. Preparation of these biological samples can be categorized as follows: Cell culture and seeding; cell fixation and permeabilization; NPC staining.

##### Cell culture and seeding

U2OS cells that endogenously express Nup96-SNAP can be obtained from Cell Line Services (cls.chop, #300444) [49]. Culture the cells using a 25 cm^2^ cell culture flask containing 6 mL culture medium in an incubator at 37°C and 5% CO_2_.

##### CAUTION

U2OS cells are sensitive to sparse culturing conditions and should not be cultured at a confluency lower than 50%.

Passage the cells every 2 – 3 days when a confluency of almost 100% is reached. To this end, aspirate all media and rinse once with sterile PBS. After removing the PBS solution again, add 1 mL of pre-warmed TrypLE and incubate for 12 min in the incubator. The cells should now all be detached from the growth surface. Resuspend the detached cells by adding 5 mL pre-warmed cell culture medium and pipetting the entire volume up and down 3 times.

The cells are now in suspension and can be used for seeding. Prepare a 6-well-plate and add a previously cleaned coverslip and 2 mL cell culture medium per well. This allows for up to 6 samples to be prepared simultaneously. Now add 150 µL of the cell suspension to each well. Shake the plate to evenly distribute the cells and return it to the incubator. After two days of incubation, this procedure should yield a confluency of 50% - 70% on the coverslip and the sample is ready for fixation.

For regular cell culture maintenance, 3 mL of the total 6 mL cell suspension has to be removed and replaced with fresh pre-warmed cell culture medium.

##### Cell fixation and permeabilization

Two days after seeding, the cells are ready for fixation. Prepare one 6-well plate containing 2 mL cell fixation solution per well and one 6-well plate containing 2 mL cell permeabilization solution per well. Transfer the coverslips to the 6-well plate containing cell fixation solution. After 30 s pre-fixation, transfer the coverslips to the 6-well plate containing cell permeabilization solution and incubate for 3 min on a shaker. Following this, transfer the coverslips back to the wells containing the cell fixation solution and incubate for 30 min on a shaker. Quench the fixative by transferring the coverslips to a 6-well plate containing 2 mL quenching solution and incubate for 5 min on a shaker. This is followed by two washing steps. To this end, aspirate the quenching solution and replace it with 2 mL PBS. Incubate for 5 min on a shaker, then repeat this washing step one more time.

##### NPC labelling

Prepare a clean surface for the labelling procedure. We recommend using a suitably sized metal plate and attaching parafilm (clean side up) on top of it using tape. For each coverslip, add a droplet of 120 µL ImageiT Signal Enhancer directly on the parafilm. Now place a coverslip onto each droplet, with the cell growth surface facing downward. Incubate at room temperature for 30 min. Transfer the coverslips back to a 6-well plate (make sure to invert the coverslip, with the cell growth surface facing upward). Rinse 3 times with 2 mL of PBS per coverslip. Replace the parafilm on the metal plate and add droplets of 150 µL labelling solution. Next, place each coverslip with the cell growth surface facing downward onto each droplet. Cover the sample and incubate for 2 - 3 hours at room temperature in the dark. To avoid drying out of the sample, it is recommended to add a Kimtech wipe soaked in dH_2_O underneath the cover.

Before imaging, transfer the coverslips back to a 6-well plate and wash the sample 3 times in 2 mL PBS per well for 5 min in the dark on a shaker.

NPC dual-color labelling: Following the steps described above, the NPC channel can be labelled as an additional target for dual-color imaging using WGA-CF680. To this end, further dilute the WGA- CF680 aliquot 1:100 in 1% BSA, place droplets of 150 µL solution on clean parafilm and place a coverslip onto each droplet with the cell growth surface facing downward. Incubate for 5 min at room temperature in the dark. Before imaging, transfer the coverslips back to a 6-well plate and wash the sample 3 times in 2 mL PBS per well for 5 min in the dark on a shaker.

## Equipment Setup

Prior to commencement of the protocol itself, ensure that all parts listed in the parts list (Supplementary Table 2) are available. For custom made parts, check all toleranced dimensions with calipers, tapped holes with their respective threaded component and dowel pin holes with correctly toleranced dowel pins. Dowel pin holes may become undersized during the anodizing process. If this occurs, the holes should be re-machined to the required tolerances. We note a few general points regarding laser safety, optical alignment and best working practices. The various aspects described are not repeated throughout the protocol. The builder is assumed to have familiarized themselves sufficiently with the notes below before proceeding.

### Conventions

Throughout the protocol, elements in the various optical paths are referred to as being situated or placed up- or downstream. Upstream refers to a position closer to the light source or earlier in the sequence than the current element or position currently considered. Conversely, downstream refers to positions further away from the light source or later in the sequence than the subject element. Naturally, upstream elements have impacts on downstream components and generally downstream elements have no impact on upstream components. When referring to alignment of components, the terms lateral and longitudinal are used to refer to positioning of the element relative to the beam axis. The two lateral axes are mutually perpendicular to each other and the beam axis. The lateral axes may thus include a height component when the beam is travelling parallel to the optical table surface. The longitudinal axis is aligned to the beam axis. Furthermore, when specifying that a given component should be introduced, this indicates that the component should be placed approximately as determined by the CAD or as otherwise indicated without achieving a final position or alignment. For example, one may be instructed to introduce a lens, implying that the lens should be oriented correctly, placed in an appropriate position to elicit an effect but without securing in position. Conversely, when instructed to install a component, this indicates that the element should be placed correctly and secured in place. Often this will concern components whose position is fixed by other pre-installed/assembled features. It is assumed throughout that introduction and installation of components takes place with reference to the CAD design. There are cases where the positioning of components may ultimately depart from the CAD design, for example, when placing lenses in positions to maintain beam collimation and conjugation of various upstream/downstream optical elements, which is affected by focal length tolerances, precise collimation states of the probe laser and uncertainty of the precise back focal plane position of the objective lens. In these cases, adjustments must be made to the positioning of downstream components accordingly.

### Mechanical assembly

#### Securing components

The various mechanical parts comprising the microscope should be fastened with appropriate screws. The specific screw to use is not included in the parts list or CAD assembly unless there is a specific reason for doing so, e.g., when providing an important mating feature. Metric screws are used almost exclusively throughout. Exceptions to the metric format are called out in the protocol. For example, UNC #4-40 screws are used as required. The builder is expected to provide a reasonable stock of appropriate screws. To some extent the builder is free to choose the screws they want to use although the screw type is fixed by the respective feature in the mechanical part. Where generic screws and other small fittings (e.g., dowel pins, hex standoffs) are included in the parts list (typically since they constitute a mating feature in a CAD assembly), the part number for a US-based vendor has been provided owing to their extensive CAD support (McMaster-Carr), similar parts can be sourced from other suppliers.

In some cases, it is necessary to use a thread adapter to affix components with different threads. For example, M4/M6 external/external thread adapters are required (see parts list). When selecting a screw length consider the depth of the threading and that 3 - 5 threads are sufficient for robust tightening. For example, for an M6x1.0 thread, 3 - 5 threads are equal to 3 - 5 mm. In the construction of the body, we specify the length of screw and dowel pin to use for best results. Where possible the addition of a washer ensures the most robust fastening and avoids damage to anodizing or components themselves resulting from over tightening of screws. Builders may seek generic information about optimal screw tightness and tighten with a torque wrench if desired (not specified in tools required). However, as a good rule of thumb, tighten the screw to finger tightness by rotating the long arm’s axis of the hex key between one’s fingers and then tighten no more than a quarter turn by rotating the short arm while holding the long arm for superior torque.

#### Use of positionally locating mechanical features

Several of the custom components feature locating features. In the case of dowel pins, install in all available positions unless otherwise informed, for example, where dowel pins may over-specify positions under conditions of tolerance stack-up, or where additional adjustment is required. In some cases, the position of a component can be fixed in one or two dimensions by a locating surface or pocket. For example, the emission path tube lens can be removed and replaced repeatedly without misaligning the system owing to a pocketed mount. In these cases, ensure that components are installed with the appropriate locating feature flush to the surface(s). In the absence of locating features, components should be installed in the position determined by screws and without biassing screws to either side of their clearance holes. For example, the optical table does not provide locating features and so assemblies that are directly affixed to the table must be manually positioned (e.g., the microscope body and the image splitter platform are the largest components affixed to the optical table).

### Optical alignment and practices

#### Handling optics

To avoid soiling surfaces of lenses, mirrors, beamsplitters and filters, wear suitable gloves when handling optical elements. Even so, one should avoid making contact with optical surfaces. This is particularly important for optics with large clear apertures and those installed in confined spaces, which are easily soiled by accident during installation. For example, the emission path tube lens should be introduced and adjusted with particular care. If soiled, optics should be cleaned immediately flowing best practices with solvents, lens tissue and without applying pressure. Flat optics are easily cleaned fully while lenses or optics in housings (e.g., filters) can be more difficult. When cleaning housed optics, use minimal solvent to prevent its ingress into the housing. Lenses and mirrors can become soiled with dust over time. Purified air dusters or bulb blowers may be used to remove dust from optics periodically while persistent soiling may require removal and replacement of the optic. In some cases, and with care, this can be performed without misalignment of the system. When securing mirrors in mounts ensure that they are firmly held but do not overtighten, which even for thick optics can result in warping. Similarly, for lenses in lens tubes secured by retaining rings, do not overtighten the rings to avoid damage to the optic and where possible, install the second retaining ring with the lens tube facing up/down such that the optic sits as flat as possible. Always use an appropriate spanner wrench (e.g., COTS-SPW602) to tighten retaining rings (COTS- SM1RR).

#### The shear plate interferometer

The shear plate interferometer is an optical device allowing judgement of the collimation state of the laser. Generally, the user is advised to follow the manufacturer notes. However, we note that additional parts are specified to aid normal alignment of the beam onto the device (see parts list/CAD files). In addition, consider that the human eye is most able to resolve fine details in the green-yellow portion of the visible spectrum. When a choice of wavelengths is available such as is the case of the laser engine(s) opt for the closest wavelength to 550 nm (e.g., 561 > 532 > 488 > 640 > 405 nm) and use minimum power or attenuate with neutral density filters to allow the eye to better resolve the interference fringes. Note for beams smaller than ca. 3.0 mm, the size of the fringes will be small enough that may make visual inspection difficult. In such a case it is helpful to use a magnifying adapter for the viewing screen. The use of the shear plate is not possible nor necessary with the NIR lasers for the focus lock system. Throughout the protocol, no reference is made to the size of beam or the specific plate to use. In each case, this is readily determined by simple considerations of input beam size and system magnification.

#### Walking the beam

Throughout the protocol it is necessary to walk the beam to achieve fixed height travel from the optical table, potentially with alignment to other alignment features, or to ensure the beam passes correctly into or out of the objective lens port. Doing so usually requires two spaced targets at the correct height/position placed downstream of the element(s) that will be adjusted. For an arbitrary height and angle input, two steerable elements are required to walk the beam. One should iteratively use the first steerable element to align to the closest target and the second to the furthest target.

#### Orienting and aligning optical elements

The spectrally discriminative optics (filters and dichroics) have dielectric coatings requiring a specific orientation of the optic, which is often indicated by a caret or arrow on the side of the optic. Consult manufacturer datasheets for correct orientation, which differs between manufacturers. Lenses have a specified orientation with their flatter (greatest radii of curvature, thus lower optical power) surface facing the beam focus. When uncertain, consult the CAD and/or Figure 2 for guidance. In the case of the tube lens, the infinite conjugate (collimated) direction is indicated by an infinity symbol. However, for lenses mounted in tubes it is easy to lose the orientation and align the lens incorrectly. For this reason, it is good practice to indicate the infinite conjugate side of the lens on the lens tube. Additionally, when aligning lenses mounted in tubes via a split clamp, lock the lens position with the clamp beforehand and loosen only to adjust collimation before locking again. Doing so ensures that the lens is correctly positioned during the alignment process. The specific degrees of freedom available will determine how a lens is aligned. Generally, one starts with a beam travelling at a fixed height from the optical table and potentially aligned to specific optical table hole features. The lens is aligned by positioning its optical axis to the beam axis. When the lens is placed correctly, the resulting beam does not deviate from its original course but may change in diameter as it propagates due to focusing by the lens. In most cases during the protocol, one is able to position lenses laterally (orthogonal to the optical axis) and longitudinally (along the optical axis). Lateral positioning is most critical for minimizing beam-deflection, while correct positioning in z is critical to achieve collimation and conjugation of various optical elements in the system. In addition, the yaw angle can be adjusted by rotating lenses on their mounting posts, while the pitch is fixed. In this case, incorrect placement/orientation leads to only small beam deflections but contributes to off-axis aberrations such as astigmatism and coma. To align a lens, place and secure iris targets up- and downstream of the intended lens position aligned to the fixed height beam. Introduce the lens and laterally position it to roughly align the beam through the downstream iris. Check for back reflections from the lens, which should be visible on the backside of the first iris. If they are not present, adjust the lens yaw until they can be seen. The objective is to align the back reflections to each other and back through the iris. The relative positions can be adjusted by lateral translation and the combined back reflections can be redirected back through the iris by adjusting the pitch and yaw. Since pitch is typically not an adjustable degree of freedom it may not be possible to achieve perfect overlap of the back reflections and vertical alignment back through the iris. In this case, assuming the input beam is indeed propagating at a fixed height, horizontal overlap of the back reflections is sufficient. The longitudinal alignment of a lens depends on whether one has a collimated or divergent output. Bringing a divergent beam into collimation is achieved by longitudinal positioning of the lens and assessing the collimation on a shear plate interferometer. Generally, in the case of the placement of lenses to focus a collimated beam, the protocol will outline steps to present the alignment as a collimation step. In some cases, trying to simultaneously ensure collimation and alignment to the optical axis presents a challenge that is best broken down into constituent parts. First the position for collimation is determined (with very rough alignment to the optical axis) and a locating feature is installed to allow the longitudinal position to be indicated for subsequent alignment to the axis. This will be indicated in the protocol.

#### Affixing dichroic beamsplitters in glue-in mounts

Improper mounting of dichroic beamsplitters whereby differential stress is applied, e.g., via a retaining screw, will result in curvature of the mirror resulting in strong aberrations e.g., astigmatism/coma in the resulting image. To avoid this occurrence, the dichroic mounts are designed to mount the elements via an adhesive only. Note, that CAD files for the dichroic mounts are provided for 1 (illumination only), 3- and 5-mm thick substrates with lateral dimensions no larger than 25.5 × 36 mm. For other lateral dimensions or thicknesses, the builder is advised to make the appropriate modification to the CAD part. It is imperative that the correct thickness is used with the correct mount for alignment purposes. Generally thicker dichroics are able to hold a tighter tolerance on surface flatness, (λ/10 desirable over the used aperture, but as low as λ/4 may prove acceptable). This is particularly important for reflection of the emitted fluorescence by the emission dichroic, where a poor surface flatness will result in aberrations in the final image. Additionally thicker substrates are more resistant to bending under stress. Nevertheless, dichroics should not be mounted in such a way that stresses are applied. Before mounting the dichroics, clean the mounts with isopropanol and clean compressed air.

Regarding choice of adhesive, A UV-cured epoxy is well-suited with the following caveats: i) the dichroic should be affixed only at two corners along the same edge of the optic, ii) the UV curing process should not be carried out under conditions of elevated temperature, e.g., under lamps. The mounting part will deform with changes in temperature and the epoxy affixes the optic very stiffly such that deformations of the mount may deform the beamsplitter. Minimal affixing points help to reduce any deformations. An alternative is to use a silicone sealant to hold the dichroic in place. The flexible sealant obviates deformations but does not provide as strong adhesion, so care must be taken to seat the dichroic correctly in the mount and in handling the mounted optic. The sealant can be easily removed, whereas epoxy should be considered a permanent solution, although the recommended epoxy can be removed with immersion in dichloromethane. Whichever solution is chosen, one should ensure that the dichroic sits flat in the mount and that adhesive does not seep under the optic. This is best achieved by holding the optic in place using light downward pressure from an optics-grade cotton tipped applicator on the dichroic during gluing.

#### The alignment laser and aligning with lasers

the alignment laser assembly includes two adjustable irises to ensure on-axis passage of the beam through the threaded assembly thus ensuring that the beam enters the microscope emission path aligned to the optical axis of the objective lens. Use the pitch/yaw (COTS-KAD11F) and X,Y mounts (COTS-CXY1) to iteratively align the laser through the first and second irises respectively. The beam incidence on the irises may be viewed through the cut-outs in the slotted lens tubes with the covers rotated appropriately. It is assumed in the protocol that the alignment of the alignment laser through the two irises is correctly established and maintained throughout. During alignment, the second of these irises may be stopped down to limit the size of the beam, which generally improves accuracy in positioning of lenses and alignment to various downstream apertures and targets. When assessing beam collimation, for example, in the axial positioning of lenses, a larger beam is preferable for readout of the collimation state and so the iris should be fully opened. When inserting optical elements into the beam path, first turn off the alignment laser and be aware that the beam path may have changed. Use laser safety screens (see parts list) as appropriate to intercept stray beams. When using laser sources to image samples, it is assumed that the microscope cover is installed. This is particularly important when aligning the illumination and focus lock beams for (total internal) reflection from the coverslip, where the beam will be launched obliquely from the objective lens constituting a particular laser safety hazard. Otherwise, the instruction to ensure the cover is correctly installed is left to the builder to exercise good judgement. When using targets for alignment purposes or the shear plate interferometer, secure these parts with appropriate base clamps to avoid losses of position due to unintended movement of the element.

#### Use of neutral density (ND) filters

The neutral density filters offer a logarithmic scale for laser attenuation. At certain stages of the protocol, it may be necessary to use ND filters to attenuate the beam, particularly when it will be incident on a camera (beam profiling cameras may include an appropriate ND filter). To avoid damage, never allow an unattenuated laser to illuminate the camera chip. Start with a high ND filter (e.g., ND6) and work downwards to achieve the required attenuation. Note that absorptive ND filters can be stacked to provide additive neutral density. If using pre-existing reflective ND filters, stacking may result in multiple reflections and is thus not recommended. The parts list provided specifies neutral densities of 1, 2 and 4. By combining these filters one may achieve effective ND1-7. When placing ND filters, ensure that their face is perpendicular to the beam optical axis to avoid steering the beam unintentionally.

#### Cleaning the objective lens

The immersion oil(s) used can leave a residue on the objectives that is not removed by mild solvents such as isopropanol alone. We recommend routine cleaning of the lens after each imaging session and when replacing the immersion oil using a mixture of 85% Petrol ether and 15% Isopropanol vol/vol. For deeper cleaning of the objective lens, other mineral or immersion oils as well as optics grade polymer cleaning sprays/solutions are effective for removal of the residue. The builder is advised to proceed with caution applying other treatments and follow usual practices such as using only high-quality lens tissue, clean air-blowers and never applying pressure when cleaning lenses, filters and mirrors.

#### Use of the Bertrand lens

The Bertrand lens allows viewing of the objective lens back focal plane. Once the Bertrand lens is correctly set up (see: Emission path alignment (Timing: 5 days)) and a fluorescent sample is mounted and imaged in focus on the camera, the back focal plane of the objective lens is visible as a bright disk in one or both channels (depending on choice of excitation wavelength/emission filters). The focused single-mode excitation spot (if present) should be visible at the center of the disk for epi illumination or offset for HILO/TIRF. The focus lock laser spot, when correctly aligned, should also appear as a spot towards the periphery of the back focal plane. It may be possible in some cases to see a dimmer reflected spot on the opposite side of the back focal plane, which also allows the Bertrand lens to be used to judge the total internal reflection condition. The presence of dark occlusions on the disk indicates the presence of bubbles in the immersion oil, which aberrate the PSF and degrade image quality. As such, the Bertrand lens can be used for quality checking and should be used whenever mounting a sample. It may also be used effectively for multi-position acquisition workflows, where bubbles may move around as the oil is spread around the coverslip by its motion. When bubbles are seen to be present, clean the objective and sample with isopropanol and lens tissue before adding more oil and replacing the sample.

### Control of the microscope and hardware

#### Using µManager and the htSMLM GUI

Throughout the protocol a multitude of instructions will be given requiring interaction with software to control hardware and acquire data. The builder is expected to familiarize themselves with the software and controls prior to continuing with the protocol. For brevity, specific instructions regarding how to achieve a given command are provided. For example, the instruction: Image fluorescent beads, assumes that the user is familiar with previewing and acquiring images using the µManager GUI. Similarly, it is assumed that the user can set laser power, region of interest (ROI) and exposure time or select filters appropriately to achieve a desired temporal resolution and image signal to noise or exposure level.

#### Making connections with cables and handling electronics

A specific list of cables required is not provided since the lengths required will depend on how the hardware and room are laid out. However, some general points are worth considering. The microFPGA front panel connectors are all SMB-type (jack/female) and so a supply of SMB cables is generally required, additionally other hardware elements use SMA (sCMOS camera) and BNC (piezo flexure stage) connectors. The builder is free to choose connectors when packaging the focus lock QPD amplifier and so cables should be determined accordingly. The builder should take care to not over flex cables, particularly those that are not removable/replaceable such as those of the XY stage, QPD stage and piezo flexure stage. Take appropriate care when handling electronic boards to avoid electrostatic damage. Ideally handle all boards using an anti-static wrist band with earthing connector.

## Protocol

### Microscope body build-up (Timing: 3 days)

The microscope body provides the core of the microscope around which the illumination, emission and focus lock optical pathways are constructed. A CAD rendering of the completed body is shown in Figure 3. The microscope body is handed: in the base configuration, illumination is provided from the back side, while emission and focus lock paths are from the right/left respectively. Back/front and left/right are defined from the normal user viewing orientation at the short edge of the optical table and are inscribed onto the body lower plate (3D-SMLM-FAB-000001). Please see Supplementary Note 8: Configuring the microscope body and optical path handedness for a guide to configuration of the microscope body including discussion of achieving a differently handed layout on the optical table. In the base configuration, shown in Figure 1, access (for example to mount samples on the microscope) is primarily from the front of the microscope along the shorter optical table edge.

**Figure 3.**
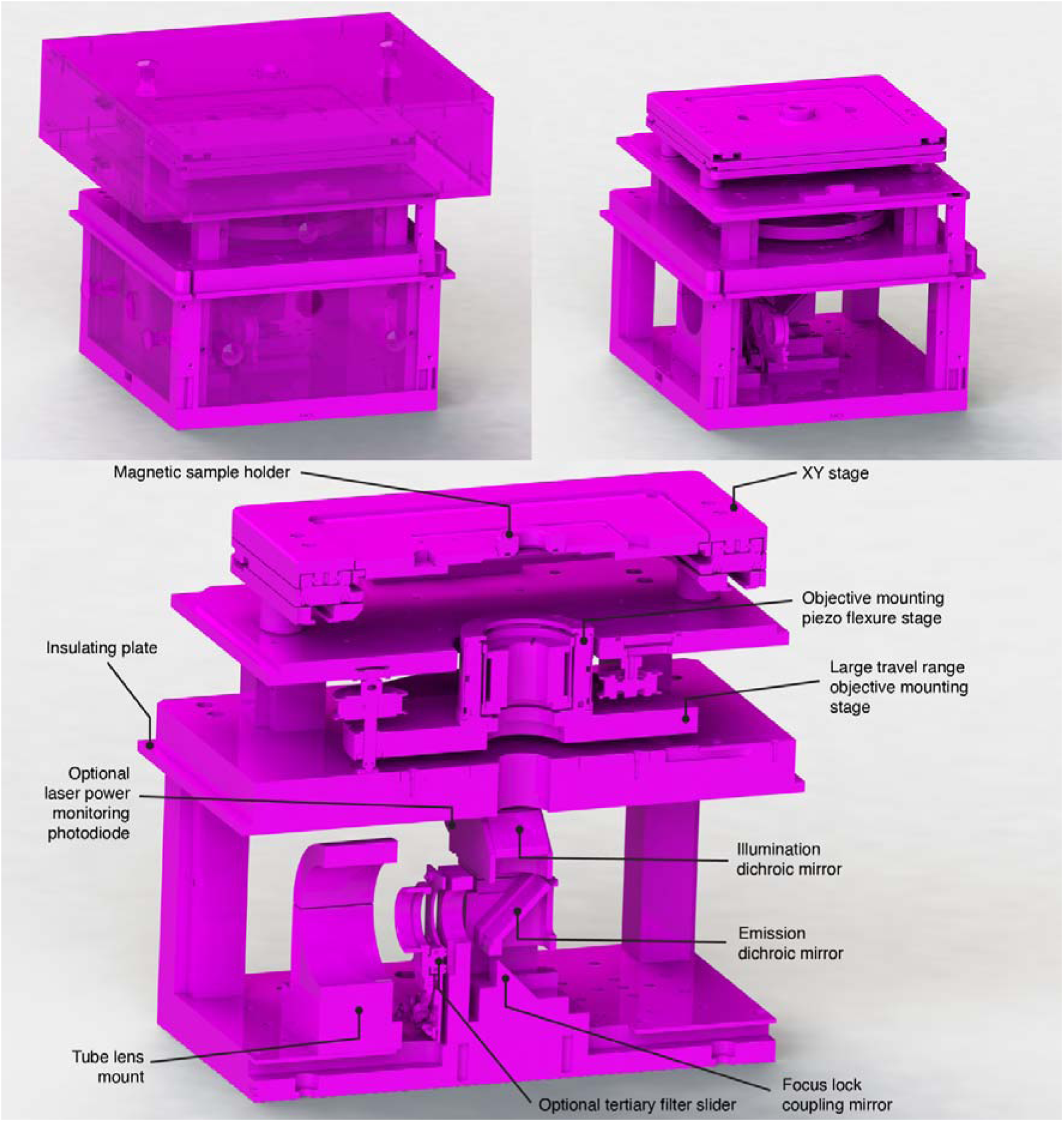
The fully assembled inverted microscope body. Top/left: The microscope body with covers installed for laser safety and to aid thermal stability. Top/right: The microscope body with covers removed. Bottom: a cut-away view of the microscope body showing its various motion control, optical and optomechanical aspects. The various optical components including high NA objective lens (omitted, mount shown) and dichroic mirrors for coupling the optical paths/in/out are situated below an XY stage. An optional filter slider can be included in the infinity space before a tube lens (omitted, mount shown) produces a primary image outside of the microscope body.

Secondary access is provided from the left side. Refer to the section: Laboratory environment for a discussion of optimal room layout.

1. OPTIONAL: Install the heating foil and/or temperature sensor in the central recess of the body middle plate (FAB-EMBL-000007). Note, the temperature sensor can still provide a useful readout of temperature in the absence of the heating foil.
2. OPTIONAL: Route the heating foil and/or temperature sensor cables along the cable channel to the slotted recess on the outside edge of the plate.
3. OPTIONAL: Cover the channels with their respective covers (FAB-EMBL-000051/52) and secure in place with 7× M2.5×6 mm countersunk screws.
4. OPTIONAL: Secure the heating foil cover (FAB-EMBL-000041) in place with 7× M2.5×8 mm countersunk screws.
5. Install the threaded bushings (COTS-F6MSSA1) in the objective piezo platform (FAB-EMBL- 000038) using a drill press.
6. Install the 6× dowel pins (COTS-91585A437) into the body middle plate (FAB-EMBL-000007) using a drill press.
7. Install the 4 mm ball bearings (COTS-9292K37) into the hollow end of the 3× actuators (FAB- EMBL-000039) using epoxy glue. Ensure that the epoxy has hardened fully before proceeding.
8. Use a small quantity of vacuum grease to lubricate the threaded section of the 3× actuators (FAB-EMBL-000039).
9. Install the 3× timing pulleys (COTS-HTPT48S2M060-A-P6.35) onto the 3× actuators and secure in place with the set screws contacting the flat surface.
10. Install the 3× actuators into the threaded bushings (COTS-F6MSSA1). Screw the actuator in just until the ball-tips protrude ca. 1 mm from the bushing. CAUTION! Take care of the orientation of the actuator in the bushing, if the bushing and actuator are incorrectly oriented, use of the actuator may act to remove the bushing from the objective piezo platform (FAB-EMBL- 000038).
11. Install the objective thread adapter (FAB-EMBL-000043) into the objective piezo flexure stage (COTS-P-726.1CD) using an adjustable spanner wrench (COTS-SPW801) to securely tighten it.
12. Install the objective piezo flexure stage onto the objective piezo platform (FAB-EMBL-000038).
13. Rest the three ball-tips of the actuators (FAB-EMBL-000039) on the three sets of pins installed on the body middle plate (FAB-EMBL-000007), orient the platform on the pins such that the cable of the objective piezo flexure stage (COTS-P-726.1CD) can be easily routed to the underlying cable channel of the body middle plate.
14. Remove the objective piezo flexure stage (COTS-P-726.1CD).
15. Install the 6× hooked springs (COTS-RZ-099l) with 6× retaining pins (COTS-91585A371) in the objective piezo platform (FAB-EMBL-000038) such that the springs pass through the clearance holes in the body middle plate (FAB-EMBL-000007).
16. Use a spring pulling hook to pull the springs fully through the body middle plate (FAB-EMBL- 000007) and install the 6× retaining pins (COTS-91585A371) such that objective piezo platform (FAB-EMBL-000038) and body middle plate are held together under spring tension.
17. Screw in the 3× actuators (FAB-EMBL-000039) such that the separation between the objective piezo platform and body middle plate is large enough to just permit 1× 7 mm optical post spacer (COTS-RS7M) to pass freely between the two components around the circumference of the objective piezo platform (FAB-EMBL-000038).
18. Insert 3× 7 mm optical post spacers (COTS-RS7M) between the objective piezo platform (FAB- EMBL-000038) and the body middle plate (FAB-EMBL-000007) close to the circumference of the objective piezo platform with approximately even spacing.
19. Lower the platform using the 3× actuators FAB-EMBL-000039 until the objective piezo platform (FAB-EMBL-000038) is supported by the 3× 7 mm optical post spacers (COTS- RS7M). One should feel the resistance to turning the actuator decrease at the point of touching. Note, it may be necessary to perform this adjustment iteratively to ensure flatness.
20. Raise the platform using the 3× actuators (FAB-EMBL-000039) just enough that the 3× 7 mm optical post spacers (COTS-RS7M) can be removed. Check that the spacing between the objective piezo platform (FAB-EMBL-000038) and body middle plate (FAB-EMBL-000007) is 7 mm throughout by moving the post spacers around the circumference. CAUTION! If the actuators are not adjusted correctly, they will not mate correctly with other components later in assembly and the objective lens will be tilted relative to the XY stage assembly.
21. Loop the timing belt (COTS-60S2M494) around the 3× timing pulleys (COTS- HTPT48S2M060-A-P6.35). Note, at this stage the belt will not be under tension and so will not hold on the pulleys. The assembly should now appear as shown in Supplementary Figure 7 (timing belt omitted).
22. Install the body upper standoffs (FAB-EMBL-000008/09/10/11) on the body middle plate (FAB- EMBL-000007) using 8× M4×16 mm dowel pins and 8× M4×20 mm socket head screws. CAUTION! Ensure that the parts are installed with the correct orientation to form 4× pairs of recesses for magnets on the four sides of the microscope.
23. Affix the illumination dichroic mount (FAB-EMBL-000026) to the top plate of its kinematic mount (COTS-KB25-M) using 1× M4×6 mm socket head screw, ensuring that the plate is situated flush with the sides of the recess marked by the corner clearance hole.
24. OPTIONAL: If wishing to make use of the laser power monitoring photodiode. Install the focus lock laser blocking filter (COTS-NF808-34) in its lens tube (COTS-SM1L03) with the caret pointing away from the externally threaded section. Secure the filter in place by threading the SM1 threaded photodiode (COTS-SM1PD1A) into the lens tube. Install the lens tube in the illumination dichroic mount (FAB-EMBL-000026).
25. Install the emission and illumination dichroic mirrors (COTS-25.5×36x3MM-DICHROIC- EM/COTS-25.5×36x3MM-DICHROIC-ILL) into their mounts (FAB-EMBL-000025/26 respectively) following the instructions in the section: Optical alignment and practices, Affixing dichroic beamsplitters in glue-in mounts. Note, if using 5 mm thick dichroics and/or a mirrored configuration of the body, install the mirrors into the respective mounts. Consult the CAD assemblies for respective part numbers and adjust the following instructions accordingly. See Supplementary Note 2: Selecting spectrally discriminative optics.
26. Install the prism mirror (COTS-MRA20-E03) into the prism mirror mount (FAB-EMBL- 000024). Take care that the mirror sits flush with the two sides of the recess sharing the corner clearance hole.
27. Install the prism mirror mount (FAB-EMBL-000024) on the optic pillar base plate (FAB-EMBL- 000022) using 2× M3×16 mm socket head screws and 2× M3×10 mm dowel pins.
28. Install the optic pillar (FAB-EMBL-000023) onto the optic pillar base plate (FAB-EMBL- 000022) using 2× M4×12 mm socket head screws and 2× M4×10 mm dowel pins.
29. Affix the emission dichroic mount (FAB-EMBL-000025) to the optic pillar (FAB-EMBL- 000023) using 2× M3×12 mm dowel pins and 2× M3×30 mm socket head screws.
30. Install the lower mounting plate of the illumination dichroic kinematic mount (COTS-KB25-M) onto the optic pillar (FAB-EMBL-000023) using 1× M4×6 mm socket head screw taking care that the magnets, balls and three sets of locating pins line up correctly with the upper mounting plate of COTS-KB25-M (affixed to FAB-EMBL-000026), in the correct orientation. Ensure that the lower mounting plate sits flush with the sides indicated by the corner clearance hole.
31. OPTIONAL: If wishing to make use of the four-position shared-path filter slider (COTS-ELL9) inside the body, install the filters (COTS-25MM-HOUSED-FILTER) individually mounted in lens tubes (COTS-SM1L03) into the corresponding SM1 thread on the filter slider. See Supplementary Note 2: Selecting spectrally discriminative optics. CAUTION!: Ensure filters are correctly oriented with the caret pointed as indicated by the filter manufacturer.
32. OPTIONAL: Install four-position shared-path filter slider (COTS-ELL9) on its mounting plate (FAB-EMBL-000027) via 4x #4-40 hex standoffs (3D-SMLM-COTS-91075A873) using 4× #4-40x⅜” socket head screws.
33. OPTIONAL: Install the mounting plate (FAB-EMBL-000027) onto the optic pillar base plate (FAB-EMBL-000022) using 1× M4×12 mm socket head screw and 2× M4×10 mm dowel pins. Note, whenever using the alignment laser (3D-SMLM-EMBL-007) for alignment of the emission path, switch the filter slider to an open position.
34. Install the body lower standoffs (FAB-EMBL-000002/3/4/5) onto the body lower plate (FAB- EMBL-000001) using 8× M4×20 mm socket head screws and 8× M4×12 mm dowel pins. Take care that the standoffs are oriented correctly and installed on the correct corner of the body lower plate.
35. Install the optic pillar base plate (FAB-EMBL-000022) on the body lower plate (FAB-EMBL- 000001) using 4× M6x10 mm socket head screws and 2× M4×12 mm dowel pins.
36. Install the body lower plate (FAB-EMBL-000001) on the optical table (COTS-M-VIS3660-PG4- 325A) using 2× M6x10 mm and 2× M6x20 mm socket head screws. Note, the shorter screws should be used to secure the plate through the counterbores in the cable channels. CAUTION! The location of this plate will determine the respective location of all other components of the microscope relative to the optical table. Ensure that it is installed at the correct position, allowing space on the table for the optical pathways.
37. OPTIONAL: Route the four-position shared-path filter slider (COTS-ELL9) ribbon cable out of the body and secure with its cover (FAB-EMBL-000050) and 4× M2.5×6 mm countersunk screws.
38. Install the tube lens mount (FAB-EMBL-000035) onto the tube lens base (FAB-EMBL-000034), securing with 4× M6x16 mm socket head screws.
39. Loosely install the tube lens retaining cap (FAB-EMBL-000036) onto the tube lens mount (FAB-EMBL-000035) using 2× M4×10 mm socket head screws.
40. Install the tube lens base (FAB-EMBL-000034) onto the body lower plate (FAB-EMBL- 000001), in an approximate nominal position securing with 3× M6x20 mm socket head screws and 3× M6 washers.
41. Install the mounted illumination dichroic (COTS-25.5×36x3MM-DICHROIC-ILL, FAB-EMBL- 000026) by reuniting the two halves of the kinematic mount (COTS-KB25-M).
42. Install 8× M4×20 mm dowel pins into the protruding end of the body lower standoffs (FAB- EMBL-000002/3/4/5). The assembly should now appear as shown in Supplementary Figure 8 (dowel pins omitted).
43. Place the insulator plate (FAB-EMBL-000006) onto the body lower standoffs (FAB-EMBL- 000002/3/4/5) such that the dowel pins pass through the clearance holes. Ensure that the plate is oriented correctly with respect to the standoffs and the body lower plate (FAB-EMBL-000001)
44. Install the body middle plate FAB-EMBL-000007 on the insulator plate (FAB-EMBL-000006), taking care to orient correctly with respect to the lower portion of the body. Secure in place with 8× M4×25 mm socket head screws.
45. Install the objective piezo flexure stage (COTS-P-726.1CD) onto the objective piezo platform (FAB-EMBL-000038).
46. Route the objective piezo flexure stage cable out of the body using the channels in the body middle plate (FAB-EMBL-000007), securing in place with the channel covers (FAB-EMBL- 000054/55) and 5× M2.5×6 mm countersunk screws. The assembly should appear as shown in Supplementary Figure 9.
47. Insert the 8× magnets (COTS-S-08-05-N) into their recesses in the body upper standoffs (FAB- EMBL-000008/09/10/11) and 4× covers (FAB-EMBL-000018), ensuring that the magnets are oriented correctly to attract and allow for interchange of the identical covers (FAB-EMBL- 000018) between the four sides. Remove the covers afterwards.
48. Glue the magnets in place using a two-part epoxy adhesive (COTS-JB-WELD) once the correct orientation is confirmed.
49. Use a drill press to install the three flanged bearings (COTS-SFL625ZZ) into the body upper plate (FAB-EMBL-000012).
50. Use a drill press to install the idler (COTS-SFD11-25) onto its mounting pin (FAB-EMBL- 000064).
51. Loosely install the idler and mounting pin onto the underside of the body upper plate (FAB- EMBL-000012) using an M4×10 mm socket head screw such that the screw and pin can be translated along the slot. Note, that there are three possible recessed slots for the screw, allowing for placement of the idler as desired. However, the recommended position is indicated by the CAD files and shown in Supplementary Figure 10.
52. Install 8× M4×12 dowel pins into top of the body upper standoffs (FAB-EMBL- 000008/09/10/11).
53. Carefully lower FAB-EMBL-000012 onto the 8× protruding pins and bosses of the 3× actuators (FAB-EMBL-000039), which mate with the bearings (COTS-SFL625ZZ) while capturing the outer flat side of the timing belt (COTS-60S2M494) between the two flanges of the idler (COTS-SFD11-25). TROUBLESHOOTING! Depending on manufacturing precision and tolerance stack up, it may be difficult to mate all dowel pins and bosses with their corresponding mating feature. The pins can be reduced in number if the parts cannot be lined up otherwise.
54. Secure the body upper plate (FAB-EMBL-000012) to the body upper standoffs (FAB-EMBL- 000008/09/10/11) using 8× M4×12 mm socket head screws.
55. Ensure that the timing belt (COTS-60S2M494) is now looped correctly around the 3× timing pulleys (COTS-HTPT48S2M060-A-P6.35, inside the belt) and the idler (COTS-SFD11-25, outside the belt).
56. Tension the timing belt (COTS-60S2M494) by sliding the idler pin (FAB-EMBL-000064) along the slotted recess in the body upper plate (FAB-EMBL-000012) and tightly secure the M4 socket head screw when an appropriate tension is achieved.
57. Check that the objective piezo platform (FAB-EMBL-000038) can be moved up and down by rotating any of the timing pulleys (COTS-HTPT48S2M060-A-P6.35). Return the platform to the nominal 7 mm spacing using 1× optical post 7 mm spacer (COTS-RS7M) placed between the body middle plate (FAB-EMBL-000007) and objective piezo platform (FAB-EMBL-000038). TROUBLESHOOTING!: If the resistance is too strong to be able to move using a hex key inserted through the actuator (FAB-EMBL-000039) for leverage with reasonable application of force, first loosen the screws affixing the body upper plate (FAB-EMBL-000012) to the body upper standoffs (FAB-EMBL-000008/09/10/11) and retry. If this rectifies the issue, re-secure the screws. If this issue persists, the FAB-EMBL-000038 platform may not be flat. Recheck the spacing between FAB-EMBL-000007 and FAB-EMBL-000038 with the 3× optical post 7 mm spacers (COTS-RS7M) as previous. Otherwise, it is acceptable to remove additional material from the section of the actuator (FAB-EMBL-000039) that mates with the bearing or remove dowel pins from the body upper standoffs/body upper plate (FAB-EMBL-000008/09/10/11/12) as necessary to relax the tolerances on fit.
58. Install the XY stage (COTS-SOM-12090) onto the body upper plate (FAB-EMBL-000012) via the 4× stage standoffs (FAB-EMBL-0000047) using 4× M6x30 mm socket head screws. Note, it is necessary to move the stages from their central position to access the counterbored holes. The assembly should now appear as shown in Supplementary Figure 11.
59. Install the sample holder support (FAB-EMBL-000044) onto the XY stage (COTS-SOM-12090) using 4× M2.5×6 mm socket head screws.
60. Install the cable channel (COTS-EMBL-000059) between the body middle and upper plates (COTS-EMBL-000007/12), securing with 1× M2.5×6 mm socket head screw and 2× M2.5×6 mm countersunk screws.
61. Route the XY stage (COTS-SOM-12090) cables out of the body using the channels on the body upper plate (FAB-EMBL-000012), cable channel (FAB-EMBL-000059) and body lower standoff (FAB-EMBL-000003) securing in place with the respective channel covers (FAB- EMBL-000056/57/58) and 7× M2.5×6 mm countersunk screws while additionally capturing the objective piezo flexure stage (COTS-P-726.1CD) cable with channel cover (FAB-EMBL- 000058).
62. OPTIONAL: Install the channel cover (FAB-EMBL-000053) on body lower standoff (FAB- EMBL-000059), capturing the heating foil and temperature sensor cables, as appropriate, using 2× M2.5×6 mm countersunk screws. The assembly should now appear as shown in Figure Supplementary Figure 12.
63. Assemble the microscope cover by affixing 2× back/front cover (FAB-EMBL-000019, 2× left/right cover (FAB-EMBL-000020), 1× upper cover FAB-EMBL-000021 and 1× lower clearance cover (FAB-EMBL-000066) using 24× M2.5×10 mm countersunk screws. Note, matching side panels should oppose each other.
64. Install the ring LED FAB-EMBL-000037 in the top plate of the upper cover (FAB-EMBL- 000021) using 4× M3×6 mm hex standoffs (COTS-98952A104) and 4× M3×5 mm socket head screws.
65. Install the 2× strain relievers (FAB-EMBL-000017) capturing the LED cable and tightening it to the left/right cover (FAB-EMBL-000020), as preferred.
66. Install 12× panel knob (COTS-24540.0021) to the covers (FAB-EMBL-000013/14/15/16, 4× FAB-EMBL-000018, FAB-EMBL-000021) using 12× M5×10 mm countersunk screws.
67. Install all covers (FAB-EMBL-000013/14/15/16, 4× FAB-EMBL-000018, FAB-EMBL-000019/20/21/66) on the microscope checking for fit and appropriate routing of the LED cable. CAUTION: certain covers (FAB-EMBL-000013/14/15/16/18) can be installed in two orientations. For FAB-EMBL-000013/14/16, which have apertures at specific heights for the various optical pathways, ensure that the panels are installed the correct way round. The assembly should now appear as shown in Supplementary Figure 13.

### Assembly of the microFPGA (Timing: 3 days)

If the microFPGA [40] has not been directly sourced in a pre-assembled state, the constituent boards must first be assembled, configured with solder bridges or jumpers, and tested for the specific use case presented. The microFPGA is then assembled with its enclosure: boards are mechanically mounted and the Breakout (Br) and custom shield (CS) are installed on the FPGA before all electrical connections are made. Note that the signal conversion board is not needed for any of the microscope components specified but adds flexibility for extension or changes of camera, which may operate at a 5V logic level that could otherwise damage the FPGA. The builder may omit this board and ignore all related protocol steps as desired, instead connecting the camera exposure input directly to the custom shield. A list of hardware and I/O used and additional capacity is given as Supplementary Table 5: Used microFPGA IO and Associated Hardware and Supplementary Table 6: Unused microFPGA IO and Potential Hardware. Note that the associated bill of materials includes pre-made cables whose ends can be clamped by a connector. When discussing the connection of pins in the protocol, the assumption is that the builder will use such a cable with both ends connectorised. When only one cable end should be connectorised, (e.g., when one end of the cable should be soldered) this is specifically called out. The cables and connectors are included in the parts list (COTS-661161122030, COTS-661002113322) Before connecting the microFPGA to a fixed power supply, it is recommended to use a variable lab power supply with current readout displayed to check current and ensure that there are no short circuits. The fully assembled microFPGA is shown in Supplementary Figure 14.

68. Assemble all boards as shown in the board files provided or have the components soldered by the PCB manufacturing facility as preferred.
69. Connect a 5V, 1A lab power supply to the P_raw5V pins on the CS (FAB-EMBL-000102) header. and check for large increases in current consumption, which can signify soldering issues.
70. Solder the direction pins on the signal conversion board (SCB, FAB-EMBL-000104) for left - right.
71. Solder the A voltage and B voltage bridges as 5V and 3.3 V respectively, thus ensuring that a 5V TTL signal from the camera when connected is scaled down to 3.3 V.
72. Connect a 7V, 1A lab power supply to the +/- pins on the SCB header. Check for large increases in current consumption, which can signify soldering issues.
73. Input a 5V voltage to the J_A1 connector and check that the output voltage on the J_A2 connector is scaled appropriately.
74. Remove the power supply.
75. To test additional unused SCB functionality, refer to the microFPGA documentation.
76. Install jumpers on the ‘1/10’ and ‘voltage & current limit’ pins on the analog conversion board (ACB, FAB-EMBL-100102) channel headers for J_A1, J_A2, J_A3, J_A4.
77. Connect a 12V, 1A lab power supply to the +/- pins on the header of the analog conversion board. Check for large increases in current consumption, which can signify soldering issues. Use an additional channel of the power supply to generate voltages in the range 0 - 10 V on pins J_A1 - J_A4 and check the scaled output on pins J_B1 - J_B4 accordingly.
78. OPTIONAL: The ACB requires additional modification relative to the base configuration supplied in the Altium project files if the builder wishes to make use of the photodiode for laser power monitoring. On the J_A4 channel, solder across R1_G and replace the resistor at R2_G with a 51 kΩ one. Note, the resistor converts the photocurrent from the photodiode to a voltage, with proportionality determined by the resistance. It may be necessary to replace this resistor when the photodiode response is calibrated in the section: OPTIONAL: Calibration of the laser power monitoring photodiode (Timing: 0.5 days), later in the protocol.
79. Remove the lab power supply.
80. Modify the microFPGA enclosure (COTS-FC-102520-GS) as per the drawings provided.
81. Mount the ACB (FAB-EMBL-000103) and SCB (FAB-EMBL-000104) to the enclosure (COTS- FC-102520-GS) base plate (FAB-EMBL-000110) using 8× M3×10 mm countersunk plastic screws, 8× M3 plastic washers, 8× M3×10mm plastic hex standoffs (female to male) and 8× M3 plastic nuts as shown in Supplementary Figure 15.
82. Mount the Br shield (COTS-BR-SHIELD) on the Au FPGA (COTS-AU-FPGA)
83. Mount the combined shield and Au FPGA on the enclosure (COTS-FC-102520-GS) base plate (FAB-EMBL-000110) using 4× M2×6mm flathead screws, 4× M2 washers, 4× M2×15mm hex standoffs and 4× M2×12mm flathead screws as shown in Supplementary Figure 15.
84. Mount the CS (FAB-EMBL-000102) onto the Br shield (COTS-BR-SHIELD)
85. Replace the connector of the power supply (COTS-GSM18E12-P1J) with the two-contact circular connector (COTS-99-0402-00-02).
86. Install the corresponding enclosure connector (COTS-09-0403-00-02) on the power supply board (Panel 3) (FAB-EMBL-000101).
87. Use a singly connectorised cable to connect the back of the power supply plug on the power supply board (Panel 3) to one of the 12 V pins on the board (P2_12V). When soldering the free-ends of the cable to the back of the power supply plug, pay attention to the polarity since the connectorised end can only be connected one way.
88. Assemble the enclosure (COTS-FC-102520-GS) side panels (FAB-EMBL-000106/7/8) and install Panels 1, 2, 3 (FAB-EMBL-000099/100/101) using 12× M3×10 mm countersunk screws, 24× M3 nuts (use half to secure the screws and the other half to act as standoffs between the enclosure and panels) and 12× M3 plastic washers as shown in Supplementary Figure 16.
89. Install the enclosure (COTS-FC-102520-GS) base plate (FAB-EMBL-000110) between the three enclosure side panels as shown in Supplementary Figure 17.
90. Install the USB-C connector (COTS-681183) in the remaining enclosure (COTS-FC-102520- GS) side panel (FAB-EMBL-000109) and connect to the USB-C connector of the microFPGA (COTS-AU-FPGA). Install the side panel on the enclosure as shown in Supplementary Figure 18.
91. Electrical connections ACB: Connect pins J1, J2, J3, J4 on Panel 2 to J_A1, J_A2, J_A3, J_A4 of the ACB.
92. Connect pins J_B1, J_B2, J_B3, J_B4 of the ACB to P_AI0, P_AI1, P_AI2, P_AI3 of the CS.
93. Electrical connections SCB: Connect pins J17 on Panel 1 to the J_A1 connector of the SCB.
94. Connect pins J_A2 on the SCB to P_Cam on the CS.
95. Electrical connections Panel 1: Connect pins P_S0 - P_S6 of the CS to pins P_S0 - P_S6 on Panel 1.
96. Connect pins P_L0 to P_L3 of the CS to pins J5 - J8 of Panel 1.
97. Connect pins P_P0 to P_P3 of the CS to pins J9 - J12 of Panel 1.
98. Electrical connections Panel 2: Connect pins J8, J9, J10, J11 of the Panel 2 to P_TTL0, P_TTL1, P_TTL2, P_TTL3 of the CS.
99. Note the P_TTL4 corresponds to the camera trigger for the passive camera mode available in the EMU GUI. If desired, connect P_TTL4 to J14 on Panel 2 and label the front panel connector TTL4.
100. Electrical connections Panel 3: Connect pins P1_12V on Panel 3 to the +/- pins on the header of the ACB.
101. Connect pins P1_7V on Panel 3 to the +/- pins on the header of the SCB.
102. Connect pins P1_5V on Panel 3 to the P_raw5V pins on the CS.
103. Install the lid (FAB-EMBL-000105) of the enclosure (COTS-FC-102520-GS) to complete the microFPGA as shown in Supplementary Figure 14.

### Configuration of the microFPGA (Timing: 2 hours)

Before the microFPGA can be used, it must be configured by uploading a configuration file to the device.

104. Install AlchitryLabs from: https://alchitry.com/alchitry-labs
105. Download the FPGA configuration binary (.bin) for the Alchitry Au from the release page of the microFPGA repository: https://github.com/mufpga/MicroFPGA/releases.
106. Connect the FPGA to the host PC.
107. Upload the configuration to the FPGA target using AlchitryLoader (installed as part of Alchitry labs).

### Setup and configuration of software and hardware for acquisition (Timing: 5 days)

Certain hardware elements require non-factory configuration for operation. For example, the piezo flexure stage controller needs to be configured in its host-software such that it provides appropriate positional feedback on the basis of the analog signal provided by the focus lock system QPD. The various piezo resonant stages used for illumination control (epi-/HILO/TIRF), back focal plane viewing and astigmatic lens positioning also require that a unique address is established for each in their host software. µManager, which ultimately provides the basis for hardware control must be installed. Communication with the various hardware devices from µManager requires that a hardware configuration is established, whereby communication with each device is defined via its device adapter (e.g., COM port etc. for serial devices). All required device adapters for the hardware elements described are available in µManager with one exception, the objective lens piezo flexure stage uses a modified version of the PI_ZStage device adapter, PI_FocusLock, a pre-compiled version of which is available on our GitHub. The htSMLM graphical user interface (GUI) is provided as a configuration for µManager’s EMU plugin. A pre-compiled version is subsequently installed and its various controls and indicators are associated with relevant hardware properties.

108. Install all manufacturer supplied software and drivers listed under: Software for software/hardware setup. Connect all hardware devices to the host PC, except for the piezo resonant stages and associated controller/distributor boards, as appropriate for the specific hardware device.
109. Test all connected hardware devices in their host software for general functionality. Return defective devices to the manufacturer for replacement.
110. Email firmware@smaract with the model and serial number of the COTS-HCU-3D stage controller to obtain firmware for ASCII communication with the device. Use SCUFirmwareUploader to upload the supplied firmware for the XY stage (COTS-SOM-12090) to allow the device to be controlled via serial commands. Connect the XY and QPD stage (COTS-2445-L) stages to the controller such that the XY stage corresponds to channels A/B and the QPD stage to channel C. Set step sizes as desired (5 µm A/B and 50 µm C step size provides sufficiently fine control of sample positioning and focusing during focus lock operation respectively). Under MENU/CONTROL MODES, set all stages to “CL” (closed-loop). Note, you may prefer to use the inverted axes configuration of the stage controller (COTS-HCU-3D) depending on personal preference. This can be carried out at any point during the protocol (MENU/KNOB CONFIG).
111. Crimp suitable length cables for the 3 - 4 piezo resonant stages using the connectors (COTS- 90327-0308) and ribbon cables (COTS-AWG 28-08G 3M) following the manufacturer guidelines. Typically, 1 m length is sufficient to position the distributor board centrally on the optical table while reaching all devices.
112. Mount the piezo resonant stage control board (COTS-ELLC2) to its base mount (FAB-EMBL- 000095) using 8× M3×12 mm hex standoffs (COTS-93655A099) (4× standoffs, then the board, then 4× standoffs on top). Mount the bus distributor board (COTS-ELLB) on top of the exposed standoff another 4× M3×12 mm hex standoffs. Secure the base to an appropriately centralized location on the optical table. Note, the boards will be covered later allowing the assembly to be located on the optical table without potential interference from the LEDs on the boards.
113. Connect the piezo resonant stage control board (COTS-ELLC2) and the bus distributor board (COTS-ELLB) and provide power to the distributor board before connecting the control board to the PC via USB.
114. OPTIONAL: Connect the four-position shared-path filter slider (COTS-ELL9) piezo resonant stages to the bus distributor (COTS-ELLB). Use the host software (Thorlabs, ELLO) to assign the device to address 3.
115. Connect the piezo resonant stages one at a time to the bus distributor (COTS-ELLB) and assign the following addresses: 0: COTS-ELL17-M, 1: COTS-ELL20-M, 2: COTS-ELL6. Note, COTS-ELL17-M should be connected last to be automatically assigned the address 0.
116. Test the piezo resonant stages in ELLO for general functionality. Return defective devices to the manufacturer for replacement.
117. Install the board cover (FAB-EMBL-000096) on the protruding 4× M3×12 mm hex standoffs from the the bus distributor board (COTS-ELLB), ensuring that the cut-outs for the various cables line up correctly and secure it in place.
118. In PIMikroMove set the parameters for the P-709.CRG controller given in Supplementary Table 7: PI MikroMove FocusLock Parameters See the manufacturer documentation for a description of these parameters, using an analog input to control the stage position, and how to write/save parameters correctly in PIMikroMove.
119. Using Windows Device Manager or other means, identify the relevant communication port for each hardware element which will use a serial protocol for communication with µManager. This includes the controller for the objective piezo focusing device (COTS-P-726.1CD), piezo resonant stages (COTS-ELL6/ELL9/ELL17-M/ELL20-M), lasers (COTS-ICHROME-MLE, COTS-IBEAM), filter wheels (COTS-FW102C), microFPGA (COTS-AU-FPGA), and XY stage (COTS-SOM-12090).
120. Ensure all software used for communication with the devices is closed.
121. Install the most recent official release of µManager 2.0.0 as necessary from https://micro-manager.org/Download_Micro-Manager_Latest_Release
122. Download the pre-compiled PI_FocusLock device adapter here: https://github.com/ries-lab/3DSMLM. Put the device adapter in the root µManager directory (e.g., C:\Program Files\micro-manager 2.0). CAUTION!: During operation, the device adapter will set some internal parameter values in the associated Physik Instrumente E-709.CRG controller.
123. Configure and save a hardware configuration in µManager including all hardware elements. See Supplementary Table 4: Micro-Manager Hardware Configuration for notes on each device including communications settings and device adapter name. TROUBLESHOOTING!: If certain hardware elements fail to connect, try extending the timeout periods before reloading the hardware configuration. If either the booster or focus lock laser fail to connect, ensure that their interlock circuits are closed.
124. Establish several µManager configuration presets following Table Supplementary Table 8: µManager Configuration Presets
125. Turn on metadata generation by µManager by navigating to Tools from the ribbon menu, then Options and ensure ‘Create metadata.txt file with Image Stack Files’ is ticked.
126. Download the pre-compiled htSMLM GUI from https://github.com/ries-lab/3DSMLM and place the .jar file in the EMU folder in the root µManager directory (e.g., C:\Program Files\micro-manager 2.0\EMU).
127. Open µManager and load the saved hardware configuration file. Ensure that all hardware devices are detected correctly.
128. Launch the htSMLM GUI by navigating to Plugins/Interface/EMU and then selecting the compiled htSMLM UI (.jar file).
129. In the htSMLM EMU plugin, configure the GUI using the Configuration wizard (Configuration/Modify configuration) as described in Supplementary Table 9. htSMLM EMU Configuration - GUI Settings. Subsequently, associate the GUI controls and indicators with the respective device property as given in Supplementary Table 10. htSMLM EMU Configuration - Hardware Bindings. In the case of a system using both the multi-mode and single-mode laser engines, created individually configured htSMLM configurations for each with the corresponding parameters.
130. Please see: https://github.com/jdeschamps/htSMLM/tree/main/guide for a guide to the various panels of the EMU htSMLM GUI and their functions.

### Testing of the microFPGA (Timing: 1 day)

Before the microFPGA can be connected to hardware, its correct function should be verified by presenting either test signals or software commands and measuring the response.

131. Power the microFPGA with the fixed 12V, 1A power supply
132. Use the lab power supply to provide voltages in the range 0 - 10 V on the microFPGA analog inputs for the QPD. Verify in the µManager Device Property Browser that the correct voltage is read out by the FPGA. Note that the analog conversion board scales the voltage to 0 - 1 V and the FPGA returns a 16-bit value (0 - 65535) for the range 0 - 1 V.
133. Change the state of the TTL channels in the µManager Device Property Browser and check with an oscilloscope that the correct voltage is read out from the front panel connectors (J8 - J11 on Panel 2).
134. Change the state of the PWM channels in the µManager Device Property Browser and check with an oscilloscope that the correct PWM sequence is read out from the front panel connectors (DO:L0-3:POW).
135. In the htSMLM GUI set the trigger mode for all laser lines to ‘Camera’, the sequences to 65535 (1111111111111111) and the FPGA synchronization mode to active camera.
136. Connect the Timing 1 signal from the sCMOS camera to an oscilloscope. Set the exposure time to 1 ms and set the camera to acquire images continuously. Check that this produces a square wave signal on the oscilloscope.
137. Connect the Timing 1 signal from the sCMOS camera to the camera exposure input of the microFPGA (DI:CAM:EX). Using the oscilloscope, check that all laser trigger channels (DO:L0- 3-EN) follow the signal.

### Making electrical connections to hardware (Timing: 0.5 days)

Proper control of the various hardware elements by the microFPGA requires that the appropriate connections are made between its I/O and those on the associated hardware including the camera, lasers/laser engine, and laser power monitoring photodiode. Connection to the QPD amplifier is omitted at this stage and follows later in the protocol, when it is required.

138. For active camera synchronization, connect the camera Timing 1 output SMA connector to the DI:CAM:EX SMB connector on the microFPGA.
139. For passive camera synchronization, connect the camera trigger input SMA connector (Ext. Trig.) to TTL4 on Panel 2 of the microFPGA.
140. Connect the laser engine ‘enable’ SMB connectors (DO:L0-3:EN) to their equivalent SMB connector on the Toptica iChrome MLE laser engine (if present in the system). Note that the laser engine defines the laser lines in reverse order of wavelength. For example, for the standard 405/488/561/640 nm configuration, 640 nm is laser 1 and 405 nm is laser 4.
141. Connect the D-SUB9 connector of the Laser Engine panel of the microFPGA to the multi-mode laser engine control box as appropriate. The D-SUB9 connector duplicates the signals DO:L0- 3:EN and DO:L0-3:POW).
142. OPTIONAL: If wishing to make use of the laser power monitoring photodiode. Connect the photodiode BNC output connector to the SMB AI:PD connector on the microFPGA. Route the cable out of the body using an unused cable channel in the body lower plate (FAB-EMBL- 000001).

### Setup of the infinite conjugate viewing camera and infinite conjugate transmission target (Timing: 0.5 days)

The camera and lens must be spaced appropriately to provide a reference to an object viewed at infinite distance. To do so, the camera and lens assembly should be taken to an appropriate location and focused to image a distant object. Once the infinite conjugate viewing camera is setup correctly, it can be unified with the infinite conjugate transmission target, which is then brought into focus on the camera by varying the distance between the target and its lens. Since the camera is setup to image an infinite conjugate object, this ensures that the lens is positioned to produce an image of the target at infinity.

143. Assemble the infinite conjugate viewing camera and infinite conjugate transmission target (3D- SMLM-EMBL-004), omitting the Ill24 mm round coverslip (Supplementary Figure 19).
144. Uncouple the infinite conjugate viewing camera from the infinite conjugate transmission target by removing the lens tube coupler (COTS-SM1S10).
145. Take the infinite conjugate viewing camera to an appropriate location to image an object > 350 m away. Typically, a window of a building or a remote location is required. The rationale for choosing this distance is described in Supplementary Note 9: Specifying an infinite conjugate source.
146. Loosen a lock ring (COTS-SM1NT) of the lens tube coupler (COTS-SM1T10, CAD configuration: Infinity Camera) and focus the camera on the distant object by threading the adjoining lens tube (COTS-SM1L35, on either side of the coupler) inwards or outwards. Once a maximally sharp image has been produced, lock the position using the lock ring(s). Check the image sharpness again after securing.
147. Produce a transmission target by taking a Ill24 mm round coverslip and repeatedly flicking the tip of a permanent marker in close proximity to the coverslip such that a spray of fine ink particles is produced thereupon.
148. Install the coverslip transmission target in the lens tube (COTS-SM1L03) of the infinite conjugate viewing target, securing it in place with 2× retaining rings (COTS-SM1RR).
149. Reunify the infinite conjugate viewing camera and the infinite conjugate transmission target by reinstalling the lens tube coupler (COTS-SM1S10).
150. Backlight the target with a white light source.
151. Loosen the lock ring (COTS-SM1NT) of the lens tube coupler (COTS-SM1T10, on the infinite conjugate transmission target side) and focus the image of the target on the camera by threading the adjoining lens tube (COTS-SM1L15 or COTS-SM1M15) inwards or outwards. Once a maximally sharp image has been produced, lock the position using the lock ring(s). Check the image sharpness again after securing.
152. Uncouple the infinite conjugate viewing camera from the infinite conjugate transmission target by unthreading the lens tube coupler (COTS-SM1S10) on the camera side.

### Emission path beam routing (Timing: 4 days)

A CAD rendering of the completed emission path is shown in Figure 4. To establish the route of the emission path, an alignment laser is aligned to the objective lens port to indicate the optical axis of the emission path. The image splitter platform and a pair of pinned alignment targets are first used as a guide to ensure proper reflection of light out of the microscope body. The image splitter platform is subsequently installed in place for imaging operation and the beam split by a dichroic mirror. The two beamlets are subsequently steered to route the corresponding image channels onto the camera with appropriate spatial separation using an alignment tool. This tool provides a reference position where the two beams from the reflected and transmitted pathway should cross each other owing to a small tilt of the two images with respect to the camera. The displacement of the image planes from the camera chip is small with respect to the depth of focus and so does not affect the optical performance across the field of view.

**Figure 4.**
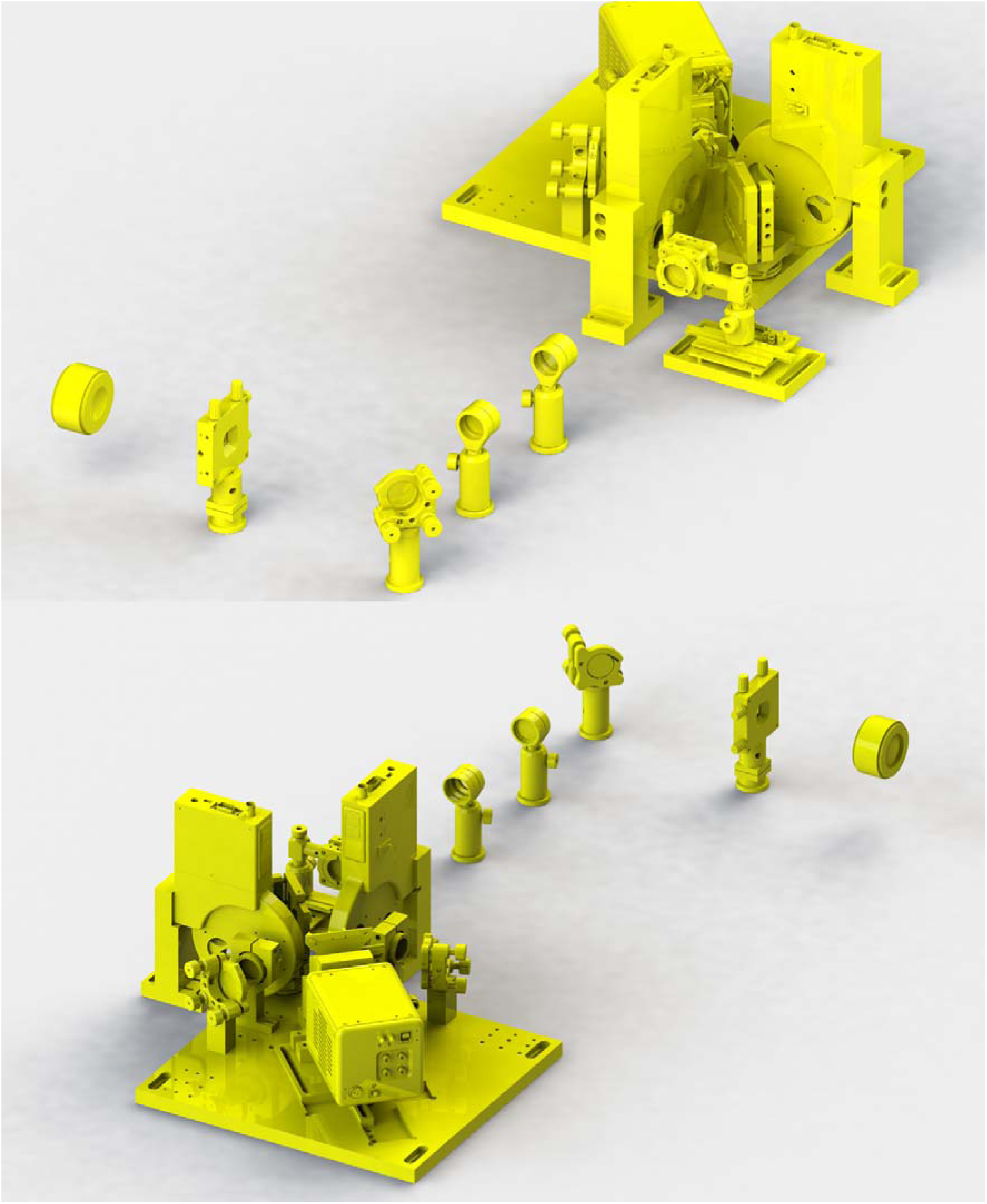
The fully assembled multi-color, 3D imaging capable emission path. Top: as viewed from the front/right of the microscope. From left-right: the tube lens, field aperture, first steering mirror, first relay lens, laser blocking filters, astigmatic lens and filter wheels either side of the image splitter dichroic mirror. Bottom: as viewed from the back/left of the microscope highlighting the image splitter platform. Following the filter wheel, the transmitted and reflected paths each comprise a second relay lens and steering mirror before the images are recombined by a knife-edge mirror (occluded in figure) and imaged onto an sCMOS camera.

153. Remove all covers (FAB-EMBL-000013/14/15/16, 4× FAB-EMBL-000018, FAB-EMBL-000019/20/21/66) from the microscope.
154. Install the pre-aligned 532 nm alignment laser assembly (3D-SMLM-EMBL-007) in the RMS thread of the objective port (FAB-EMBL-000043).
155. Introduce the image splitter platform (FAB-EMBL-000077) (omitting all mounted components) as indicated in Supplementary Figure 20. Ensure that the side facing the right side of the microscope body sits flush against the body lower plate (FAB-EMBL-000001).
156. Install two alignment targets (FAB-EMBL-000081) with two irises (COTS-SM1D12D) at positions spaced as far apart as possible along the emission path axis (the alignment targets should be placed in the transmitted pathway of the image splitter as shown in Supplementary Figure 20).
157. Using the alignment laser as a guide, translate the image splitter platform (FAB-EMBL-000077) along the edge of the body lower plate (FAB-EMBL-000001) such that the beam reflected by the emission path dichroic mirror (COTS-25.5×36x3MM-DICHROIC-EM) aligns laterally with the distal iris before securing the platform in position.
158. Check if the alignment laser passes the proximal iris in the lateral direction. TROUBLESHOOTING!: Depending on tolerance stack up, seating of the emission path dichroic mirror in its mount (FAB-EMBL-000025) and flatness of the levelled objective piezo platform (FAB-EMBL-000038), it is possible that the beam will not pass the proximal iris perfectly on-axis. Such lateral misalignment is not deleterious to system performance but makes assessment of vertical alignment more difficult. If this is the case, loosen the screws securing the image splitter platform thus allowing the irises to be individually aligned to the beam by sliding the image splitter platform during subsequent alignment steps using the body lower plate (FAB- EMBL-000001) as a linear guide.
159. Check that the laser passes correctly through the proximal and distal irises in the vertical direction repositioning the splitter platform as described above as necessary. TROUBLESHOOTING!: If the beam does not pass through the vertical centers of each iris, first ensure that the actuator screws (FAB-EMBL-000077) are seated correctly in contact with the three pairs of contact pins to center the objective piezo platform (FAB-EMBL-000038) with respect to the clear aperture through the microscope body and check that the alignment laser emission is well-centered on the two irises of its assembly (3D-SMLM-EMBL-007). If the issue persists the likely cause is incorrect seating of the dichroic mirror in its mount (FAB-EMBL- 000025). Depending on the adhesive used, one may reinstall the optic and check whether this improves alignment. If these steps do not rectify the issue, the dichroic mount includes features allowing for vertical realignment. Remove the mounted emission dichroic (COTS- 25.5×36x3MM-DICHROIC-EM/FAB-EMBL-000025) from the optic pillar (FAB-EMBL- 000023), remove the lower/center dowel pin from the emission dichroic mount (FAB-EMBL- 000025) and reinstall the mount onto the optical pillar (FAB-EMBL-000023) with just one dowel pin. The rotational axis of the remaining dowel pin is centered on the reflective face of the emission dichroic and the clearance holes for the M3×35 mm socket head screws are loosely specified to allow a small range of rotation. Loosen the screws just enough to allow rotation under light force and rotate the emission dichroic mount until the beam passes as closely as possible through the center of the distal iris target (FAB-EMBL-000081/COTS-SM1D12D) on the image splitter platform (FAB-EMBL-000077). Tighten the screws once this condition is achieved. The beam should pass the proximal iris to within ± 0.5 mm. If this condition is not met, repeat the procedure.
160. Remove the image splitter platform (FAB-EMBL-000077) but leave the alignment targets in place.
161. Install the lower section of the kinematic magnetic mount (COTS-SB1-M) on the image splitter platform (FAB-EMBL-000077). Note, the rotational orientation should be set such that when the upper section is installed, its release lever should be accessible so check the resulting orientation of the upper section and secure the lower section accordingly.
162. Install the knife-edge mirror (COTS-MRAK25-P01) secured with the clamp (COTS-PM3-M) on its pedestal (FAB-EMBL-000075) as well as the camera image alignment target (FAB-EMBL- 000093, COTS-SM1D12D, COTS-SM1L10) and the steering mirror pedestals FAB-EMBL- 000079/FAB-EMBL-000080) on the image splitter platform (FAB-EMBL-000077).
163. Install the image splitter platform (FAB-EMBL-000077) on the optical table as shown in Supplementary Figure 21.
164. Place the first emission path steering mirror (COTS-RS2P4M, COTS-RS5M, COTS-POLARIS- K1, COTS-BB1-E02) such that the alignment laser beam is centered on the mirror. Position and steer the mirror such that the beam remains centered on the mirror and passes the two alignment targets (FAB-EMBL-000081) on the image splitter platform (FAB-EMBL-000077) through the irises (COTS-SM1D12D). Secure the mirror in place using a pedestal post clamp (COTS-PS-F).
165. Pre-configure the actuators of the two steering mirror mounts (COTS-POLARIS-K1) for the transmitted and reflected paths to nominal positions using a ⅛” post height spacer (3D-SMLM- PS-0.125) inserted between the steerable and mounting sections of the steering mirror mount and adjusting each actuator in an iterative manner. This ensures that the mirrors are centered with regard to the system optical axis when installed later. Note, the later addition of the splitting dichroic (COTS-25.5×36x3MM-DICHROIC-SP) elicits a ca. 1 mm optical axis shift (for a 3 mm thick dichroic) resulting in only a minor path length difference between the two arms which is adjusted for by later alignment of the steering mirror in the transmitted path.
166. Remove the alignment targets (FAB-EMBL-000081) and install the steering mirror for the transmitted path (COTS-POLARIS-K1, COTS-BB1-E02) on its pedestal (FAB-EMBL-000080, Supplementary Figure 22). The beam should reflect from the mounted mirror onto the subsequent knife-edge mirror. Steer the mirror such that the reflected beam passes the camera image alignment target (FAB-EMBL-000093) through the iris (COTS-SM1D12D).
167. Glue the splitting dichroic mirror (COTS-25.5×36x3MM-DICHROIC-SP) into its mount (FAB- EMBL-000069) as described previously and install the mount into the steerable kinematic mount (COTS-KM200S). See Supplementary Note 2.
168. Replace the 2× pre-installed actuators of the steerable kinematic mount (COTS-KM200S) with the alternative 2× hex-drive actuators (COTS-F25SS050)
169. Install the two alignment targets (FAB-EMBL-000081) with irises (COTS-SM1D12D) as far apart as possible in two available positions of the reflected (angled) pathway of the image splitter platform (FAB-EMBL-0000077).
170. Install the upper section of the kinematic magnetic mount (COTS-SB1-M) and place the mounting plate for the image splitter dichroic kinematic mount (FAB-EMBL-000068) on the top. Since the orientation of the mounting plate will determine the initial orientation of the splitter dichroic, secure the mounting plate loosely such that the upper plate can be rotated by hand on the magnetic mount under light pressure.
171. Use a suitable machined flat component as a guide to orient the mounting plate for the splitter dichroic kinematic mount (FAB-EMBL-000068) relative to the underlying image splitter platform (FAB-EMBL-0000077). When correctly positioned the sides of the mounting plate and splitter platform facing away from the reflected pathway (closest to the optical table edge) should be flush with each other. Secure in place when a correct orientation is achieved.
172. Install the steerable kinematic mount (COTS-KM200S) housing the dichroic (COTS- 25.5×36x3MM-DICHROIC-SP)/FAB-EMBL-000069 onto the mounting plate for the image splitter dichroic kinematic mount (FAB-EMBL-000068) ensuring that the rear side of the mount is flush with the locating surface.
173. Use the hex adjustment screws of the steerable mirror mount (COTS-F25SS050, COTS- KM200S) to reflect the incident beam through the irises (COTS-SM1D12D) of the two alignment targets (FAB-EMBL-000081), optimizing preferentially for the distal target. The beam should pass the proximal iris to within ± 0.5 mm. The assembly should now look as shown in Supplementary Figure 23.
174. Remove the alignment targets (FAB-EMBL-000081) and install the steering mirror (COTS- POLARIS-K1, COTS-BB1-E02) for the reflected path on its pedestal (FAB-EMBL-000079) following the process for the transmitted path. Use the steering mirror adjusters to align the beam to the camera image alignment target (FAB-EMBL-000093) as previously described (Supplementary Figure 24).

### Emission path alignment (Timing: 5 days)

Having routed the beam through the emission path, the tube lens and relay lens pair must be positioned to produce primary and secondary images of the objective focal plane (the design focal plane for which the objective produces a conjugate image at infinity). The tube lens mount is aligned to the optical axis using an alignment assembly and positioned for telecentricity by passing the alignment laser through the tube lens (to produce a focus at the back focal plane) and subsequently collimating the beam with the objective lens. The objective and tube lens thus provide a 4f imaging system yielding a ca. 92× magnification primary image. Spacing and alignment of the first relay lens is similarly achieved by using the tube lens to generate a focus from an infinite conjugate source and collimating with the relay lens accordingly. To aid alignment, the second relay lenses of the reflected and transmitted paths of the image splitter feature mounts with circumscribed degrees of freedom: a dowel pin slot and a pair of dowel pins in the base allow lateral translation but not reorientation of the mount. The tube lens is removed from its base adapter to produce a focus from the first relay lens thus again presenting the longitudinal positioning of the second relay lens as a collimation operation. Once correctly positioned, the emission path and image splitter are set up to produce two secondary images on the camera with ca. 61× magnification. The addition of blocking filters for the laser lines used for illumination/focus lock and various emission filters allows for fluorescence images to be collected in subsequent steps. Initially, the camera is installed in a nominal position and then focused using a reference to an infinite conjugate object.

175. Remove the upper section of the kinematic magnetic mount (COTS-SB1-M) with dichroic mirror and associated components (COTS-25.5×36x3MM-DICHROIC-SP, COTS-KM200S, FAB-EMBL-000068/69). Install one alignment target with iris (FAB-EMBL-000081, COTS- SM1D12D) as shown in Supplementary Figure 25. The alignment laser should be aligned to the target. TROUBLESHOOTING! If the beam is no longer aligned with the target, return to step 155 to check alignment out of the body and repeat subsequent alignment steps.
176. Install the tube lens alignment target (3D-SMLM-EMBL-002) in the tube lens mount (FAB- EMBL-000035), securing it with the retaining cap (FAB-EMBL-000036) as shown in Supplementary Figure 26.
177. Position the tube lens mount on its base (FAB-EMBL-000034/35/36) such that the laser passes through the ground-glass alignment target and subsequent iris. Note, the base (FAB-EMBL- 000034) should remain loosely affixed to the body lower plate (FAB-EMBL-000001) during this operation. Firmly secure the base in place.
178. Remove the tube lens alignment target (3D-SMLM-EMBL-002) and install the tube lens (COTS- TTL165-A) at an arbitrary position along the axis of the mount (FAB-EMBL-000035) without securing the retaining cap (FAB-EMBL-000036) such that the lens may be moved back and forth. CAUTION: The tube lens has a correct orientation with the infinite conjugate (objective lens) side indicated by the arrow on the housing. CAUTION: Take care not to touch optical surfaces of the tube lens.
179. Check that the beam still passes the iris target on the image splitter platform within an allowable margin of ± 0.5 mm. TROUBLESHOOTING!: If the lateral beam walk-off is > ± 0.5 mm, the tube lens mount should be realigned. If the problem persists, secure a component with a flat side against the tube lens mount to define the angular alignment and reposition the tube lens mount along this axis such that the laser passes the target in the horizontal direction. If the vertical beam walk-off is > ± 0.5 mm and all previous alignment steps have been completed correctly it is possible that the tube lens is mounted at an incorrect height. Remove the tube lens mount (FAB- EMBL-000035) from the tube lens base adapter (FAB-EMBL-000034). Remove the tube lens (COTS-TTL165-A). Install a second alignment target with iris (FAB-EMBL-000081/COTS- SM1D12D) in the transmitted path of the splitter platform and check the beam height on the two irises. Return to step 176 if the beam does not pass both irises to within ± 0.5 mm. If the beam passes the irises then the tube lens height may be at fault. Check the centration of the lens in its housing by reinstalling the tube lens on its mount (COTS-TTL165-A/FAB-EMBL-000035) and rotating the lens. If the laser spot moves noticeably between several rotational positions of the tube lens, the lens elements may not be well centered in their housing and should be exchanged with the manufacturer. Otherwise, the tube lens may not be mounted at the correct height. Check the dimensions of the tube lens mounting components (FAB-EMBL-000034/35/36) and remachine/shim as necessary.
180. Remove the alignment laser assembly (3D-SMLM-EMBL-002) from the objective port (FAB- EMBL-000043) and replace it with the objective port alignment target assembly (3D-SMLM- EMBL-006) with the beam profiling camera installed on the distal end.
181. Use the pitch/yaw (COTS-KAD11F) and XY (COTS-CXY1) mounts of the alignment laser assembly to align through the two irises (COTS-SM1D12D) before removing the 2× slotted lens tubes (SM1L30C), 2× irises (COTS-SM1D12D), lens tube (COTS-SM1S10) and thread adapter (COTS-SM1A4). Install the thread adapter (COTS-SM1A2) into the exposed SM1 port of the XY mount and install the assembly onto the tube lens (COTS-TTL165-A) via the SM2 internal thread of the thread adapter (see CAD 2. Emission Path Build-up. F. Tube Lens Focusing, Supplementary Figure 27).
182. Use the pitch/yaw (COTS-KAD11F) and XY (COTS-CXY1) mounts of the alignment laser assembly to align through the two irises (COTS-SM1D12D) of the objective port alignment target assembly (3D-SMLM-EMBL-006).
183. Remove the objective port alignment target assembly (3D-SMLM-EMBL-006) and replace it with the objective lens.
184. CAUTION! When illuminating the tube lens with the alignment laser, a small and divergent beam will exit the objective lens and even small misalignments may result in a strong deviation from the optical axis and constitute a laser safety hazard accordingly. Wear laser safety eyewear whenever assessing alignment through the objective lens. If the laser is launched substantially off-axis, use the pitch/yaw (COTS-KAD11F) and XY (COTS-CXY1) mounts of the alignment laser assembly to approximately center the output with respect to the objective lens such that the beam emerges approximately on-axis.
185. Move the tube lens (COTS-TTL165-A) along the axis of its mount to collimate the beam by minimizing the laser spot size on the ceiling or optical table enclosure. Secure the tube lens in place with the retaining cap (FAB-EMBL-000036).
186. Remove the alignment laser assembly (3D-SMLM-EMBL-002) from the tube lens and the objective lens from the objective port (FAB-EMBL-000043). Reinstall the 2× slotted lens tubes (SM1L30C), 2× irises (COTS-SM1D12D), lens tube (COTS-SM1S10) and thread adapter (COTS-SM1A4) of the alignment laser assembly.
187. Install the four covers (FAB-EMBL-000013/14/15/16) on the body lower standoffs (FAB- EMBL-000002/3/4/5) using 8× M4×6 mm socket head screws and taking care to install the correct cover on each side of the microscope body. The covers on the side of the body where the illumination, emission, and focus lock paths are coupled in/out should have their aperture centers at 110.4, 80 and 48.4 mm from the table surface respectively.
188. Reinstall the alignment laser assembly (3D-SMLM-EMBL-002) in the objective port (FAB- EMBL-000043) and realign through the two irises.
189. Adjust for minor walk-off resulting from installation of the tube lens by using the first steering mirror (COTS-POLARIS-K1/COTS-BB1-E02) of the emission path to align the beam to the alignment target (FAB-EMBL-00081) installed in the transmitted path of the image splitter platform (FAB-EMBL-000077).
190. Set up the shear plate (3D-SMLM-EMBL-005) just upstream of the image splitter platform (FAB-EMBL-000077) such that the diverging alignment laser is incident on it.
191. Introduce and align the first relay lens (COTS-AC254-300-A-ML, COTS-LMR1-M, COTS- PH50E-M, COTS-TRA30-M) to collimate the alignment laser focused by the tube lens using the back reflection method (see Orienting and aligning optical elements) with the installed alignment target (FAB-EMBL-000081, COTS-SM1D12D) as the downstream target and an additional target placed upstream of the lens (e.g., 3D-SMLM-EMBL-003, adjusted to the correct beam height). The relay lens has a correct orientation with the infinite conjugate (collimated) side indicated by the arrow on the housing. If the alignment is performed correctly the associated walk off should be negligible for the two pathways. It will be necessary to move the shear plate (3D-SMLM-EMBL-005) in and out of the beam path to assess collimation and alignment individually.
192. Remove the shear plate assembly (3D-SMLM-EMBL-005).
193. To adjust for minor walk-off, use the second steering mirror for the transmitted path (COTS- POLARIS-K1/COTS-BB1-E02) to align the beam to the camera alignment target (FAB-EMBL- 000093).
194. Replace the upper section of the kinematic magnetic mount (COTS-SB1-M) with its dichroic mirror and associated components (COTS-25.5×36x3MM-DICHROIC-SP, COTS-KM200S, FAB-EMBL-000068/69).
195. Remove the alignment target with iris (FAB-EMBL-000081, COTS-SM1D12D) from the transmitted pathway of the image splitter platform (FAB-EMBL-000077) and reinstall as far downstream as possible in the reflected pathway.
196. Adjust the alignment of the reflected alignment laser to the alignment target (FAB-EMBL- 000081, COTS-SM1D12D) on the image splitter platform (FAB-EMBL-000077) using the steerable kinematic mount for the image splitter dichroic mirror mount (COTS-KM200S) and the second steering mirror (COTS-POLARIS-K1/ COTS-BB1-E02) to align the beam to the camera alignment target (FAB-EMBL-000093).
197. Remove the alignment target with iris (FAB-EMBL-000081, COTS-SM1D12D) from the reflected pathway of the image splitter platform (FAB-EMBl-000077) and reinstall it as far downstream as possible in the transmitted pathway.
198. Install the second relay lens mounts (FAB-EMBL-000078) in the transmitted and reflected pathways. Loosely secure with the screws just enough that the mount can still be translated along the axis of the dowel pin slot.
199. Install the second relay lenses (COTS-AC254-200-A) approximately half way down the length of their lens tubes (COTS-SM1M10) using the two retaining rings (COTS-SM1RR) to secure them place. CAUTION: the relay lenses have a correct orientation with the finite conjugate side (towards the camera) facing the side of least curvature (flattest). Indicate the orientation on the lens tube with a permanent marker with an arrow pointing away from the side of least curvature (towards the infinite conjugate).
200. Insert the second relay lenses mounted in lens tubes (COTS-AC254-200-A, COTS-SM1M10) into the second relay lens mounts (FAB-EMBL-000078) in the reflected and transmitted paths with the arrow pointing towards the image splitter dichroic assembly. Secure the lens tubes in an arbitrary longitudinal position by gently closing the split clamp of the respective second relay lens mount. CAUTION: too much pressure could deform the lens tube and the lens contained therein, tighten just enough to prevent motion of the lens. Ensure that the lens tube is correctly centered in the mount with three points of contact. The assembly should now appear as shown in Supplementary Figure 28.
201. Laterally translate the mounted lens in the reflected path to align the beam to the iris of the camera image alignment target (FAB-EMBL-000093/COTS-SM1D12D) and secure in place. Depending on previous alignment steps, some vertical walk off is possible. This will be corrected when viewing the beam foci on the camera.
202. Repeat the alignment process (step 201) for the transmitted path lens. Alignment of the transmitted path is more challenging since only a fraction of a percent of the laser power will be transmitted by the long pass image splitter dichroic mirror (COTS-25.5×36x3MM-DICHROIC- SP). Nevertheless, it should be possible to translate and align the lens as above. CAUTION! Do not remove the dichroic to increase the visibility of the alignment laser before attempting alignment since replacement of the dichroic will shift the beam.
203. Remove the camera image alignment target (FAB-EMBL-000093) and release the second relay lens tubes (COTS-SM1L10) by opening the split clamp on the second relay lens mount (FAB- EMBL-000078).
204. Remove the mounted emission tube lens (FAB-EMBL-000035/36, COTS-TTL165-A) from its base adapter as a single piece. CAUTION: do not remove the base (FAB-EMBL-000034) that is affixed to the microscope body lower plate (FAB-EMBL-000001) nor the tube lens retaining cap (FAB-EMBL-000036) from the tube lens mount (FAB-EMBL-000035). Removing the tube lens assembly correctly will allow it to be replaced later without requiring realignment. The collimated beam from the alignment laser should now be focused by the first relay lens and approximately collimated by the second relay lens of the transmitted path.
205. Set up the shear plate (3D-SMLM-EMBL-005) to assess the beam collimation for the transmitted pathway by placing the assembly downstream of the knife-edge mirror. Slide the lens tube housing of the second relay lens of the reflected path (COTS-AC254-200-A, COTS- SM1L10) to collimate the beam. TROUBLESHOOTING! If the position of the second relay lens to achieve collimation is substantially shifted relative to the nominal position, problems may arise. First, if the lens is placed too far upstream, it may occupy the space required to later install the filter wheel. Second, if the lens is placed too far downstream, the lens tube (COTS-SM1L10) in which it is mounted may result in vignetting of the image and/or spatial variations in PSF shape owing to blocking of the reflected light from the subsequent steering mirror. If either of these cases are apparent, it is necessary to adjust the position of the entire image splitter platform (FAB-EMBL-000077) to lie further upstream or downstream respectively. For small adjustments (up to 25 mm), one may reposition the image splitter platform without compromising the alignment. In this case, use two fixed kinematic stops (COTS-KL01L) to indicate the position of the side of the platform running parallel to the transmitted path optical axis. The platform can then be adjusted to the correct position along the defined axis before being fixed in position. Check the alignment of the laser through the splitter platform using the various alignment targets (FAB-EMBL-000081/93 as necessary). For larger adjustments (> 25 mm), it is advisable to follow the above procedure but return to step 163 to realign the emission path more thoroughly thereafter.
206. Repeat the process of collimating the beam using the second relay lens (COTS-AC254-200-A, COTS-SM1L10) for the transmitted path. Note, if the beam is too dim to judge collimation, remove the upper section of the kinematic magnetic mount (COTS-SB1-M) with its dichroic mirror and associated components (COTS-25.5×36x3MM-DICHROIC-SP, COTS-KM200S, FAB-EMBL-000068/69) and replace them once collimation has been achieved.
207. Install the Bertrand lens assembly (COTS-#32-963, COTS-SM1AD20) in its piezo resonant slider (COTS-ELL6).
208. Install the Bertrand lens piezo resonant slider (COTS-ELL6) on the Bertrand lens piezo resonant slider mount (FAB-EMBL-000074) and in turn install that on the camera base plate (FAB- EMBL-000077).
209. Adjust the longitudinal position of the Bertrand lens (COTS-#32-963, COTS-SM1AD20) approximately to the nominal position and lightly secure the lens in place with the locking ring (COTS-SM1NT).
210. Switch the Bertrand lens piezo resonant slider (COTS-ELL6) to the open position.
211. Mount the sCMOS camera (COTS-C15440-20UP) to the camera mounting plate (FAB-EMBL- 000071) via the camera c-mount fastener (FAB-EMBL-000073).
212. Install the mounted sCMOS camera (COTS-C15440-20UP) onto the camera base plate (FAB- EMBL-000074). The assembly should appear as shown in Supplementary Figure 29.
213. Rotate the camera such that its upper side faces up by loosening the camera c-mount fastener (FAB-EMBL-000073) and tightening once the desired orientation has been achieved. With the camera base plate (FAB-EMBL-000074) secured to the optical table, the camera can be rotated to a precise orientation using a bubble level (COTS-LVL01) placed on top. The rationale for choosing this orientation is discussed in Supplementary Note 10: Correct orientation of the sCMOS camera.
214. Remove the alignment laser assembly (3D-SMLM-EMBL-002) from the objective port (FAB- EMBL-000043) and replace it with the infinite conjugate transmission target (3D-SMLM- EMBL-004) using an RMS to SM1 thread adapter (COTS-SM1A4) to couple the lens tube (COTS-SM1S10) to the port (Supplementary Figure 33).
215. Install the sCMOS camera/Bertrand lens assembly to the nominal position on the image splitter platform (FAB-EMBL-000077) and secure it in place as shown in Supplementary Figure 30. The camera base plate (FAB-EMBL-000074) should sit flush against one edge of the image splitter platform. Consult the CAD assembly for guidance.
216. Backlight the target with a white light source and image the target on the sCMOS camera, adjusting exposure time appropriately for the supplied light source to not saturate the camera. Note, that the magnification from the target to the camera is 1.1×, thus delivering a view of the central ca. 13.6 × 13.6 mm of the target (for the specified Hamamatsu Orca Fusion BT camera).
217. Loosen the screws securing the camera base plate (FAB-EMBL-000074) to the image splitter platform (FAB-EMBL-000077).
218. Observe the target on the camera and focus the camera by translating the camera base plate (FAB-EMBL-000074) against its linear guiding surface of the image splitter platform (FAB- EMBL-000077, see CAD assemblies) such that the target appears maximally in focus. Fine adjustment can be performed by installing a ball-tipped kinematic adjuster (COTS-KL02L) on the image splitter platform as needed. Secure the camera base plate in place once a fine focus has been achieved. The camera is now positioned to sharply focus the infinite conjugate image produced by the objective lens.
219. Install an effective ND 6 filter (COTS-NE20A-A, COTS-NE40A-A) onto the accessible internal SM1 thread of the first relay lens (COTS-AC254-300-A-ML) to near-fully attenuate the alignment laser beam.
220. Remove the infinite conjugate transmission target (3D-SMLM-EMBL-004) from the objective port (FAB-EMBL-000043) and replace it with the alignment laser assembly (3D-SMLM-EMBL- 002).
221. Preview the image from the sCMOS camera (1 ms exposure time) and decrease the total neutral density until the laser spot can be seen on the camera for the reflected path (the transmitted beam will be dimmer).
222. Use the image display and the second steering mirror (COTS-POLARIS-K1, COTS-BB1-E02) of the reflected path to steer the beam center to the X,Y pixel positions given in Supplementary Table 11. Field of View Center Positions on Camera.
223. Appropriately block the reflected path using a laser safety screen (COTS-TPSM1-M).
224. Decrease the neutral density further until the laser spot can be seen on the camera image for the transmitted path.
225. Use the second steering mirror (COTS-POLARIS-K1, COTS-BB1-E02) of the transmitted path to steer the beam center to the xy pixel positions given in Supplementary Table 11. Field of View Center Positions on Camera.
226. Remove the alignment laser assembly (3D-SMLM-EMBL-002) from the objective port (FAB- EMBL-000043) and replace it with the objective lens.
227. Install the blocking filters for the excitation and focus lock lasers (2× COTS-25MM-HOUSED- FILTER, COTS-SM1L03, 1× COTS-LMR1-M, COTS-PH50E-M, COTS-TRA30-M) just downstream of the first relay lens (AC254-300-A-ML), avoiding locations that will block later installation of other components and secure using a pedestal post clamp (COTS-PS-F) as shown in Supplementary Figure 31. See Supplementary Note 2: Selecting spectrally discriminative optics.
228. Install all filters in the transmitted and reflected path filter wheels (COTS-FW102C), ensuring that each is installed in the correct positions for the GUI control bindings (or amend these bindings appropriately). Ensure that the filters are correctly oriented when the filter wheels (check with the filter manufacturer for the correct orientation of the arrow/caret relative to the direction of light propagation). See Supplementary Note 2: Selecting spectrally discriminative optics.
229. Install the filter wheels (COTS-FW102C) in their mounts (FAB-EMBL-000076/000111) and install the mounted filter wheels in position ensuring that the correct wheel is placed in each path. The assembly should now appear as shown in Supplementary Figure 32. TROUBLESHOOTING! If the image splitter platform has been moved from its nominal longitudinal position to account for tolerances in lens focal lengths, the filter wheel mount bases (FAB-EMBL-000111) may not align to the optical table holes when the filter wheel aperture is aligned to the emission path when the upper part of the mount (FAB-EMBL-000076) is affixed to the base in the nominal position shown in Supplementary Figure 32. The position can be adjusted by releasing the mount from its base and securing in an offset position.

### Basic epi-illumination alignment (Timing: 4 days)

To aid initial focusing and quality control of the microscope in the following section, the illumination path is set up first for simple basic epi-illumination without the additional advanced features of homogenization/TIRF/HILO/variable field size that will follow. Nevertheless, it is necessary to consider some aspects of these features *en route*. For example, to ensure proper operation of the refractive beam shaper, the beam(s) from the single mode sources need to be carefully expanded to a diameter of 6 mm and subsequently routed through the illumination path to the objective port using various alignment targets. The illumination tube lens is subsequently aligned to the illumination path optical axis and focused to achieve a collimated output from the objective lens. A CAD rendering of the fully assembled illumination path including optional components (e.g., booster laser, multi-mode laser path and associated parts) and those that will be installed in later sections is shown in Figure 5.

**Figure 5.**
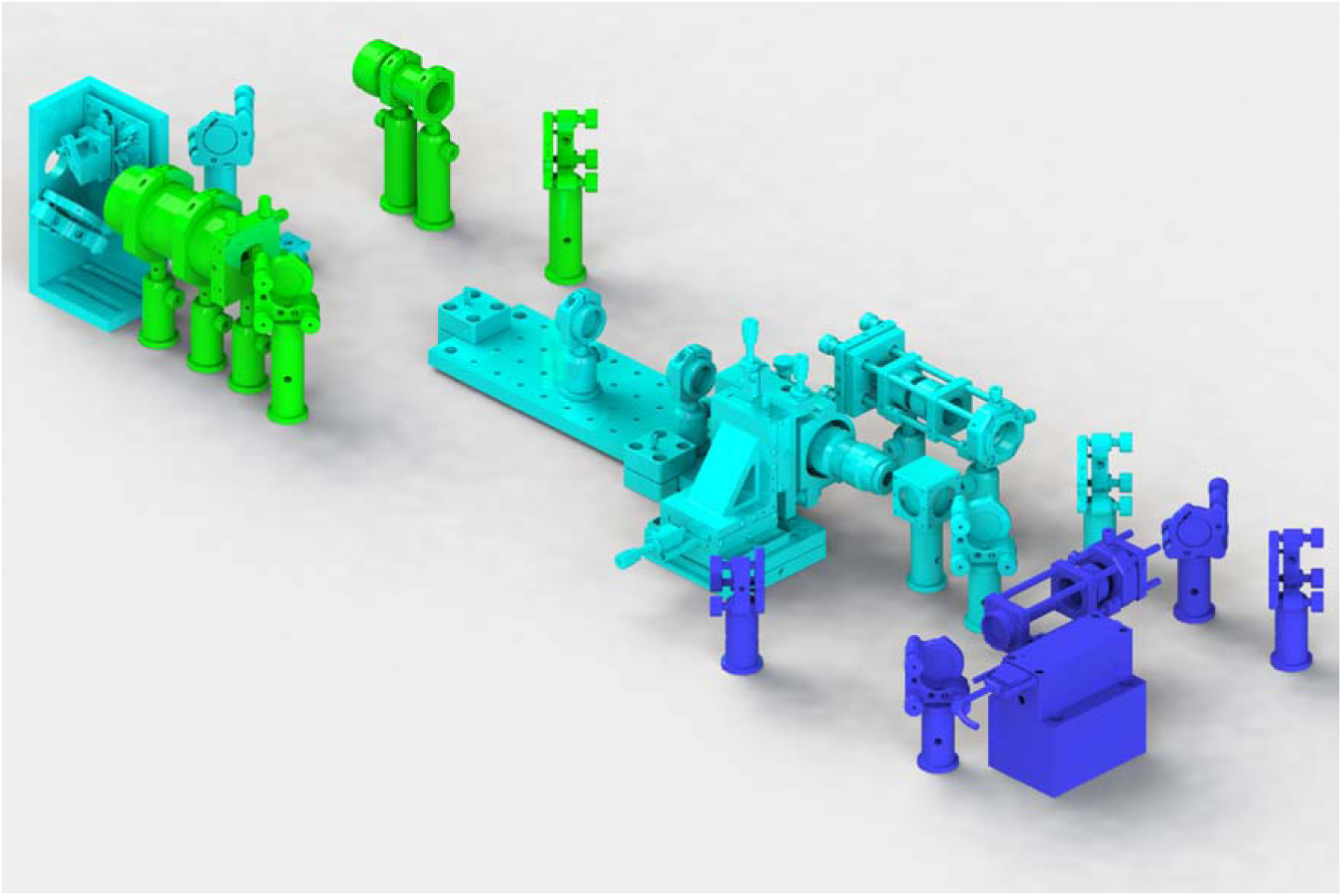
The fully assembled illumination paths showing the different optional modules. Cyan: single-mode laser engine path. Blue: single-mode booster laser path. Green: multi-mode laser engine path. The modules are compatible with each other allowing for the expanded set of capabilities offered by each and allowing for easy expansion of the system as desired.

230. Pre-adjust all three-actuator steering mirror mounts (COTS-POLARIS-K1, COTS-POLARIS- K1E3) for the illumination path to nominal spacing using a ⅛” post height spacer (3D-SMLM- PS-0.125).
231. Build up the single-mode beam launch assembly (3D-SMLM-EMBL-001-003-001) for the single-mode laser engine omitting the two collimating lenses (COTS-AC254-075-A-ML) as shown in Supplementary Figure 34. Secure the assembly on the optical table using 2× pedestal post clamps (COTS-PS-F).
232. Place the single-mode laser engine (COTS-ICHROME-MLE) on its heat dissipating plate at a suitable location and avoid moving it from this position for the remainder of the protocol. Ensure that the single-mode fiber is long enough to reach the FC/APC fiber terminator (COTS-S1FCA) of the single-mode beam launch without tension or tight turns (radius of curvature > 10 cm). Plug the fiber into the Fiber In port on the laser engine.
233. Optimize the alignment of the single-mode lasers into the fiber using the COOL^AC^ auto-calibration function of the TOPAS iChrome software.
234. Install the fiber in the FC/APC fiber terminator (COTS-S1FCA) of the single-mode beam launch assembly, ensuring the key is properly aligned to the slotted recess. Adjust the fiber kinematic mount (COTS-KC1-T-M) to center the diverging beam through the cage system using drop-in cage alignment targets (COTS-CPA1). Use the bronze locking nut (COTS-LN2580) to lock the actuator positions.
235. Install the collimating lenses (COTS-AC254-075-A-ML) with correct orientation but arbitrary spacing. Setup the shear plate assembly (3D-SMLM-EMBL-005) downstream of the beam launch and move the two collimating lenses to collimate the 561 nm beam to an arbitrary diameter. Note, if the laser engine was configured with alternative laser lines, select a line that is centered in the wavelength range.
236. OPTIONAL: if the specified configuration uses an additional booster laser, it is necessary to fix the polarization orientation out of the fiber at this stage. Install the mounted polarization beam splitter (COTS-PBS251, COTS-CCM1-4ER-M, COTS-RS10M, COTS-RS2P4M) after the collimator such that the input (mixed polarization) side is placed where the laser is incident and the horizontal polarization side is opposite to it. Loosen the retaining ring (COTS-SM1RR) securing the FC/APC fiber terminator (COTS-S1FCA). Using a piece of paper to view the beam, rotate the fiber terminator to minimize the power reflected by the polarization beam splitter. Secure the retaining ring.
237. OPTIONAL: Install the booster laser (3D-SMLM-COTS-IBEAM-SMART) on its mount (3D- SMLM-FAB-EMBL-000087) and position the assembly on the optical table to direct the laser into the polarization beam splitter on the intended side of incidence. CAUTION: The beam path is not well defined at this time. Take extra care to intercept stray beams using a laser safety screen (COTS-TPSM1-M).
238. OPTIONAL: Check that the booster laser is efficiently reflected down illumination path by the polarization beam splitter. If the beam is primarily transmitted by the beam splitter then the orientation is not correct. Orient the polarization beam splitter (COTS-PBS251) appropriately in its mount (COTS-CCM1-4ER-M) to efficiently reflect the booster laser and repeat the procedure with the rotation of the FC/APC fiber terminator (COTS-S1FCA) to transmit the single-mode laser engine fiber output.
239. Check the collimated beam diameter on the camera beam profiler (COTS-LASERCAM-HRII). Adjust the spacing between the two collimating lenses (COTS-AC254-075-A-ML) to change the effective combined focal length (decreasing the spacing between the lenses will result in a longer focal length and thus a larger collimated beam diameter) and adjust the divergence/convergence by translating the lenses as a pair to produce a collimated Ill6 ± 0.05 mm (1/e^2^ criterion) beam. Switch between the beam profiler and shear plate as necessary to judge beam diameter and collimation respectively.
240. Install an iris (COTS-SM1D12D) in the XY mount (COTS-CXY1) of the single-mode beam launch and stop it down to a 1 - 2 mm aperture diameter. Use the camera beam profiler to center the iris on the collimated beam such that the diffraction pattern has maximum radial symmetry. When performing alignment operations (aligning the beam to the optical axis of a lens), where a smaller beam is optimal, leave the iris stopped down. When assessing collimation (aligning a lens longitudinally to achieve a minimally divergent output), where a larger beam is optimal, open the iris fully. Similarly for assessment of the beam profile or for imaging, the iris should be fully open.
241. Install the objective port alignment target assembly (3D-SMLM-EMBL-006) in the objective port (FAB-EMBL-000043) with the beam profiling camera installed on the distal end (Supplementary Figure 35). For the following alignment steps, use the camera image to center the beam on the irises.
242. Install the pair of steering mirrors (COTS-BB1-E02, COTS-POLARIS-K1, COTS-RS5M, COTS-RS2P4M) downstream of the single-mode beam launch using 2× pedestal post clamps (COTS-PS-F) as shown in Supplementary Figure 36. Check that the beam is well-centered on the mirror faces and that the second mirror is positioned with its face centered with respect to the respective line of tapped holes on the optical table surface.
243. Use a pair of 80.4 mm beam height alignment targets (3D-SMLM-EMBL-003) to ensure that the beam travels at the correct height and approximately along the respective hole pattern of the optical table.
244. OPTIONAL: Install the polarization beam splitter assembly (COTS-PBS251, COTS-CCM1- 4ER-M, COTS-RS10M, COTS-RS2P4M). CAUTION: Ensure that the polarization beam splitter is oriented to correctly pass the single-mode laser engine and reflect the booster laser given its polarization and intended direction of incidence.
245. Assemble the illumination periscope and inclination control assembly (3D-SMLM-EMBL-001- 003-007, see CAD and parts list for all components, Supplementary Figure 37). CAUTION! Ensure that the M6x30 mm dowel pin (3D-SMLM-COTS-91585A668) is installed to limit the stage motion. This pin provides a safety feature which, when the system is correctly aligned, prevents the illumination lasers from being inadvertently launched obliquely away from the short optical table edge closest to the microscope body. Nevertheless, follow all laser safety guidelines noted in: Laboratory environment. If the illumination control prism mirror mount (FAB-EMBL- 000086) contacts the pin, it may be necessary to rehome the stage (COTS-ELL17-M) using its software (Thorlabs ELLO).
246. Move the inclination control stage (COTS-ELL17-M) to the position that it just contacts the pin and record this position. Rehome the stage and move the stage to: recorded position - 0.75 mm. This defines the nominal position required for non-inclined epi-illumination. Store and save this value in the EMU configuration for Two-state device 2, 3 and 6 - Off values such that turning off the HILO/TIRF/Multi-mode controls, switches the microscope to the default case of epi-illumination.
247. Introduce and align the third steering mirror of the single-mode laser engine illumination path (COTS-BB1-E02, COTS-POLARIS-K1, COTS-RS5M, COTS-RS2P4M, Supplementary Figure 38), ensuring that the laser is centered on the mirror and that the beam is reflected parallel to the hole pattern on the optical table. Use a pair of 80.4 mm beam height alignment targets (3D- SMLM-EMBL-003) aligned to the hole pattern to ensure that the beam travels at the correct height and along the table holes. Position the mirror laterally along the incoming beam direction to align to the first target and angle the mirror to align to the second target in an iterative manner. Secure the mirror in position using a pedestal post clamp (COTS-PS-F) once the beam is aligned to the two targets. CAUTION! If the beam is tilted with regard to the table hole pattern, correct alignment through the subsequent mirrors will be compromised.
248. Introduce the illumination periscope and inclination control assembly (3D-SMLM-EMBL-001- 003-007) in position on the optical table and lightly secure with screws through the slotted counterbores. Install an iris (COTS-SM1D12D) into the entrance aperture in the entrance plate of the assembly (FAB-EMBL-000084) and position the assembly such that the beam passes the iris and is incident close to the center of the subsequent mirror (COTS-POLARIS-K1E3, COTS- BB1-E02). CAUTION! Try to position the illumination periscope and inclination control assembly such that its faces are aligned to the hole pattern on the optical table. Adding additional screws through the counterbores may assist in maintaining reasonable alignment.
249. The beam should now pass into the illumination periscope and inclination control assembly (3D- SMLM-EMBL-001-003-007) and reflect off the upper prism mirror (COTS-MRA20-E02) and into the microscope body.
250. Use the lower steering mirror (COTS-POLARIS-K1E3, COTS-BB1-E02) to align the beam to the second target of the objective port alignment target (3D-SMLM-EMBL-006). The beam should approximately pass the first target in the left/right direction (with respect to the microscope body). To correct for any misalignment here adjust the position of the beam on the first target by using an equal rotation of all three adjusters of the lower steering mirror to translate the mirror face forwards or backwards as necessary and use the topmost adjuster alone to then realign to the second target.
251. For back/front alignment (with respect to microscope body) adjust to the first iris by adjusting the position of the inclination control stage (COTS-ELL17-M) and correct at the second iris using the bottom/left adjuster screw of the illumination periscope and inclination control assembly steering mirror (COTS-POLARIS-K1E3, COTS-BB1-E02). If it is necessary to move the stage for alignment, ensure that the beam remains well centered on the prism mirror (COTS- MRA20-E02) face (± 3 mm) and use the final stage position as the new nominal position for non-inclined epi-illumination. Store and save this value in the EMU configuration for Two-state device 2, 3 and 6 - Off values.
252. Remove the beam profiling camera from the objective port alignment target (3D-SMLM-EMBL- 006). The laser should now be incident on the ceiling or optical table enclosure above the microscope. Mark the position of the beam using a suitable marker. This reference is useful for realignment of the illumination to the objective during routine maintenance. Replace the beam profiling camera when finished.
253. Install the illumination tube lens (COTS-AC254-400-A) approximately half way down the length of its lens tube (COTS-SM1M10) using the two retaining rings (COTS-SM1RR) to secure it in place. CAUTION: the tube lens has a correct orientation with the finite conjugate side (closest to the objective lens port) facing the side of least curvature. Indicate the orientation on the lens tube with a permanent marker with an arrow pointing away from the side of least curvature (towards the infinite conjugate).
254. Assemble the illumination tube lens (COTS-AC245-400-A) on its pedestal post (COTS-PH40E- M, COTS-TRA30-M) with the lens tube split clamp (COTS-SM1RC-M) securing the lens tube (COTS-SM1M10). Introduce the tube lens to the beam path with the arrow pointing upstream as shown in Supplementary Figure 39.
255. Use the back reflection method to align the illumination tube lens in its nominal longitudinal position, placing one 80.4 mm beam height alignment target (3D-SMLM-EMBL-003) upstream of the lens and using one iris of the objective port alignment target (3D-SMLM-EMBL-006). Secure the illumination tube lens assembly in position with a pedestal post clamp (COTS-PS-F)
256. Remove the objective port alignment target (3D-SMLM-EMBL-006) from the objective port (FAB-EMBL-000043) and replace it with the objective lens. CAUTION! A small and divergent beam will exit the objective lens and even small misalignments may result in a strong deviation from the optical axis and constitute a laser safety hazard accordingly. Wear laser safety eyewear whenever assessing alignment through the objective lens. If the laser is launched substantially off-axis, such that it is no longer incident within ± 10 mm of the mark made in step 252 adjust the steering mirror of the illumination periscope and inclination control assembly (COTS- POLARIS-K1E3), to re-center the beam with respect to the objective lens such that the beam is well-centered on the mark.
257. Loosen the split clamp (COTS-SM1RC-M) just enough to allow motion of the tube lens in its lens tube (COTS-AC254-400-A, COTS-SM1M10). Slide the lens tube to collimate the beam by minimizing the laser spot size as viewed on the ceiling or optical table enclosure. Repeat the previous step as necessary to ensure that the beam remains incident within ± 10 mm of the mark. TROUBLESHOOTING!: If the range of motion is insufficient to achieve collimation, release the pedestal post clamp (COTS-PS-F), center the lens tube in the split clamp and manually position the assembly to achieve collimation. Subsequently realign the tube lens as previous but in the new nominal upstream/downstream position for collimation.
258. The microscope is now set up for epi-fluorescence imaging with a Gaussian illumination field.

### Focusing the microscope (Timing: 1 day)

The emission path has been set up to image an object at infinity and basic epi-illumination established thus allowing the microscope to be focused to image the native focal plane of the infinity-corrected objective in fluorescence mode. A fluorescent bead sample is used to mechanically position the objective lens to a coarse focus before focusing more finely with the objective piezo flexure stage. Note that owing to the image-splitter geometry, the two image channels are mirrored relative to each other. Next the bead sample is used to focus the Bertrand lens and to provide an initial adjustment of the objective correction collar (if present), to check the PSF shape more generally, and to check that all color channels appear similarly focused on the camera without the need for channel-specific refocusing.

259. Place a drop of immersion oil onto the objective after checking the applicator nozzle for clearly visible bubbles. Blot away oil from the nozzle as necessary with tissue to remove bubbles. Repeat this process whenever mounting a sample on the microscope.
260. Mount a fluorescent bead sample on the microscope. Refer to: Fluorescent bead samples in the reagent setup section for a detailed description of the sample preparation. Note, when mounting samples on the microscope, use 4× M2×6 mm socket head screws to secure the sample holder FAB-EMBL-000045 in position. Failing to do so can result in greater drift, errors in repositioning and small variations in focus height between a secured and unsecured holder.
261. Move the objective piezo flexure stage (COTS-P-726.1CD) to the middle of its travel range (50 µm).
262. Select an appropriate combination of laser line and emission filter in the reflected/transmitted path filter wheels. If the shared path filter wheel has been installed, switch it to an open position. Illuminate and image the fluorescent bead sample.
263. Focus the image of the fluorescent beads on the camera using the mechanically-translatable objective piezo platform (e.g., rotating the timing pulley (COTS-HTPT48S2M060_A_P6_35). Start in a downward direction (rotating anti-clockwise as viewed from above) and if the focus is not found, move upward carefully to not break the coverslip and/or damage the objective lens. TROUBLESHOOTING: If the two channels appear in focus at different positions of the camera, return to step 204 and ensure that the alignment laser beam is collimated in the two pathways.
264. Once an approximate focus has been achieved, focus more finely using the objective piezo. If the piezo position at focus is within the last 20 µm travel range from either limit, mechanically refocus to better center the focus height with regard to the objective piezo travel range.
265. Ensure the beads are sharply in focus and switch the Bertrand lens into the emission path. Installed in its nominal position, the Bertrand lens should produce an approximately focused image of the objective back focal plane on the camera (potentially in both sides of the camera chip depending on the emission filters used).
266. Translate the Bertrand lens in its piezo resonant slider (COTS-ELL6) using the threaded mount (COTS-SM1AD20) such that the periphery of the back focal plane image appears maximally sharp. Lock the lock ring (COTS-SM1NT) to secure the position of the lens. If bubbles are present in the oil, dark occlusions may also be present on the back focal plane image. If this is the case, clean the objective and remove/replace oil. Continue with the protocol only when no bubbles are present. Repeat this step whenever mounting samples on the microscope.
267. Switch the Bertrand lens out of the emission path.
268. Observe the general PSF shape when focusing through the beads above and below the focal plane. Adjust the correction collar of the objective lens (if present) to achieve the best possible symmetry above and below the focus by visual inspection. Note, adjusting the correction collar will introduce defocus, which should be corrected by focusing as usual. When the collar is incorrectly positioned, on one side of the focus, the PSF will appear as a set of concentric rings, while on the other, the PSF will quickly fade into the background signal. Rotating the collar allows the ring/fade sides to be switched from above to below the focus and vice versa. The ideal position for the collar is the transition between rings above and below focus where spherical aberrations are minimized.
269. Acquire a z-stack of the fluorescent bead sample (over a range of ± 1 µm, 0.05 µm spacing) using the µManager multi-dimensional acquisition wizard.
270. Use ImageJ to load the image stack, set the voxel dimensions in (Image/Properties, 0.105, 0.105, 0.05 µm, height, width, depth). Reslice the stack (Image/Stacks/Reslice) with start at ‘Top’ selected and no other boxes checked. Make a maximum intensity projection of the resliced stack (Image/Stacks/Z Project) to view a projection of the beads as seen from the side. Check the PSF symmetry in z (now the vertical direction in the resulting image). Adjust the correction collar and repeat this process as appropriate. The correction collar position should be in the range 0.165 - 0.175 mm. TROUBLESHOOTING! If the final correction collar position is outside the range 0.165 - 0.175 mm then it is possible that the infinite conjugate viewing camera and/or infinite conjugate transmission target are not set up accurately enough to provide appropriate infinite conjugate references. In this case, return the collar to its nominal 0.17 mm position and then focus the camera (as in step 218) as well as the objective (using the objective piezo) to find a piezo/camera position where the spherical aberration is minimized. Return to step 263 to check that the two channels are evenly focused.
271. Switch to other appropriate combinations of laser line and emission filter in the reflected/transmitted path filter wheels. Check that the beads are focused at the same axial position for all emission bands to within ± 100 nm. TROUBLESHOOTING! If an even focus for all emission bands cannot be achieved, it is possible that the objective lens is still not imaging at its native (infinity corrected) focal plane. Place a camera (e.g., the camera used for the infinite conjugate viewing camera but without the additional components at the primary image produced by the emission path tube lens (COTS-TTL-165-A) and check whether the color performance is similar and whether the spherical aberrations are similarly well corrected when focused to the same plane as the sCMOS camera (secondary image). Note, it is useful to be able to remove/replace the camera at the primary image to be able to check that the focus has not drifted using the sCMOS camera, If the performance is comparable, it is likely that the objective is not imaging its native focal plane. Return to step 218 to refocus the sCMOS camera and recheck the color performance and spherical aberrations. If the color performance is substantially different between the primary and secondary images, it is likely that the relay lenses which produce the secondary images are incorrectly set up. Return to step 204 to reconfigure the emission path, taking additional care to ensure beam collimation through the use of the shear plate.
272. Remove the objective lens from the objective port (FAB-EMBL-000043).

### Field variable and homogeneous epi-illumination alignment (Timing: 5 days)

The basic epi-illumination system established in previous steps provides the basis for the addition of field-variable and illumination homogenizing modules. Several exchangeable Keplerian telescopes are aligned to the single-mode laser path to allow the user to quickly optimize the illumination field size for a given application. Each telescope has an equal 4f track length ensuring that various conjugates are maintained through the optical path upon exchange. Note that builders wishing to omit the exchangeable telescope system can do so either by choosing one telescope to set a fixed non-unit magnification or omitting the relay entirely and positioning the iris at the conjugate image plane one focal length upstream of the illumination tube lens. In this case, some modification of the protocol steps and parts used will be necessary. An iris is installed upstream of the telescopes, providing a field aperture to allow clean-up/cropping of the illumination field independent of the telescopic magnification. Subsequently, a refractive beam shaper is introduced to flatten the Gaussian output. It should be noted that while the beam shaper is achromatic in design, unavoidable small spectral variations (± 10 %) in the beam diameters from the single-mode laser engine, results in a homogenization efficiency that scales accordingly. Nevertheless, when correctly adjusted, the beam homogeneity for all wavelengths is better than the case of a Gaussian beam truncated at 80% peak intensity and achieves ca. 10 × greater power throughput/efficiency. Subsequent alignment of the beam shaper requires careful pitch/yaw and lateral alignment to produce a homogenized output and the use of the beam profiling camera to assess the homogeneity of the output.

273. Install the shear plate assembly (3D-SMLM-EMBL-005) at the objective port (FAB-EMBL- 000043) using an RMS to SM1 thread adapter (COTS-SM1A4) and lens tube (COTS-SM1S10)
274. Install the kinematic magnetic bases (COTS-AKP-BF, COTS-AKP-BC, COTS-AKP-BV) with the telescopic relay breadboard (COTS-M-SA2-04X12, 3× COTS-AKP-TF) in the nominal position on the optical table (Supplementary Figure 40).
275. Leaving the kinematic magnetic bases (COTS-AKP-BF, COTS-AKP-BC, COTS-AKP-BV) in place, assemble additional breadboards (COTS-M-SA2-04X12, 3× COTS-AKP-TF) to mate with the kinematic magnetic bases for each additional magnifying telescope (as specified by the builder).
276. For the 0.5, 1 and 2× telescopic relays, install all lenses (2× COTS-AC254-040-A, 2× COTS- AC254-060-A, 2× COTS-AC254-080-A) approximately half way down the length of their lens tubes (COTS-SM1M10) using two retaining rings (COTS-SM1RR) to secure them in place. For each lens tube, use a permanent marker to add an arrow pointing towards the most curved side of the lens and the effective focal length.
277. For the 2.85× telescopic relay, install the lenses COTS-AC254-075-A and COTS-AC254-050-A into one lens tube (COTS-SM1M10) with the flatter surface of COTS-AC254-075-A in contact with the more curved surface of COTS-AC254-050-A. Take extra care when placing lenses in contact, first use clean compressed air (COTS-LAB16/LAB16-EU) to ensure neither contact surface nor the lens tube have any debris that may cause scratches. Use two retaining rings (COTS-SM1RR) to hold the first lens in place and install the second lens by installing it from below the lens tube with its retaining ring (COTS-SM1RR) using an SM1 spanner wrench (COTS-SPW602) to thread the retaining ring carefully into the lens tube, stopping when there is resistance signifying contact of the two lenses. Use a permanent marker to add an arrow pointing towards the more curved side of the COTS-AC254-075-A lens and the effective focal length (31.2 mm for the combined lens pair). Install the lens COTS-#49-769 into a lens tube, secure it in position and mark it as described in the previous step.
278. Assemble all lens tube mounted lenses with their split clamp/pedestal post assembly (COTS- SM1RC-M, COTS-PH30E-M, COTS-TRA20-M) and secure the split clamp with the lens tube centered.
279. For the 1× magnification (2× COTS-AC254-060-A) telescopic relay, introduce the downstream lens assembly on the breadboard (COTS-M-SA2-04X12) with the arrow pointing downstream and position to approximately align and collimate the beam as viewed on the shear plate. Mark the nominal longitudinal position (i.e., along the beam axis) of the pedestal post base (e.g., with colored tape).
280. Remove the shear plate assembly (3D-SMLM-EMBL-005) from the objective port (FAB- EMBL-000043) and replace it with the objective port alignment target (3D-SMLM-EMBL-006) with the beam profiling camera installed.
281. Use the back reflection method to align the lens (AC254-060-A) approximately in the longitudinal position marked, placing one 80.4 mm beam height alignment target (3D-SMLM- EMBL-003) upstream of the lens and a second placed downstream, close to the illumination tube lens (COTS-AC254-400-A). Check alignment through one iris of the objective port alignment target (3D-SMLM-EMBL-006). If the alignment has been performed correctly the beam should pass this iris on its axis to within ± 0.25 mm. Secure the pedestal post (COTS-PH30E-M) in position with a pedestal post clamp (COTS-PS-F)
282. Remove the objective port alignment target (3D-SMLM-EMBL-006) from the objective port (FAB-EMBL-000043) and replace it with the shear plate assembly (3D-SMLM-EMBL-005).
283. Loosen the split clamp (COTS-SM1RC-M) securing the lens tube (COTS-SM1M10). Translate the lens tube to achieve collimation. Secure the split clamp.
284. Remove the shear plate assembly (3D-SMLM-EMBL-005) from the objective port (FAB- EMBL-000043) and replace it with the objective port alignment target (3D-SMLM-EMBL-006) with the beam profiling camera installed.
285. Install the shear plate assembly (3D-SMLM-EMBL-005) downstream of the breadboard (COTS- M-SA2-04X12).
286. For the 1× magnification (2× COTS-AC254-060-A) telescopic relay, introduce the upstream lens assembly on the breadboard (COTS-M-SA2-04X12) with the arrow pointing upstream and position to approximately align and collimate the beam as viewed on the shear plate. Mark the nominal longitudinal position of the pedestal post base as previous.
287. Move the shear plate assembly out of the beam path (3D-SMLM-EMBL-005).
288. Use the back reflection method to align the lens (COTS-AC254-060-A) approximately in the longitudinal position marked, placing one 80.4 mm beam height alignment target (3D-SMLM- EMBL-003) upstream of the lens and a second placed downstream, close to the lens installed previously on the breadboard (COTS-AC254-060-A). Check alignment through one iris of the objective port alignment target (3D-SMLM-EMBL-006). If the alignment has been performed correctly the beam should pass this iris on its axis to within ± 0.25 mm. Note that the beam will be divergent after passing the illumination tube lens (AC254-400-A). Secure the pedestal post (COTS-PH30E-M) in position with a pedestal post clamp (COTS-PS-F)
289. Return the shear plate assembly to the beam path (3D-SMLM-EMBL-005).
290. Loosen the split clamp (COTS-SM1RC-M) securing the lens tube (COTS-SM1M10). Translate the lens tube to achieve collimation. Secure the split clamp.
291. The 1× telescope is now aligned (Supplementary Figure 41) and the lenses positioned to conjugate the front focal plane of the upstream lens with the sample plane. Repeat this process for each telescopic relay breadboard (COTS-M-SA2-04X12, 3× COTS-AKP-TF, see Supplementary Table 12: Field variable illumination telescopic relays for a list of lens part numbers, which differ between telescopic relays) as above. CAUTION!: these alignment steps must be carried out with precision, otherwise the exchange of the platforms will result in differential misalignments of the illumination path.
292. Switch to the 1× telescope.
293. Assemble the beam shaper assembly (3D-SMLM-EMBL-001-003-005: FAB-EMBL-000082, COTS-KBM1-M, COTS-ULM-TILT, COTS-M-462-XZ-M, COTS-AJS100) but omit the beam shaper (COTS-PISHAPER-6_6_VIS) itself. Adjust the pitch/yaw (COTS-ULM-TILT) and XZ stages (COTS-M-462-XZ-M) to neutral positions.
294. Install the beam shaper assembly on its base (COTS-KBM1-M). The assembly should appear as shown in Supplementary Figure 42
295. Position the beam shaper base (COTS-KBM1-M) on the optical table such that the iris (COTS- SM1D12D) is positioned approximately 60 mm upstream of the upstream lens of the 1× telescope (COTS-AC254-060-A) and aligned to the beam (Supplementary Figure 43).
296. Remove the objective port alignment target (3D-SMLM-EMBL-006) from the objective port (FAB-EMBL-000043) and install the objective lens in its place.
297. Mount and image a fluorescent dye sandwich sample. Refer to: Fluorescent dye sandwich in the reagent setup section for a detailed description of the sample preparation. Focus on the upper side of the lower coverslip in brightfield mode by illuminating with the ring LED (FAB-EMBL- 000037) and imaging with all filter wheels/sliders switched to an open position. It should be possible to focus on some particulate matter on the coverslip surface.
298. Image the dye sandwich in fluorescence mode. Stop down the iris (COTS-SM1D12D) to observe the effect of the blades cropping the illuminated field of view on the camera image.
299. Longitudinally position the beam shaper mount base (COTS-KBM1-M) on the optical table such that the cropped edges of the illumination field are maximally sharp in the image while maintaining approximate alignment of the iris to the laser source.
300. Center the iris (COTS-SM1D12D) on the beam by inspecting the image and translating the beam shaper mount (FAB-EMBL-000082) with the XZ stage (COTS-M-462-XZ-M).
301. Install the beam shaper (COTS-PISHAPER-6_6_VIS) in its mount (FAB-EMBL-000082) as shown in Supplementary Figure 44. CAUTION! The beam shaper has the same M27x1.0 thread at its entrance and exit apertures. Ensure that the beam shaper is threaded into the mount in the correct orientation as shown in the CAD assembly (the graduated tick marks on the housing should be outside of the pitch/yaw (COTS-ULM-TILT) stage when the beam shaper is installed in the correct orientation.
302. Remove the 1× telescope and install the beam profiling camera to profile the beam output close to the beam shaper exit aperture.
303. Ensure that any irises cropping the beam are fully open and observe the 561 nm beam profile. It is likely that the beam shaper is not well-aligned to the beam resulting in a highly non-uniform intensity. The four alignment adjusters of the pitch/yaw (COTS-ULM-TILT) and XZ stages (COTS-M-462-XZ-M) should be used as two pairs (yaw/X) and pitch/Z) to achieve an even illumination spot. Typically, an edge or ring of slightly higher intensity may be present towards the periphery of the spot. Iteratively, use the angular adjustment to evenly distribute the brighter region up/down and left/right and the translation to provide an even intensity across the spot. The degree of homogeneity is highly dependent on the beam quality and some small variation across the field is to be expected.
304. Remove the camera beam profiler and replace it with the shear plate (3D-SMLM-EMBL-005). Adjust the longitudinal spacing of the beam shaper elements by releasing the locking ring and screwing the two sections in/out before re-locking to provide optimum collimation of the output beam. Check and adjust the alignment as above.
305. Remove the upper plate of the beam shaper mount base (COTS-KBM1-M) with all affixed components.
306. To aid periodic realignment of the single-mode excitation path, install two 80.4 mm beam height kinematic magnetic alignment targets (3D-SMLM-EMBL-008) as indicated in Supplementary Figure 45 (precise longitudinal placement is not critical) and align them to the laser. Secure the targets in place using pedestal post clamps (COTS-PS-F).
307. Remove the top plate of the magnetic mount (COTS-KB25M) with the upper section of each target. Use a permanent marker to indicate on each which is the upstream/downstream target for future use.
308. Fully open any irises present in the illumination path.
309. Reinstall the 1× telescope and upper plate of the beam shaper mount base (COTS-KBM1-M) with all affixed components.
310. Install the illumination laser clean-up filter in a lens tube (COTS-SM1L03) with the intended propagation of light moving from the external to internal threaded sections. See Supplementary Note 2: Selecting spectrally discriminative optics. Install the lens tube into the iris (COTS- SM1D12D) of the beam shaper assembly (3D-SMLM-EMBL-001-003-005) as shown in Figure Supplementary Figure 46.
311. Image the fluorescent dye sandwich sample and check that the specified change in the illumination field diameter is visible when switching between telescopes without repositioning the field of view on the camera (within an allowable error of ± 2 µm, ca. ± 20 pixels) relative to the position of the illumination field without any telescope installed or a consensus position shared by two or more telescopes. Note that without a telescope installed, the beam profile at the object will not be homogeneous but it should still be possible to determine its center. TROUBLESHOOTING! If any of the telescopes result in a center position that does not meet this criterion, return to step 285 to realign them to the beam.
312. Check the homogeneity of the illumination spots at all wavelengths. Since the input beam diameter and divergence will vary with wavelength, the degree of homogenization will also vary accordingly.
313. Partially close the iris (COTS-SM1D12D) mounted to the beam shaper mount (COTS-EMBL- 000082) to sharply apodize the illumination spot at its periphery such that no dimly illuminated beads are visible. Lock the iris position.

### OPTIONAL: Installation of the booster laser (Timing: 2 days)

A free-space single-mode booster laser can be combined with the single-mode laser engine at the polarization beam splitter installed previously to provide additional laser power at a single wavelength. The free-space Booster laser requires spatial filtering to clean the beam profile (since the beam shaper requires a high-quality Gaussian input). The spatial filter serves the dual purpose of expanding the booster laser to the required 6 mm beam size. The booster laser expansion and co-alignment with the single-mode laser engine through the beam shaper are facilitated by use of the beam profiling camera. The profile and co-alignment are subsequently validated with a fluorescent bead sample.

314. Pre-adjust all three-actuator steering mirror mounts (COTS-POLARIS-K1) for the illumination path to nominal spacing using a ⅛” post height spacer (3D-SMLM-PS-0.125).
315. Assemble the booster laser spatial filter (3D-SMLM-EMBL-001-003-002) omitting the collimating lenses (2× COTS-LA1986-A), the pinhole (COTS-P15K) and the focusing lens and associated optomechanics (COTS-LA1540-A-ML, COTS-CP33/M, COTS-SM1A6) as shown in Supplementary Figure 47. Install the assembly on the optical table using 2× pedestal post clamps (COTS-PS-F).
316. Install the mounted booster laser (3D-SMLM-COTS-IBEAM-SMART, 3D-SMLM-FAB- EMBL-000087) on the optical table.
317. Install the first pair of steering mirrors (COTS-POLARIS-K1, COTS-BB1-E02, COTS- RS2P4M, COTS-RS5M) to route the beam to the spatial filter with the laser incident on the center of the mirror faces as shown in Supplementary Figure 48. Secure the mirrors in place using 2× pedestal post clamps (COTS-PS-F)
318. Align the booster laser on-axis through the spatial filter cage system using the two steering mirrors and a pair of 30 mm cage drop in alignment targets (COTS-CPA1).
319. Install the pinhole (COTS-P15K) and the focusing lens with its associated optomechanics (COTS-LA1540-A-ML, COTS-CP33/M, COTS-SM1A6).
320. Position the lens (COTS-LA1540-A-ML) and pinhole (COTS-P15K) relative to each other to spatially filter the beam. Sliding the lens in its cage plate (CP33-M) along the cage rods (COTS- ER6) allows axial alignment, while the pinhole can be moved via its XY mount (COTS-CXY1) for lateral alignment. The quality of the alignment can be judged by the transmitted beam circularity and power throughput, which can be measured using the laser power meter (COTS- PM100D) with its SM1 externally threaded sensor (COTS-S121C) mounted to the last cage plate (COTS-CP33-M) of the spatial filter. With correct alignment > 60 % power throughput is achievable. Note, positioning the cage plate for correct focusing through the pinhole is easier if using a ball-tipped kinematic adjuster (COTS-KL02L) on an optical post to make adjustments.
321. Install the collimating lenses (2× COTS-LA1986-A) and shape the beam to a IZ6 ± 0.05 mm (1/e^2^ criterion), as described for the single-mode laser engine in step 239.
322. Remove the upper plate of the beam shaper mount base (COTS-KBM1-M) with all affixed components as well as the 1× telescope.
323. Reinstall the two 80.4 mm beam height kinematic magnetic alignment targets (3D-SMLM- EMBL-008) by reunifying the top and bottom plates of the magnetic mount (COTS-KB25M). CAUTION!: Ensure that the correct target is placed on the corresponding pedestal following the markings made previously.
324. Install the second pair of steering mirrors (COTS-POLARIS-K1, COTS-BB1-E02, COTS- RS2P4M, COTS-RS5M) as shown in Supplementary Figure 49 to route the collimated output from the spatial filter via the polarization beam splitter (COTS-PBS251 mounted in COTS- CCM1-4ER-M) to the two targets with the laser incident on the center of the mirror faces. Secure the mirrors in place using 2× pedestal post clamps (COTS-PS-F).
325. Remove the top plate of the magnetic mount (COTS-KB25M) of each 80.4 mm beam height kinematic magnetic alignment targets (3D-SMLM-EMBL-008) with the upper section of each target.
326. Replace the upper plate of the beam shaper mount base (COTS-KBM1-M) with all affixed components.
327. Switch off the booster laser and switch on the 561 nm laser of the single-mode laser engine.
328. Repeat the procedure of aligning the beam shaper using the beam profiling camera as well as the pitch/yaw (COTS-ULM-TILT) and XZ stages (COTS-M-462-XZ-M) as necessary.
329. Switch on the booster laser and regulate the power of both sources to achieve approximately equal intensity at the beam profiling camera and such that when both sources are present, the camera does not saturate.
330. Switch off the single-mode laser engine.
331. Align the booster laser to the beam shaper using the pair of steering mirrors upstream of the polarization beam splitter (COTS-POLARIS-K1, COTS-BB1-E02, COTS-RS2P4M, COTS- RS5M). Much as for the alignment using the pitch/yaw (COTS-ULM-TILT) and XZ stages (COTS-M-462-XZ-M), use the mirror actuators in pairs to adjust the x and y beam profiles.
332. Switch on the 561 nm laser of the single-mode laser engine. Block/unblock the booster laser periodically using a laser safety screen (COTS-TPSM1-M) to check for co-alignment between the two beams on the beam profiling camera. Once the booster laser output is well-flattened and well-aligned to the beam from the single-mode laser engine, the alignment is complete. Otherwise repeat the previous steps to optimize the alignment of the two lasers.
333. Replace the 1× telescope.
334. Image a fluorescent bead sample.
335. Check that the illumination spots from the booster laser and single-mode laser engine are overlaid in the resulting camera image with a centration better than ± 1 µm ca. ± 10 pixels).

### OPTIONAL: HILO/TIRF setup (Timing: 1 day)

Having installed all TIRF compatible single-mode sources, it is possible to establish various settings for HILO/TIRF illumination. Delivering HILO/TIRF illumination requires that the illumination lasers are shifted from the center of the objective back focal plane towards its periphery such that the collimated beams are launched obliquely by the objective lens to just avoid (HILO) or fulfil the total-internal reflection criterion (TIRF). This is accomplished via a positional shift of the illumination inclination control mirror/stage. In the case of a silicone oil objective lens, the refractive index contrast between the immersion oil and the sample is too small to fulfil the TIRF condition at the angle defined by the maximum NA of the objective lens. In this case, one need only configure the stage position for HILO illumination.

336. Mount and image a fluorescent bead sample. Refer to: Fluorescent bead samples in the reagent setup section for a detailed description of the sample preparation and: Focusing the microscope for a description of setting up the microscope to image fluorescent beads. Ensure that the bead sample is immersed in water.
337. View the back focal plane using the Bertrand lens, check for the presence of bubbles.
338. Ensure that the cover is correctly installed on the microscope to intercept obliquely launched beams.
339. Move the illumination inclination control stage (COTS-ELL17-M) in either direction from its nominal position to observe the shift in the beam position at the back focal plane. Translation of the stage results in an equal translation of the beam at the back focal plane. Since the back focal plane diameter is in the range 4.5 - 5.5 mm depending on the specific objective, a translation of 2.25 - 2.75 mm should be sufficient to translate the beam to the back focal plane periphery.
340. Move the beam towards the BFP periphery until further motion results in attenuation of the laser spot. The laser should now be reflected at the coverslip/sample boundary and should pass back through the objective lens and should be visibly apparent passing back down the illumination path laterally offset a few mm from the illumination beam on its way into the microscope. For example, the incoming illumination and outgoing total internal reflected beams are focused intermediate between the two lenses of the illumination telescope, which facilitates their viewing (at low laser power) with paper. The reflected beam may also be visible on the opposite side of the back focal plane image. Check also that the beads remain visible on the coverslip without major changes in the illumination spot shape or homogeneity. TROUBLESHOOTING! If the illumination field homogeneity degrades under the TIR condition, this typically signifies that the illumination laser is not focused to the back focal plane of the objective lens. If so, return to step 256 to refocus the illumination tube lens (COTS-AC254–400-A) as described.
341. The angle determined is the maximum angle for TIRF illumination. Store and save the stage position value in the EMU configuration for the Two-state device 2 - On value, such that turning on the TIRF control, switches the microscope to TIRF illumination.
342. Determine the critical (minimum) angle for TIRF illumination by returning the beam towards the BFP center until the total internal reflected beam is no longer present. The transition between the TIRF/non-TIRF regime is gradual, requiring some judgement as to where TIRF does/does not occur. Try to determine this position as the centermost beam position for which the total internal reflected beam intensity does not noticeably decrease. Record the critical stage position for later use. Note, between the maximum and critical (minimum) angles for TIRF illumination there are many usable angles that will result in a different penetration depth of the evanescent field. When imaging in TIRF mode, it is advisable to tune the TIRF angle to achieve optimum results.
343. The stage position for HILO is judged as a largest offset that does not elicit a strong TIRF response. Determine the appropriate position as above but find a limiting value where the TIRF condition is not met. Store and save this value in the EMU configuration for the Two-state device 3 - On value, such that turning on the HILO control, switches the microscope to HILO illumination.
344. The precise positions for TIRF and HILO will likely need adjustment on a sample-by-sample basis and/or when re-aligning the microscope. However, the saved positions will provide a useful starting point. When imaging a sample, the maximum TIRF angle is that for which the image gets dim and below the critical angle the image gets brighter with less contrast as background appears from regions for which the TIRF evanescent wave does not penetrate.

### OPTIONAL: Installation of the multi-mode laser engine (Timing: 2 days)

The multi-mode laser engine provides a conveniently homogenized option for epi-illumination and a comparatively inexpensive alternative to the beam shaper and single-mode laser engine. The multi-mode lasers are coupled into the illumination path by moving the illumination inclination control mirror (for HILO/TIRF illumination) fully out of the beam path, thus allowing the multi-mode lasers to pass into the microscope body. Note, if building the microscope with the option of multi-mode illumination only, the entire illumination periscope and inclination control assembly can simply be omitted. The alignment laser is used to define the route of the beam and to align/focus the retrofocus tube lens to the back focal plane of the objective. A slit is installed as a field aperture (placed conjugate to the object plane) with the assistance of the infinite conjugate viewing camera. Having established the multi-mode path, the multi-mode fiber is introduced and focused relative to its projecting lens to image the fiber tip onto the slit and in turn onto the sample. Note, the path length from the beam launch to the adjustable slit will determine the precise magnification of the square fiber tip and the illuminated field of view accordingly. While exact placement of the beam launch, adjustable slit and intervening mirrors is not critical, deviation from the intended placement/path length should be minimized. The multi-mode laser engine is described elsewhere [35] and the protocol assumes that a pre-built and working engine is available.

345. Pre-adjust all three-actuator steering mirror mounts (COTS-POLARIS-K1) for the illumination path to nominal spacing using a ⅛” post height spacer (3D-SMLM-PS-0.125).
346. Move the illumination inclination control mirror/stage (COTS-MRAE02, COTS-ELL17-M) to 0 mm such that the mirror clears the exit aperture of the illumination periscope and inclination control assembly (3D-SMLM-EMBL-001-003-007). This defines the nominal position required for multi-mode illumination. Store and save this value in the EMU configuration for the Two-state device 6 - On value.
347. Remove the objective lens from the objective port (FAB-EMBL-000043). Replace the objective lens with the objective port alignment target (3D-SMLM-EMBL-006).
348. Assemble the multi-mode illumination beam launch (3D-SMLM-EMBL-001-003-008) and place it in the position indicated on the optical table (see Supplementary Figure 50). CAUTION! When installing the lenses 2× COTS-AC254-040-A into the lens tube (COTS-SM1M20). The flatter surface of one lens should be placed in contact with the more curved surface of the other. Take extra care when placing lenses in contact, first use clean compressed air (COTS-LAB16/LAB16- EU) to ensure neither contact surface nor the lens tube have any debris that may cause scratches. Use two retaining rings (COTS-SM1RR) to hold the first lens in place and install the second lens by installing it from below the lens tube with its retaining ring (COTS-SM1RR) using an SM1 spanner wrench (COTS-SPW602) to thread the retaining ring carefully into the lens tube, stopping when there is resistance signifying contact of the two lenses. Use a permanent marker to add an arrow pointing towards the outward facing curved side of the COTS-AC254-040-A lens.
349. Install the laser clean-up filter 5 - 10 mm from the end of the lens tube furthest from the fiber mount (COTS-SM1M20) using two retaining rings (COTS-SM1RR) ensuring that the caret/arrow is pointing in the correct direction as defined by the manufacturer. See Supplementary Note 2: Selecting spectrally discriminative optics. Note for configurations using both single- and multi-mode sources, the clean-up filter (which will have been previously installed in the single-mode path) can instead be accommodated in the shared path inside the microscope body (i.e., just upstream of the illumination dichroic, COTS-25.5×36x3MM- DICHROIC-ILL) by mounting it with the additional components: COTS-PH50E-M, COTS- TRA30-M, COTS-LMR1-M, COTS-PS-F (The components are listed but not enumerated in the parts list and not included in the CAD assembly).
350. Remove the lens tube and affixed components (COTS-SM1M20, COTS-SM1ZM, COTS- SM1FC, 2× COTS-AC254-040-A) by releasing the split clamps (COTS-SM1RC-M). Install the alignment laser assembly (3D-SMLM-EMBL-007) through the split clamps. Note, it may be necessary to partially disassemble the alignment laser assembly to do so). Secure the alignment laser in the split clamps. Make sure the alignment laser is aligned through the two irises of the assembly.
351. Reorient the beam launch such that the alignment laser output travels approximately parallel with the hole pattern of the optical table. Secure the beam launch on the table using 2× pedestal post clamps (COTS-PS-F).
352. Introduce the two steering mirrors (COTS-BB1-E02, COTS-POLARIS-K1, COTS-RS3P4M, COTS-RS10M) with the beam centered on the mirror faces and positioned as indicated in Supplementary Figure 51 such that the beam passes approximately on-axis through the two irises (COTS-SM1D12D) of the objective port alignment target (3D-SMLM-EMBL-006). Secure the mirrors on the table using 2× pedestal post clamps (COTS-PS-F). Use the mirror actuators to more accurately align the beam through the irises.
353. Assemble the retrofocus tube lens (3D-SMLM-EMBL-001-003-009) with the lens tube in a nominal position in the split clamps. and introduce it in the beam path (Supplementary Figure 52). Align the tube lens using the back reflection method, placing one 110.4 mm alignment target (3D-SMLM-EMBL-003) upstream of the tube lens and using one iris (COTS-SM1D12D) of the objective port alignment target (3D-SMLM-EMBL-006) as a downstream target.
354. Remove the objective port alignment target (3D-SMLM-EMBL-006) from the objective port (FAB-EMBL-000043) and replace it with the objective lens. CAUTION! When illuminating with the alignment laser, a small and divergent beam will exit the objective lens and even small misalignments may result in a strong deviation from the optical axis and constitute a laser safety hazard accordingly. Wear laser safety eyewear whenever assessing alignment through the objective lens. Some misalignment is acceptable at this stage, when the alignment laser is replaced by the multi-mode laser, some realignment will be necessary in any case.
355. Ensure that the alignment laser is not cropped by irises and loosen the split clamps (COTS- SM2RC-M) securing the lens tubes of the retrofocus tube lens (COTS-SM2L20, COTS- SM2M25). Focus the tube lens by minimizing the laser spot size from the objective lens when viewed on the ceiling of the room or enclosure of the optical table (as appropriate).
356. Remove the alignment laser assembly (3D-SMLM-EMBL-007) from the multi-mode illumination beam launch (3D-SMLM-EMBL-001-003-008) taking care not to move the split clamps (COTS-SM1RC-M) from their aligned position.
357. Remove the objective port alignment target (3D-SMLM-EMBL-006) from the objective port (FAB-EMBL-000043) and replace it with the infinite conjugate viewing camera (3D-SMLM- EMBL-004, using COTS-SM1A4, COTS-SM1S10 to couple the assembly to the objective port).
358. Install the slit in its nominal position (COTS-SP60, COTS-PH50E-M, COTS-TRA50-M) as shown in Supplementary Figure 53. Backlight the slit with an incoherent white light source and position the slit such that the aperture edges appear as sharp as possible on the infinite viewing camera while ensuring that the slit is oriented perpendicular to the optical axis of the tube lens. Secure the slit in place using a pedestal post clamp (COTS-PS-F).
359. Open the slit (COTS-SP60) using the 4 adjusters until the slit blades are just visible in the camera image (or fully open, whichever comes first, which will depend on the choice of camera).
360. Reinstall the multi-mode illumination beam launch (3D-SMLM-EMBL-001-003-008) lens tube with affixed components (COTS-SM1M20, COTS-SM1ZM, COTS-SM1FC, 2× COTS-AC254- 040-A) through the split clamps of the assembly (COTS-SM1RC-M).
361. Install the multi-mode fiber of the multi-mode laser engine (FAB-EMBL-000094) in the fiber terminator (COTS-SM1FC) and switch on the multi-mode fiber agitation unit.
362. Rotate the lens tube (COTS-SM1M20) in the split clamps (COTS-SM1RC-M) to orient the square fiber such that its image is aligned with the slit axes. Note, the multi-mode beam may not be aligned to the slit itself at this stage. Secure the split clamps.
363. Remove the infinite conjugate viewing camera (3D-SMLM-EMBL-004) from the objective port (FAB-EMBL-000043) and replace it with the objective port alignment target (3D-SMLM- EMBL-006).
364. Use the two steering mirrors (COTS-BB1-E02, COTS-POLARIS-K1, COTS-RS3P4M, COTS- RS10M) to align the multi-mode beam through the irises of the objective port alignment target (3D-SMLM-EMBL-006).
365. Remove objective port alignment target (3D-SMLM-EMBL-006) from the objective port (FAB- EMBL-000043) and replace it with the infinite conjugate viewing camera (3D-SMLM-EMBL- 004).
366. Install an ND6 filter at the end of the multi-mode illumination beam launch (3D-SMLM-EMBL- 001-003-008) lens tube (COTS-SM1M20). In subsequent steps, decrease the neutral density to observe the beam on the infinite conjugate camera without saturating.
367. Focus the fiber in the focusing mount (COTS-SM1ZM) such that the secondary image of the beam at the slit (COTS-SP60) (on the infinite conjugate camera) appears maximally uniform. TROUBLESHOOTING! If the square beam does not look uniform, regardless of the fiber focus position, this signifies that the beam is being clipped. Check the beam profile throughout its propagation with a piece of paper to detect where the beam is being clipped and return to earlier alignment steps as appropriate.
368. Remove the infinite conjugate viewing camera (3D-SMLM-EMBL-004) from the objective port (FAB-EMBL-000043) and replace it with the objective lens. CAUTION! When illuminating with the alignment laser, a small and divergent beam will exit the objective lens and even small misalignments may result in a strong deviation from the optical axis and constitute a laser safety hazard accordingly. Wear laser safety eyewear whenever assessing alignment through the objective lens. If the laser is launched substantially off-axis, such that it is no longer incident within ± 10 mm of the mark made in step 252 adjust the downstream of the two steering mirrors (COTS-BB1-E02, COTS-POLARIS-K1), to re-center the beam with respect to the objective lens such that the beam is well-centered on the mark.
369. Mount and image a fluorescent dye sandwich sample using the multi-mode laser engine. Refer to: Fluorescent dye sandwich in the reagent setup section for a detailed description of the sample preparation.
370. Observe the shape of the illumination field. Focus the fiber in its mount (COTS-SM1ZM) to maximize the uniformity of the square illumination field. TROUBLESHOOTING! If the beam is not uniform but was previously uniform when viewed on the camera, this signifies clipping of the beam by the back aperture of the objective lens. Return to earlier alignment steps as appropriate.
371. Partially close the slit using the 4 adjusters to sharply truncate the illumination field at the periphery of the square region to achieve maximally uniform illumination.
372. Further rotate the fiber and subsequently realign (following steps 363 - 365) as necessary to better align the square fiber image to the slit.
373. Switch back and forth between the illumination inclination control mirror/stage (COTS- MRAE02, COTS-ELL17-M stage) positions for the single- and multi-mode laser engine illumination to check that the centers of the two illumination fields are well aligned to within ± 2 µm (ca. 20 pixels). TROUBLESHOOTING!: If the two fields are not fully aligned. Open the slit (COTS-SP60) and use the upstream steering mirror in the multi-mode path to reposition the square illumination region with respect to that of the single-mode laser. Repeat the process of truncating the square region with the slit as previously described.

### OPTIONAL: Calibration of the laser power monitoring photodiode (Timing: 0.5 days)

Having an active readout of the laser power is recommended and helpful for two primary reasons. Firstly, to monitor the proper functioning and internal alignment of the illumination lasers. For example, the Toptica iChrome MLE laser engine included as part of the single-mode illumination system features an automated scheme for internal realignment, which can boost the output power greatly depending on the degree of misalignment. Secondly, for reproducible blinking kinetics, it is helpful to be able to achieve a defined intensity (typically stated in kW/cm^2^), which can be calculated for a known transmission through the objective lens and measured illumination field size. The photodiode was installed previously but with all lasers in place it is necessary to calibrate the photocurrent generated vs. laser power.

374. Remove the objective lens from the objective port (FAB-EMBL-000043) and replace it with the photodiode sensor (COTS-S121C) of the laser power meter (COTS-PM100D) coupled to the port with a thread adapter (COTS-SM1A4).
375. For each laser wavelength, calculate the product of the maximum laser power available at that wavelength and the responsivity of the laser power monitoring photodiode (COTS-SM1PD1A) at the wavelength. In the case of a single-mode booster laser, consider the combined output of it and the corresponding line of the single-mode laser engine. Since multi-mode and single-mode illumination is not used in tandem, do not add sources together and rather consider them separately. Determine which laser line will thus produce the highest signal on the photodiode at maximum power.
376. Illuminate with this laser line at full power (combining single-mode sources as above) and check the voltage on the AnalogInput3 channel in µManager. The microFPGA will convert the voltage to an unsigned 16-bit value (0 - 65,535). Check that the voltage does not saturate, if it does halve the laser power and recheck. Continue halving until the voltage does not saturate. Now increase the laser power until the detector approaches saturation (> 80%, above ca. 52,500). If the maximum laser power does not approach saturation of the detector (> 80%) determine the increase in load resistance that would be necessary to do so. Otherwise determine the decrease in load resistance that would be necessary such that at maximum power approaches saturation of the detector (> 80%). Modify the resistance at R2_G on the ACB by the determined factor (or as close as possible given discrete resistance values) without resulting in saturation.
377. Check the measured signal again on the AnalogInput3 channel in µManager for maximum laser power and repeat the above steps as necessary to achieve the optimum load resistance.
378. For each laser line, vary the laser power stepwise between the minimum and maximum in 10 steps, recording the AnalogInput3 channel and the reading on the laser power meter (COTS- PM100D). CAUTION!: Ensure that the laser power meter is configured for the calibrated laser wavelength each time.
379. For each laser line, plot the 16-bit value from the AnalogInput3 channel (x) against the reading on the laser power meter (y) to determine the offset and gradient from a linear fit. Input the offsets and gradients as comma separated values in ‘Powermeter - offsets’ and ‘Powermeter - slopes’ respectively in the EMU properties in µManager. The values correspond to lasers 0,1,2,3 respectively.

### 2D imaging quality checks and pixel size calibration (Timing: 1 day)

Having set up the microscope with functional emission and illumination paths, it is necessary to perform various checks and calibrations. The 2× or 2.85× telescopic relay is typically used to produce the largest possible field of view and thus the most thorough test of image quality. Furthermore, since the potential for crosstalk between the two channels is greatest, the large field of view provides the limiting case for setting of slits to restrict the spatial extent of the illumination source and resulting images. The image quality and slit positioning is assessed using a fluorescent bead sample, which also provides the basis for calibrating the effective pixel size of the camera.

380. Switch to the highest magnification telescope available (typically 2× or 2.85× depending on system configuration).
381. Mount and image a fluorescent bead sample. Refer to: Fluorescent bead samples in the reagent setup section for a detailed description of the sample preparation and: Focusing the microscope for a description of setting up the microscope to image fluorescent beads. Ensure that the bead sample is immersed in water.
382. Introduce the emission path slit (COTS-SP60, COTS-RS2P4M, COTS-KB25-M, COTS- RS05P4M) in its nominal position and position it such that the image truncation arising from the slit blades appear in sharp focus in the image. Secure the slit in place using a pedestal post clamp (COTS-PS-F). The assembly should appear as shown Supplementary Figure 54.
383. Close the slit blades from the left and right and check for the sharpness on both sides of the image to ensure that the slit is oriented perpendicular to the emission path optical axis. Adjust alignment as necessary and fully open the slit blades.
384. View the fluorescent bead sample in both channels, adjusting contrast as necessary. Close the slit with the left/right blade such that the image is truncated from the center of the camera outward in both directions (since the channels are mirror images). Close the slit only minimally until there is no crosstalk/spatial overlap between the two channels.
385. Acquire a z-stack of the fluorescent bead sample (over a range of ± 1 µm, 0.05 µm spacing) using the µManager multi-dimensional acquisition wizard.
386. Use ImageJ to load the image stack, set the voxel dimensions in (Image/Properties, 0.105, 0.105, 0.05 µm, height, width, depth). Reslice the stack (Image/Stacks/Reslice) with start at Top selected and no other boxes checked. Make a maximum intensity projection of the resliced stack (Image/Stacks/Z Project) to view a projection of the beads as seen from the side. Check the PSF symmetry in z (now the vertical direction in the resulting image). Adjust the correction collar and repeat this process as appropriate. Repeat this process, with start at Left selected to produce an orthogonally oriented projection.
387. Check the two resliced/projected z-stacks for uniformity of the PSF. For oil objectives, it is possible that some non-uniformity is visible towards the FOV periphery (particularly for the largest FOV defined by use of the 2.85× telescope) but the PSF should look uniform otherwise. For the silicone objective lens (maximum compatible telescopic magnification of 1×), it is likely that the PSF will show coma aberrations in one or both views (the PSF will have a curved banana-like shape). TROUBLESHOOTING! For the oil immersion objectives if the PSF does not appear spatially uniform up to at least a IZ50 µm field. It is likely that either the lens is damaged or that the emission path has not been aligned correctly. Recheck the beam from the alignment laser traverses the emission path correctly, including good centration on mirrors and lenses and return to step 153 to realign the emission path as necessary. For the silicone oil objectives, it is more difficult to make assertions regarding the expected image quality and the builder is recommended to test the alignment with a regular oil immersion objective lens. In either case it is possible to check whether aberrations arise from the objective lens by carefully unscrewing the lens to reorient it (refocusing as necessary) and checking whether the deformed PSF rotates in accordance.
388. Position a single fluorescent bead in the center of the FOV and acquire an image, while recording the XY stage position from its controller. Move the bead to the peripheries of the FOV in X (e.g., for a ± 35 µm range when using the 2× magnification telescope). Acquire images at these positions, again noting the XY stage position.
389. Repeat this process for steps in Y.
390. Open the images in ImageJ and either manually follow the position of the bead or determine the bead center analytically (e.g., by applying a threshold to the images and determining image centroid for a bounding box including the bead). Calculate the bead displacement in each case (in pixels) and use the recorded stage position to calibrate for nm per pixel for both the X and Y axes. The pixel size should be ca. 105 nm in X and Y (assuming a system magnification of 61× and a camera pixel size of 6.5 µm)
391. Record the X and Y pixel size for later input in the SMAP Camera Manager.
392. Switch back to the 1× magnification telescope.

### 3D imaging alignment (Timing: 2 days)

To perform 3D imaging, the astigmatic lens must be introduced to the emission path, aligned and adjusted to provide an appropriate amount of astigmatism. The astigmatic lens consists of a pair of cylindrical lenses of equivalent power and opposite sign. Introducing a relative rotation between the zero-power axes of the lenses results in a variable amount of astigmatism being introduced, which allows the axial position of emitters to be encoded via the resulting lateral PSF asymmetry and orientation. The astigmatic lens is placed at a conjugate of the back focal plane of the objective and is thus expected to introduce minimal distortion associated with anisotropic magnification. The correct placement of the astigmatic lens is assessed by imaging an aperture target onto the camera with the help of the Bertrand lens, which has been pre-configured to image the back focal plane and its conjugates. The alignment of the astigmatic lens proceeds via use of the alignment laser. The lens placement is further assessed by measuring the magnification anisotropy and minimized by repositioning the lens as necessary.

393. Construct the astigmatic lens assembly (3D-SMLM-EMBL-001-002-001) but omit the cylindrical lenses (COTS-LJ1516RM, COTS-LK1002RM).
394. Remove the objective lens from the objective port (FAB-EMBL-000043) and replace it with the alignment laser (3D-SMLM-EMBL-007).
395. Install 2× irises (COTS-SM1D12D) on the rotational mount (COTS-CRM1T-M) of the astigmatic lens assembly separated by a slotted lens tube (COTS-SM1L30C).
396. Move the piezo resonant stage (COTS-ELL20-M) to 50 mm and secure the assembly in its nominal position via the base plate (FAB-EMBL-000067) as shown in Supplementary Figure 55.
397. Use the stage position and height of the optical post assembly (COTS-PH30-M, COTS-TRA20- M) to align the beam to the downstream iris and rotate the two optical posts in their holders (COTS-TRA20-M/COTS-TRA40-M in COTS-PH30-M/COTS180-M respectively) to align the beam to the upstream iris. Turn off the alignment laser.
398. Taking care not to reposition components, remove the 2× irises (COTS-SM1D12D) and slotted lens tube (COTS-SM1L30C) from the rotational mount (COTS-CRM1T-M).
399. Place a cage alignment target (COTS-CPA2) in between the two rotation mounts (COTS- CRM1T-M, COTS-CRM1PT-M) of the assembly and supported by the 2× cage rods (COTS- ER1).
400. Use two fixed kinematic stops (COTS-KL01L) to indicate the position of the short side of the astigmatic lens assembly base plate (FAB-EMBL-000067) closest to the optical table edge.
401. Switch the Bertrand lens into the emission path.
402. Switch the filter wheels (COTS-FW102C) to an open position for transmission imaging.
403. Backlight the target with a white light source and image it on the sCMOS camera via the second relay lenses (COTS-AC254-200-A) and Bertrand lens (COTS-#32-963). TROUBLESHOOTING! If the aperture does not appear on the camera image, check the alignment of the alignment laser through its irises and return to realign from step 395. Note, that the magnification from the target to the camera is 0.3×, and the target aperture is 5 mm. As such, the image of the target should be 1.5 mm in diameter or ca. 230 pixels in the camera image.
404. Loosen the screws securing the base plate (FAB-EMBL-000067) and translate the assembly along path defined by the fixed kinematic stops (COTS-KL01L), such that the aperture appears as sharply defined as possible in the camera image. Secure the screws when the correct longitudinal position has been found.
405. Remove the cage alignment target (COTS-CPA2).
406. Install the cylindrical lenses (COTS-LJ1516RM, COTS-LK1002RM) in their respective rotation mounts (COTS-CRM1T-M, COTS-CRM1PT-M). Note, some disassembly of the 30 mm cage section of the astigmatic lens assembly (3D-SMLM-EMBL-001-002-001) will be required.
407. Repeat steps 395 - 398 to realign the assembly to the alignment laser in the corrected longitudinal position. Record the final piezo resonant stage position. Save this value in the EMU configuration for the Two-state device 1 - On value. Set the Two-state device 1 - Off value to 0, such that turning on/off the 3D control moves the astigmatic lens in and out of the beam path.
408. Remove the alignment laser (3D-SMLM-EMBL-007) from the objective port (FAB-EMBL- 000043) and replace it with the objective lens.
409. Mount and image a fluorescent bead sample and set the 3D property to off, checking that the stage repositions to move the cylindrical lenses out of the emission path. Refer to: Fluorescent bead samples in the reagent setup section for a detailed description of the sample preparation and: Focusing the microscope for a description of setting up the microscope to image fluorescent beads
410. Set the 3D property to on and observe the PSF shape when focusing through the beads.
411. Carefully rotate the upstream rotation mount (COTS-CRM1T-M) until the PSF appears as close to the 2D case (i.e., without the astigmatic lens) as possible. This corresponds to a position where the cylindrical lens axes are aligned and the astigmatism introduced is negligible.
412. Rotate the downstream rotation mount (COTS-CRM1PT-M) using the micrometer screw to introduce mild astigmatism such that the PSF at focus appears as a faint cross. Note, it may be necessary to refocus slightly as the amount of astigmatism is adjusted.
413. Focusing up and down should result in a PSF that extends in one axis on one side of the focus and in an orthogonal axis on the opposite side. Carefully rotate the upstream rotation mount (COTS-CRM1T-M) and compensate with an equal motion of the downstream cylindrical mount (COTS-CRM1PT-M) to achieve a similarly mild astigmatism but such that the extension axes are aligned to the pixel grid of the camera. Note, for large movements of the downstream rotation mount, manually adjust via the mount itself rather than using the micrometer, which should remain close to the center of its travel range.
414. Repeat the process of calibrating and recording the X and Y pixel size from steps 388 - 391

### Focus lock path build-up and alignment (Timing: 2 days)

To be able to perform SMLM imaging experiments and assess drift/vibrations, it is necessary that the focus lock system is installed. Setting up the focus lock system requires aligning a near-infrared (808 nm) laser to the optical axis of the objective lens, focusing it to the back focal plane and then shifting the focused spot to the periphery of the back focal plane to fulfil the condition for total internal reflection. The reflected beam is subsequently routed to the QPD where an axial drift of the objective-coverslip spacing is detected as a lateral shift of the reflected beam. When setting up the microscope with a silicone oil objective, start with 20× higher resistance to provide higher detector sensitivity. In the case of a silicone oil objective lens, the refractive index contrast between the immersion oil and the sample is too small to fulfil the total internal reflection condition at the angle defined by the maximum NA of the objective lens. In this case, one can still focus lock via Fresnel reflection of the tilted beam at the interface. In this case, the reflected signal will be much dimmer than for total internal reflection. The focus lock path needs to be constructed in a stable manner with minimized beam path lengths since any drift in the beam pointing will result in focus drift. A CAD rendering of the fully assembled focus lock path is shown in Figure 6.

**Figure 6.**
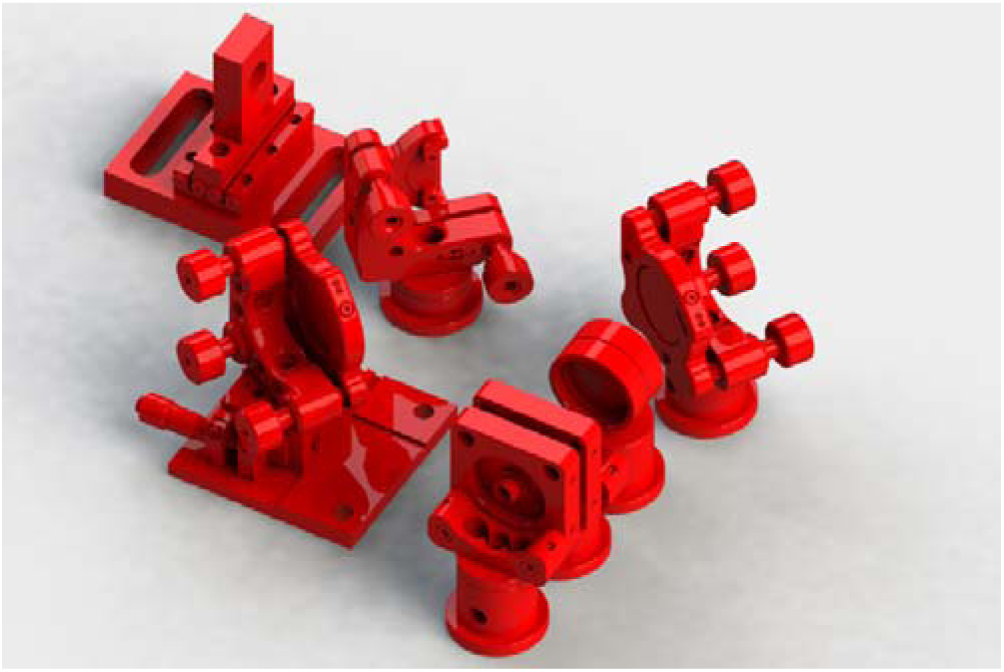
The fully assembled focus lock path. A fiber coupled NIR (808 nm) laser is connected to the fiber terminating adapter (bottom) and focused by a lens (occluded) onto the objective back focal plane (BFP) via a pair of steering mirrors. The second steering mirror is on a manually adjustable translation stage to offset the beam at the BFP to achieve the condition for total internal reflection at the coverslip sample interface (for oil objectives) or to optimize for Fresnel reflections (for silicone oil objective lenses). The returned beam is picked off by a D-shaped mirror and captured on a quadrant photodiode (QPD). The QPD measures focal drift as a positional offset of the returned beam and sends a corrective signal to the objective lens piezo flexure stage accordingly. Mounting the QPD on a motorized stage allows refocusing during focus lock operation by translating the QPD.

415. Remove the objective lens from the objective port (FAB-EMBL-000043) and replace it with the objective port alignment target (3D-SMLM-EMBL-006) with the beam profiling camera installed on the c-mount port at the distal end.
416. Install the focus lock laser (COTS-IBEAM-SMART-PT) on its mount (FAB-EMBL-000088) and install the mounted laser on the optical table. The precise location is arbitrary but make sure that it is placed close enough to the left side of the microscope body where the focus lock system is situated to avoid damage to the fiber.
417. Assemble the focus lock laser beam launch (3D-SMLM-EMBL-001-004-001) and secure it in position on the optical table using 2× pedestal post clamps (COTS-PS-F). Install the fiber from the focus lock laser in the terminating adapter (COTS-SM1FCA).
418. Use the beam viewer card (COTS-VRC2) to view the 808 nm beam and collimate the laser as best as possible by visual inspection by translating the lens (COTS-AC127-019-B-ML) relative to the fiber tip. CAUTION! The 808 nm laser is not visible and so presents an acute laser safety hazard, wear appropriate laser safety eyewear when aligning the laser.
419. Assemble the focus lock beam offset mirror assembly (3D-SMLM-EMBL-001-004-002) and install it in place on the optical table. Adjust the mirror mount (COTS-POLARIS-K1) to the nominal orientation/position and the translation stage (COTS-MS1S-M) to 6 mm as readout via the micrometer screw.
420. Introduce the first steering mirror (COTS-POLARIS-K1, COTS-BB1-E03, COTS-RS1P4M) with the beam centered on its aperture such that it reflects onto the center of the downstream mirror. Secure it in place with a pedestal post clamp (COTS-PS-F). The assembly should appear as shown in Supplementary Figure 56.
421. Use the steering mirrors to align the beam into the microscope body and direct it through the two irises (COTS-SM1D12D) of the objective port alignment target (3D-SMLM-EMBL-006), using the beam profiling camera to assess centration of the beam at each iris. Note, it may be necessary to reorient the downstream mirror on the stage by releasing the mirror mount (COTS-POLARIS- K1) and securing it once it has been appropriately rotated.
422. Remove the objective port alignment target (3D-SMLM-EMBL-006) from the objective port (FAB-EMBL-000043) and replace it with the objective lens. CAUTION! When the objective is in place and the focus lock laser is emitting, a highly divergent beam will exit the objective lens and even small misalignments may result in a strong deviation from the optical axis and constitute a laser safety hazard accordingly. Wear laser safety eyewear whenever assessing alignment through the objective lens and use the beam viewer card (COTS-VRC2) to check the path of the beam out of the objective lens.
423. Using the beam viewer card (COTS-VRC2) to view the beam out of the objective, focus the lens (COTS-AC127-019-B-ML) of the beam launch to collimate the output by minimizing the spot size far from the front of the objective lens. Secure the cage plate (COTS-CP32-M) mounting the lens in place. CAUTION! When collimated, a minimally divergent beam will exit the objective lens and even small misalignments may result in a strong deviation from the optical axis and constitute a laser safety hazard accordingly.
424. Check the path of the focus lock laser out of the objective. If the laser is launched substantially off-axis (no longer incident within ± 10 mm of the mark made in step 252 as viewed using the beam viewer card, COTS-VRC2), return to realign the beam through the objective port alignment target (3D-SMLM-EMBL-006) without adjusting the laser focus. TROUBLESHOOTING!: If the laser cannot be aligned to launch the beam close to the objective lens axis, use the first steering mirror to reposition the beam such that the laser aligns to the mark made in step 252.
425. Mount and image a fluorescent bead sample to establish the focal position. Refer to: Fluorescent bead samples in the reagent setup section for a detailed description of the sample preparation and: Focusing the microscope for a description of setting up the microscope to image fluorescent beads. Periodically adjust the focus as necessary.
426. Ensure that the cover is correctly installed on the microscope to intercept obliquely launched beams.
427. Translate the translation stage (COTS-MS1S-M) toward the front of the microscope such that the beam travelling parallel to the objective optical axis is translated towards the front side of the objective lens back focal plane. CAUTION! Avoid translating the stage towards the back of the microscope, which will result in the beam launching obliquely out of the optical table on the proximal side.
428. While translating the stage, check for total internal reflection using the beam viewer card (COTS-VRC2) to see the reflected beam spatially offset by 4 - 5 mm from the input beam towards the back side of the microscope, ensuring that the card does not block the input beam. Translate the stage to maximize the power of the reflected beam, which will be substantially dimmer than the input (particularly for the reduced refractive index contrast present when using a silicone oil objective). Note that the beam viewer card will dim with repeated exposure and needs to be charged by e.g., room lights. TROUBLESHOOTING!: If the reflected beam cannot be seen, use the Bertrand lens to confirm that the focus lock beam is focused at the periphery of the back focal plane. With great care one may also remove the cover of the microscope to check the beam angle out of the objective lens. CAUTION! The focus lock laser will propagate out of the objective lens at an oblique angle. Make sure to intercept the beam with the beam viewer card. When translating the stage correctly, the laser spot should be launched at ever more oblique angles until it can no longer be seen emerging from the objective when the total internal reflection condition is met. In the case of difficulties aligning the focus lock system using a silicone objective lens, it is highly recommended to complete the alignment with an oil objective first.
429. Install the pickoff mirror (COTS-PFD10-03-P01, COTS-KM100DL, COTS-RS4M, COTS-RS05P4M), such that the input beam passes into the microscope body, while steering the reflected beam parallel to the hole pattern along the long-edge of the optical table. Note, the beam will be diverging. Secure the mirror in place with a pedestal post clamp (COTS-PS-F).
430. Assemble the QPD translation stage assembly (3D-SMLM-EMBL-001-004-003) but omit the QPD itself (COTS-SD197-23-21-041).
431. Center the QPD stage (COTS-2445-L) in its travel range.
432. Install the QPD translation stage assembly (3D-SMLM-EMBL-001-004-003) in place on the optical table such that the returned focus lock beam passes centrally through its aperture as judged using the beam-viewer card (COTS-VRC2).
433. Install the QPD (COTS-SD197-23-21-041) in its mount (FAB-EMBL-000091). Re-center the QPD stage (COTS-2445-L) in its travel range if necessary. The assembly should appear as shown in Supplementary Figure 57 (QPD omitted).

### Setup of the focus lock amplifier (Timing: 1 day)

Measuring the focus lock QPD signal on the microFPGA aids in determining a correct working voltage range for the feedback to the objective piezo flexure stage (COTS-P-726.1CD). The conversion of the photocurrent from each of the QPD quadrants to a voltage is handled by the QPD amplifier board but requires the installation of four resistors (of equal resistance) to provide an output in the range 0 - 10 V. The focus lock laser is attenuated by a neutral density filter (ND 1), allowing it to be operated at 30 – 50% of its total power, where it typically is most stable, while reducing the incident laser light on the objective and sample. Having set the resistance appropriately, the QPD is translated on its stage and two potentiometers adjusted to appropriately scale the positional output to the range 0 - 10 V as required by the analog conversion board of the microFPGA and the objective piezo flexure stage. The values for resistors provided below assume the use of an oil objective. When setting up the microscope with a silicone oil objective, start with 20× higher resistance.

434. Package the QPD amplifier board as desired (the following assumes that 3 SMB female connectors are provided for the X, Y and SUM signals). Provide an appropriate power supply to the board and make connections to the QPD (COTS-SD197-23-21-041). The two LEDs should be on.
435. Connect the X signal SMB connector of the QPD amplifier to an appropriate SMB T-splitter and connect one of the T-splitter arms to the analog input BNC connector of the Physik Instrumente E-709 controller. Connect the other to the AI:QPD:X connector on the microFPGA.
436. Connect the Y and SUM SMB connectors of the QPD amplifier to AI:QPD:/Y/SUM connectors on the microFPGA.
437. Install 4× 51 kΩ resistors on the amplifier board (in the positions indicated in the board datasheet). Set the focus lock laser power to 30 mW. Check again for return of the focus lock laser using the beam viewer card (COTS-VRC2).
438. Install the focus lock laser ND 1 filter (COTS-NE10A-B, COTS-LMR1-M, COTS-RS1P4M) and secure it in place with a pedestal post clamp (COTS-PS-F). Note the filter can be unscrewed from its mount as necessary for realignment of the focus lock system (Supplementary Figure 58).
439. Check the LEDs on the QPD amplifier board. If both LEDs are off, the incident intensity is too low and the resistors should be exchanged for those with higher values (the amplification is linearly proportional to the resistance). Conversely, if both LEDs are on, the resistance should be decreased. Establish a resistance for which only one LED remains on over a range 5 - 50 mW.
440. Check the SUM signal in the QPD tab of the EMU GUI. adjust the laser power to bring the green bar to approximately half of its full height.
441. Connect the X channel of the QPD amplifier to an oscilloscope. Cover the QPD so that no light is incident on it. Adjust the X and Y offset potentiometer to adjust the signal to 5 V.
442. Uncover the QPD and translate the QPD stage (COTS-2445-L) back and forth, using the LEDs to check that the beam is still incident on the QPD and note the change in the X voltage on the oscilloscope. Use the X gain potentiometer to scale the signal to the range 0 - 10 V (or as close as possible) across the range of motion. Note, if translating the stage does not elicit a change in the X signal, check the Y signal as well. If the Y signal shows a response, then the X and Y channels have been switched at some point in assembling and connecting the electronics.
443. Repeat the process for the Y voltage and Y gain potentiometer, instead using the D-shaped steering mirror mount (COTS-KM100DL) to move between the Y extremes. Recenter the red point on the 2D graph display in EMU.
444. Connect the X and Y signals as necessary to the microFPGA front panels and again monitor the signals in EMU. Center the QPD stage (COTS-2445-L) in its travel range.
445. Use the D-shaped steering mirror mount (COTS-KM100DL) to center the red point on the 2D graph display.

### Focus locking (Timing: 1 day)

Having aligned the focus lock system and setup the amplifier, it is now possible to engage the focus lock and check that the microscope can be focused by moving the QPD stage.

446. Bring the fluorescent bead sample into focus in the image.
447. Ensure that the SUM signal from the focus lock laser is sufficient to half fill the green bar in the QPD tab of the EMU GUI. Adjust the focus lock laser power as necessary. Center the focus lock laser spot on the QPD (X axis) by translating its stage.
448. Engage the focus lock (‘Lock’ in the EMU htSMLM GUI). Note, that the focus lock will only work over a z-range for which the laser spot remains incident on the QPD. Engaging the focus lock with the focus lock laser spot not visible on the display, will lead to a sudden drift of focus typically away from the true focal position (as viewed from the z stage position). TROUBLESHOOTING! If the focus begins rapidly shifting (as viewed from the z stage position) after engaging the focus lock, it is likely that the piezo controller parameters have been set incorrectly. Recheck that the parameters are as described in Supplemental Table 7.
449. Adjust the locked focus position by repositioning the QPD stage (10 – 100 µm increments) such that the beads move in and out of focus. Return the microscope to a sharp focus using the QPD stage.

### Mode-hopping and amplifier tuning (Timing: 2 days)

The focus lock diode laser may undergo a process of transient mode-hopping depending on the current flow and temperature of the unit. This hopping may result in instability of the focus lock system. The laser will have stable regions defined by the set laser power and other performance-stabilizing parameters. To minimize the impact of mode hopping, first the stability of the z-position is checked by viewing the reflected beam on the emission path camera to identify mode hops and correlating with the reported objective piezo position. Secondly a usable parameter space of laser settings, which eliminate or suppress mode-hopping, is determined.

450. Set up the microscope for focus lock operation as previously described. Reduce the focus lock laser power to the lowest setting eliciting a stable single LED response. Switch off all imaging lasers and set the transmitted filter wheel and shared path filter slider (if present) to an open position.
451. Ensure the FINE property is switched off in the focus lock tab of the EMU htSMLM GUI.
452. Remove the NIR laser blocking filter from the emission path and/or the neutral density filter (COTS-NE10A-B) from the focus lock path if the focus lock laser is not visible on the emission path sCMOS camera. The focus lock laser should now be visible on the camera. Adjust the exposure time such that the counts are < 10% of the saturation threshold of the camera.
453. Capture sequences of images at 10 frames per second for 5 minutes, sequentially increasing the focus lock laser power in steps of 5 mW at a time. When increasing the laser power make sure not to saturate the QPD.
454. Mode-hopping is visible as an instability in the beam profile. During periods of visible mode-hopping, look for instabilities in the objective piezo monitoring signal (saved in the metadata as ‘PIZStage-Position’) using the plugin ‘Process/Images/DisplayImageTagsTiff’ in SMAP to view the z-focus position for each image sequence. If the instability is on the order of ± 10 nm (the resolution of the piezo flexure stage), the mode-hopping has negligible influence on subsequent imaging.
455. If larger mode hops are observed for certain laser operating powers (and laser operating temperature), establish a laser power range that can be safely used without mode hops and use this range for future operation of the focus lock system. TROUBLESHOOTING!: If it is not possible to prevent mode hops to the extent necessary for optimal focus locking, it is possible to scramble the output modes using the FINE feature of the focus lock laser. To do so, switch the FINE property on starting with settings of FINE A = 90 %, FINE B = 10 % recheck the instability as before. Try small variations in these values to minimize mode hops and seek advice from the laser manufacturer.
456. Once an appropriate operating range of laser power has been established, set the QPD amplification (by exchanging resistors) such that the lower power limit of this range results in ca. 25 % saturation of the QPD signal. This lower limit provides an optimal operating power given considerations of laser lifetime, safety and stable operation. Note, that the signal strength may vary for cases of smaller refractive index contrast between the sample and coverslip and as such, it may be necessary to re-optimize for these cases.
457. Reinstall the focus lock neutral density filter if it was previously removed from the focus lock path.
458. Assuming it was not previously removed (COTS-NE10A-B), release the NIR laser blocking filter from its mounting lens tube (COTS-SM1L03). Install the filter in the exposed SM1 port of the sCMOS camera c-mount adapter (FAB-EMBL-000073) securing it in place with a retaining ring (COTS-SM1RR). Note to install the filter as such, it is necessary to remove the Bertrand lens stage on its pinned mount (FAB-EMBL-000074), which can be replaced once the filter is secured. Installing the blocking filter in front of the camera is necessary to prevent a NIR LED in the Bertrand lens stage (COTS-ELL6) causing an artificially high background on the camera for the localization imaging that follows in the protocol.
459. Depending on the specific choices of spectrally discriminative optics in the system it may be possible to see the focus lock laser spot on the camera. To check, set the focus lock laser to the highest operating power and set the sCMOS camera exposure time to 200 ms or the longest expected exposure time for operation. If the laser spot is visible above the background level of the camera (ca. 100 counts), install the focus lock laser cleanup filter (COTS-FB810-10) in a lens tube (COTS-SM1L03) and install onto the focus lock laser neutral density filter (COTS-NE10A- B). This blocks out of band emission from the focus lock laser. Note, the cleanup filter bandwidth is narrow (10 nm full-width at half maximum) and so the edges are close to the nominal laser wavelength (808 nm). As such, small rotations of the filter with respect to the incident light may result in partial or complete extinction as the edges move to lower wavelengths. Appropriate rotation of the optic via the post mounting the neutral density filter (COTS-RS1P4M) should allow the laser line to pass while maximally blocking lower wavelengths. Once out of band emission has been blocked, the focus lock laser spot should no longer be visible on the sCMOS camera image.

### Setup and configuration of software for analysis (Timing: 1 day)

Analysis software is required to determine the positions of the single-molecule activation events in the camera frames to then reconstruct a super-resolution image. There are various complete software packages for this task [56–58] that are compatible with this microscope. Here we detail the installation of SMAP [28], developed in our group, that provides additional functionality to characterize the microscope. In the following sections, we then describe how to use SMAP for single-molecule fitting and analysis. For a more detailed description, please consult the User Guide in the Help menu of the SMAP window.

460. If you have access to Matlab (R2022a or newer) including the toolboxes: Optimization, Image processing, Curve fitting, Statistics and Machine Learning, we recommend installing the Matlab version of SMAP from www.github.com/jries/SMAP, where you also find the installation instructions.
461. Alternatively, you can install the compiled stand-alone version of SMAP following the link on www.github.com/jries/SMAP.
462. Start SMAP.m in Matlab or the SMAP application when using the standalone version. Consult ‘Getting started’ in the ‘Help’ menu to familiarize yourself with SMAP. Here you also have access to the User_Guide.
463. Install Micro-Manager 1.4.22 or later from https://micro-manager.org. Note that this is necessary even if you have Micro-Manager 2.x installed, as only version 1.4 is compatible with SMAP.
464. As described in the User_Guide, Chapter ‘2.2 Installation’, add the Micro-Manager path to SMAP:Preferences:Directories.
465. Add the main camera to SMAP (see User_Guide Chapter ‘5.1 Camera Manager’ for details). First, duplicate the file cameras_Hamamatsu.mat in the SMAP/settings directory and rename it to e.g., cameras_local.mat. In the SMAP menu / Preferences… select the File tab and in the ‘CameraManager file’ field select the duplicated file. Press ‘Save and Exit’. Second, open the Camera Manager from the SMAP menu. With ‘load images’ load any image file that you acquired with your camera. When asked if to add a new camera click ‘no’. In the first list click (second line, next to the OrcaFusion name) in the ID column. A list will open with all metadata properties. Select a field that uniquely describes your camera, for instance HamamatsuHam_DCAM-CameraID. Save and close the Camera Manager. Do not add your local camera file to the git repository. Third, by clicking on the camera name ‘OrcaFusion’ you can check if the correct parameters are set: EMon: 0; cam_pixelsize_um: 0.105, 0.105 (you can overwrite these values with the correct pixel sizes that were calibrated in step 414); conversion: 0.24; offset: 100. These parameters are correct for the Hamamatsu Fusion BT camera for a system magnification of 61× provided by the optical scheme described. For other cameras and changes to the emission path optics, please amend these values accordingly.

### Performing PSF calibration (Timing: 3 days)

Having set up the focus lock system, it is now possible to perform an accurate 3D calibration for subsequent image reconstruction using fluorescent beads. Focus locking is necessary in this case, since the calibration requires that a large number of individual bead-derived PSFs are averaged, which in turn requires imaging over a large region of the coverslip, which may vary slightly in height. An iterative process of optimizing correction collar and astigmatism settings is performed to attain the best possible 3D imaging performance. Subsequently, we will describe how to perform a PSF calibration with SMAP and use this for 3D fitting. This procedure is used later to check for drifts and vibrations and to fit the example data for performance characterization.

466. Mount a fluorescent bead sample. Refer to: Fluorescent bead samples in the reagent setup section for a detailed description of the sample preparation.
467. Use 1% laser power to locate a region of the sample with ca. 5 - 20 beads that are well separated. Sharply focus the beads in the image. Acquire a z-stack from -1 µm to 1 µm around the focus with a spacing of 0.02 µm using the µManager multi-dimensional acquisition wizard.
468. Iterate this process, adjusting the correction collar and amount of astigmatism (as previously described) between iterations to achieve optimal 3D localization precisions over the desired axial range. It should be possible to achieve localization precisions better than 5/15 nm laterally/axially over a range of ± 400 nm as judged from the CRLB plot from the 3D calibration output. The minima of the CRLB x and y plots should be approximately 400 nm apart in z height for both the oil and silicone oil objectives. In general, stronger astigmatism is associated with improved z localization precision but also a larger distance between the x and y minima resulting in a reduced effective operating z range. Results from a typical 3D calibration are shown in Figure 7. Note, this procedure is only required the first time and after exchanging the objective, realigning aspects of the emission path or if the collar/astigmatism are adjusted for other reasons.
469. Once optimal settings have been determined, repeat the calibration step over 5 - 10 regions of interest (using the µManager Stage Position List to mark individual positions) to acquire several bead stacks for a more precise 3D PSF model. To this end, adjust the laser power to optimize between good signal-to-noise ratio and low bleaching and low flickering of the fluorescent beads. A good starting point is 0.5 kW/cm^2^. This process should be repeated for each imaging session.
470. Start SMAP and open the plugin: Analyze:sr3D:calibrate3DsplinePSF. Press Run.
471. Using ‘select Camera Files’ load the bead stacks you just acquired.
472. Set the distance between the frames to 20 nm. All other settings can be left at default. Press ‘Calculate bead calibration’.
473. Check the Results window (see Figure 7): Under ‘Files’ the beads should be identified, but no background noise (if beads are not identified reduce the ‘relative cut-off’ parameter, if background is identified as beads reduce it). In PSFx and PSFz the individual bead profiles should envelop the average. In ‘validate’ the central part should consist of parallel straight lines. ‘CRLB’ calculates the achievable localization precision for a hypothetical photon count of 5000 and a background of 50 photons. An asymmetry of the z localization precision about the plot’s z center (ca. z = 0 nm assuming that the z-stack is well centered on the focus where the elongation in either axis is at a minimum) often hints at the presence of spherical aberrations. A peak in the z localization precision at or close to z = 0 nm suggests that the astigmatism is too weak. The bead calibration is saved in the folder of the bead stacks with a file name ending in ‘_3Dcal.mat’.

**Figure 7.**
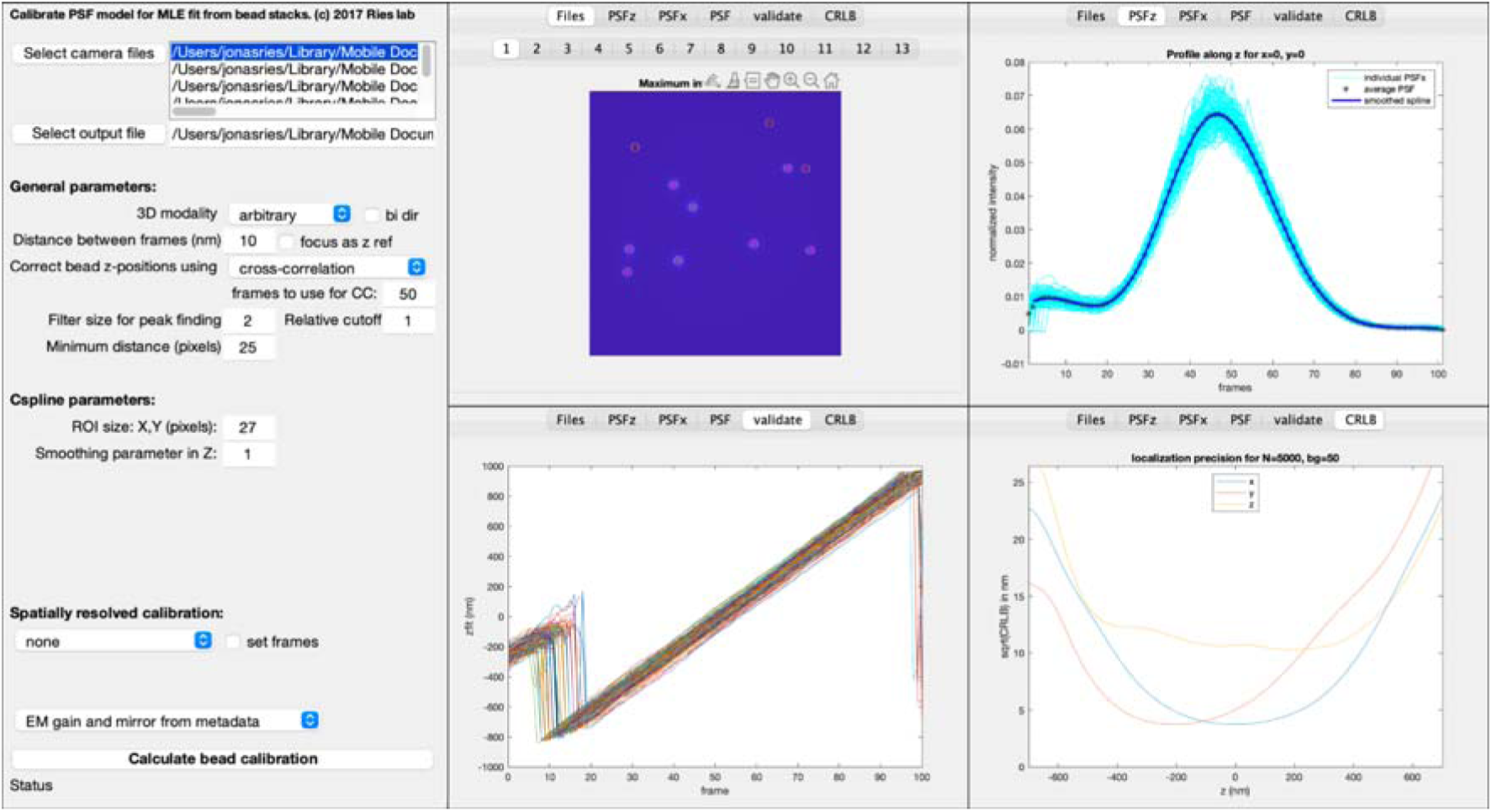
3D calibration of the microscope based on fluorescent bead z-stacks. The SMAP GUI with default parameters and examples of the results window for a suitable bead calibration. Left: interface for setting various fitting parameters. Top middle: a viewer for the detected beads in each z-stack. Top right: The spline fit to the averaged PSF along the z-axis. Bottom middle: The bead calibration is validated by comparing the fitted z position to the expected one (given by the frame). Bottom right: The 3D localization precision expected for a theoretical localization with N = 5000 photons and a background of 50 photons/pixel. The z range over which the x, y and z localization precisions remain acceptable defines the maximum depth of the 3D volume that can be imaged.

### 3D and 2D Fitting with SMAP (Timing: 1.5 day)

Here we describe how to use SMAP to fit imaging data. In 2D, single PSFs originating from beads or a biological sample, can be approximated by a 2D Gaussian function. To extract additional information such as the z-position and the color when using multiple channels an experimental PSF model is generated and used for fitting 3D and multi-color data (see globLoc [16] for more detail on generating a global PSF model for multi-channel data).

How to acquire these data is detailed below in the sections: 2D imaging of nuclear pore complexes (Timing: 5 days), 3D single-color imaging of nuclear pore complexes (Timing: 5 days) and 3D, multi-color imaging of nuclear pore complexes (Timing: 5 days).

474. In the SMAP main window select the ‘Localize: Input Image’ tab and make sure that the ‘fast_simple’ workflow is selected (otherwise load it with the ‘Change’ button).
475. With ‘load images’ load the Micro-Manager tiff stack that you want to fit (see section: Performing PSF calibration (Timing: 3 days) for instructions on how to acquire this data).
476. If the camera has not been added to the Camera Manager or if the camera is not recognized, set the camera parameters in the dialog ‘set Cam Parameters’. Afterwards check ‘lock’ to prevent this data from being overwritten.
477. In the ‘Fitter’ tab select ‘Spline’ as a PSF model and load the ‘*_3Dcal.mat’ file that you generated above with ‘Load 3D cal’.
478. Press ‘Preview’ (you can select the frame that is fitted for testing next to the ‘Preview’ button). True localizations should be marked, background noise should not. You can adjust the sensitivity of the peak finder in the ‘Peak finder’ tab by changing the value next to the ‘dynamic (factor)’ menu.
479. Once you are happy with the preview, press ‘Localize’ to start the localization process. Once the fitting is done, the results are automatically saved in the folder of the tiff files and a super-resolution image is reconstructed that can be explored.
480. For 2D fitting of data acquired without the astigmatic lens, follow the steps of this section with the following change: in the ‘Fitter’ tab select ‘PSFxy’ as PSF model, no calibration file is needed.
481. For dual-channel 3D fitting, first calibrate the dual-channel PSF as described above, but with the following modifications: As 3D modality choose ‘global 2 channel’. Under ‘global fit parameters’ check ‘make T’ and select ‘right-left mirror’ and main Channel ‘d/r’. The resulting calibration will be shown in a new window, and the Matlab file as well as the figure are saved automatically to the imaging folder. TROUBLESHOOTING! In the calibration figure, the results for each channel are shown (‘first channel’ and ‘second channel’). For each tab the quality of the calibration can be assessed as described in step 474. Two additional tabs can be inspected for dual-color data. In the ‘transformation’ tab, the transformation of channel 1 onto channel 2 is evaluated. Check ‘transformation:pos’ tab to check that most beads found in channel 1 are matched to a specific bead in channel 2. Beads for which no matching was possible are marked as ‘+’ and should be significantly fewer than matched beads marked as ‘o’. Ensure that most beads across the entire field of view are paired. The ‘CRLB’ and ‘validation’ for the global transformation are found in ‘global cal’. Similar as described in step 474, the ‘validation’ tab should show a central part with straight and parallel lines. ‘CRLB’ should show a symmetric localization precision for z, which can be expected to reach approximately 10 – 15 nm. Similarly, x and y localization precision should be symmetrical, and at a low value around 5 nm. In the PSF tabs, the cross-section through the PSF model along x (‘PSFx’), y (‘PSFy’), and z (‘PSFz’) are shown. The traces for PSFs calculated for individual beads (light blue) should follow the smoothed PSF model (dark blue) with minimal deviations. Should the spread between individual beads and the model be too wide, the bead calibration has to be optimized. In the ‘validation’ tab check that zfit for all beads is linear with the frame and that all lines are parallel to the zfit for the average model (black line). Deviations at the extreme frames are expected and result from the bead calibration spanning a larger z range than that which is useable. Experimental reasons for imperfect bead calibration include the presence of an air bubble when imaging, poor choice of bead regions such as overlapping beads, flickering and bleaching of beads because of high laser powers or saturation of the camera. It is thus recommended to re-mount the sample and acquire a fresh bead stack. Check if the model properly fit the individual beads (‘global cal’/’validate’) and that all curves fall onto the same linear relationship for most of the frames. Check the cross-section through the model along the z-axis to see if the model well represents the individual beads (‘global cal’/’PSFz’). Repeat for the model along the x-axis (‘global cal’/’PSFx’).
482. For fitting dual-channel data navigate in SMAP to the ‘Localize’ tab. Check if the workflow ‘fit_global_dualchanneĺ is selected. If not, use the ‘Change’ button to select the file ‘fit_global_dualchannel.txt’ to load this workflow.
483. Fitting is analogous to the steps described above but with the following differences: In the ‘Peak Finder’ tab select the PSF ‘*_3Dcal.mat’ file you created with ‘load T’. In the ‘Fitter’ tab select ‘Spline’ and load the same file again with ‘Load 3D cal’. Check ‘Global fit’. Make sure only x, y and z are checked and N and bg remain unchecked. Select ‘Ch1’ in ‘main xy’.
484. Start the fitting process with ‘Localize’.
485. After fitting assign the colors with the plugin ‘Process/Assign2C/Intensity2ManyChannels’ (press ‘Info’ for instructions). To render a specific color, you need to specify the color in the Render tab (0 is unassigned colors). You can add a layer to display the second color.
486. To fit 2D dual-channel data, you need to first generate a transformation file. For this, first fit both channels as described above for 2D data with the fast_simple workflow. Then use the plugin ‘Process/Register/Register Localizations’ and use as a target ‘right’ and ‘left-right’ at mirror. Press ‘Run’ and if the transformation is good save it with ‘save T’.
487. For fitting, follow the steps above (including changing the workflow to fit_gobal_dualchannel’, and in the ‘Fitter’ tab specify ‘PSF free’. In the ‘Peak Finder’ tab load the transformation file ‘*_T.mat’ you just generated.

### Post-processing and rendering with SMAP (Timing: 1.5 days)

The localization workflow discussed above extracts the coordinates of single fluorophores from the camera frames together with additional attributes, such as the localization precision or the likelihood as a measure for goodness of fit. In the following we discuss how to merge localizations in consecutive frames, stemming from the same fluorophore, into a single localization, how to render a super-resolution image, how to filter out poor localizations, and how to perform drift correction.

488. Load a fitted data set (‘*_sml.mat’ file) in the file tab using the ‘Load’ button. An overview image should appear in the SMAP window.
489. Localizations in consecutive frames stemming from the same fluorophore activation event are automatically merged (‘grouped’) into a single localization according to the parameters in the ‘File’ tab under ‘Grouping’. You can change the parameters and perform the grouping again by pressing the ‘Group’ button. You can later select whether to display these grouped localizations or the original frame-wise localizations with the ‘group’ checkbox in the ‘Render’ tab.
490. By pressing ‘Render’ in the render tab you can render a super-resolution image. You can zoom into the image with the mouse wheel or track pad or the ‘+’ and ‘-’ buttons in the ‘format’ area of the SMAP window and move around by right clicking in the new position in the super-resolution window, by dragging the image to a new location or by left clicking in the overview image.
491. You can change the appearance of the super-resolution image by adjusting the parameters in the ‘Render’ tab and display several layers as described in the User_Guide, Chapter ‘7.1 The render GUI’.
492. SMAP allows for versatile filtering of localizations that are used for rendering and further analysis. To access the filtering window of SMAP press the button ‘OV->filter’. In the list, you can select any property of localizations, look at the histogram and decide on minimum and maximum values (check ‘filter’ to apply the filter). The following filters are used regularly and are found directly in the ‘Render’ tab: a) localization precision to filter out poor localizations from dim fluorophores; b) z position to only display a slice in z; c) for 2D data instead the fitted size of the PSF can be used as a filter to filter out out-of-focus localizations for optical sectioning; d) the frames to filter out the beginning where localizations might be too dense, or the end that does not contain useable localizations; e) LLrel, a normalized log-likelihood measure that describes the goodness of fit to filter out localizations that were close to other localizations and thus poorly localized.
493. Even a stable microscope will show drift on the nanometer scale over the measurement time of minutes to hours. This drift can be corrected with the plugin: ‘Process:Drift:drift correction x,y,z’, which you also find in the ‘Process’ tab. SMAP uses an approach called ‘redundant cross-correlation’ that splits the data into equal windows in time and then compares super-resolution images reconstructed from each time window with those of all other time windows to estimate the displacement. From all pairwise displacements a drift trajectory is then calculated and used to correct the data. In the plugin you need to select if only to ‘correct xy-drift’ or also to ‘correct z-drift’. The main parameter you need to choose is the number of time windows called ‘timepoints’. More time points lead to a smoother curve, but if the data is sparse, at some point the noise increases. Press ‘Run’ to perform drift correction. Then check the tabs ‘dxy/frame’ and ‘dz/frame’. Except for outliers, the colorful curves should envelope the black curve. If the noise is too large, reduce the number of time windows. As drift correction is automatically applied, you need to press ‘Undo’ before performing the drift correction again with altered parameters.
494. To render 3D data in 3D you can use the plugin ‘Analyze:sr3D:Viewer 3D’.

### Performing focus lock, drift and vibration checks (Timing: 2 days)

At this juncture, the microscope is capable of focus locking and 3D imaging. Before continuing it is first necessary to validate the focus lock performance by checking the axial drift of a fluorescent bead sample over 24 hours. The measurement is then repeated to check for medium-term instability over 10 minutes and subsequently the presence of short-term vibrations is checked for a single bead over a few seconds. To do so, it is necessary to introduce water cooling and switch off the camera fan. The instructions provided assume the use of the base configuration Hamamatsu Fusion BT sCMOS camera.

495. Power down the sCMOS camera.
496. Install the water-cooling protection cap of the sCMOS camera to avoid ingress of water.
497. Provide suitable water cooling to the camera via the inlet/outlets and check for leaks once circulation commences.
498. Power up the sCMOS camera and turn off the fan with the DCAM API configurator.
499. Acquire camera frames continuously in the manufacturer supplied HoKaWo software and monitor the temperature of the circulating coolant and camera itself to ensure that the cooling provided is sufficient to support the thermal load of the camera. If the camera is able to maintain its -8 °C operating temperature over a period of 4 hours, the cooling is adequate. If the camera emits a constant tone during operation, the camera has overheated and image acquisition will cease. In this case, switch off the camera and resume with a higher load cooling system.
500. Close the HoKaWo software when the testing is complete.
501. Add the additional settings to the µManager presets as described in Supplementary Table 13: Micro-Manager Configuration Presets for Camera Cooling, in order that sensor cooling is set to its maximum setting with the sensor fan set to off. CAUTION! The sCMOS camera is now reliant on water cooling to maintain the functionality of its Peltier element. Check the sensor temperature in HoKaWo periodically.
502. Mount and image a fluorescent bead sample. Refer to: Fluorescent bead samples in the reagent setup section for a detailed description of the sample preparation and: Focusing the microscope for a description of setting up the microscope to image fluorescent beads.
503. Acquire bead stacks and generate the PSF calibration as described in the section: Performing PSF calibration (Timing: 3 days) steps 466 - 467 and 469.
504. Set up for focus lock operation and engage the focus lock as described previously.
505. Turn on the 3D property to move the astigmatic lens into the emission path.
506. To measure long-term drift, image the bead sample once every 5 minutes for 24 hours (289 time points) using the µManager multi-dimensional acquisition wizard. For this experiment, use the ‘normal’ imaging mode preset.
507. Check the objective piezo flexure stage (z stage) position over the 24 hours of imaging by displaying ‘PIZStage-Position’ using the plugin ‘Process/Images/DisplayImageTagsTiff’ in SMAP. Some motion of the stage will be apparent to correct for the drift between the coverslip and objective lens.
508. Check the associated images to make sure that the focus remains stable throughout.
509. To measure medium-term instability, repeat the drift measurement but instead image the bead sample continuously with an exposure time of 25 ms for 10 minutes.
510. For the two datasets, fit the bead time-lapse date for drift analysis as described in step 493 in the section: Post-processing and rendering with SMAP (Timing: 1.5 days).
511. Use the ‘Analyze:measure:Drift_Analysis’ plugin in SMAP to display the datasets over time. For the long-term drift measurements, drift values up to 200 nm / hour in x,y and 50 nm / hour in z are acceptable and can usually be corrected during post-processing. The drift observed generally results from changes in laboratory temperature, which is to be expected. However, look for sudden jumps of the position that indicate mode jumps of the focus lock laser (see section: Mode-hopping and amplifier tuning (Timing: 2 days), which are deleterious for imaging. For the medium-term instability measurements, peak-to-peak amplitudes up to 5 nm are acceptable. Larger instabilities often result from a failure to properly isolate the microscope from air currents. If this is the case, add additional enclosure around the microscope.
512. Move to another field of view and locate a single isolated bead, position the bead at the vertical center of the camera chip and crop to a suitable ROI (e.g., 24 × 24 pixel) to encompass the bead and its surroundings while maintaining a small ROI size to achieve the best possible frame rate and small dataset size.
513. Change to the ‘fast’ imaging mode preset and set the exposure time to 0.5 ms (thus setting a maximum frame rate of 2,000 frames per second).
514. Increase the laser power as necessary to ensure that the bead appears as bright as possible without saturating given the much shorter exposure time.
515. Use the µManager multi-dimensional acquisition wizard to image the single bead field of view for 10,000 frames at the maximum frame rate of the camera (define an interval of zero between image acquisitions in µManager).
516. Fit the bead time-lapse data for vibration analysis as described in steps 474 - 481 in the section 3D and 2D Fitting with SMAP (Timing: 1.5 day).
517. Use the ‘Analyze:measure:Vibrations’ plugin to display the position of the bead vs time and a Fourier analysis of the position. In this frequency spectrum you can identify the prominent vibration frequencies and by zooming in the position vs time plot one can estimate the amplitude of vibrations. Peak-to-peak amplitudes up to 5 nm are acceptable.

### 2D imaging of nuclear pore complexes (Timing: 5 days)

Having minimized drift and vibrations within the system, we can next benchmark the 2D imaging capabilities of the set up. Here, we want to test the effective resolution achievable on a biological sample. To this end, we require a well characterized and homogenous biological structure that can serve as a reference standard. The Nuclear Pore Complex (NPC), responsible for transfer of molecules across the nuclear membrane, has been thoroughly studied with complementary methods including cryogenic Electron Microscopy (cryoEM) [61]. It comprises ∼30 proteins, of which Nup96 is our preferred candidate for benchmarking. This protein is present in 32 copies within the NPC, and distributed symmetrically across the structure, with 16 copies forming the cytoplasmic ring, and 16 copies forming the nucleoplasmic ring. Each ring contains 8 corners formed by two Nup96 proteins. Two Nup96 proteins in one corner are 12 nm apart, two corners are 42 nm apart, the entire ring has a diameter of 107 nm and the two rings are separated by 50 nm. This 8-fold symmetrical distribution of Nup96 makes it an ideal candidate to estimate the resolving power of the set up, and resulting images and downstream computational analyses allow for the identification of potential issues and more targeted troubleshooting [49]. We note that for the following sections concerning biological validation of the microscope performance, the builder should refer to Supplementary Note 2: Selecting spectrally discriminative optics to aid in selection of appropriate filters and dichroics suited to the far-red ratiometric dSTORM scheme. Since we are acquiring 2D data, one should follow the instructions in section 474 - 480 for 2D fitting. Fitting can be performed after the acquisition has finished. Alternatively, it can be performed ‘online’ to assess the reconstructed image in real-time during acquisition. This early evaluation offers the ability to detect any issues or artefacts during imaging and make necessary adjustments or abort the acquisition and restart with optimized settings. To investigate image quality (both during and after acquisition) visual inspection can be complemented with a statistical analysis of the image. It is important to understand the acquired data quality to identify potential points for troubleshooting. The following steps will outline a brief quality assessment. For a more in-depth analysis, please refer to the SMAP documentation, which further covers the calculation of Fourier Ring Correlation based resolution, geometric analyses and calculation of the effective labelling efficiency.

518. Refer to: Biological sample preparation for a detailed description as guidance in carrying out the following sample preparations. Seed U2OS Nup96-SNAP cells that have been in culture for at least 2 passages onto a coverslip.
519. After incubating seeded cells for 2 days, perform fixation and permeabilization.
520. Perform NPC labelling targeting the Nup96-SNAP tag with SNAP-Surface BG-AF647.
521. Place the coverslip onto the sample holder and add imaging buffer. For prolonged imaging (up to 24 hours), fill the entire chamber of the magnet ring with imaging buffer and seal it using clean parafilm.
522. Set up single-color imaging by manually removing the image splitter dichroic from the emission path. (COTS-SB1-M, COTS-25.5×36x3MM-DICHROIC-SP, COTS-KM200S, FAB-EMBL- 000068/69).
523. Set the exposure time to 30 ms in µManager.
524. Place a drop of immersion oil onto the objective and then mount the sample on the microscope.
525. Set up EMU for imaging as follows: a) Monitor stage position; b) monitor Focus Lock; c) turn on Focus Lock laser; d) set 640 nm laser to 1% e) set appropriate emission filter in the transmitted path of the image splitter (typically for AlexaFluor647, this is a 676/37 bandpass filter).
526. Switch to live imaging stream and focus on the coverslip. Typically, some residual dye or other autofluorescent particles are present on the coverslip to aid in focusing.
527. Double check the back focal plane (BFP) image using the Bertrand lens for potential air bubbles in the immersion oil, which appear as dark occlusions on the round back focal plane image. TROUBLESHOOTING! In case of an air bubble, unmount the sample, clean the objective and coverslip underside and mount again.
528. Lock the focus (steps 446 - 449).
529. Continue to use 1% 640 nm laser while searching for a ROI. If bleaching is still too strong, set the laser triggering to pulse the laser by selecting the Rising trigger mode in the Trigger tab of the htSMLM UI. Select an appropriate pulse length (shorter than the camera exposure time) to generate sufficient signal for focusing, while minimizing bleaching. A good starting point is a 10,000 µs pulse length.
530. To achieve the best resolution, image NPCs located in the nuclear membrane closest to the coverslip. Ideally, select nuclear membrane areas that are flat, with most NPCs on the same focal plane. Start by slowly moving the focus up in z until most structures are in focus. TROUBLESHOOT: Should you struggle getting most NPCs in the ROI in focus, you might want to select a different nucleus where these structures are more homogeneously located. Some blurring at the edge of the nucleus is expected, as there the nuclear membrane bends upwards and out of focus.
531. STORM imaging: Decide between slow STORM (long acquisition times, high data quality) and regular STORM regime (short acquisition times, decreased data quality). a) *Slow STORM regime:* slowly turn off most fluorophores using approximately 0.2 kW/cm^2^ laser power density until the overall signal intensity in the ROI is not decreasing anymore. This usually takes between one to two minutes and can be determined by eye. Next, acquire images at 100 ms exposure time and approximately 6 kW/cm^2^ laser power. b) *Regular STORM regime:* Directly acquire images at 30 ms exposure time and approximately 20 kW/cm^2^ laser power density.
532. Image for approximately 80,000 frames (slow STORM regime) or 40,000 frames (regular STORM regime), until you reach the maximum 405 nm pulse length (10,000 µs), and the density of activated fluorophores drops. Adjust the 405 nm pulse length manually during the experiment to keep the density of localizations constant. Alternatively, you can use the automatic feedback of the 405 nm pulse length. Adjust the activation parameters to ensure signals are not overlapping. Start with the following parameters: Set activation density to ‘Get N:’ 6 blinks per frame, at a ‘sd coeff’ of 4.0, ‘Feedback’ of 0.05; ‘Average’ at 1.0. Automatically detect background using the ‘auto’ button. You can also set the background manually to a level so that the measured number of activations reflect the visual impression.
533. Use SMAP to fit the acquired data and reconstruct and render the localized image of 2D NPCs. See steps 460 - 465 for a description on how to set up SMAP, and steps 475 - 481 for a description of the localization procedure.
534. Follow steps 475 - 477 to set up image localization.
535. OPTIONAL. In order to perform the fitting in real time, parallel to the image acquisition, additionally tick ‘Online Analysis’ in the ‘Localize:Input Image’ tab. Leave other settings on default
536. Set up ‘Localize:Peak Finder’ according to step 474 - 478. TROUBLESHOOTING: Depending on the signal intensity of the detected peaks, the background cut off has to be adjusted. The ‘dynamic (factor)’ cut off ranges from 1 to 3. In the case of low intensity fluorophores, visual inspection of the preview image might show that too many emitters are undetected. In that case, it is advisable to lower the cut off until the desired emitters are included. As lowering this cut off will ultimately result in inclusion of background signal, an optimal balance between true detection and detection of background has to be established. In the case of bright fluorophores, the cut off can be increased. Double check new cut off settings in different frames using the slider next to the ‘Preview’ button.
537. The preview image also allows for restricting the ROI for which fitting is performed. To this end, draw the desired ROI in the preview image and press ‘ROI to include’ to include the selected ROI for fitting or ‘ROI to exclude’ to exclude the selected ROI from fitting. Double-click to confirm the selected region.
538. To perform 2D fitting, select ‘PSF free’ in the ‘Fitter’ tab as a PSF model instead of ‘Spline’. Using this setting, each detected signal is fitted with a 2D Gaussian, assuming a Gaussian-distributed PSF.
539. In the ‘Localize:Localization’ tab, the frequency of rendering new localization data can be adjusted. According to the default setting, during ‘online’ fitting, new data is rendered every 60 s.
540. Run the localization of the acquired data. The localization data will automatically be saved to the imaging folder after the acquisition is finished.
541. Load a previously saved .sml file into SMAP via the ‘File’ tab.
542. Even in good samples there will be localizations of poor quality that need to be removed from further analysis. We recommend initial filtering based on lateral localization precision and relative log likelihood as described in the SMAP Documentation. Common cut-off values are localization precision (locprec) < 20 nm and relative log likelihood (LLrel) > -1 and can be set in the ‘Render’ tab. For 2D data we also filter by the size of the PSF (PSFxnm < 175 nm) to remove out-of-focus fluorophores.
543. After this first filtering step, perform a drift correction in SMAP (step 493). Go to the ‘Process’/’Drift’ tab in SMAP and run the drift correction with the default settings. For 3D data additionally check ‘correct z drift’. TROUBLESHOOTING: Make sure that all of the curves scatter without too much error (tens of nm) around the black curve (see Supplementary Figure 59). Large scattering indicates that the redundant displacements are not compatible with each other and that the drift correction did not work correctly.
544. Next, analyze dye photophysics in the ‘Analyze’/’Measure’ tab and select ‘Statistics’. By default, the plugin evaluates only localizations currently depicted in the field of view in SMAP that meet the current filtering criteria. Press Run, and the resulting analysis will open in a separate window.
545. *Photons:* For AF647 you can expect approximately 6,000 to 8,000 photons per localization, for regular and for slow STORM respectively.
546. *Localization precision* can be expected to peak at 5 +/- 2 nm.
547. The average *on-time* (meanall) should be kept in a range of 2 – 2.5 frames.
548. To assess the *background*, make sure to perform the analysis on ungrouped data. Background is expected to peak at 100 photons/pixel.
549. Visual inspection of individual NPCs can aid in evaluating the data qualitatively. A circular shape and noticeable corners (preferably 8) are desired traits of each NPC. Note that an average labelling efficiency of 60% (desired target) leads to most NPCs displaying 6-8 corners, as each corner is occupied by 4 target proteins. For a quantitative approach to assess effective labelling efficiency see SMAP Documentation.
550. TROUBLESHOOTING
  - A low number of photons could be caused by 1) bad imaging buffer (this can be caused either by errors in buffer preparation, or due to the acidification of the imaging buffer over time. See section reagent preparation); 2) lower effective laser intensity than expected. In the latter case, check that the laser power is as expected using the laser power meter (COTS- PM100D/COTS-S121C) just before the illumination lasers pass into the microscope body. If the laser power is as expected, prepare a fresh aliquot of the imaging buffer and return to step 521 for repeating the biological sample imaging.
  - Poor localization precision could be caused by 1) low number of photons; 2) most structures being out of focus. 3) high background.
  - On-time out of the suggested range could be caused by 1) deterioration of the imaging buffer; 2) A discrepancy between the actual and expected effective laser intensity.
  - High background could be caused by 1) high intracellular and out of focus signal from unspecifically bound dye or dye in solution indicating errors in sample preparation; 2) deteriorated imaging buffer 3) other sources of light incident on the camera. For example, refer to point 458 regarding the NIR blocking filter. Isolate the camera better from sources of light.
  - Low number of localizations could be caused by 1) decreased data quality according to the points above or 2) low dye concentrations or poor cell permeabilization during sample preparation.
  - Low labelling efficiency could be caused by 1) low number of localizations; 2) cell culture related issues (e.g., prolonged culturing of cells past three months); 3) problems with the reagents or labelling protocol.

### 3D single-color imaging of nuclear pore complexes (Timing: 5 days)

3D imaging is performed analogously to 2D imaging of nuclear pore complexes, but with the astigmatic lens in the beam path. Additionally, a PSF calibration is required to allow 3D fitting, as described in steps 466 - 473. Localization of the acquired frames in SMAP is set up as described in steps 474 -479, using ‘Spline’ in the ‘Fitter’ tab as a PSF model. Filtering of the 3D data as described in step 542 can be further extended by filtering in z. It is recommended to choose a range well within the linear region of the bead calibration, as shown in the ‘validate’ tab of the calibration results window (see step 473). We recommend that not too stringent filtering is applied and to include a range of ± 400 nm. By accessing the filtering window (see step 492), the distribution of z positions can be assessed. Here, ensure that your z-filtering range includes the majority of localizations, otherwise adjust accordingly. Drift correction is performed as described in step 543, additionally ticking ‘correct z drift’.

To gain a more intuitive understanding of the z information in the super-resolved image, the localizations can be color coded depending on their z-position. To this end, go to the ‘Render’ tab and change ‘Colormode’ from ‘normal’ to ‘z’. Change the ‘LUT’ to ‘jet’ to recreate images shown in Figure 8.

**Figure 8.**
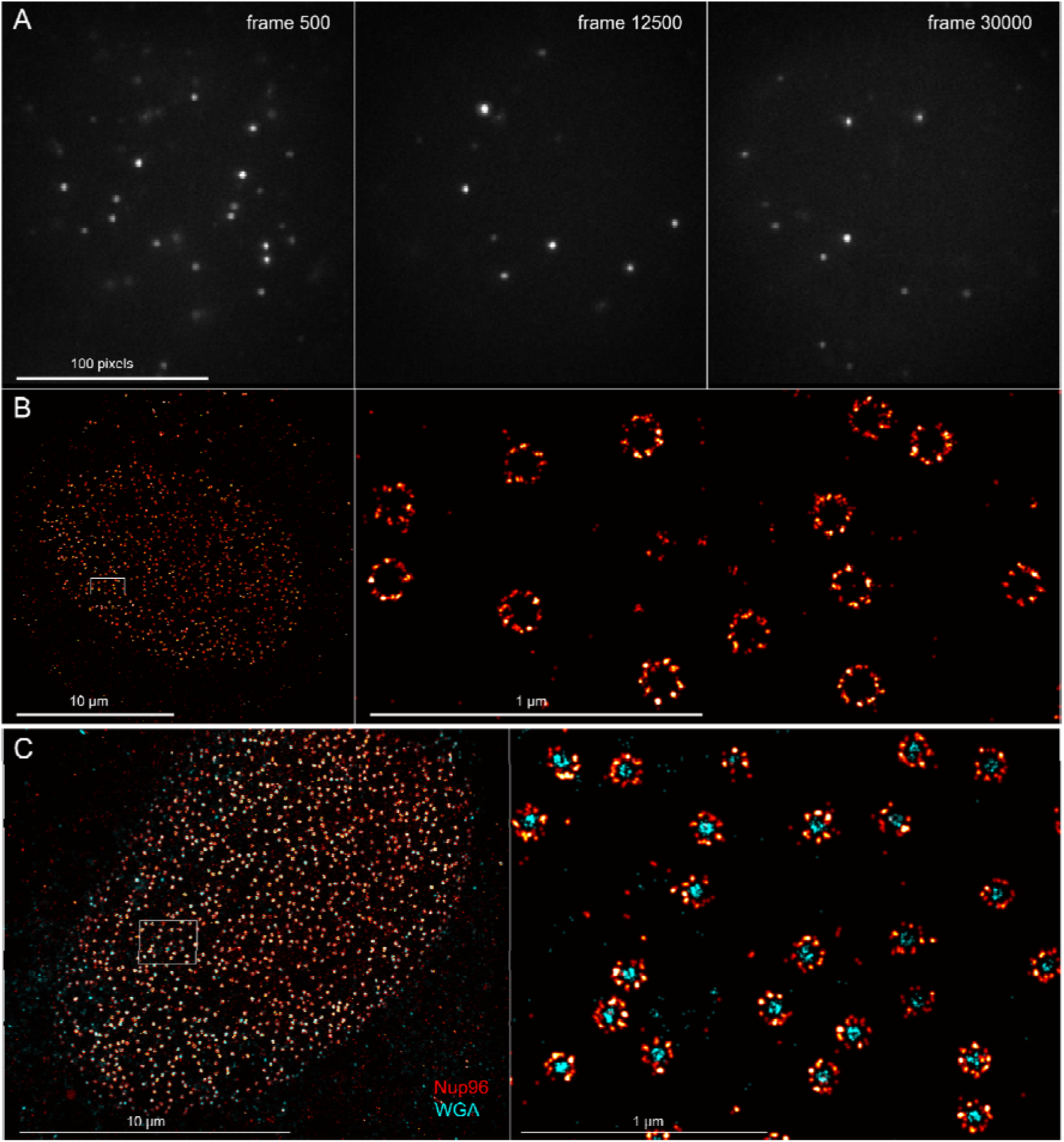
2D imaging of the NPC. Here Nup96-SNAP is labelled with Alexa Fluor 647 and imaged with slow STORM. A: Raw camera frames show bright single molecules with a sufficiently low density. B: Single-color overview and zoom into individual NPCs. C: Dual-color overview and zoom, here additionally the central channel is labelled with WGA-CF680.

### 3D, multi-color imaging of nuclear pore complexes (Timing: 5 days)

The aim of the following procedure is to generate dual-color images of the NPC protein Nup96- SNAP-AF647 together with the NPC channel which can be targeted using WGA-CF680. The two far red dyes used in this example result in highly overlapping emission spectra. To distinguish both fluorescent species, a ratiometric imaging approach is applied, where signal from both fluorophores is split into short- and longwave components by a dichroic and imaged simultaneously on two separate regions of the camera chip. Assignment of a given localized emitter to a specific fluorophore is then based on the intensity ratio between both camera channels. To fit this multi-channel data, a global PSF calibration is required (see GlobLoc [16]). This calibration calculates the transformation between both channels and generates a dual-channel PSF model (step 481 - 485). While imaging is performed analogous to 3D imaging in single color, specific adjustments have to be made which are detailed in the following steps.

551. Mount and image a fluorescent bead sample. Refer to: Fluorescent bead samples in the reagent setup section for a detailed description of the sample preparation and: Focusing the microscope for a description of setting up the microscope to image fluorescent beads.
552. Set up the microscope according to step 523 - 525.
553. Before imaging, ensure that the microscope is set up for 3D dual-color imaging. This includes the installation of the imaging splitter dichroic (COTS-SB1-M, COTS-25.5×36x3MM- DICHROIC-SP, COTS-KM200S, FAB-EMBL-000068/69) on the lower half of its kinematic magnetic mount (COTS-SB1-M).
554. Select the 676/37 and 685/70 bandpass filters for the reflected and transmitted paths respectively.
555. Activate the 3D property to position the astigmatic lens in the emission path for 3D imaging.
556. The beads should now be visible in two spatially separated camera regions. In the reflected channel (shortwave, nominally left on the camera image), the beads will appear dim but visible. In the transmitted channel (longwave, nominally right on the camera image), the beads will appear brighter. The transmitted channel being brighter will provide the basis of the precise localization in later steps while the reflected and transmitted channels taken together provide the basis of the fluorophore’s ratiometric ‘fingerprint’. TROUBLESHOOTING! If the relative brightness of the beads in the two channels is reversed, it is possible that the camera has been installed in the wrong orientation. Refer to step 213 to rectify this and return to step 466 to repeat the 3D calibration. If the beads cannot be seen in one channel or the relative brightness between the channels is similar, check the orientation and part number of all dichroics and other spectrally discriminatory elements (notch/emission filters etc.) in the emission path.
557. Perform steps 466 - 469 to acquire several bead stacks for a more precise 3D PSF model.
558. Generate a bead calibration according to step 481.
559. After performing a satisfactory 3D bead calibration and transformation, next mount the biological sample labelled for Nup96-SNAP-AF647 and WGA-CF680 (see section: Biological sample preparation). TROUBLESHOOTING: WGA-CF680 staining can require additional titration. The initial dilution of 1:100 might have to be adjusted with every aliquot. Here it should be optimized between the majority of NPC being labelled (high concentration of WGA- CF680), while minimizing background (low concentration of WGA-CF680). Background is caused by WGA binding to the plasma membrane, which in turn generates out of focus signal. This can be reduced by further diluting the WGA-CF680 aliquot. Low WGA-CF680 concentration can in turn decrease the number of NPCs labelled, thus the right concentration should be carefully titrated and empirically determined.
560. Imaging of 3D dual-color biological samples is performed as described in steps 526 - 532 for 2D single-color imaging.
561. Perform 3D dual-color fitting according to step 481 - 484. Here, compared to 2D and 3D fitting, the workflow has to be changed to ‘fit_global_dualchannel’ and the previously generated PSF calibration file for multi-channel imaging has to be loaded.
562. After color assignment, post-processing can be performed as described in section Post-processing and rendering with SMAP.

## Anticipated Results

A correct implementation of the protocols as described, is expected to yield a fully functional 3D multi-color SMLM system with potential for automated high-throughput applications. Diffraction-limited resolution and 3D localization precision approaching theoretical limits will have been demonstrated across a wide field of view and used to provide exemplary 2D and 3D images of nuclear pore complexes via single and multi-color dSTORM workflows (Figure 8 and Figure **9**). In this regard, the nuclear pore sample provides an ideal standard with which to benchmark performance and a strong basis to apply the microscope confidently to a range of biological studies.

**Figure 9.**
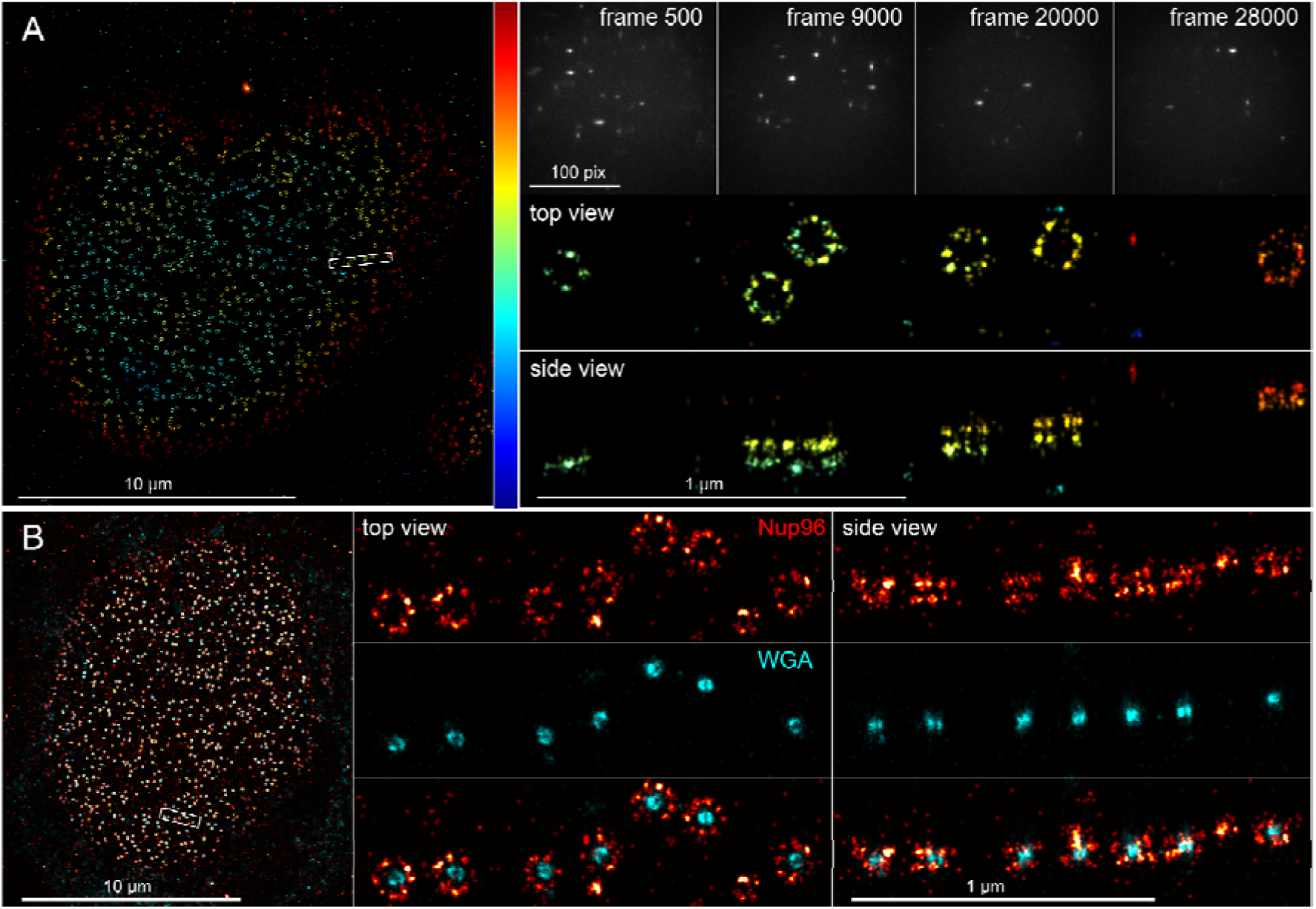
3D imaging of the NPC. Here Nup96-SNAP is labelled with Alexa Fluor 647 and imaged with slowSTORM. A: Overview, raw camera frames and single-color zoom into individual NPCs. The localizations are color-coded by their z-position with blue corresponding to z = -300 nm and red corresponding to z = +300 nm. B: Dual-color 3D imaging.

NPC data can further allow for the correction of z-dependent image distortions. The expected ring separation for NPCs is 49.3 nm. By determining the ring separation in the data set using advance geometric model fitting, the deviance from this expected value can be calculated and the z-scaling of the data corrected accordingly [49,62].

## Acknowledgements

We thank Christian Kieser (EMBL Electronic Workshop) for help with construction of and documentation for the microFPGA. We thank Arthur Milberger (EMBL Mechanical Workshop) for providing all mechanical drawings. We thank Joran Deschamps (Human Technopole, Milan, Italy) for providing the EMU htSMLM user interface and continued support in various aspects of microscope control. This work was supported by the European Research Council (CoG-724489) and the European Molecular Biology Laboratory.

## Footnotes

1 Note intensity is used here to refer to power per unit area, more rigorously termed irradiance in radiometry. We use the term intensity throughout as this is a more common convention in microscopy

## Notes

https://github.com/ries-lab/3DSMLM

### Competing Interest Statement

The authors have declared no competing interest.

