## Supplementary Information for "Automated 3D multi-color single-molecule localization microscopy"

### Supplementary Note 1: Using the parts list and understanding part names

The parts list (Supplementary Table 2) is broken down into three main sections. The first, Assemblies, describes the content of each CAD assembly for ease of navigation. The Assemblies are named with the format ProjectID-Institution-XXX. For example, 3D-SMLM-EMBL-001 is the master assembly for the microscope. Sub-assemblies reference the parent assembly, for example, 3D-SMLM-EMBL-001-001 is the microscope body, which is one of the major sub-assemblies of the microscope. This continues down the hierarchy. For example, 3D-SMLM-EMBL-001-001-001 is the emission path tube lens, which itself is a sub-assembly of the body 3D-SMLM-EMBL-001-001, which is a sub-assembly of the master 3D-SMLM-EMBL-001. The second section, Custom, includes all custom parts of the microscope with the nomenclature ProjectID-Institution-FAB-XXXXXX (where FAB denotes a fabricated component). For example, the bottom plate for the microscope body is 3D-SMLM-EMBL-000001. The material to use for the manufacture of each part is listed in the table. Note, that this information is not necessarily included in the associated CAD file. The parts list also includes a more detailed description of the material that has been used in the EMBL workshop to fabricate the components; other specifications with similar performance can be used.

The third section, COTS (commercial off-the-shelf), includes all commercially available parts with the nomenclature ProjectID-COTS-XXX. For example, the part 3D-SMLM-COTS-AC254-060-A is an achromatic doublet lens. The parts list includes the supplier in the table. For example, for 3D-SMLM-COTS-AC254-060-A, both the supplier and manufacturer are Thorlabs. Where parts are not directly sourced from their manufacturer, the manufacturer of the component is included as additional information. In some cases, the part number may not match exactly to the manufacturer designation. For example, many Thorlabs parts use the ‘/M’ suffix to denote a metric threaded part. Since ‘/’ is reserved for file path designations, the part number suffix is amended to ‘-M’ e.g., TRA20/M is listed as 3D-SMLM-COTS-TRA20-M. The COTS parts are further subdivided into groups describing their general function, for example, lenses and mirrors are ‘Optics’ while a mounting post for these components would be ‘Optomechanics’. In some cases, it may be hard to source a given component from the listed vendor. For generic items such as basic electronic components, screws and fixings, it is acceptable to source these from existing stocks or other manufacturers, paying attention to associated tolerance implications (e.g., in the case of dowel pins).

The parts list for the FAB and COTS parts provides numbers of each part required and also whether they belong to the base configuration of the microscope or one of the illumination configurations, which are interchangeable, thus aiding the purchase of the correct components for the desired configuration. In addition, certain parts are required as assembled alignment tools. In this case, the same component may be shared between tools (e.g., a ring-adjustable iris 3D-SMLM-COTS-SM1D12D) providing that all tools that are needed concurrently have access to the required part.

In the protocol rather than referring to the full part name, for FAB and COTS parts we omit the ProjectID (3D-SMLM-) and refer to the parts in shorthand as e.g., FAB-EMBL-000001 and COTS-TRA20-M. Assemblies, including the various assembled tools are referred to in long form as e.g., 3D-SMLM-001.

### Supplementary Note 2: Selecting spectrally discriminative optics

The microscope detailed herein, requires the use of several dichroic mirrors for the functions of coupling various light paths into the microscope body, selecting spectral windows for imaging on the camera and cleaning up and blocking laser sources, which unchecked, will corrupt images. We include in this note a guide to selecting these optics for the microscope and include filters, which are suitable to use in most cases. As a general note, the use of the optics described is assumed in the protocol section for 2D, 3D and multi-color imaging of the biological samples. For other tasks such as viewing fluorescent bead samples, other options may also suffice. We include the OEM name but also note that the items are more readily sourced in some geographical regions via distributors. Note that the multi-mode laser engine (if present) requires additional internal spectrally discriminative optics to combine the various laser lines. However, we consider only the output of the laser engine from the multi-mode fiber.

First consider the dichroic mirrors: DM1/2/3. The thickness considerations for dichroic mirrors are discussed in: Affixing dichroic beamsplitters in glue-in mounts. For this discussion we assume the use of 3 mm thick substrates only. DM1 should reflect the illumination source(s) and efficiently pass the emitted fluorescence and focus lock laser (808 nm). In the base configuration including laser lines at 405, 488, 561, and 640 nm, a quadband excitation dichroic (TIRF or imaging grade) is an optimal choice (Semrock: Di03-R405/488/561/635-t1-25x36/Di03-R405/488/561/635-t3-25x36). This optic reflects the four laser lines and a narrow surrounding band to minimally attenuate the emitted fluorescence close to the laser line while transmitting the focus lock laser. DM2 should reflect the emitted fluorescence out of the body while also passing the focus lock laser. In this case, a long pass dichroic is recommended (Semrock: Di03-R785-t3-25x36). DM3 is the image splitter dichroic, which contributes to the selection of spectral bands for the two camera channels. In this case, one typically uses a long pass dichroic. In the case of ratiometric multi-color dSTORM imaging, which is commonly performed to discriminate up to 4 far-red dyes (emission maxima 650 - 700 nm) a long pass dichroic with an edge at 665 nm is preferred (Chroma, T665LPXR, custom: 3 mm thick). For splitting of typical green/red fluorescent protein (GFP/RFP) channels an edge at 560 nm (Chroma, T560LPXR, custom: 3 mm thick) is suitable and for splitting GFP or RFP from a far-red channel an edge at 635 nm is suitable (Chroma, T635LPXR, custom: 3 mm thick). Many other options are suited and should be selected on the basis of fluorophores to be separated.

Next, consider emission filters. For the standard setup with the 665 nm long pass dichroic installed in the image splitter. The reflected path serves typical GFP/RFP bands as well as the shortwave portion of the far-red fluorescence. For GFP and RFP respectively, one should opt for broad bandpasses of the type 525/50 (center/width) (Semrock: FF01-525/50-25) and 600/50 (Semrock: FF01-600/52-25). For the shortwave portion of the far-red, a narrower 676/37 bandpass has been proven to work well (Semrock: FF01-676/37-25). This filter is also useful for single-color imaging, which is performed without the image splitter dichroic and so should ideally be available in both the reflected and transmitted channels. The longwave far-red channel is best served by a very broad 685/70 bandpass (Chroma, ET685/70m). For far-red ratiometric imaging, deviating from the choice of filters listed should be avoided to attain optimal results.

In addition to the emission filters, it is necessary to provide cleanup of the laser sources and blocking of the photodetectors (sCMOS camera/laser power monitoring photodiode) to avoid off band emission (from lasers) and more generally minor reflections (which can still be brighter than the fluorescent signal) from impinging on the detector. In the case of the single- and multi-mode laser lines at 405, 488, 561 and 640 nm, a laser cleanup filter is used (Semrock, FF01-390/482/563/640-25). The emission path should include a notch filter at all excitation wavelengths (Semrock, NF03-405/488/561/635E-25). Typically, a short pass filter is sufficient to block the focus lock laser Semrock, FF01-758/SP-25) in the emission path but a more costly notch filter can also be used to ensure that the focus lock laser is not visible on the camera for long exposures or at high power (Semrock, NF03-808E-25). Finally, the laser monitoring photodiode requires a filter to block the focus lock laser, in this case a notch filter is recommended (Thorlabs, NF808-34) as described in the protocol.

### Supplementary Note 3: Using non-specified objective lenses

For various reasons, one may wish to use an objective lens that is not specified for use in the system. The effective pixel size magnified into the object space is ca. 106 nm (theoretical), using a f = 1.8 mm (100×) Olympus objective lens. Deviating strongly from the stated focal length is not recommended as the effective pixel size will scale accordingly. Therefore, the use of 60× lenses is precluded or requires the redesign of the emission path. In any case, the available laser power and emission path optics have been specified to provide appropriate intensity and minimal field aberrations over the maximum field of view of the microscope (⌀77 μm). In the case of alternative 100× lenses, the 60 mm parfocal length favored by Nikon can be accommodated by raising the sample-positioning XY stage by 15 mm and adjusting the bore diameter thread specifications of part FAB-EMBL-000043 (Nikon options are typically M25x0.75). Since the stages will in this case be raised up, the side plates for the microscope cover (parts: FAB-EMBL-000019/20) should be extended to accommodate the increased height and position the ring LED source appropriately for transmission imaging. Note, that using Leica/Zeiss lenses would require a more fundamental overhaul of the architecture to incorporate the manufacturer specified tube lens, which corrects for fixed chromatic aberrations in the objective lenses and for this reason is not recommended.

### Supplementary Note 4: Using non-specified single-mode lasers and laser-engines

For builders wishing to use pre-existing or otherwise non-specified laser systems (for example to achieve higher intensity at the sample) as excitation/activation sources, there are additional concerns to consider beyond basic expectations of minimal deviation from the stated wavelength, power stability and temperature control. Firstly, one must ensure that µManager device adapters providing the required functionality are available. In addition, for the laser exposure to follow the camera exposure signal, the laser (or associated acousto-optic modulators) must be digitally modulatable at > 1 kHz. The illumination scheme utilizing a refractive beam shaper also presents additional requirements in that the beam quality, diameter and divergence are tightly specified for correct functioning. For free-space lasers, it is advisable to construct a spatial filter to clean the beam-profile. For reference the optional booster laser uses such a configuration to achieve a clean profile and a subsequent variable focal length compound collimator to achieve the required beam diameter. Fiber-coupled lasers can be expanded to the required diameter following a similar scheme to variably collimate the diverging fiber output. In either case, the builder will need to re-design the sections of the illumination path prior to the beam-shaper, ensuring that the associated device footprint is still small enough to be accommodated on the optical table.

### Supplementary Note 5: Using non-specified XY stages

For builders wishing to use pre-existing or otherwise non-specified XY stages, first one should consider whether the preferred option is sufficient for the task at hand. In this regard, the stage should be capable of measuring and holding a stable position at rest (at a scale < 10 nm) and for automated multi-position acquisition should have a bi-directional repeatability < 5 µm. It is recommended to first ensure that the stages have pre-existing µManager device adapters. One must also consider what additional modifications are required to the body to accommodate the stages. For example, taller stages can be accommodated up to a point by reducing the size of (or eliminating) the spacer (for the COTS-SOM-12090, FAB-EMBL-000047). The stage should be supported on the top plate, without encroaching on other hardware, which limits the stage footprint. Additionally, the cover for the microscope, which greatly aids in isolating the microscope from cross currents, is sized for the maximum travel range of the specified XY stage and may require modification. Moreover, the specified XY stage shares a controller with the 1D stage used for positioning the quadrant photodiode of the focus lock system, which needs to be accounted for in any decision to omit the specified option.

### Supplementary Note 6: Using non-specified cameras

For builders wishing to use pre-existing or otherwise non-specified cameras, we refer to the section: Imaging camera, which lists many cameras that are and are not well-suited to use. Here we note explicitly the basic requirements of the camera. First, the camera should be supported by a µManager device adapter. The camera should ideally have high quantum efficiency, low noise, and a well-linearized response across all pixels. This largely excludes inexpensive industrial/machine vision cameras, limiting one to scientific options. The camera should be capable of being triggered (passive camera mode) and/or should provide a TTL signal indicating when the camera is exposing (active camera mode). The camera pixel size should be in the range 5.5 - 7.5 µm at the design magnification to achieve an object space pixel size of 90 - 122 nm. If either the pixel size is not 6.5 µm or the number of pixels is not 2,048 or 2,304 then the X,Y pixel positions provided in Supplementary Table 11. Field of View Centre Positions on Camera are invalid and should be determined from first principles. Likewise, one must consider whether the camera chip is large enough to accommodate the two image channels. Furthermore, if the camera to be used operates in a rolling shutter mode, it must be oriented in such a way that the same fluorophore is imaged synchronously in the two camera channels.

### Supplementary Note 7: Camera data rate

The Hamamatsu Fusion BT camera included in the reference configuration is capable of acquiring images at an equivalent of ca. 1 GB/s. Transferring the data at this rate requires a PCI Express CoaXPress frame-grabber card at substantial additional expense. However, this is not typically necessary for SMLM imaging. Under the design conditions, the maximum size of the field of view on the camera assuming a rectangular region of interest is given by the largest illumination field size (⌀77 μm) multiplied by the emission path magnification (61.1×) giving 4.7 mm or 723 pixels (6.5 μm pixel size). Given that the splitter provides two such images on the camera, which are placed side by side at the center of the camera chip, the effective maximum field of view is 723 × 1,446 pixels. With 16-bit encoding and assuming that the frame rate is limited only by the exposure time (not always the case, but for sake of argument this limitation provides a worst possible case), for a very short exposure time of 10 ms, the associated data rate is ca. 200 MB/s, which is well within the bandwidth of a USB 3.0 connection. For builders seeking to extend the reported system to larger field of views or to operate the camera at higher frame rates, the CoaXPress frame grabber may be required and the IT infrastructure described under the section: Computational infrastructure, may require expansion commensurate with the increased data rate.

### Supplementary Note 8: Configuring the microscope body and optical path handedness

If the builder wishes to achieve a different handedness from the base configuration with illumination from the back side, emission to the right and focus lock to the left, this can be achieved by replacing the central optical pillar mounting the various mirrors/dichroic beamsplitters with mirrored versions of the parts (provided as additional part files). In this manner, the emission and focus lock path orientations will stay the same, while the illumination will flip from the back to the front side. The base plate of the microscope has the required dowel pin and threaded hole features to accommodate the mirrored parts. It is also possible to achieve a different handedness by routing the emission path to the left side and focus lock correspondingly to the right. The builder will need to specify the orientation before having parts fabricated. Depending on the ultimate desired layout on the optical table, it may be necessary for the builder to adapt other handed parts accordingly. For example, flipping the illumination orientation will require mirroring the emission and focus lock paths to stay within the specified table footprint, each of which has handed parts associated with them. Given the technical challenge associated with making the required changes to the part files and assemblies, we strongly recommend following the base configuration specified. If the builder wishes to build the system in a mirrored configuration, it is necessary to make the required changes to the CAD files and assemblies before having parts made.

Note that this protocol has been developed over several generations of the microscope using slightly different versions of the XY stage. To maintain back compatibility, we have left mounting points available for all options, but specify that the newer stage included in the core design is used. The body features additional configurability in the form of a recess for a heating foil and temperature sensor, which, provided proper control, allows the top half of the body to be heated either to maintain an elevated temperature for imaging or to ensure that the body does not provide a heatsink for stage top environmental control chambers operating at elevated temperatures. If the heating foil is to be used, we recommend that part FAB-EMBL-000006 is made of plastic to act as an insulator between the upper and lower sections of the microscope body. If the heating foil is omitted, this part should be made of aluminum to maintain the correct spacings between various components in the body. Since its implementation requires only a few parts, we recommend that builders who are unsure whether heating functionality is desired follow the base configuration. Otherwise, a configuration with the associated recesses and channels absent is available in the CAD assembly. Finally, body dichroic mirror mounts are provided for 1 (illumination only), 3- and 5-mm thick substrates. Both 3 and 5 mm thick dichroics have been tested to provide excellent performance for all dichroics, while TIRF-grade 1 mm thick dichroics are suitable for coupling in illumination only. It is critical that the correct mounting part is selected in either case to ensure that the reflected optical axes are positioned correctly.

### Supplementary Note 9: Specifying an infinite conjugate source

A rearrangement of the thin lens equation provides an estimate of the discrepancy between the lens-camera spacing found from the finite object distance and the true lens-camera spacing for a truly infinite conjugate, Δz:

$\Delta z = \left[ \frac{1}{\left( 1/f \right)-(1/S1)} \right]$ -f	(1)

Where f is the focal length of the lens used, S1 is the distance of the object from the lens. Focusing in this manner provides an adequate model of infinity when Δz is much smaller than the depth of focus at the camera. The depth of focus is dependent on the effective numerical aperture of the imaging lens under operating conditions. In the protocol, the infinite conjugate viewing camera will be used up to its full NA defined by the mechanical size of the lens. The effective focusing NA can thus be estimated as:

$NA=\frac{d_{clear,lens}}{2f}$	(2)

For the 1” diameter lens (d_clear,lens_ = 23 mm when mounted), f = 200 mm lens used we arrive at an NA of 0.0575 Estimating the depth of focus, Δf as:

$\Delta f=\frac{\lambda}{{NA}^{2}}$	(3)

Where 𝝀 is the wavelength of light. For 510 nm light, the depth of focus is ca. 0.309 mm. Assuming that we want to be accurate to within ± 20% of the depth of focus, we find that the allowable shift is ca. 0.12 mm (equivalent to ca. 20% of the thread pitch used in the lens tube coupler that focuses the infinite conjugate camera and transmission target). From equation 1 we find that the object needs to be at least ca. 335 m away from the lens. Such distances are not readily encountered indoors so it is suggested that the camera is taken to a window of a building and used to image a suitably distant object (clouds/trees/buildings all work well).

### Supplementary Note 10: Correct orientation of the sCMOS camera

The sCMOS camera acquires images by a rolling shutter mechanism with sequential exposure and readout of pixel rows from the vertical center of the chip upward and downward. The achievable frame rate is thus related to the centration of the exposure region with respect to the vertical center and the height in rows of the exposed region. The camera chip is much larger than required given sampling and field of view limitations (see Overview of the automated 3D multi-color single molecule localization microscope), allowing two images to be placed side-by-side on the camera chip and resulting in an effective field of view on the camera that is extended in one direction. To achieve the highest frame rates, the shorter dimension should ideally be oriented vertically on the camera chip. Furthermore, there are notable cases where emission from a single-molecule needs to be captured in the two image-splitter channels synchronously. For example, in ratiometric dSTORM multiple spectrally overlapping dyes can be spectrally separated on the basis of ratiometry. The rolling shutter architecture thus places constraints on how the two channels are oriented on the camera, requiring that they are placed side to side (since a given pixel row will expose across its length synchronously whereas a pixel column will not). As such, it is clear that both frame rate and exposure constraints suggest that the two channels should be oriented horizontally and so the camera should be oriented with its upper side facing up. CAUTION: Although the camera could be inverted such that the upper side is facing down, the position of images on the chip will not be as expected in the protocol and so this arrangement should be avoided.

### Supplementary Figures

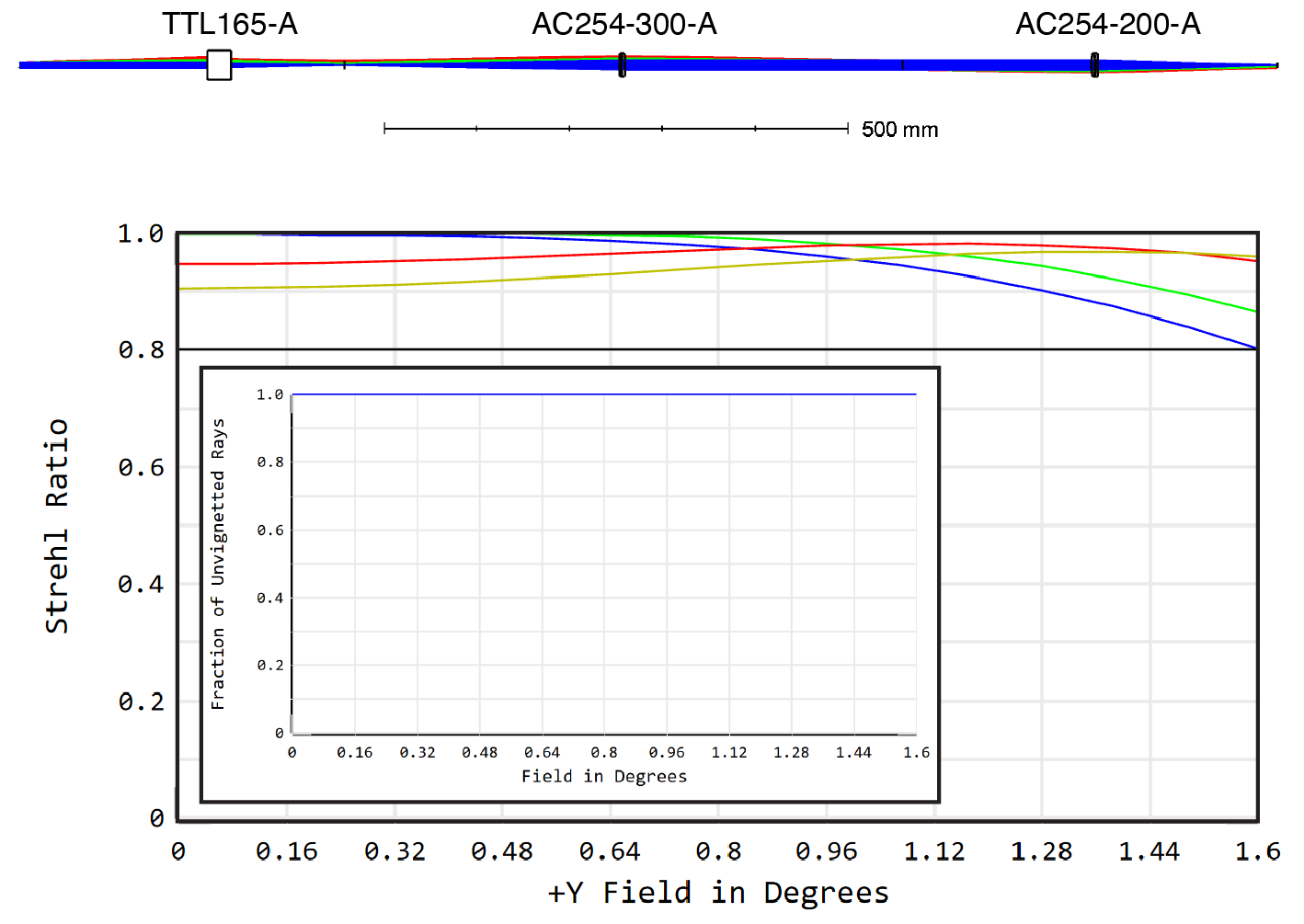

Supplementary Figure 1: Ray tracing optical simulation of the emission path imaging forming optics (Zemax, OpticStudio). Top: optical pathway comprising a widefield tube lens, and two achromatic doublets (all Thorlabs). The optical simulations were performed for a 5.4 mm entrance pupil (the largest for all objective lenses considered in the protocol) and a maximum field angle of 1.6 degrees, which is equivalent to an object height of 50 µm under paraxial assumptions (field of view of ⌀100 µm). Rays are traced from the entrance pupil (left, object at infinity) to the secondary image (right, camera chip) Bottom: the optical performance of the system as defined by the Strehl ratio at 515 (blue), 580 (green), 660 (red), 700 (gold) nm. A Strehl ratio of above 0.8 is defined as diffraction limited performance, illustrating that the optical pathway is diffraction limited over the field of view and wavelength range required. Inset: The vignetting plot shows that no rays are vignetted by the optical system.

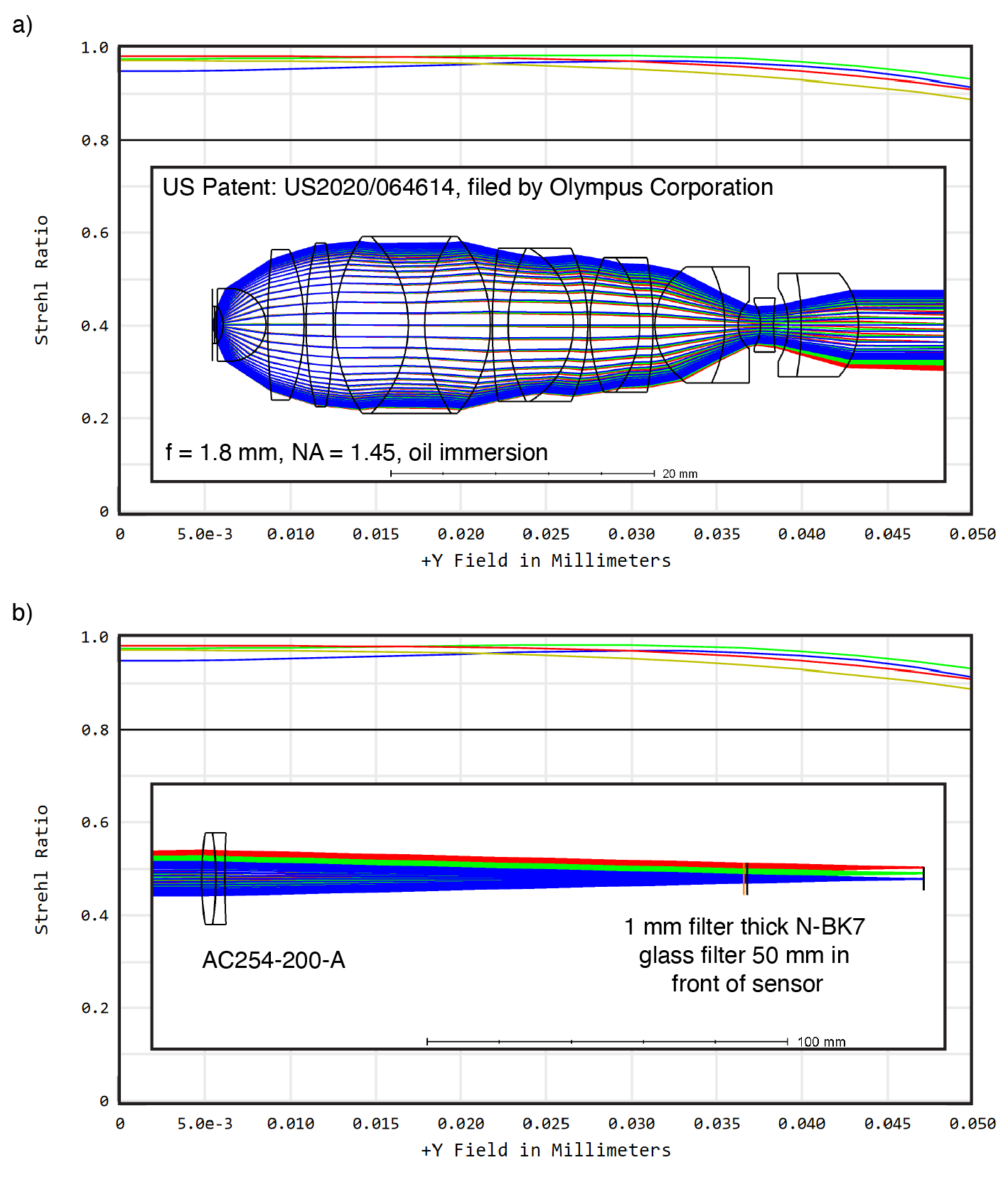

Supplementary Figure 2: a) Ray tracing optical simulation of the emission path imaging forming optics of Supplementary Figure 1 (Zemax, OpticStudio) with the addition of a high NA oil objective lens with similar specifications to the objectives recommended in the protocol. The optical simulations were performed for a NA of 1.45 (defining an entrance pupil of 5.22 mm under paraxial assumptions) and an object height of 50 µm (field of view of ⌀100 µm). The optical performance of the system is shown by the Strehl ratio at 515 (blue), 580 (green), 660 (red), 700 (gold) nm. A Strehl ratio of above 0.8 is defined as diffraction limited performance, illustrating that the optical pathway is diffraction limited over the field of view and wavelength range required for a typical high NA objective lens. The slightly improved performance relative to the results presented Supplementary Figure 1 is a result of smaller effective entrance pupil, deviations from paraxial assumptions and potentially a counterbalancing of aberrations between the objective lens and other elements. b) The addition of a standard optical filter placed in the secondary image space in front of the camera sensor. The change in the Strehl ratio is negligible, highlighting that the infrared blocking filter placed in front of the camera has no detrimental impact to the image formation.

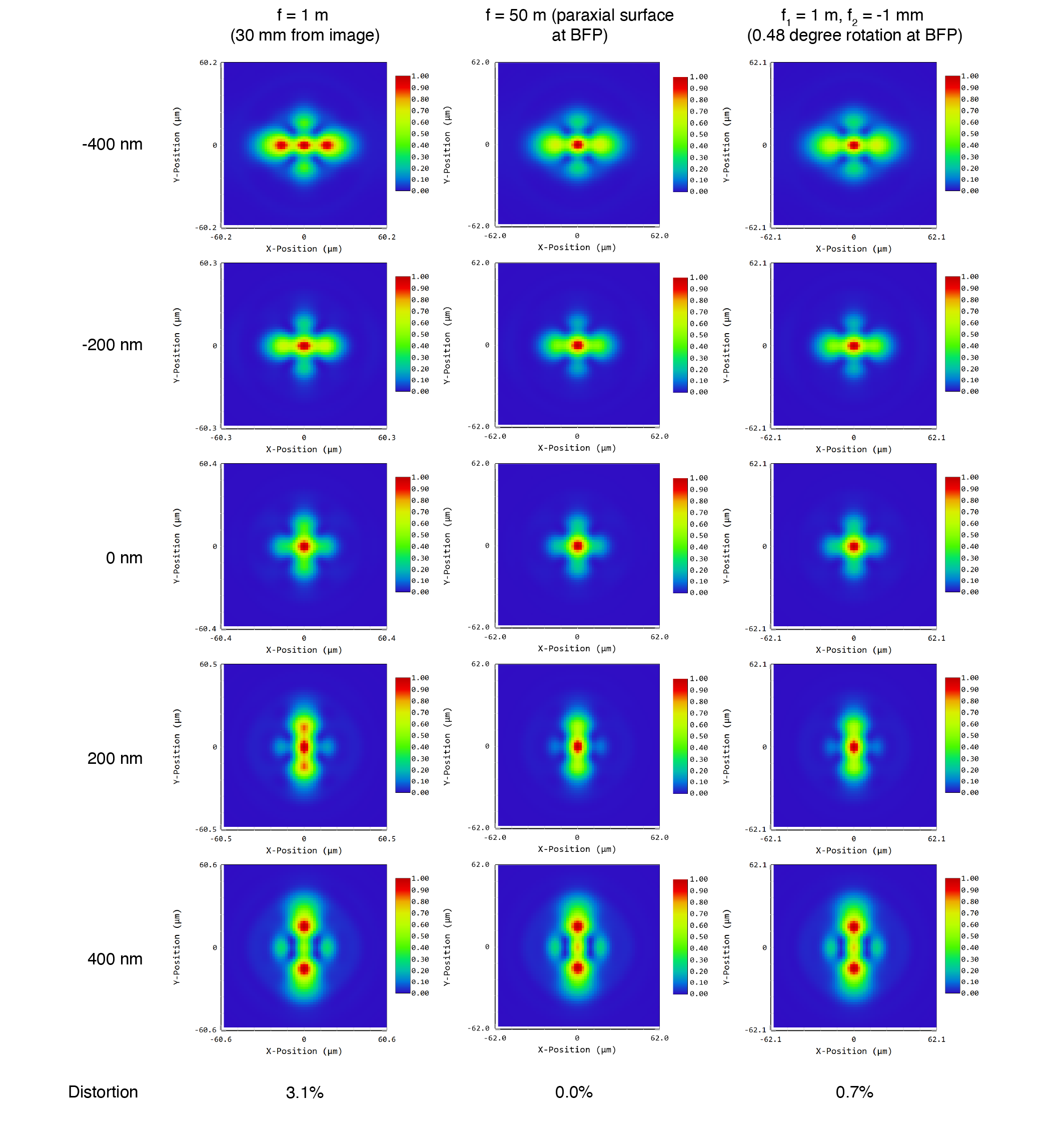

Supplementary Figure 3: 3D point spread function simulations for the emission path modelled in Supplementary Figure 2 with the addition of various elements to introduce astigmatism. Left: a paraxial surface with optical power equivalent to a f = 1 m cylindrical lens, placed 30 mm from the image plane/camera chip. Centre: a paraxial surface with optical power equivalent to a f = 50 m cylindrical lens, placed conjugate to the back focal plane of the objective lens (at the intermediate focus shared by the lenses AC254-300-A, AC254-200-A). Right: a pair of cylindrical lenses placed either side of the back focal plane (f1 = 1 m, Thorlabs, LJ1516RM, f2 = -1 m, Thorlabs, LK1002RM) and rotationally offset by 0.48 degrees relative to each other. The microscope is focused 400 nm into a sample with identical refractive index and dispersion as the immersion oil (for real samples, there will be an increased contribution from spherical aberrations). The PSF is shown at different depths (from -400 nm at the coverslip boundary to +400 nm). The elongation of the PSF is clearly seen above and below the focus. The three simulations give similar results as viewed from the shape of the PSF. However, the distortion is greatly reduced when placing the astigmatic element at the back focal plane. A single infinitely thin surface at the back focal plane (center) is not a realizable configuration and the rotationally offset cylindrical lenses (right) provide an ideal solution allowing adjustment of the precise amount of astigmatism while reducing distortion to negligible levels.

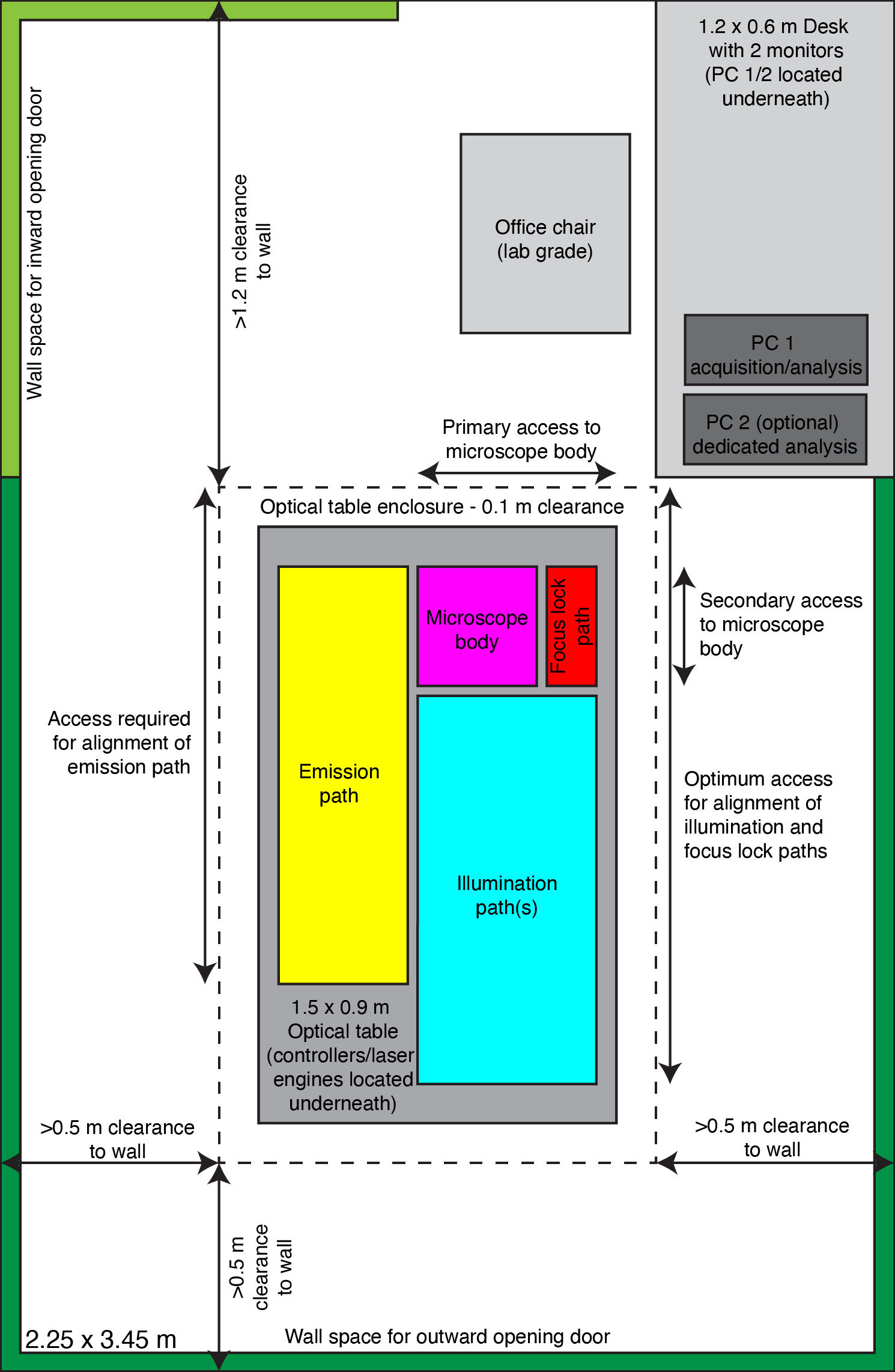

Supplementary Figure 4: An optimized layout of the microscope in a typical room (2.25 × 3.45 m). The user can easily switch between operation of the microscope and PC by pivoting on a lab grade office chair. The optical table can be accessed from all four sides with a minimum clearance to the wall of 0.5 m and a clearance around the optical table of 0.1 m, which is sufficient for an optical table enclosure. Access is only required from the two long edges and the short edge where the body is situated. Outward opening doors can be accommodated at most positions along the walls (dark green), while an inward opening door could be located in the corner shown (light green). All controllers and other large electronics can be accommodated beneath the optical table and on top of a table enclosure as appropriate. Power and compressed air can be provided either from the walls (using suitable floor channeling) or from the floor/ceiling as the room allows.

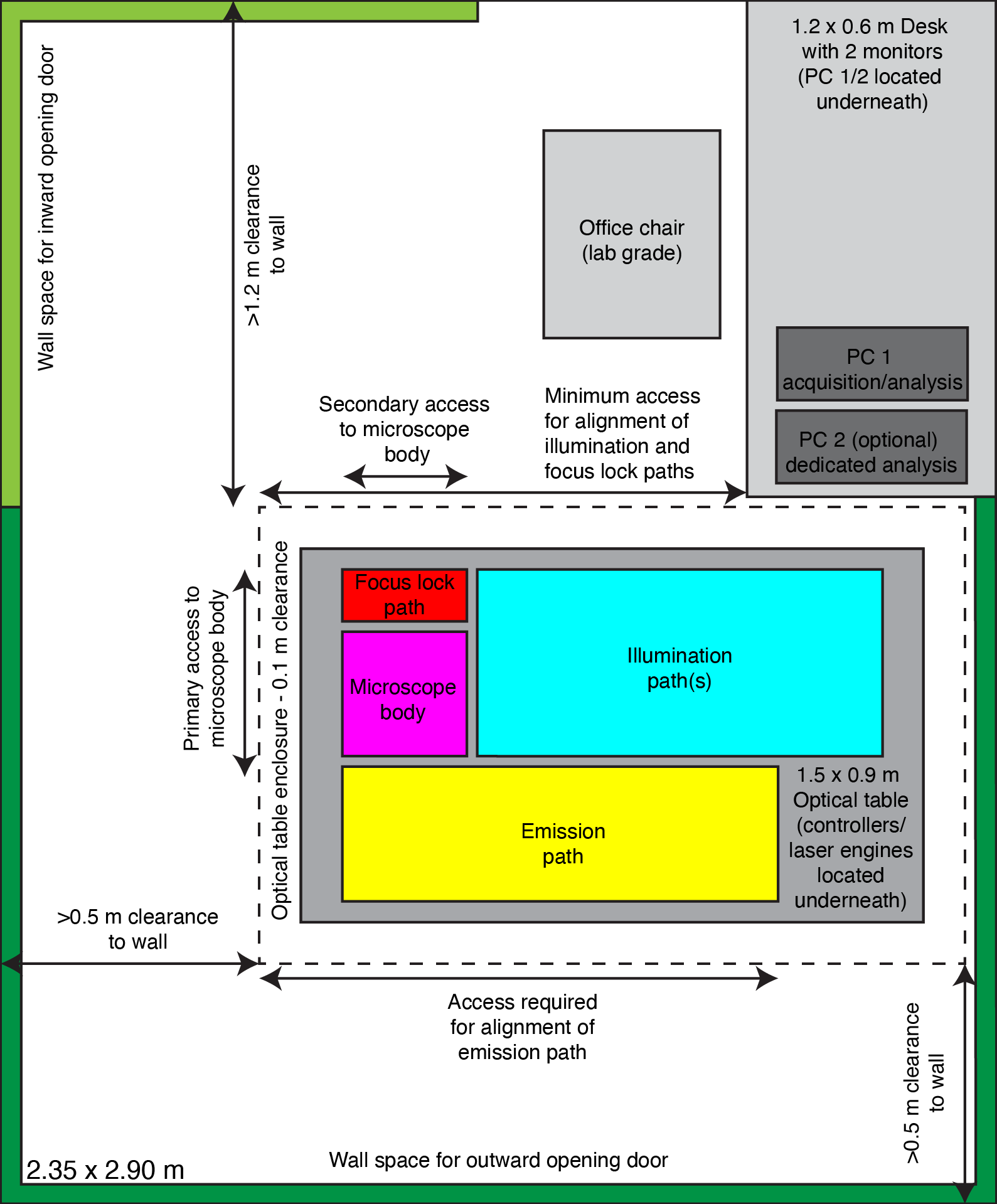

Supplementary Figure 5: An optimized layout of the microscope in a small room (2.35 × 2.90 m). Relative to the ideal layout shown in Supplementary Figure 4, this layout allows the microscope to be accommodated in a smaller room or against one wall in a larger shared lab space with access via the secondary body access point and at the cost of slightly reduced access to the illumination path. However, all components for which precise manual alignment is necessary are located in the accessible section.

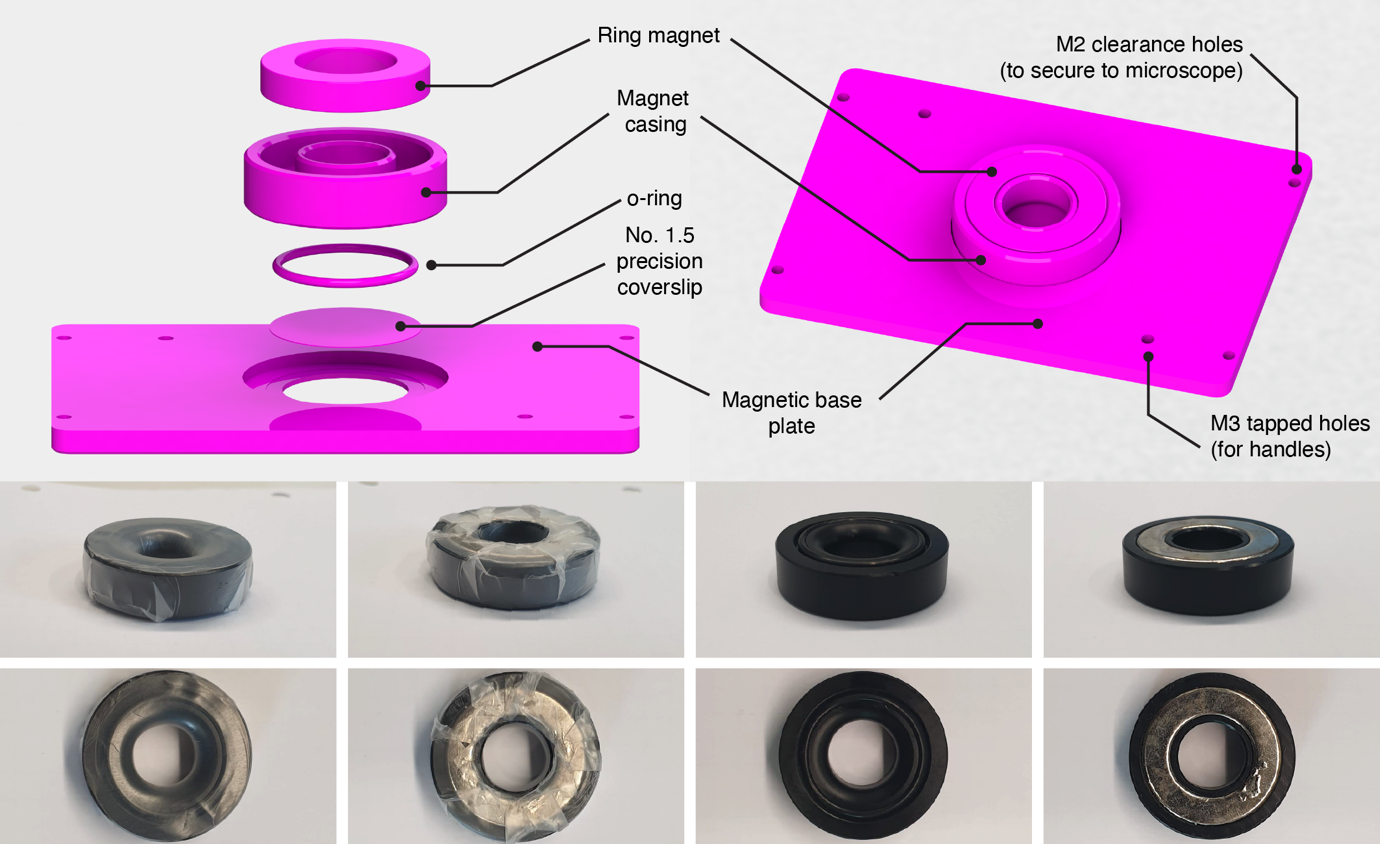

Supplementary Figure 6. The EMBL-SMLM sample holder. Top: CAD rendering of the sample holder. Left: an exploded view of the sample holder showing the constituent components. Right: an assembled view of the sample holder. Bottom: The ring magnet installed in its casing. Left: uncovered with the O-ring installed. Right: covered with parafilm.

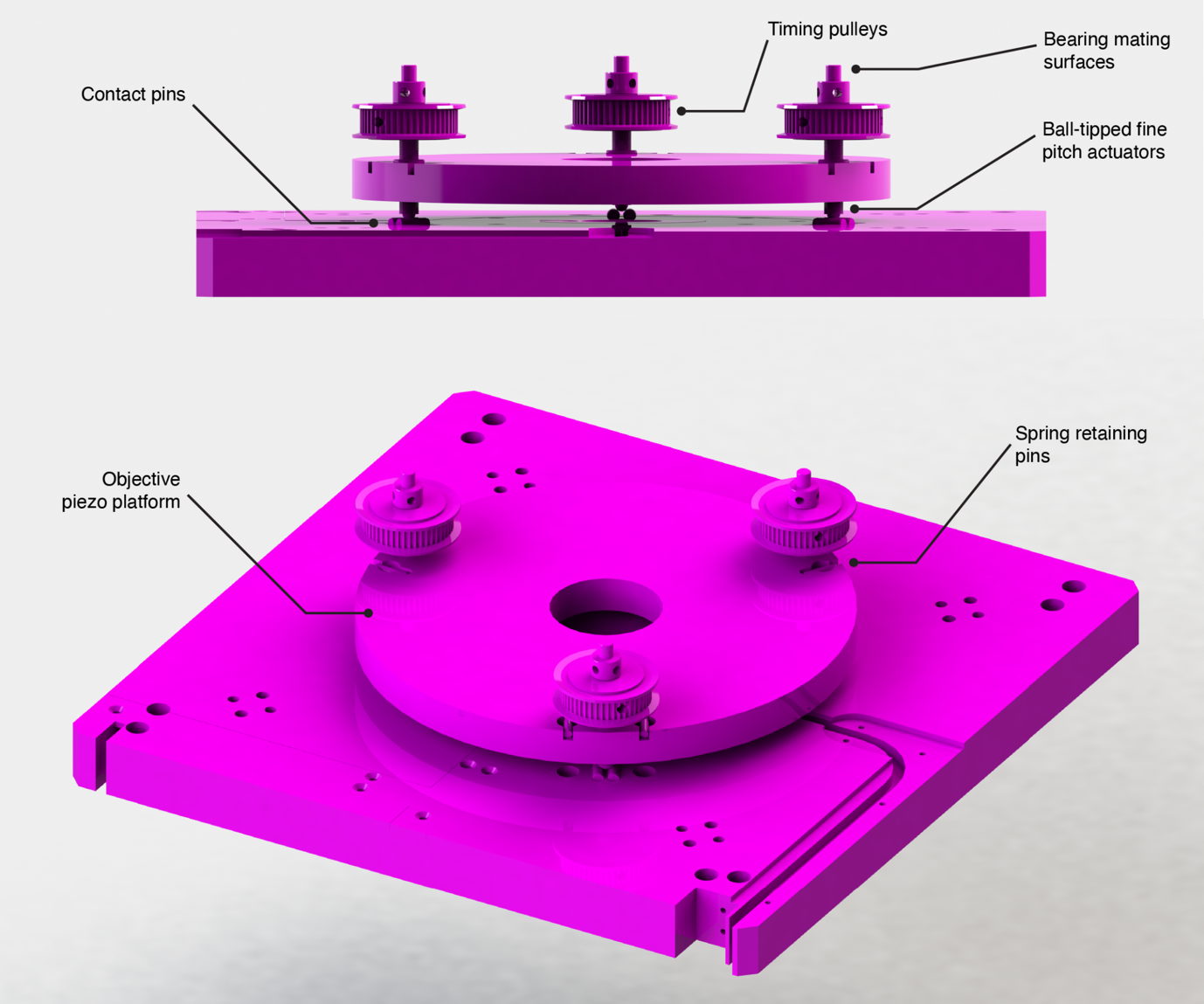

Supplementary Figure 7: Assembly of the middle section of the EMBL-SMLM microscope body. Top: a side view showing the components comprising the objective piezo platform (springs and timing belt omitted). Bottom: a top view highlighting the pins used to retain the springs and the uncovered cable channel for the piezo flexure stage.

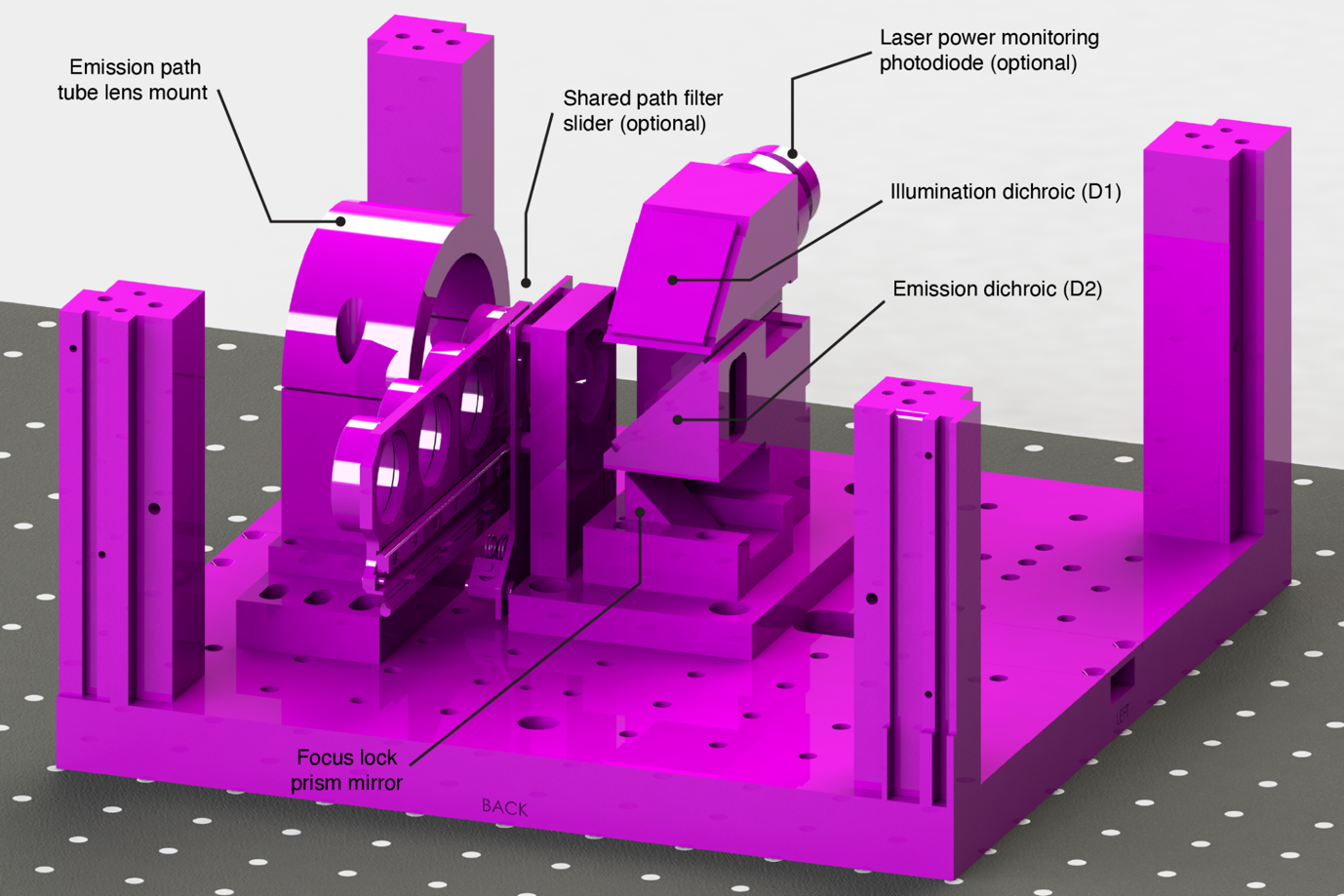

Supplementary Figure 8: Assembly of the lower section of the EMBL-SMLM microscope body (back/left view).

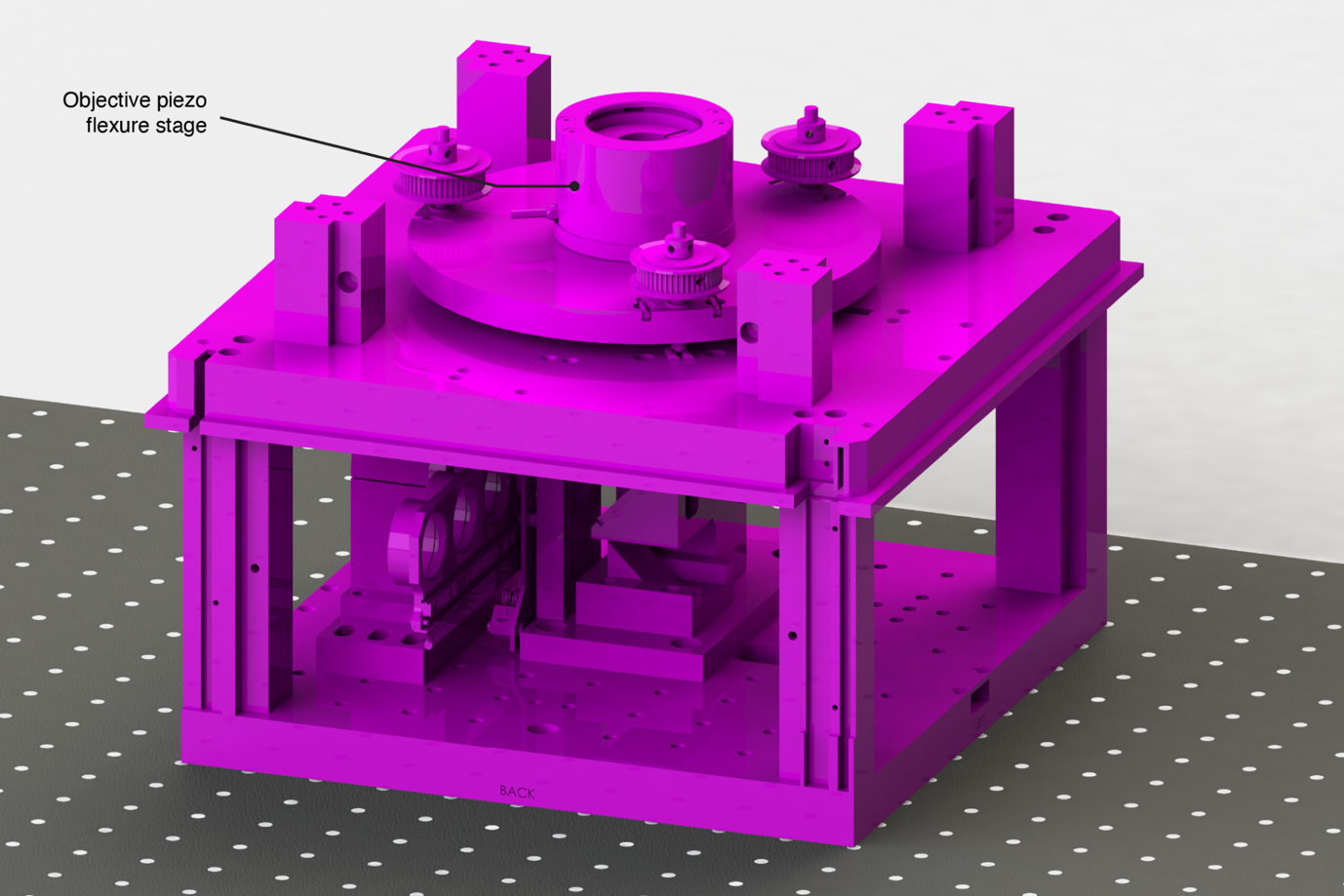

Supplementary Figure 9: Installation of the middle section of the EMBL-SMLM microscope body on top of the protruding standoffs of the lower section (back/left view). An insulating plate is placed between the two sections for thermal isolation.

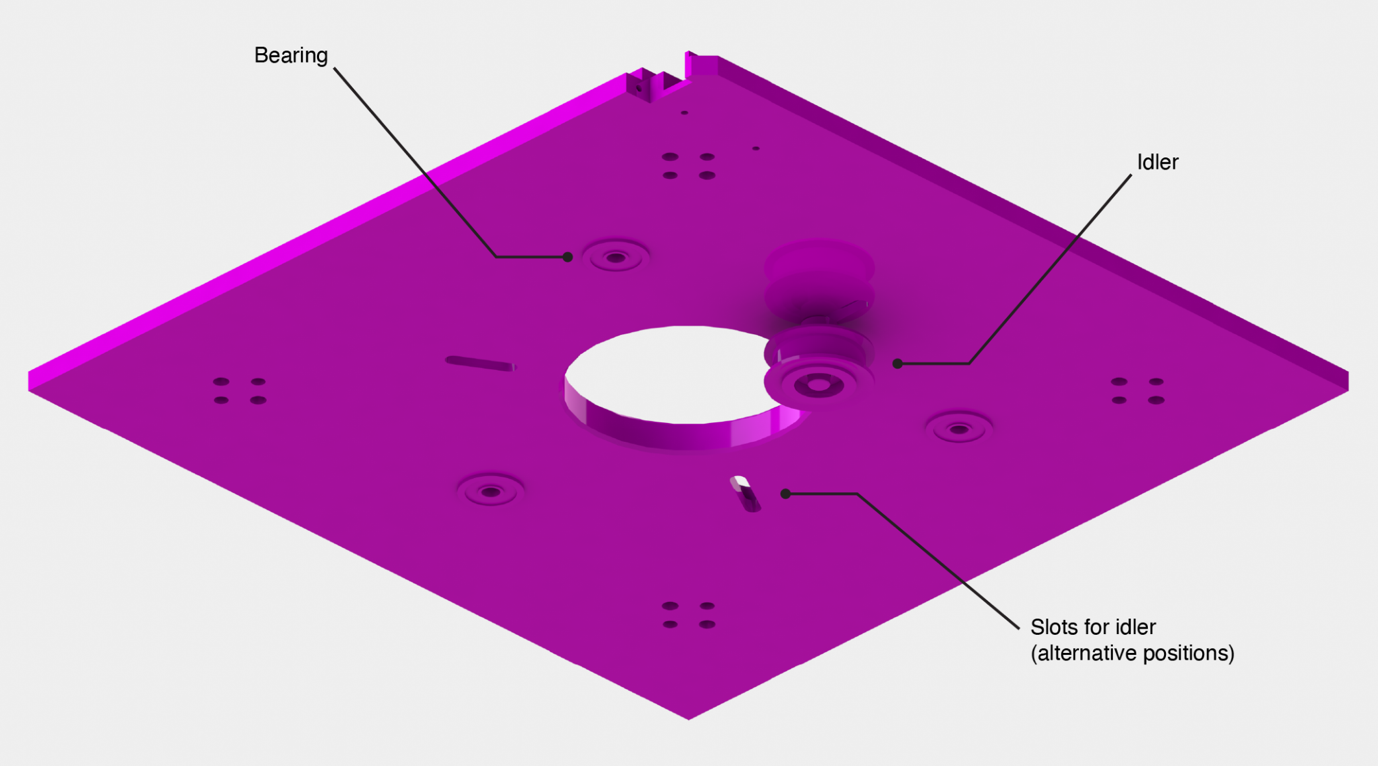

Supplementary Figure 10: The upper plate of the EMBL-SMLM microscope body, which mounts the XY stage above and mounts additional components of the objective piezo platform. The idler is positioned outside the timing belt to provide correct tensioning and is shown in the optimal location. The three bearings mate to the three fine pitch actuators shown in Supplementary Figure 7.

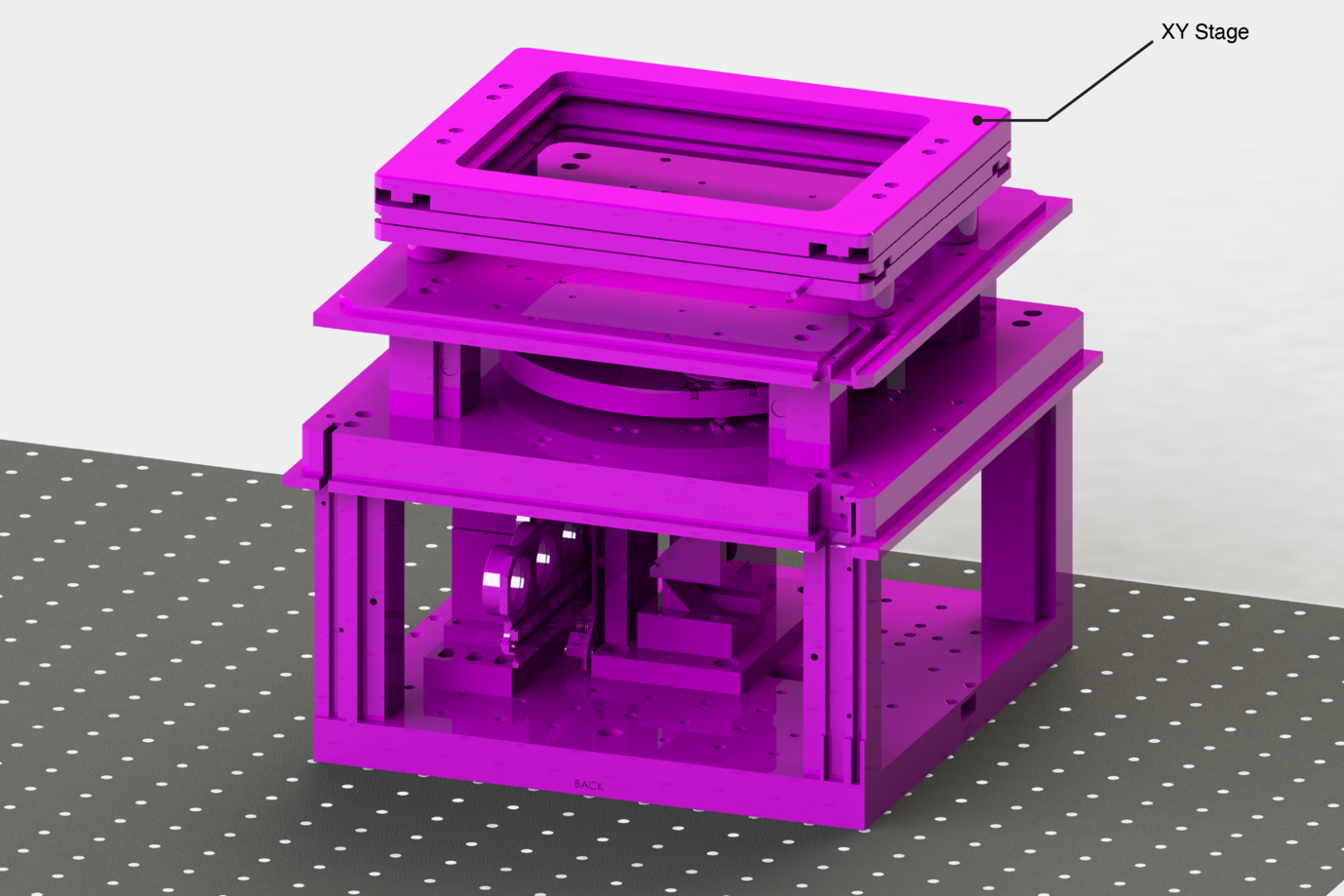

Supplementary Figure 11. The upper, middle and lower sections of the EMBL-SMLM microscope body. The upper plate and XY stage of the microscope body are mounted on top of the protruding standoffs from the middle section shown in Supplementary Figure 8.

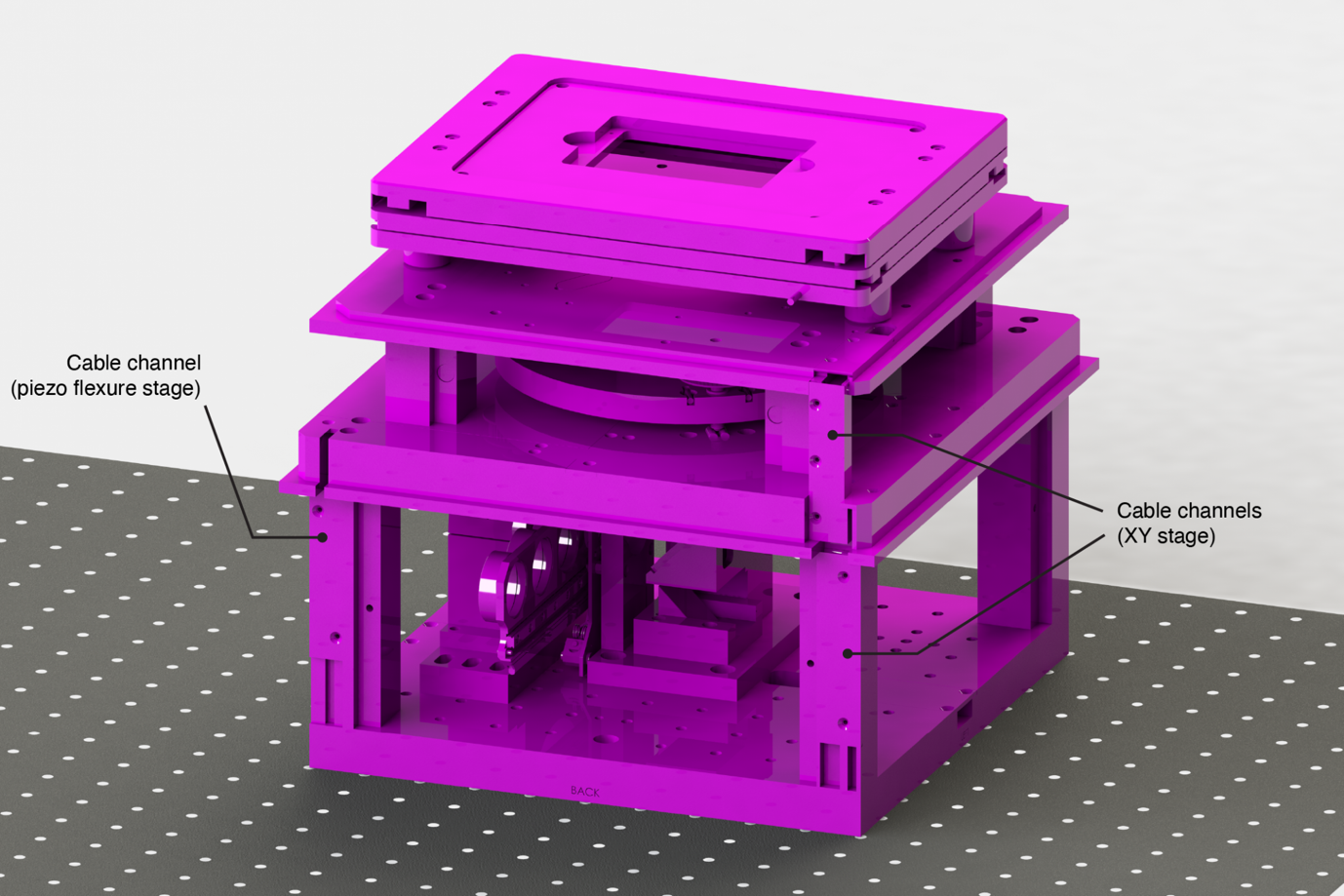

Supplementary Figure 12. Installation of additional cable routing channels on the 3D-SMLM microscope body.

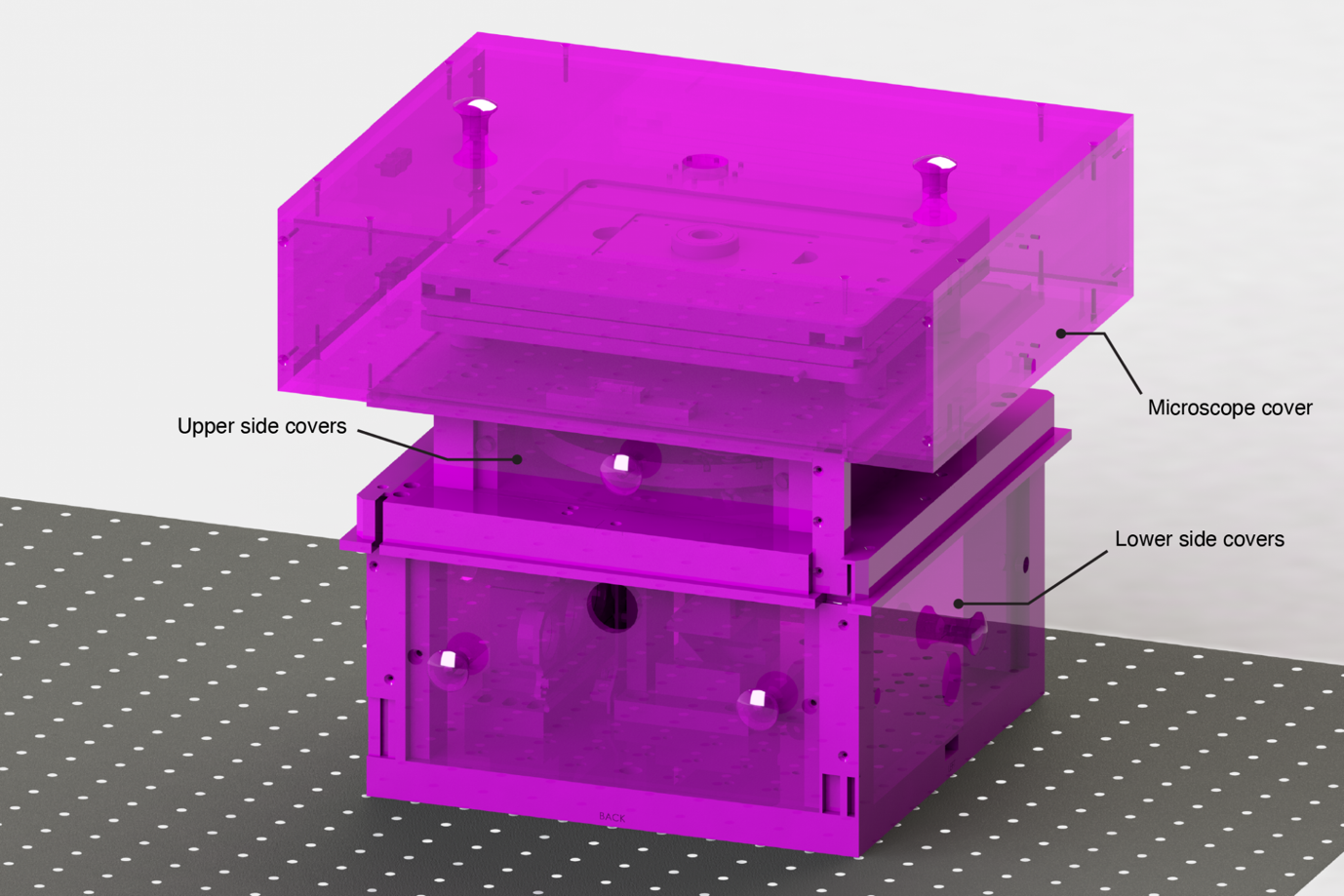

Supplementary Figure 13. Installation of the covers completes the assembly of the EMBL-SMLM microscope body

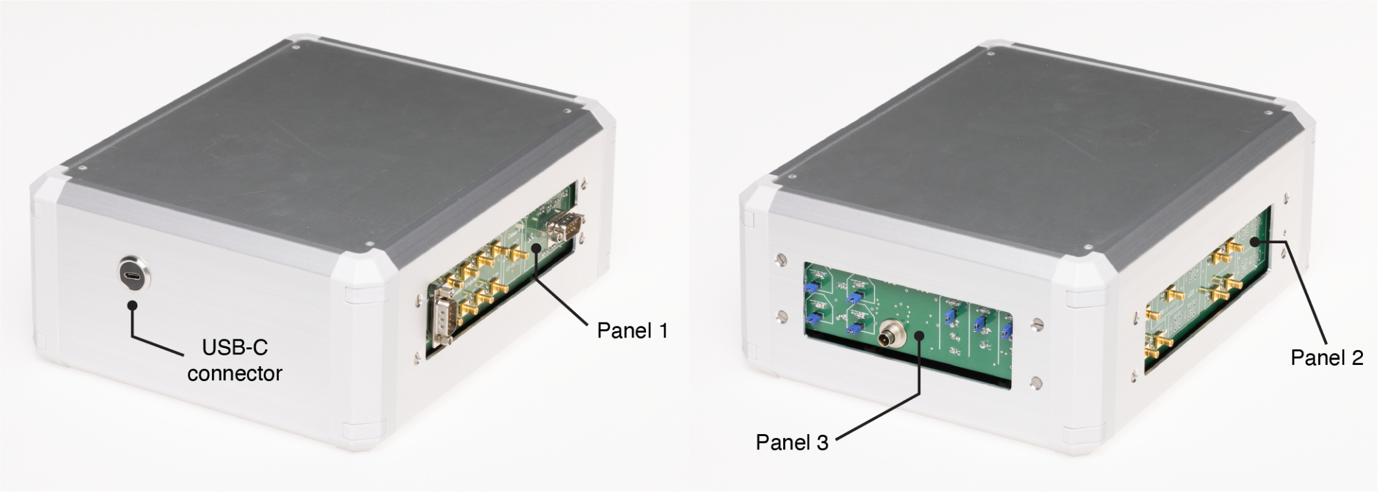

Supplementary Figure 14. A fully assembled microFPGA unit showing the three front panel boards and USB-C connector.

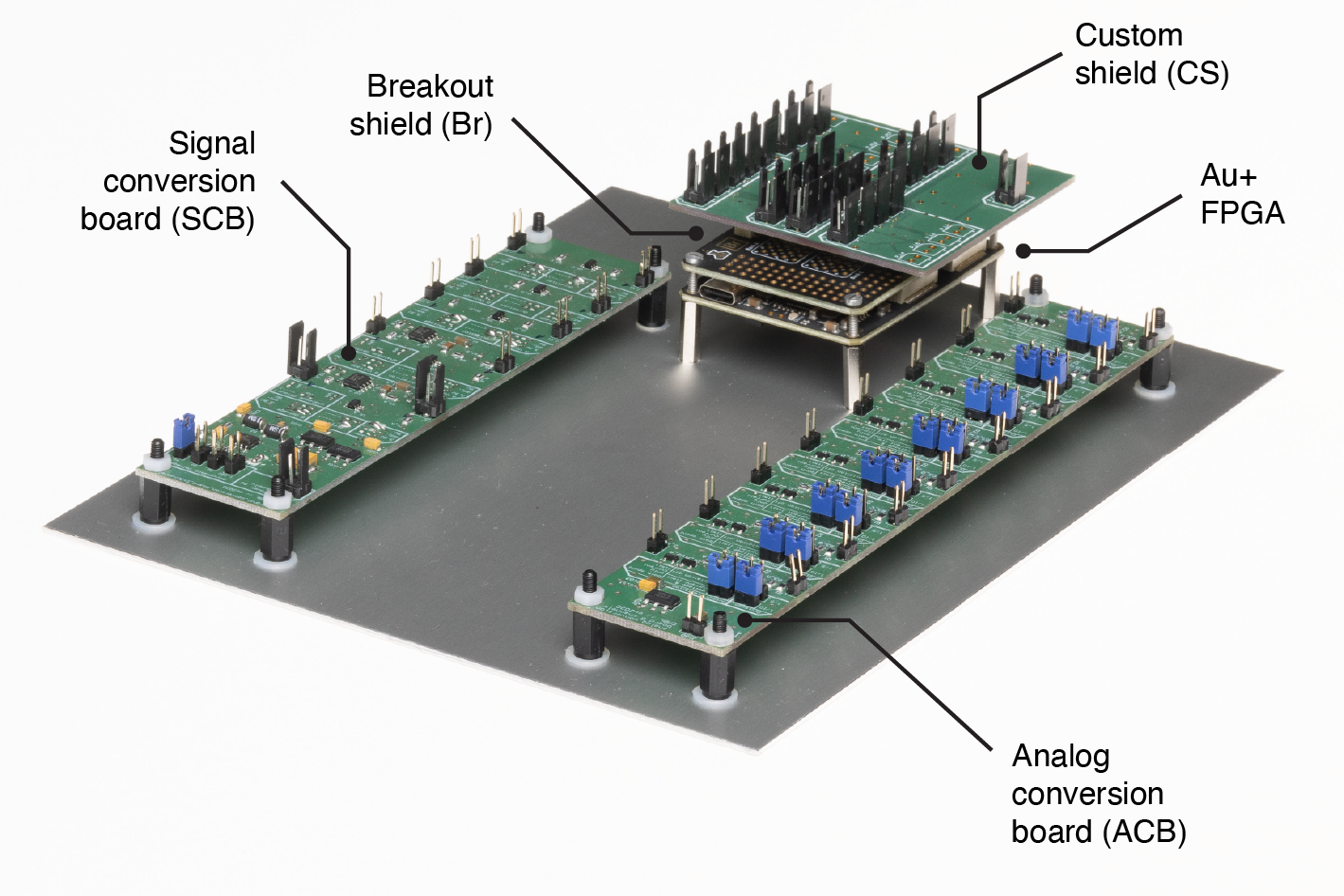

Supplementary Figure 15. Assembly of the microFPGA. The signal and analog conversion boards, Alchitry Au FPGA, FPGA shield and FPGA custom shield are mounted to the base plate via various hex standoffs.

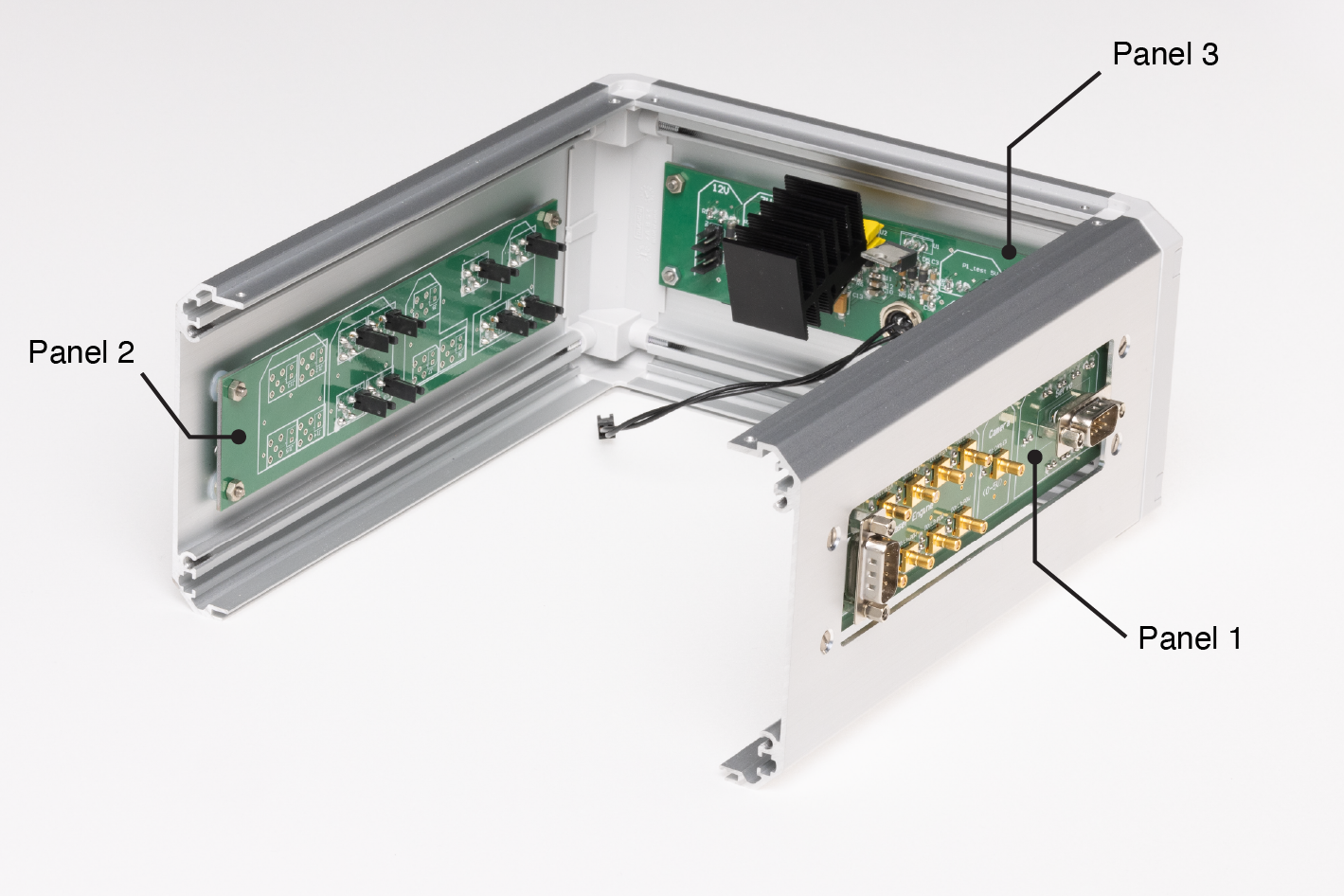

Supplementary Figure 16. Assembly of the microFPGA. The front panel boards are mounted to the modified side panels of the housing.

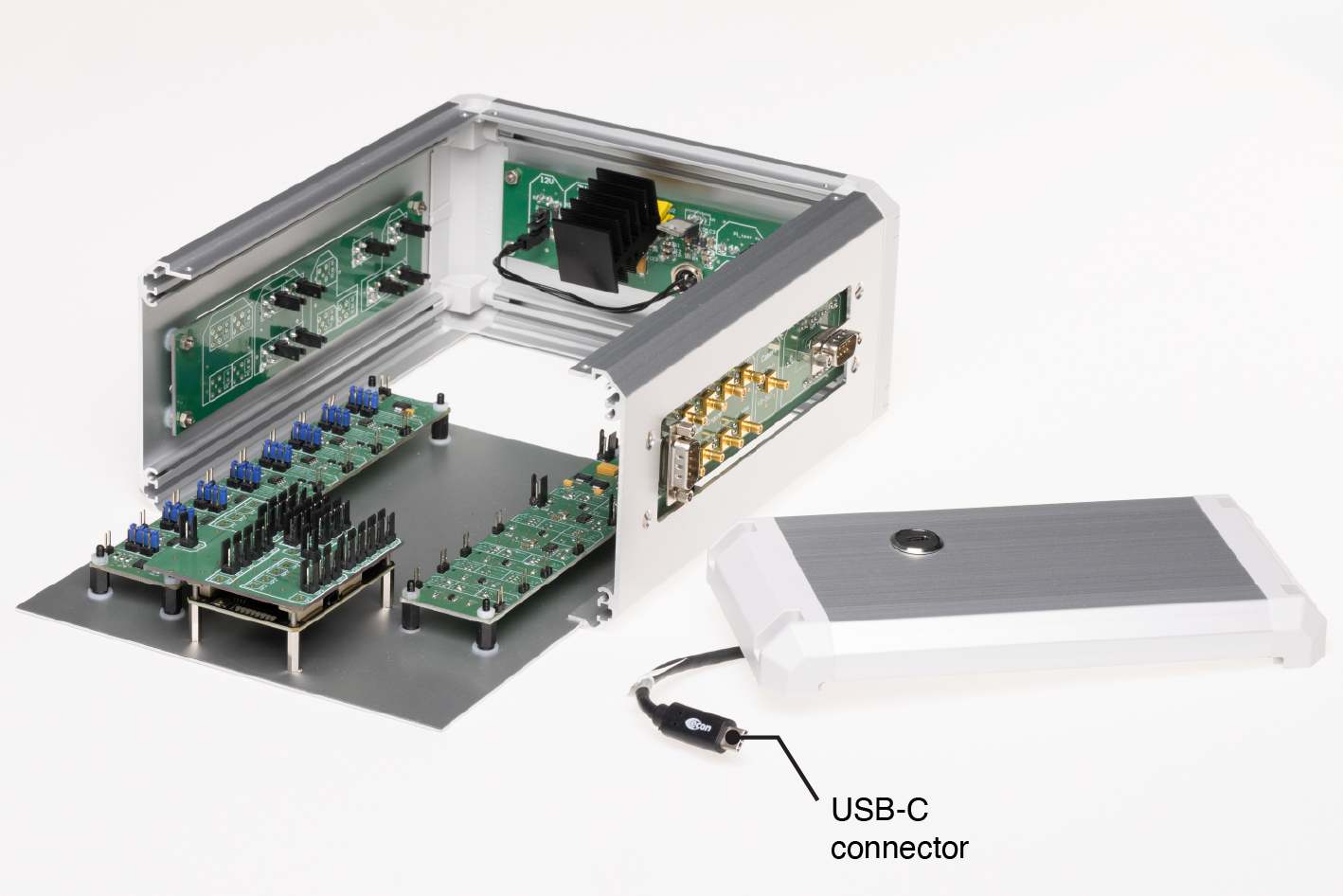

Supplementary Figure 17. Assembly of the microFPGA. The USB connector is installed in the one remaining side panel and the base plate is mounted inside the housing.

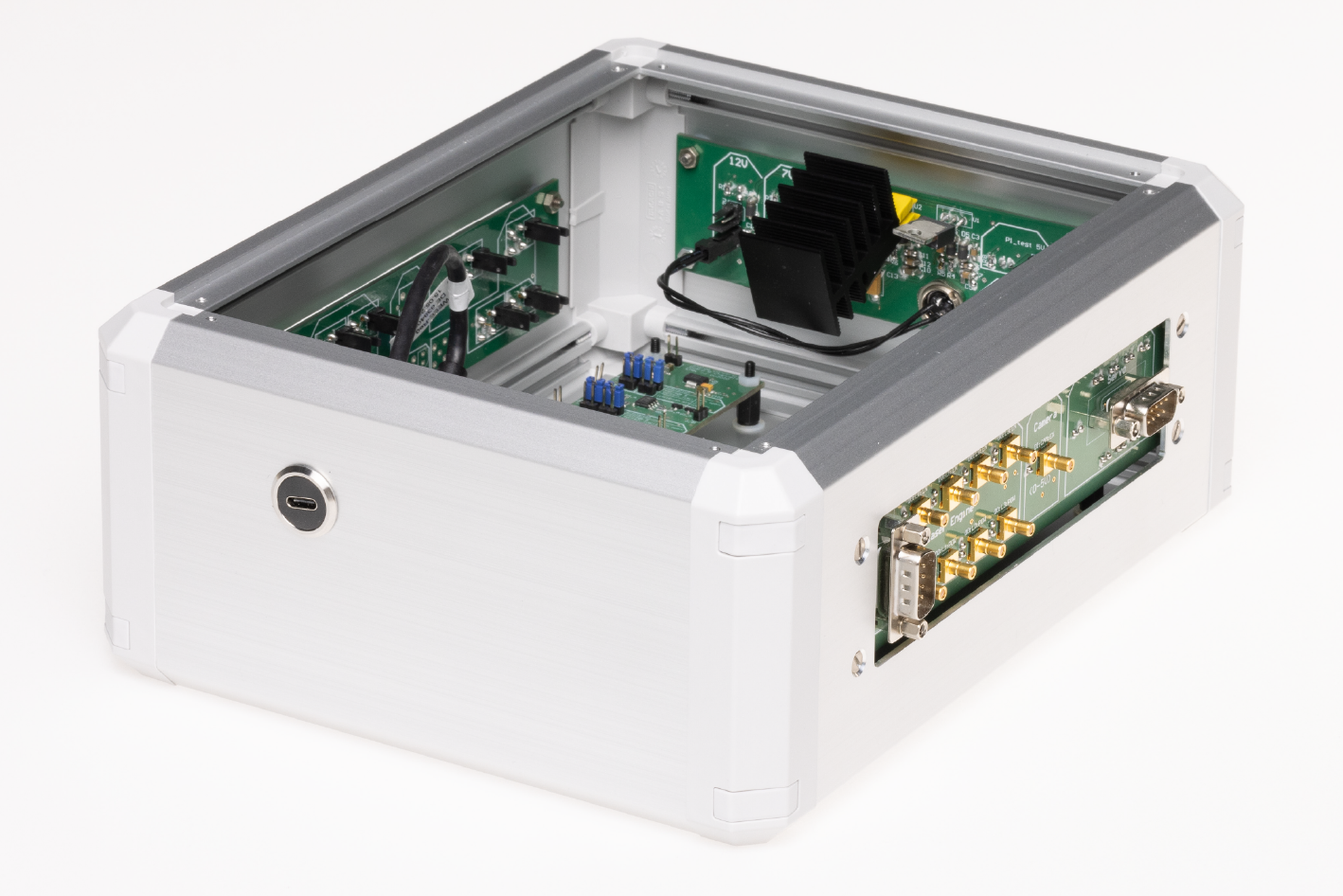

Supplementary Figure 18. Assembly of the microFPGA. The side panel with the USB-C connector is mounted to the box and connected to the Alchitry Au FPGA.

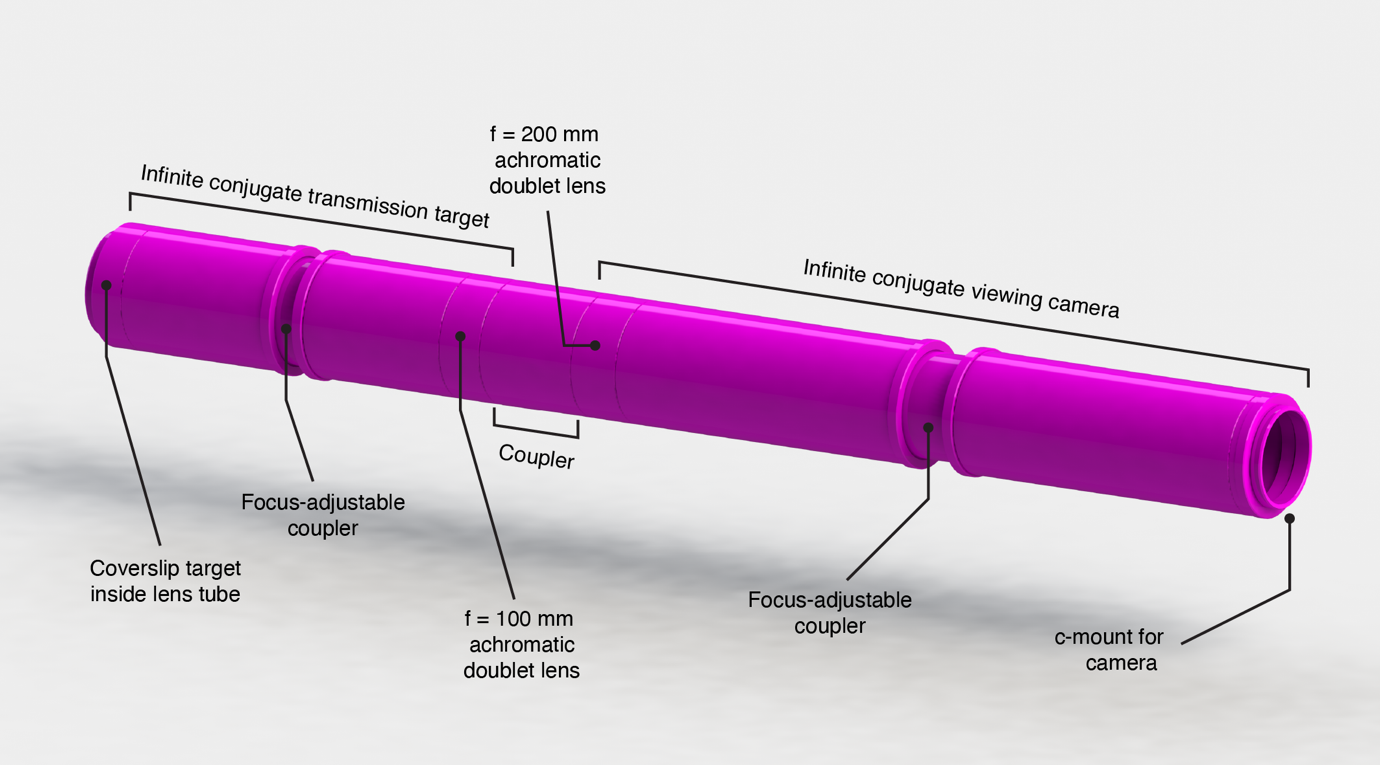

Supplementary Figure 19. The coupled infinite conjugate transmission target and infinite conjugate viewing camera assemblies. The camera to be supplied by the builder is omitted but a c-mount adapter provides appropriate spacing for the adjustable focus.

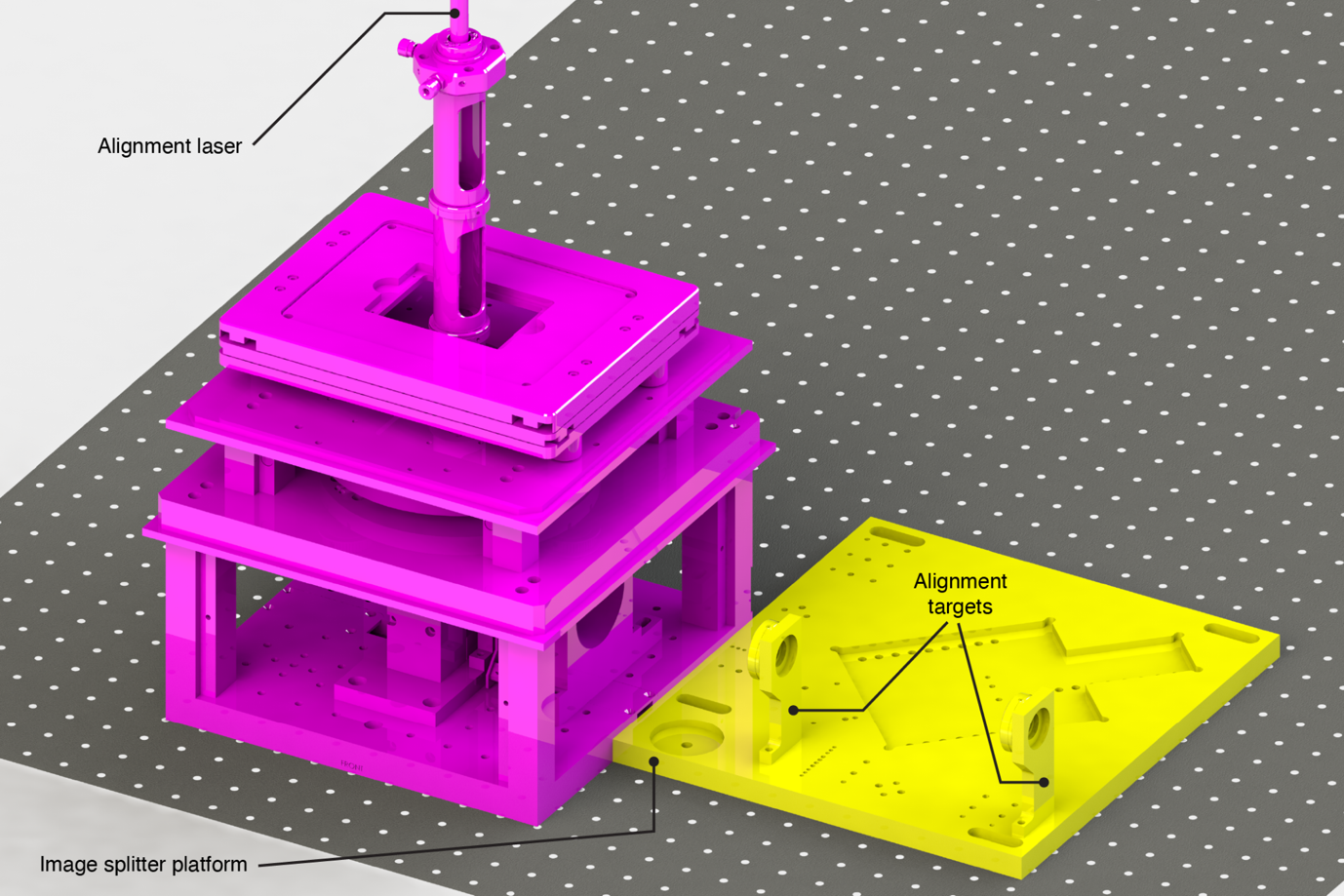

Supplementary Figure 20. Installation of the alignment laser and image splitter platform with alignment targets to align the emission path out of the EMBL-SMLM microscope body. The laser is used as a proxy for the emission path optical axis defined by the objective lens.

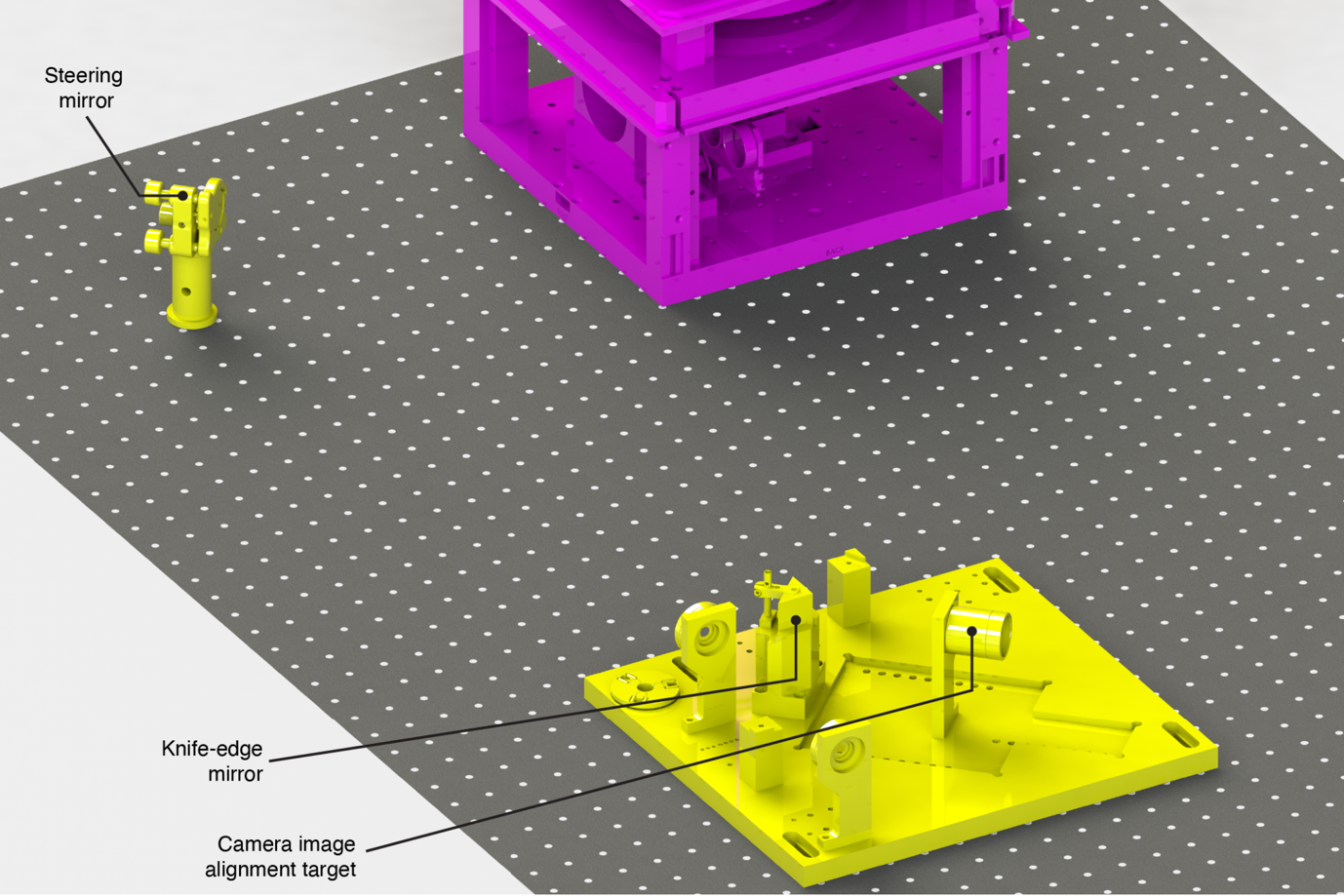

Supplementary Figure 21. Placement of the first steering mirror in the emission path to align through the alignment targets of the image splitter platform. The image platform is installed in its nominal position for imaging with the knife-edge mirror and camera image alignment target pre-installed.

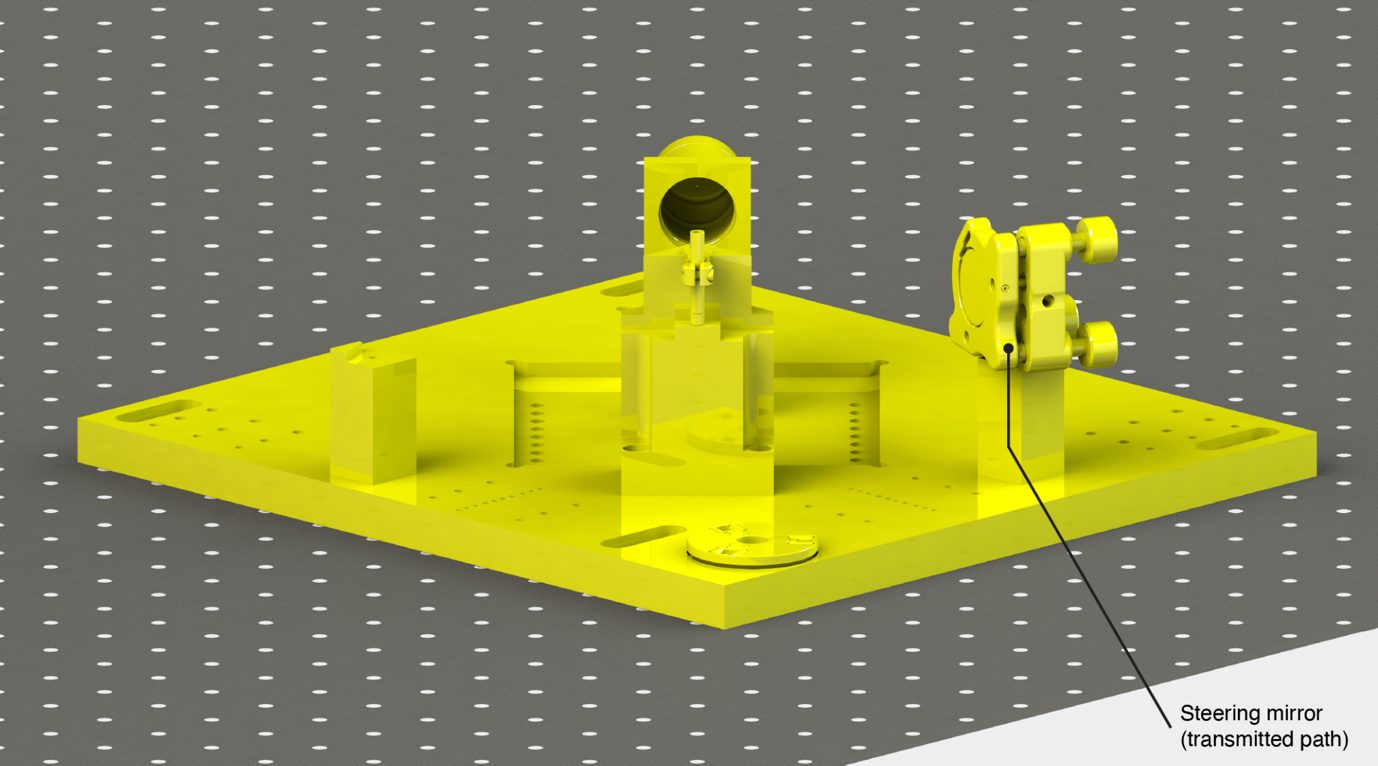

Supplementary Figure 22. Installation and alignment of the emission path second steering mirror in the transmitted path of the image splitter platform. The mirror is steered to route the alignment laser through the camera image alignment target.

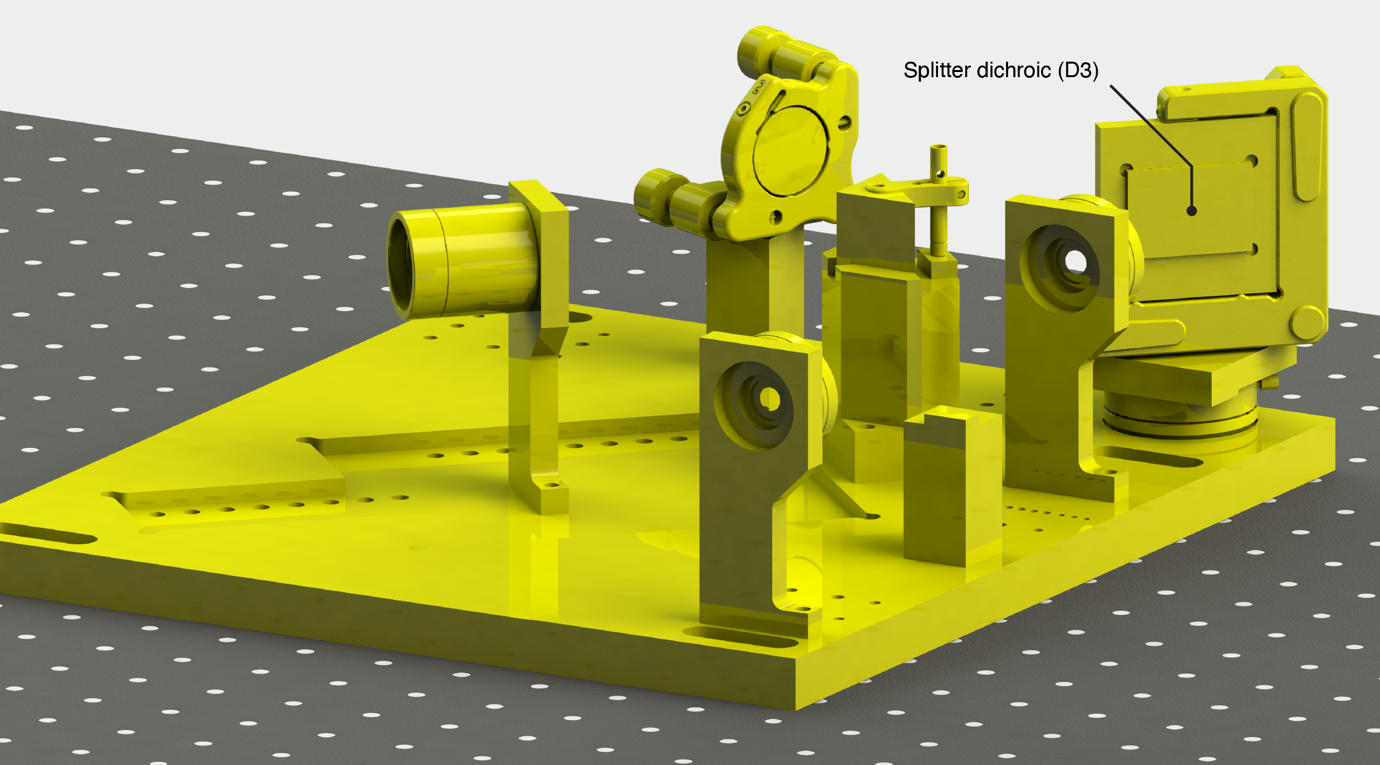

Supplementary Figure 23. Installation and alignment of the image splitter dichroic mirror. The dichroic is steered to route the alignment laser through the alignment targets.

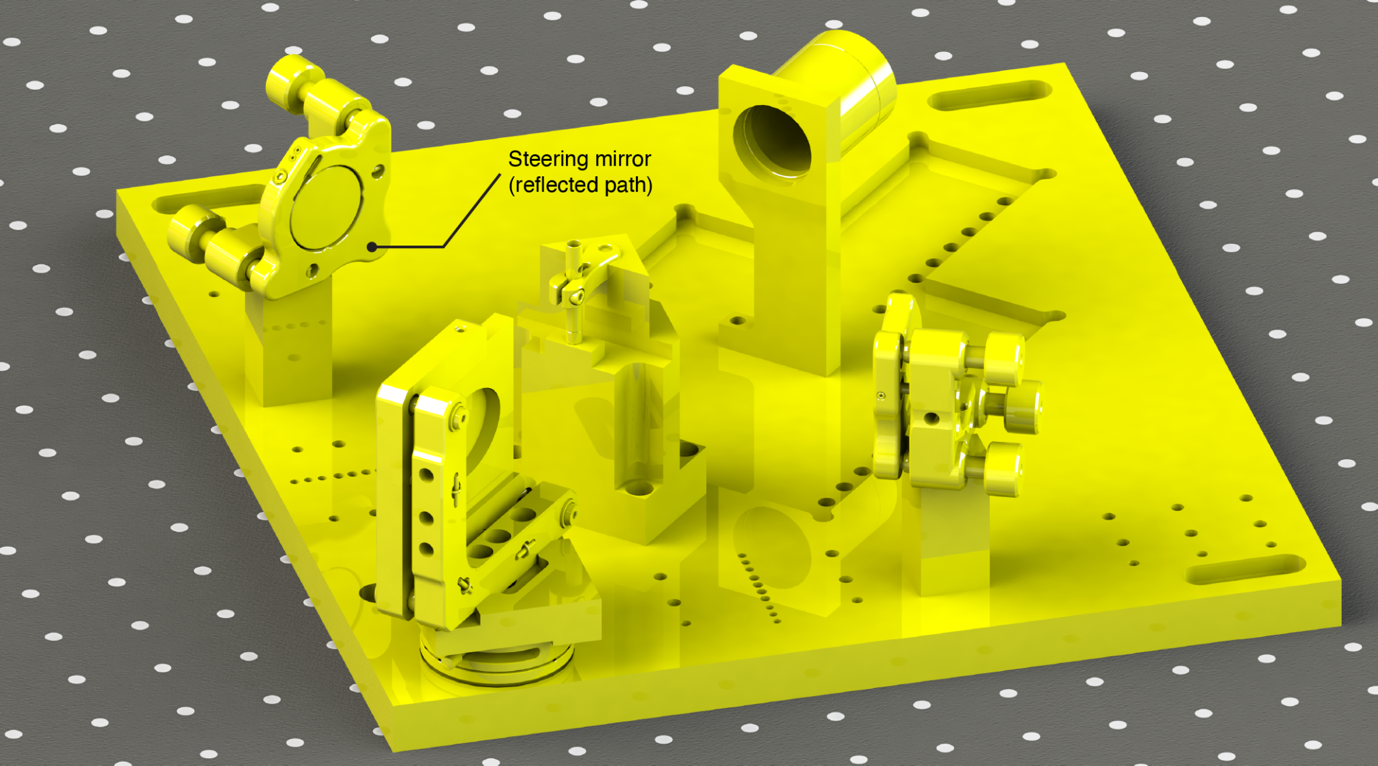

Supplementary Figure 24. Installation and alignment of the emission path second steering mirror in the reflected path of the image splitter platform. The mirror is steered to route the alignment laser through the camera image alignment target.

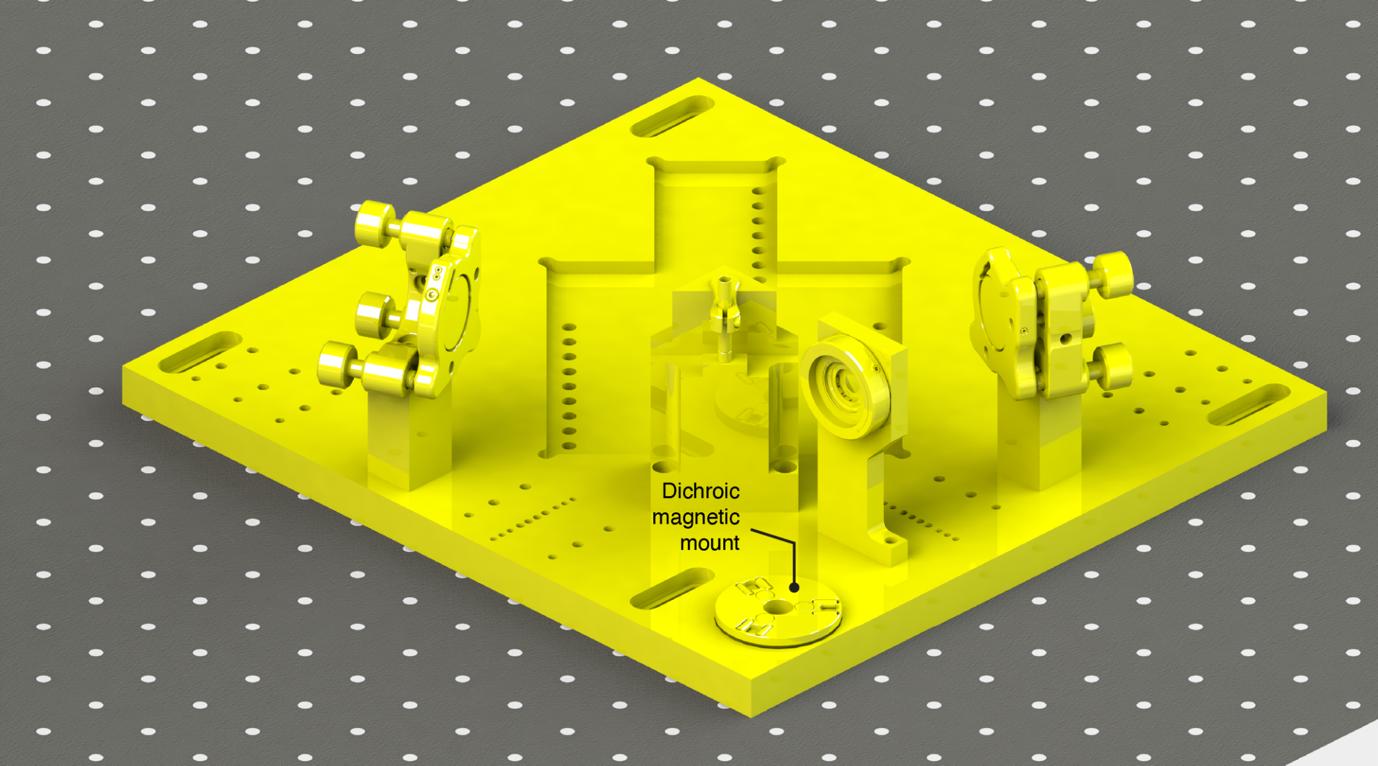

Supplementary Figure 25. The image splitter dichroic mirror is removed to use the transmitted path of the image splitter to judge alignment of the emission path tube lens further upstream.

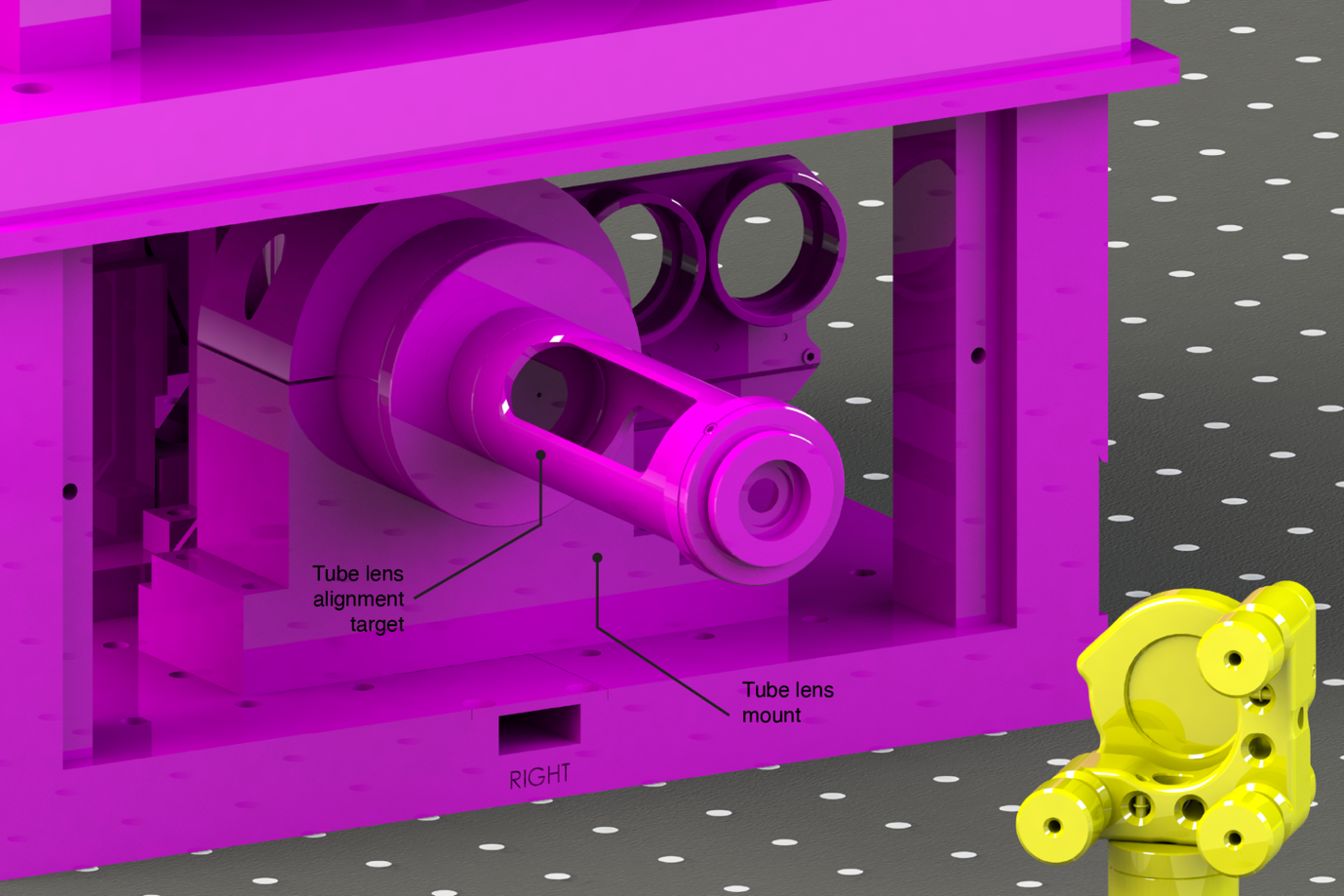
Supplementary Figure 26. The tube lens mount is aligned to the alignment laser using the tube lens alignment target comprising a spaced ground glass alignment disk and adjustable iris.

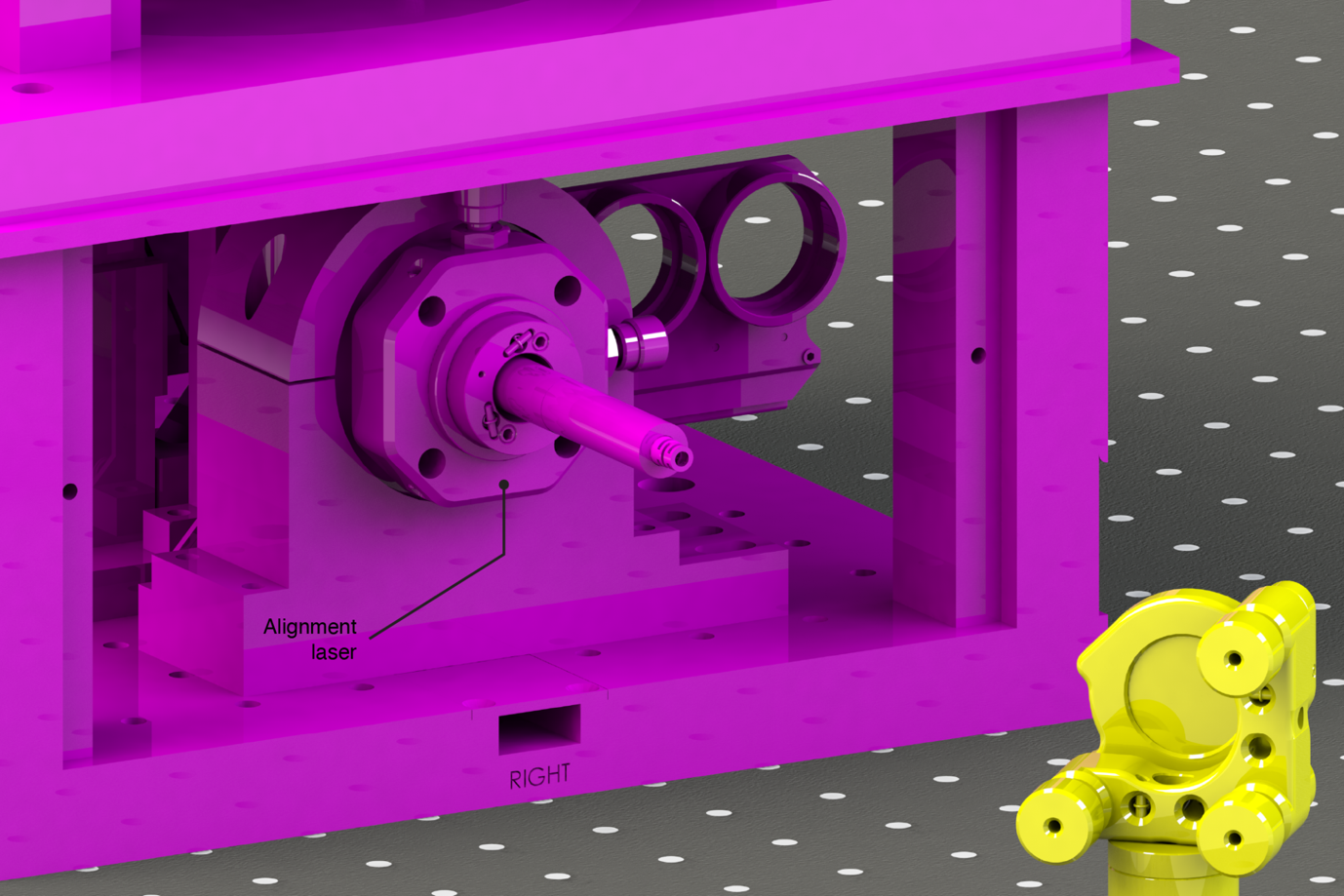

Supplementary Figure 27. The emission path tube lens is installed in its mount (occluded). The alignment laser is installed on the protruding thread of the tube lens. The tube lens is then positioned along the axis of the mount to focus to the back focal plane of the objective lens, thus producing a collimated output.

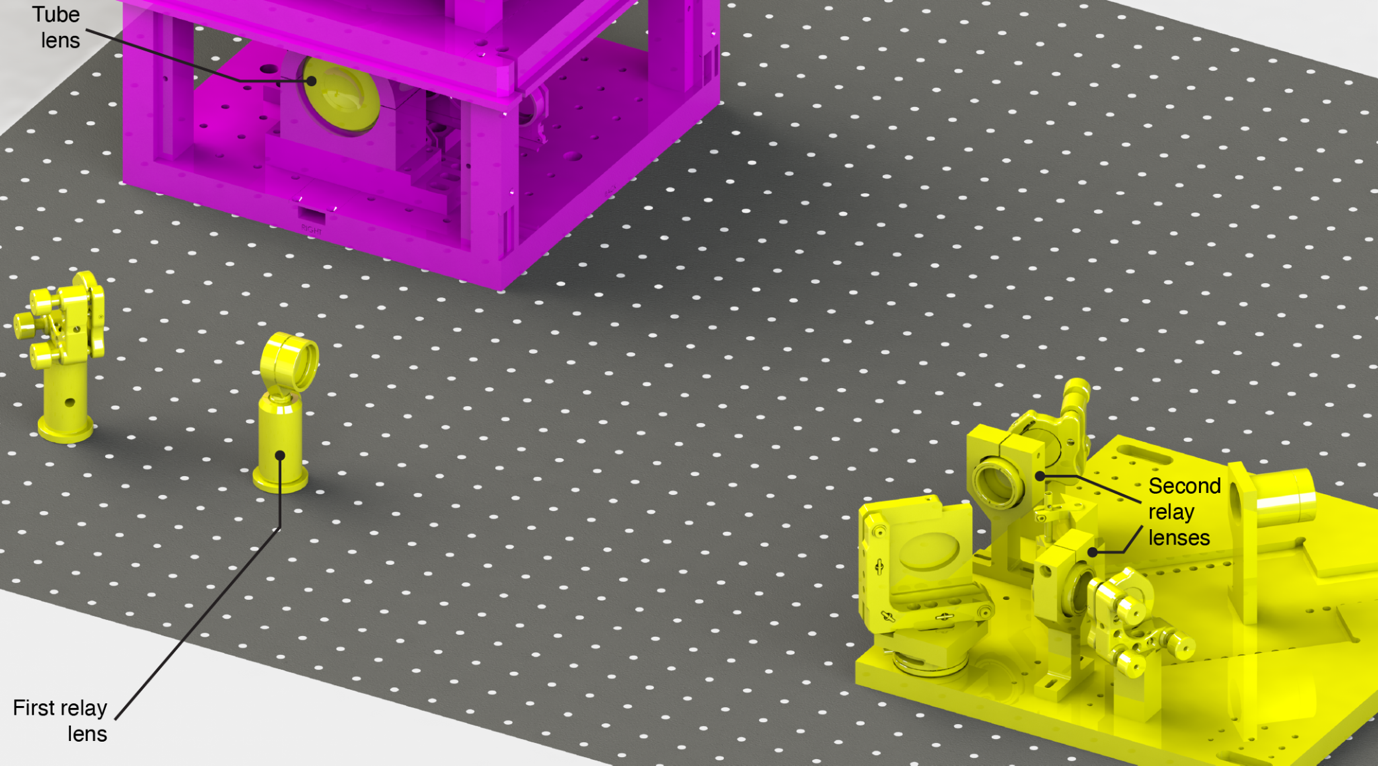

Supplementary Figure 28. Installation of the relay lenses in the emission path. With the alignment laser returned to its primary location at the objective lens port, the tube lens can be seen in its mount .

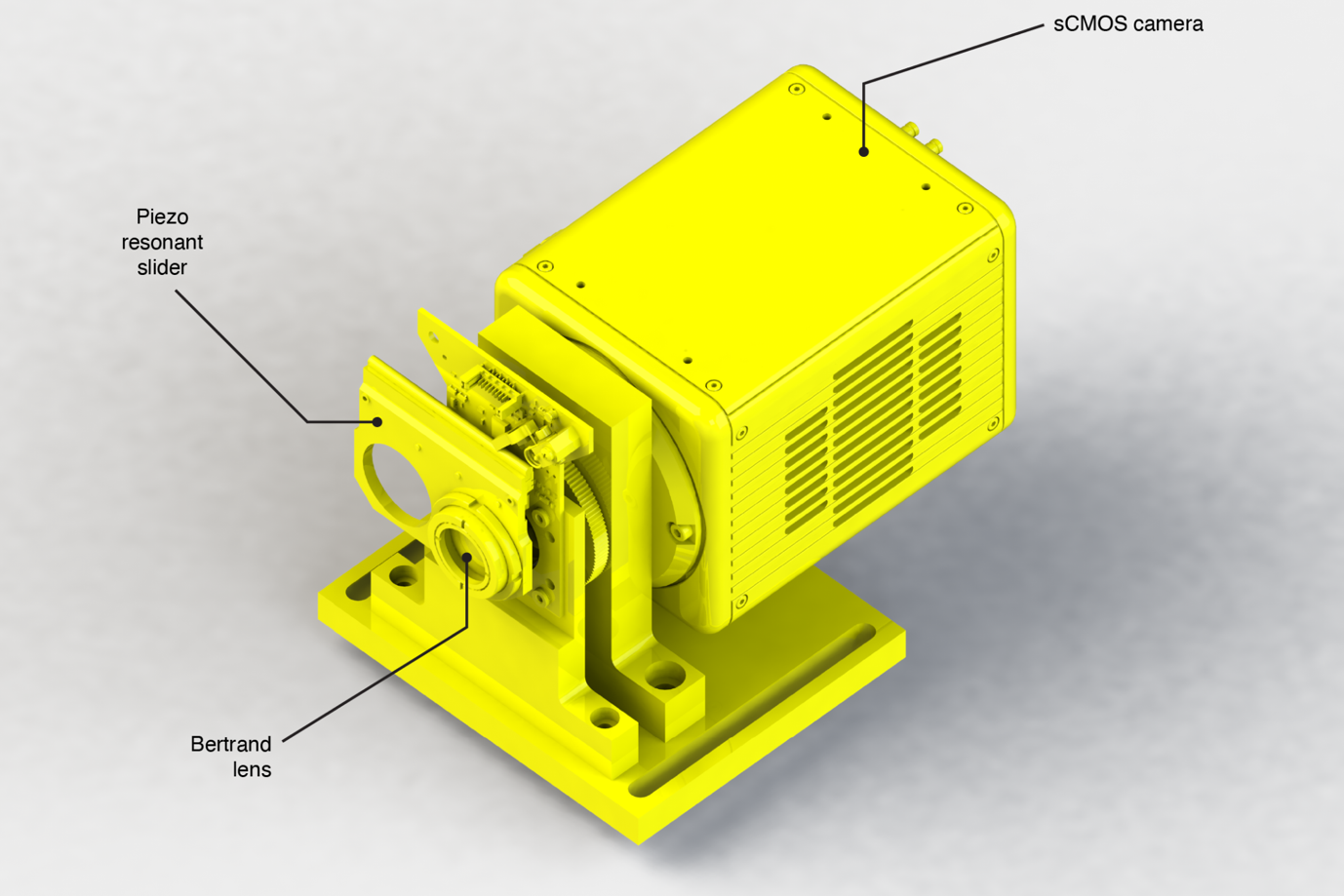

Supplementary Figure 29. The sCMOS camera and Bertrand lens assembly. The Bertrand lens is nominally spaced to produce an image of an infinite conjugate object on the camera but the focus can be adjusted. The Bertand lens can be moved in/out of the emission path using the two-position piezo resonant slider.

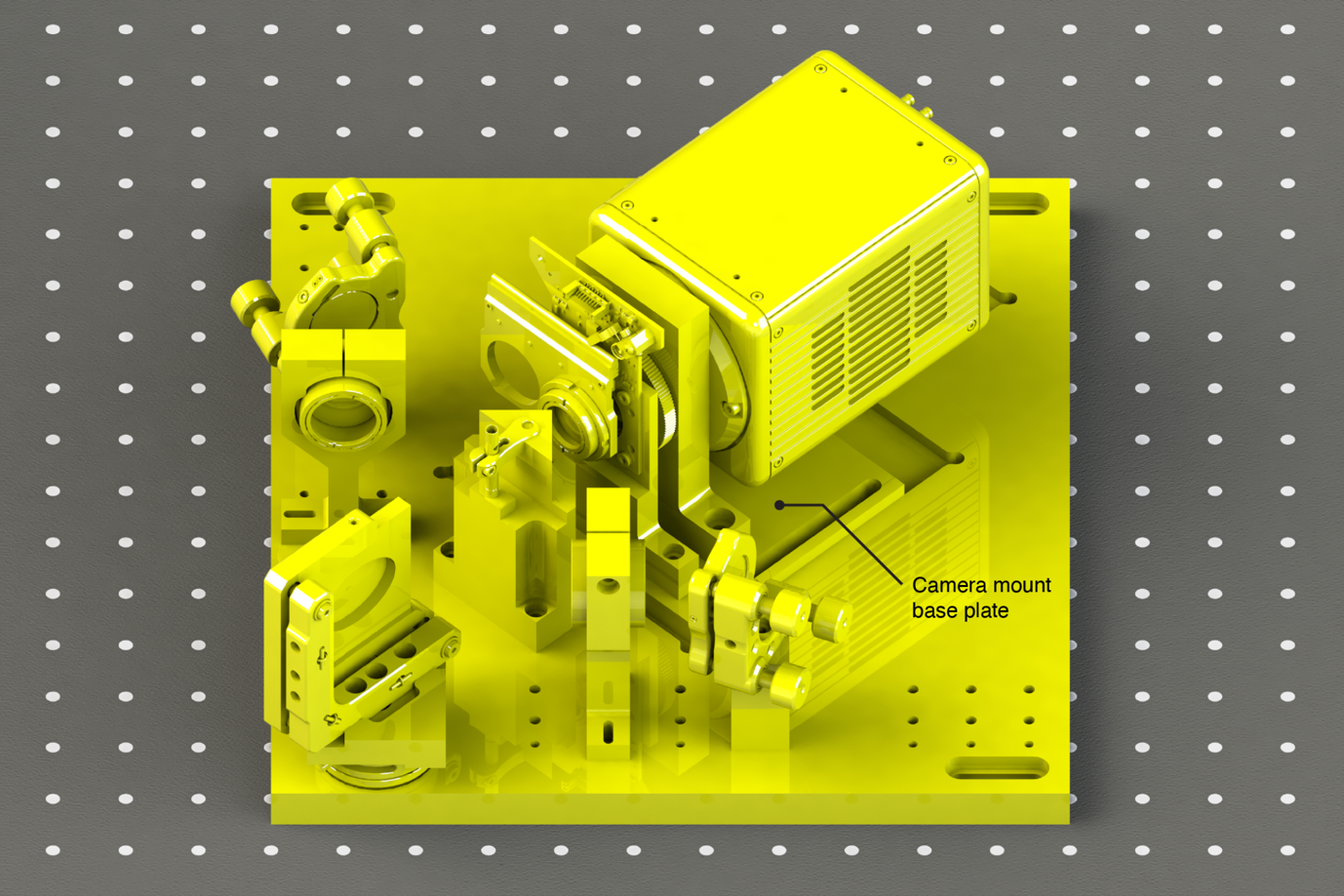
Supplementary Figure 30. The sCMOS camera and Bertrand lens installed in the nominal position on the image splitter platform.

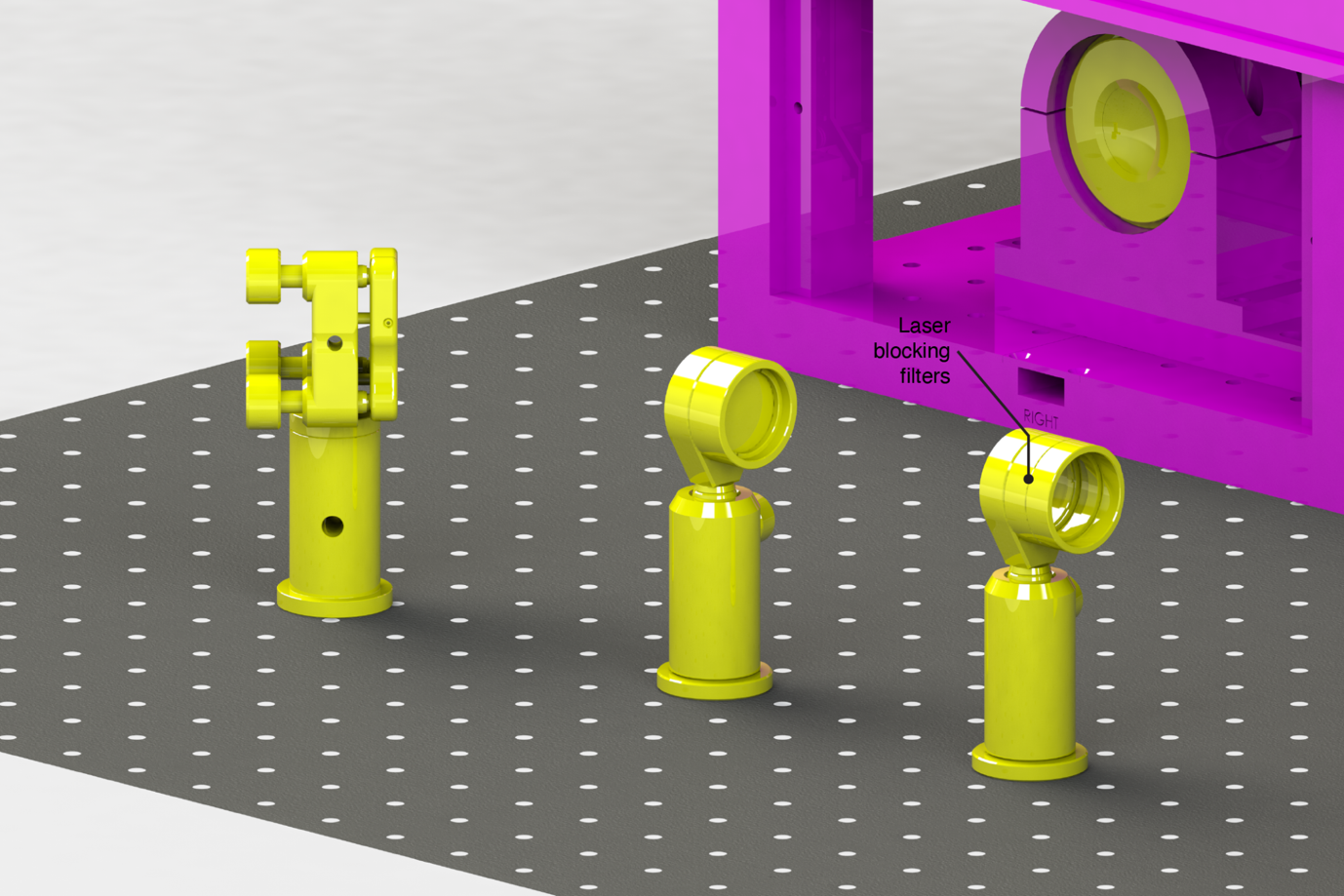
Supplementary Figure 31. Installation of the laser blocking filters in the secondary infinity space of the emission path. Two filters are installed in lens tubes to block the activation/excitation and focus lock lasers respectively.

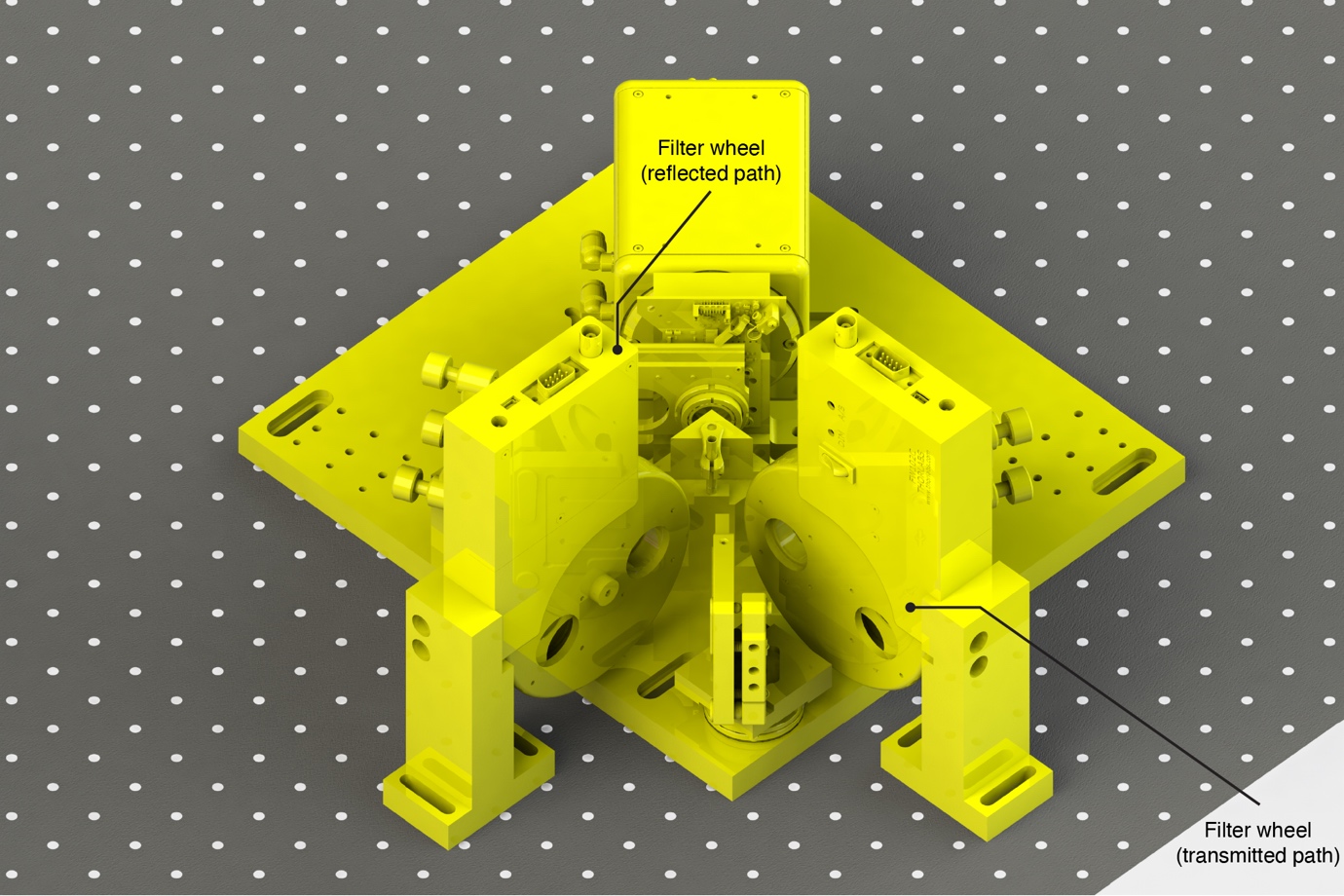

Supplementary Figure 32. Installation of the filter wheels in the image splitter transmitted (longwave) and reflected (shortwave) paths.

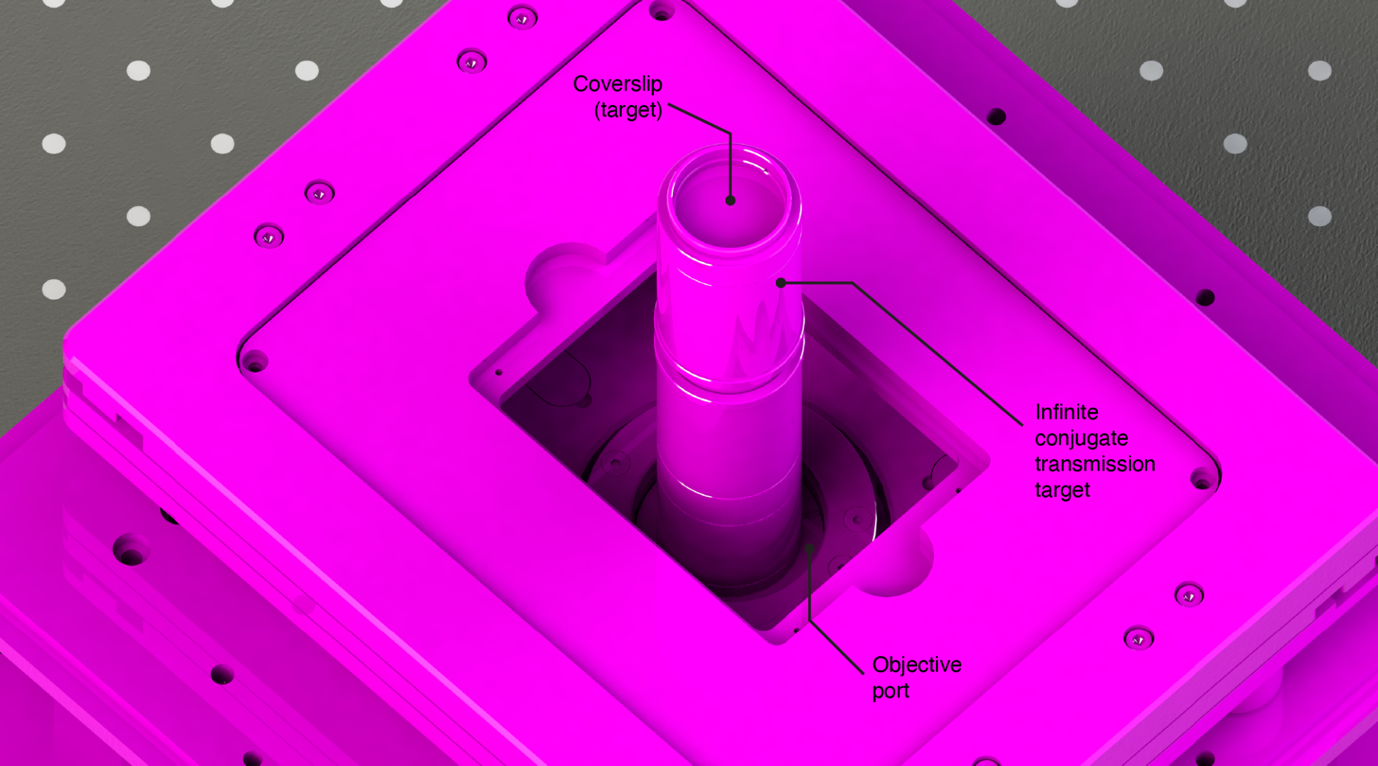

Supplementary Figure 33. Installation of the infinite conjugate transmission target at the objective port provides a reference to an optical infinity to correctly position the camera to focus at the native infinite conjugate focus of the objective lens.

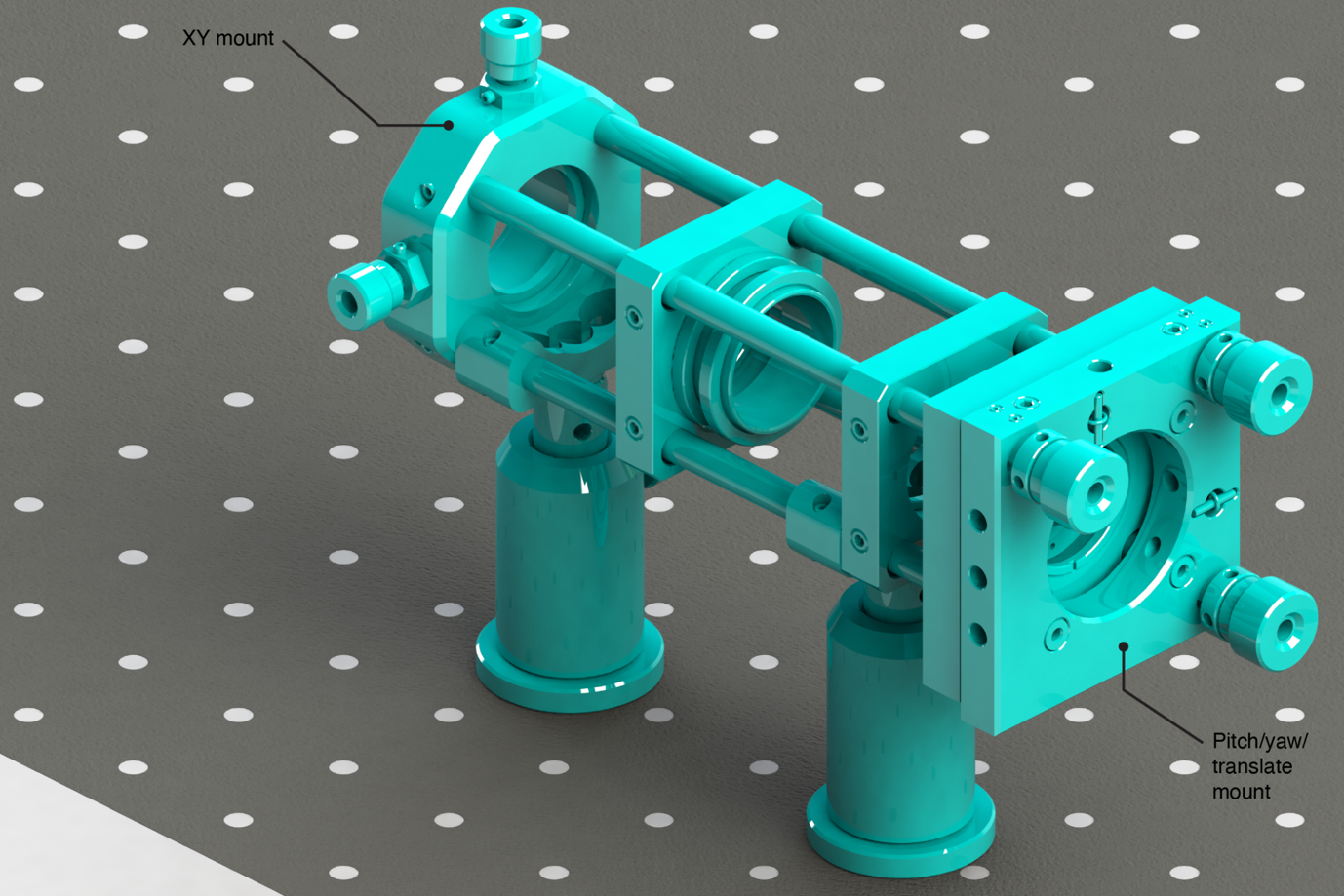

Supplementary Figure 34. The single-mode laser engine beam launch (with lenses omitted). The fiber (omitted) is mounted to a terminating adapter and the diverging output is aligned through the cage system using the pitch/yaw/translate mount.

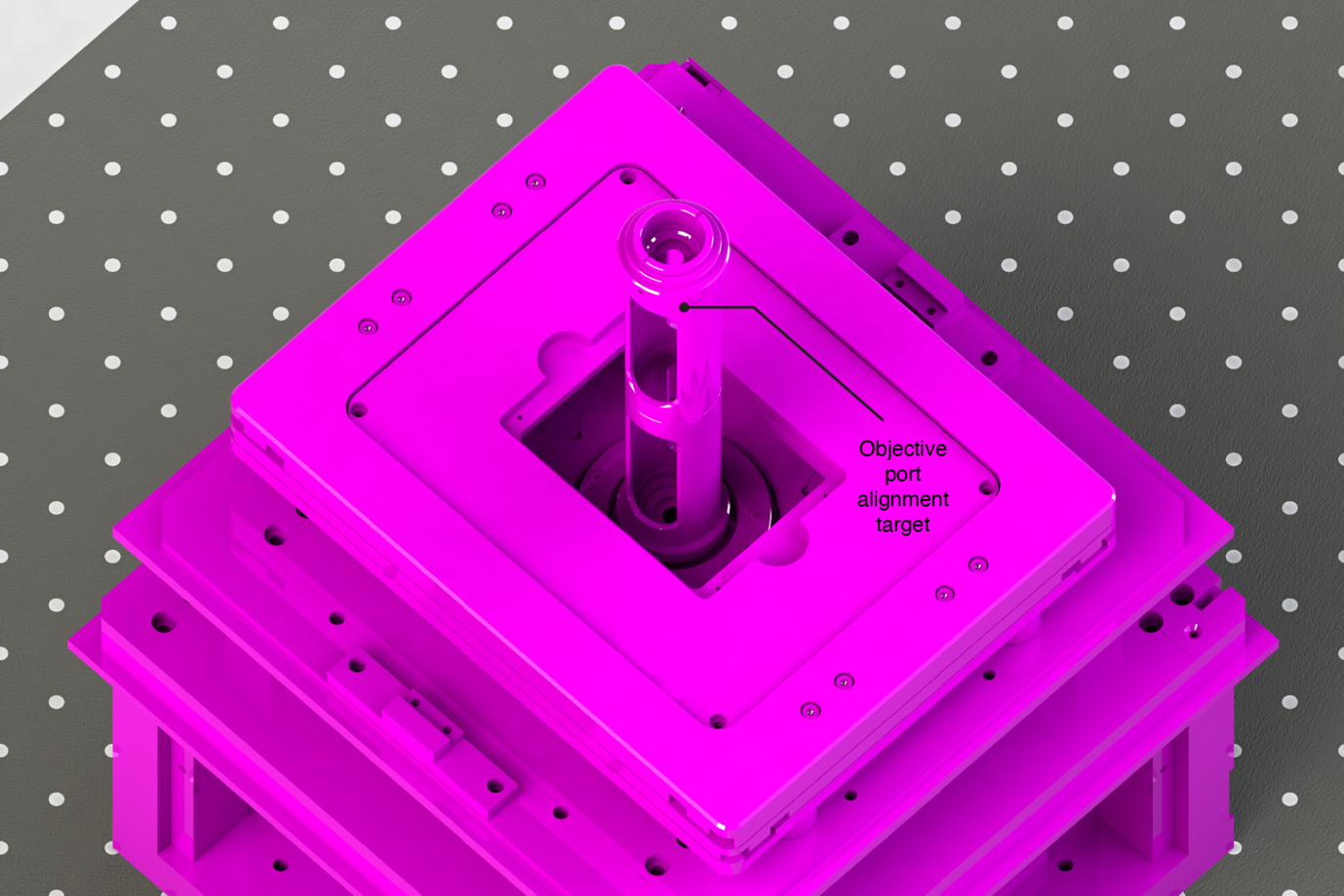

Supplementary Figure 35. Installation of the objective port alignment target at the objective port. The target comprises two spaced irises to ensure co-axial alignment of the illumination lasers with the objective lens axis. The target features a c-mount on the distal (upper) end, allowing it to be coupled with a beam profiling camera (not shown), which greatly aids alignment precision.

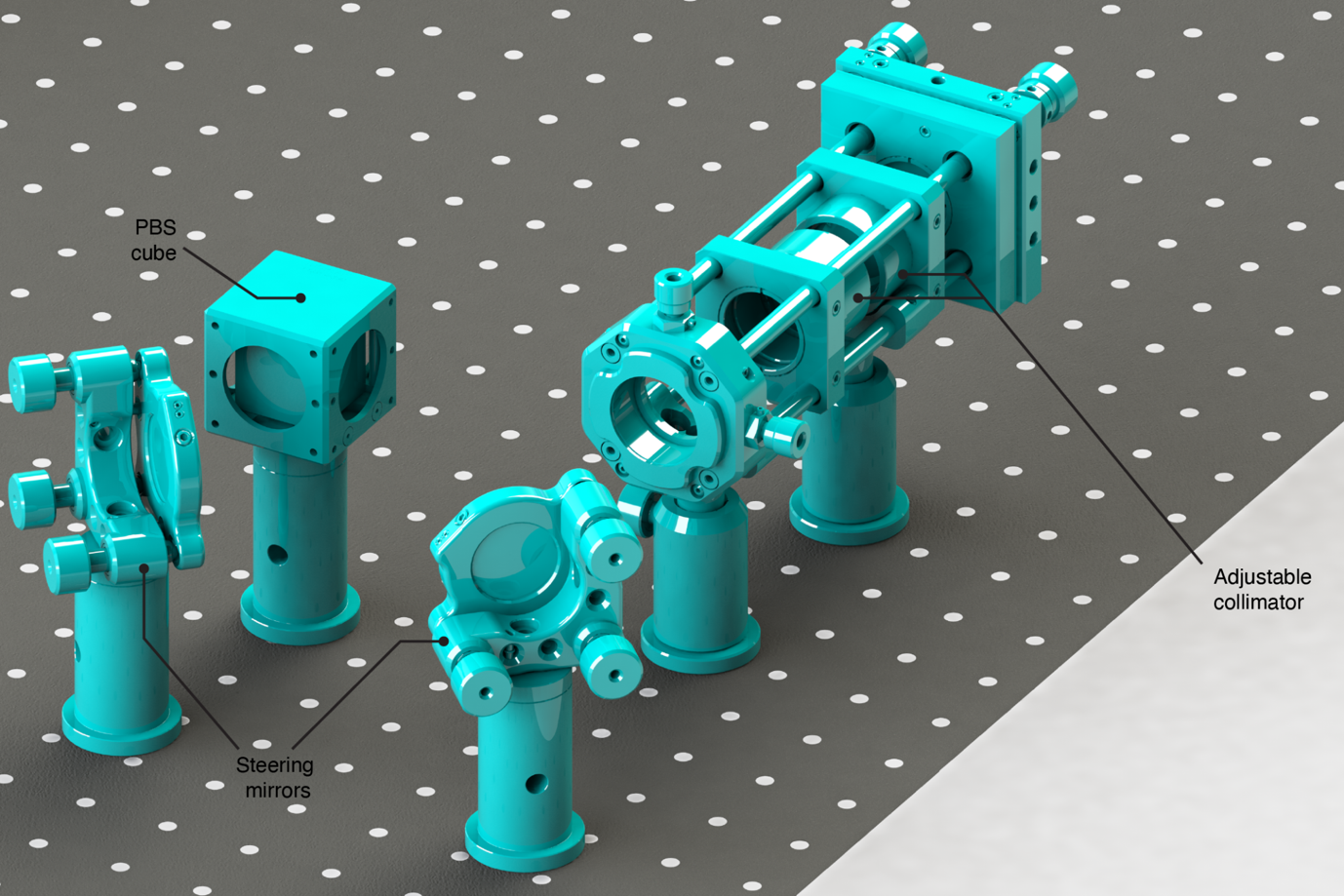

Supplementary Figure 36. Placement and installation of the first steering mirror pair in the single-mode laser engine illumination path. The completed single-mode laser engine beam launch is shown with the pair of achromatic doublets comprising the adjustable collimator. The polarization beam splitter (PBS) cube is included for configurations including a booster laser and must be installed prior to alignment of the booster laser to correctly align and set the polarization of the single-mode laser engine path.

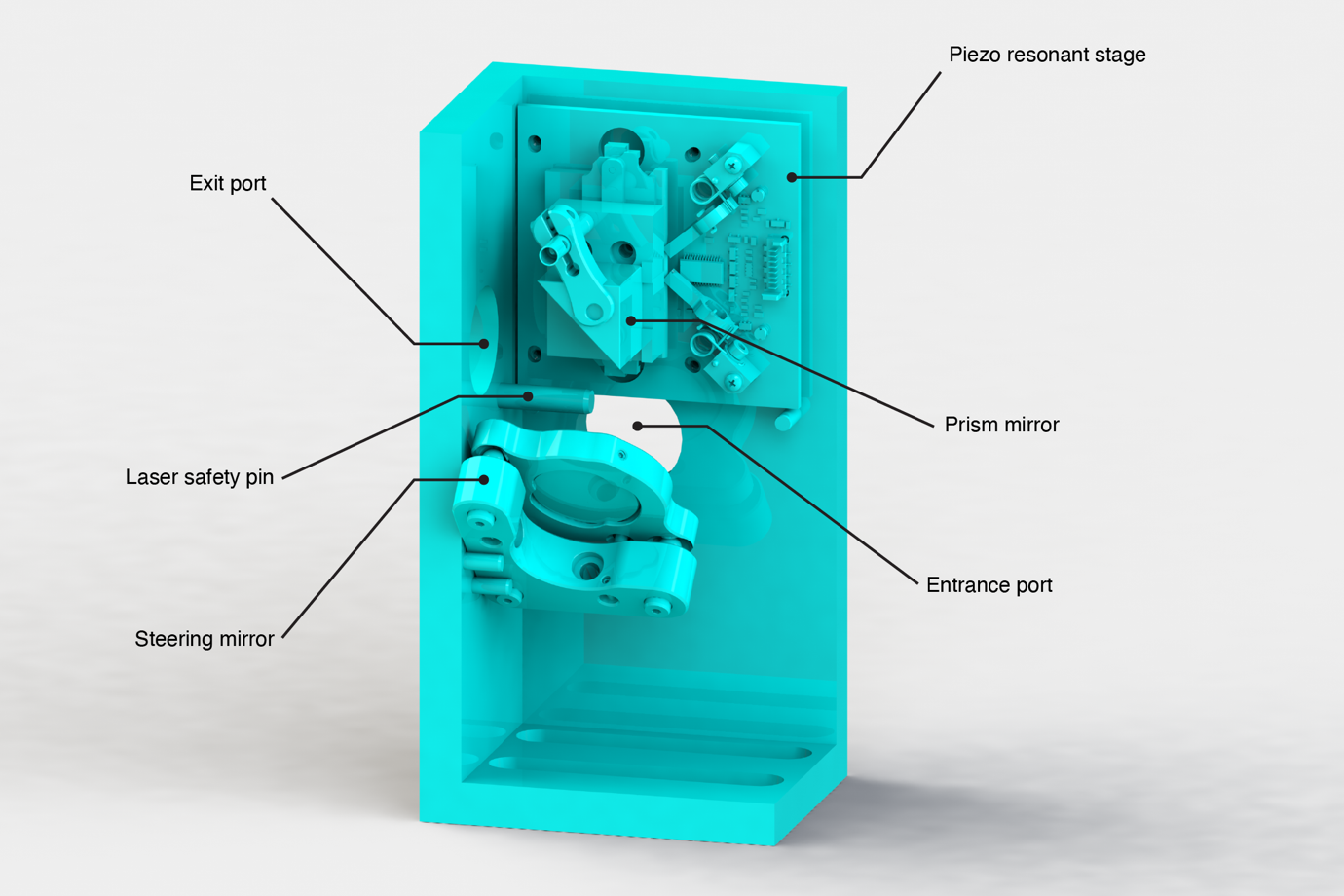

Supplementary Figure 37. The assembled illumination inclination control system, which allows switching between epi-/HILO/TIRF illumination modes and switching between single- and multi-mode laser sources. The laser safety pin limits the motion of the prism mirror to prevent launching a heavily inclined illumination beam out of the proximal side of the optical table.

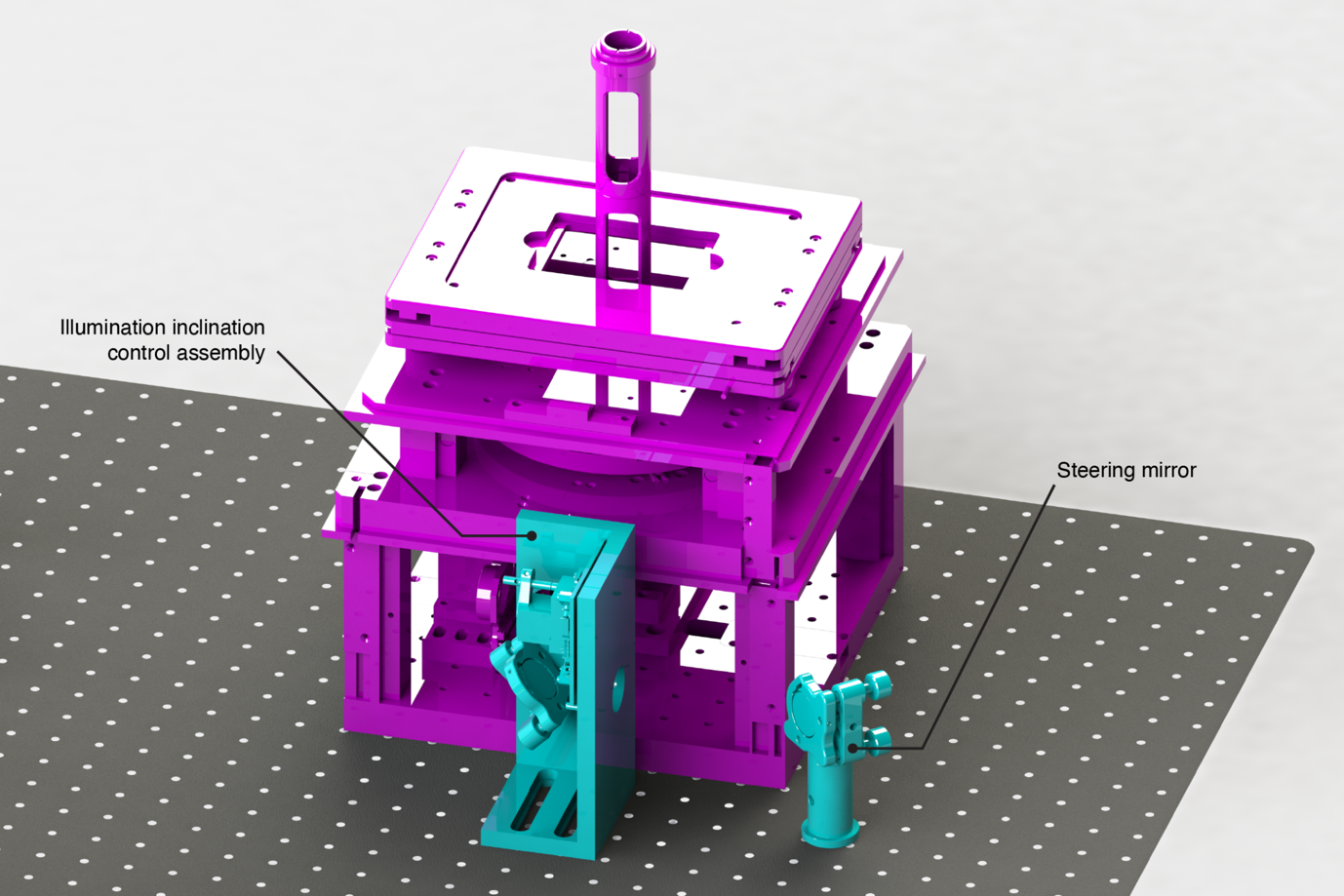

Supplementary Figure 38. Placement and installation of the illumination inclination control system and upstream steering mirror.

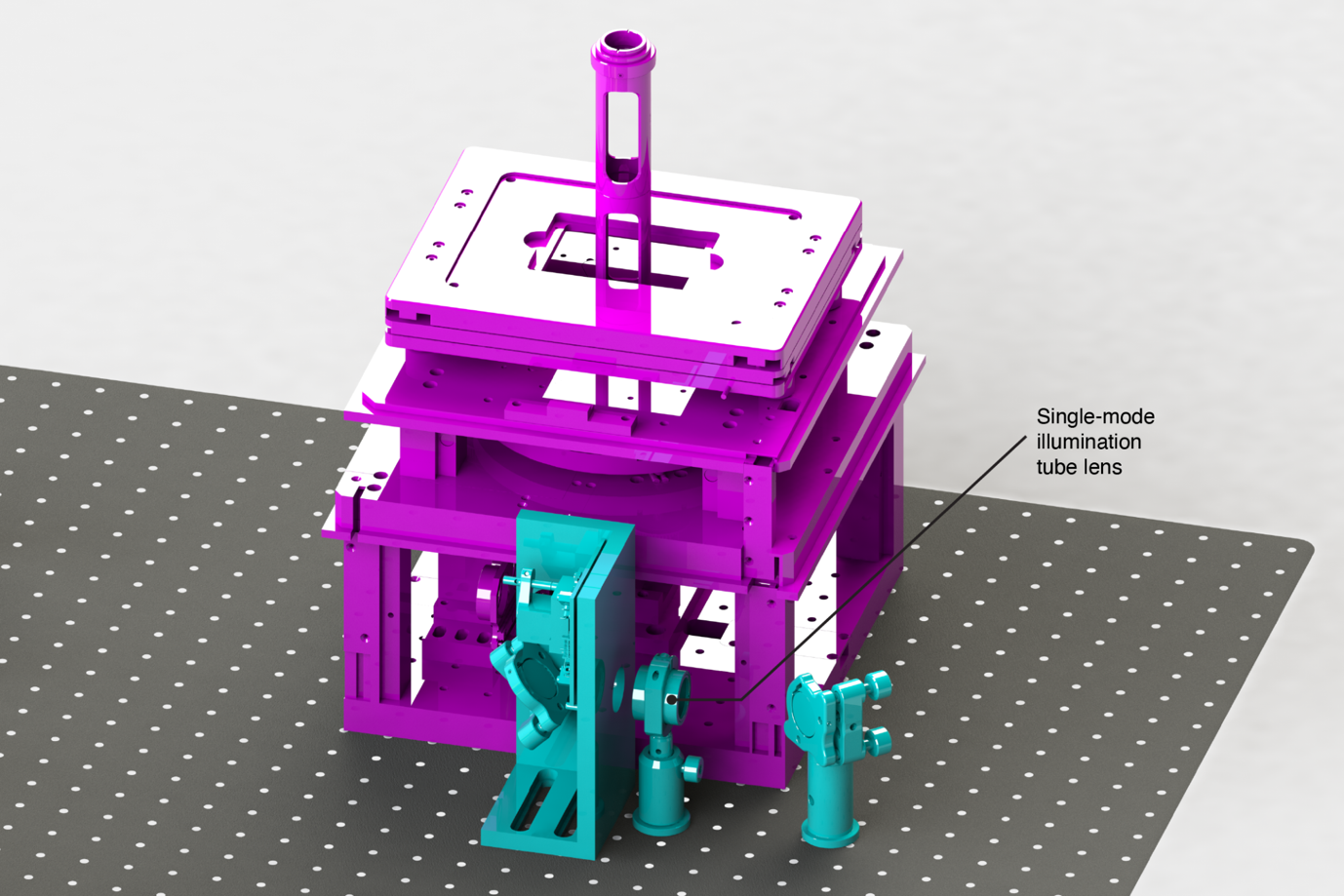

Supplementary Figure 39. Alignment and installation of the tube lens for the single-mode illumination path.

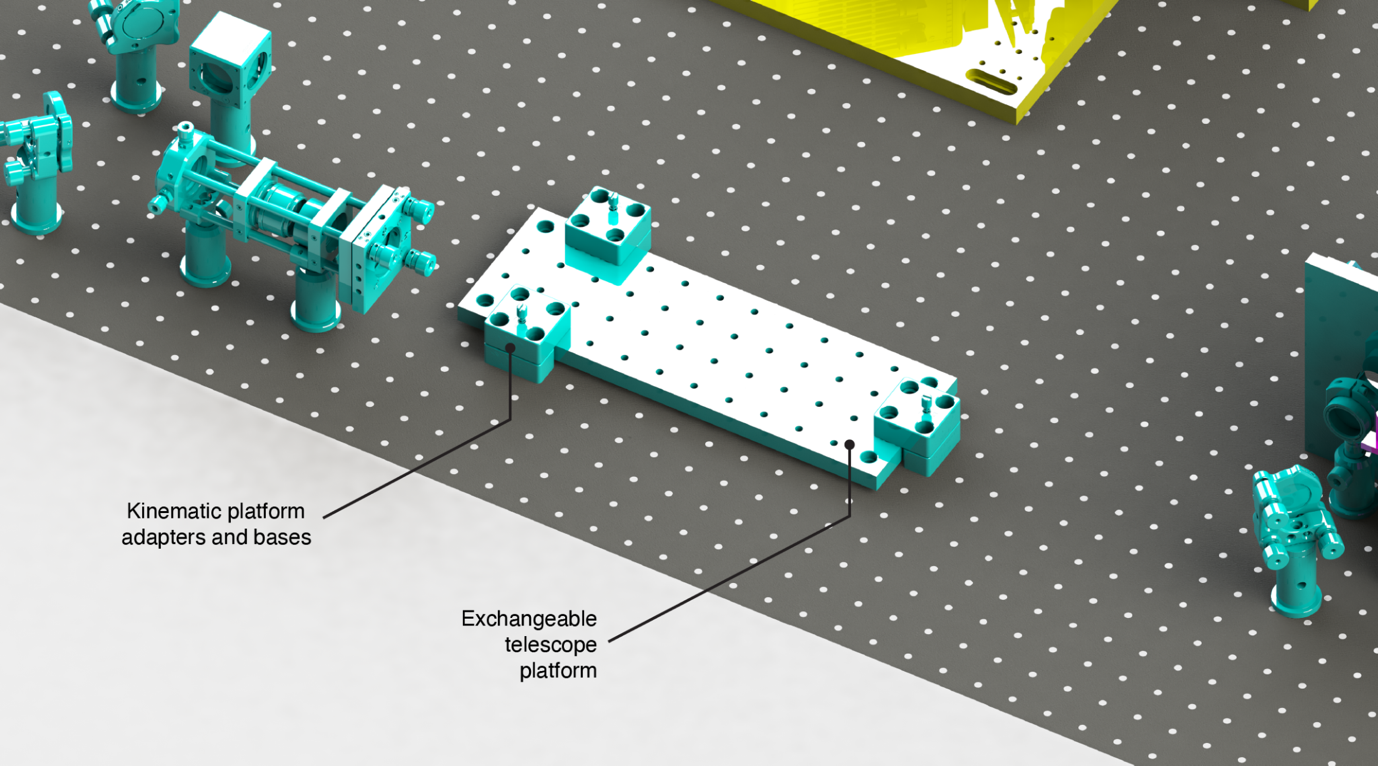

Supplementary Figure 40. Installation of an exchangeable platform, which hosts one of four telescopes used to control the size of the illumination field. The platforms can be exchanged without misalignment via the use of lockable kinematic mounts, which ensure repeatable positioning.

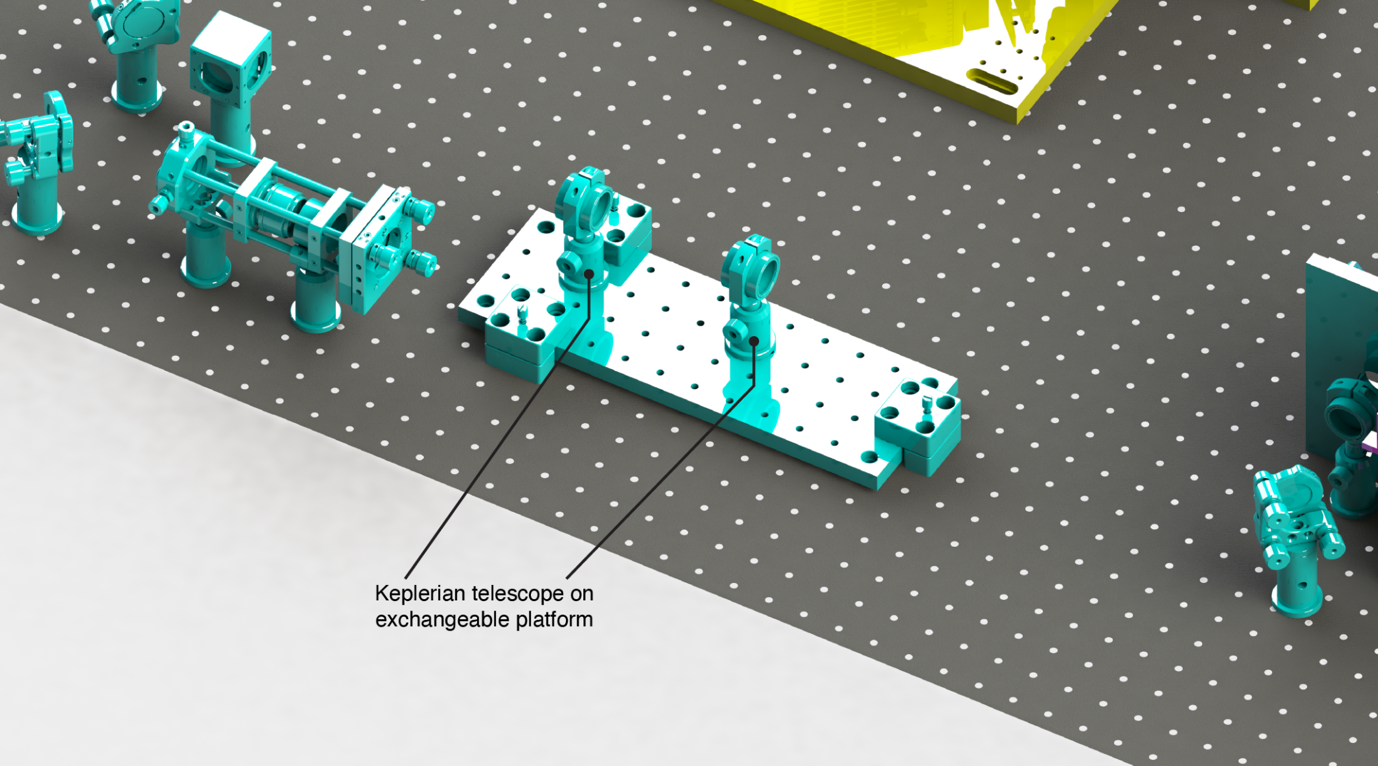

Supplementary Figure 41. One telescope (Keplerian type) installed on an exchangeable platform. In the case shown, two f = 60 mm achromatic doublets are installed, yielding a magnification of 1× and 4f conjugation of their entrance and exit pupil respectively

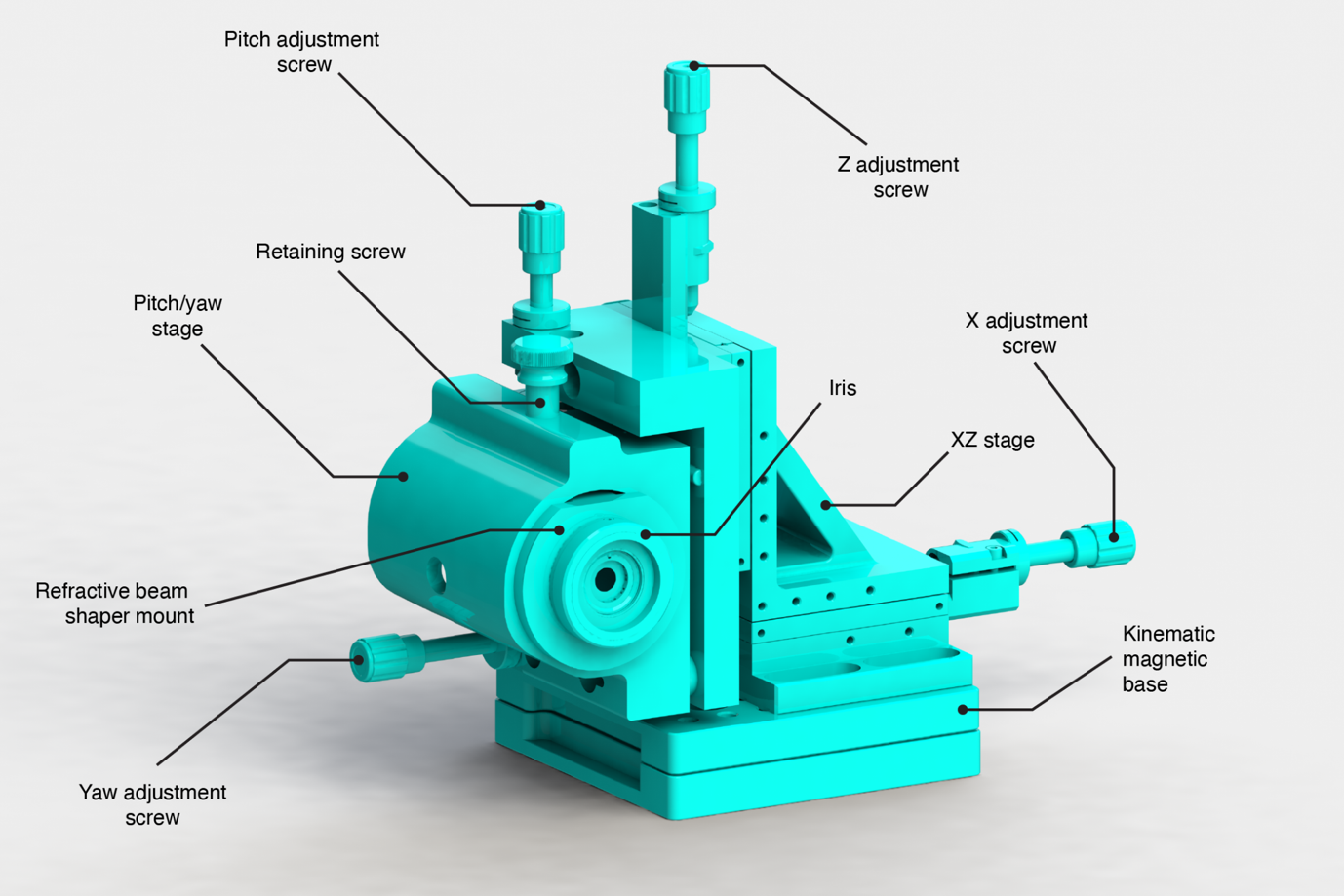

Supplementary Figure 42. The assembled refractive beam shaper mount including XZ, and pitch/yaw stages. The beam shaper mount is retained by a screw of the pitch/yaw stage.

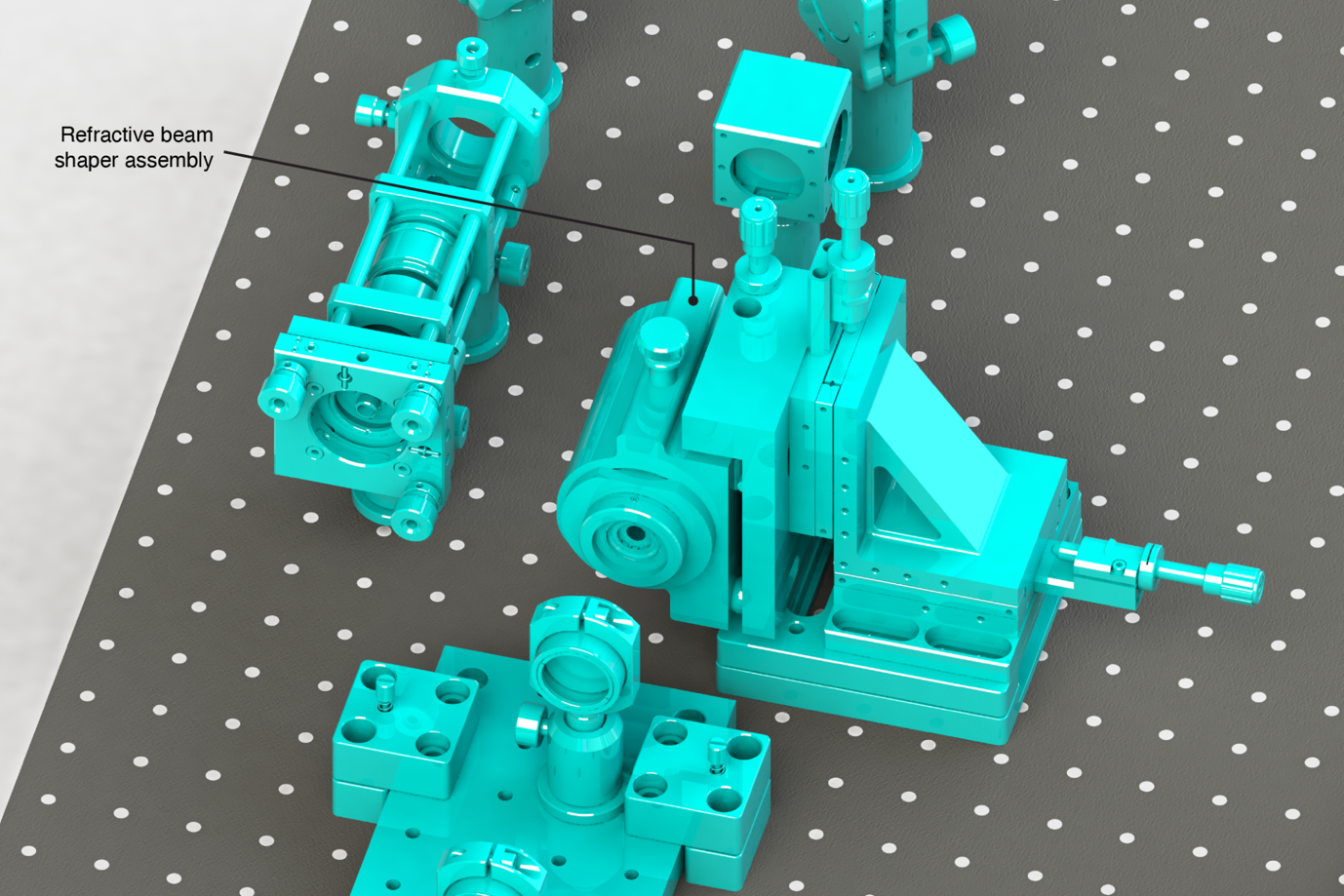

Supplementary Figure 43. Installation of the refractive beam shaper mount. An iris at the exit aperture of the beam shaper (omitted) constitutes the illumination system field aperture when correctly spaced with regard to the downstream telescope.

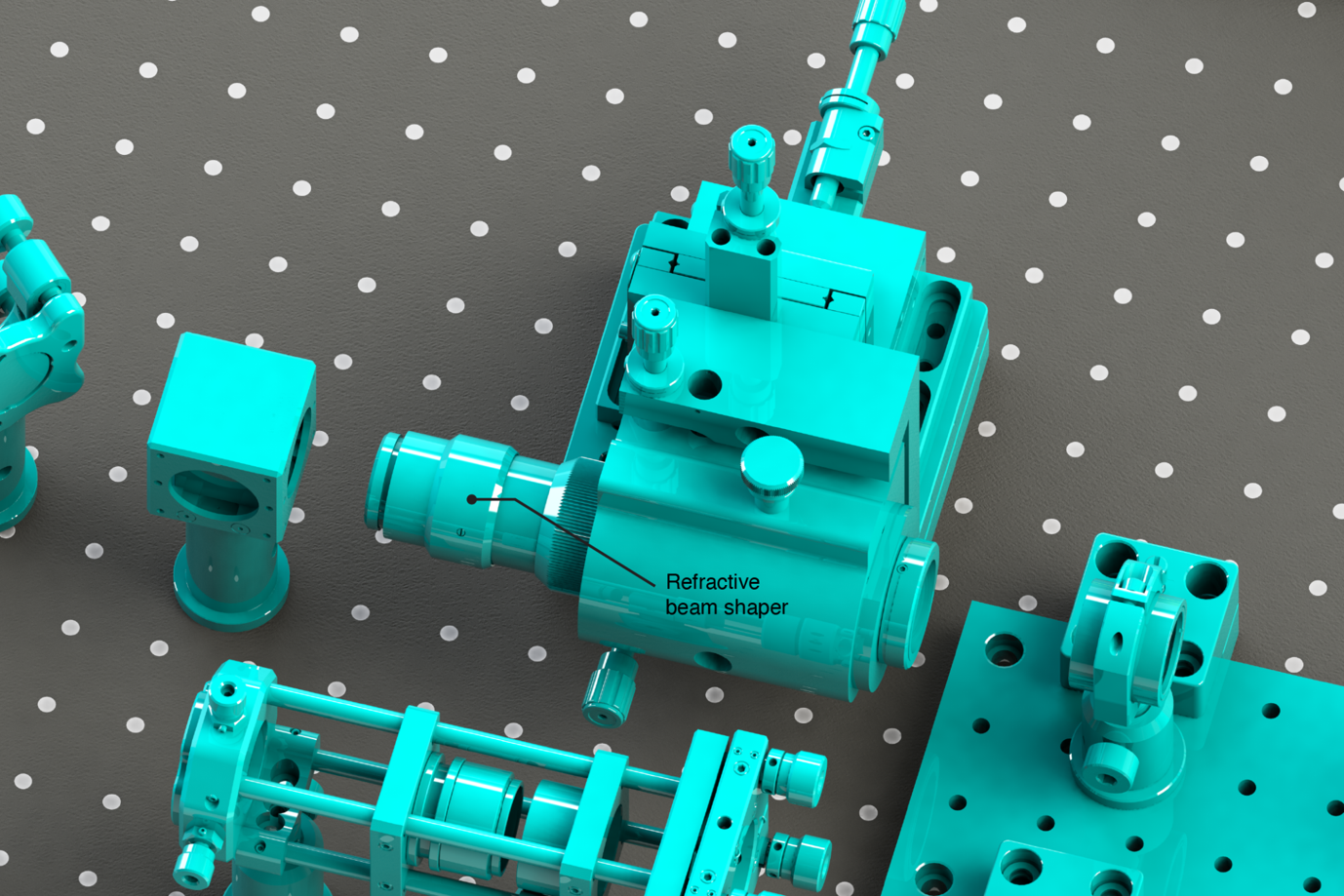

Supplementary Figure 44. Installation of the refractive beam shaper in its mount.

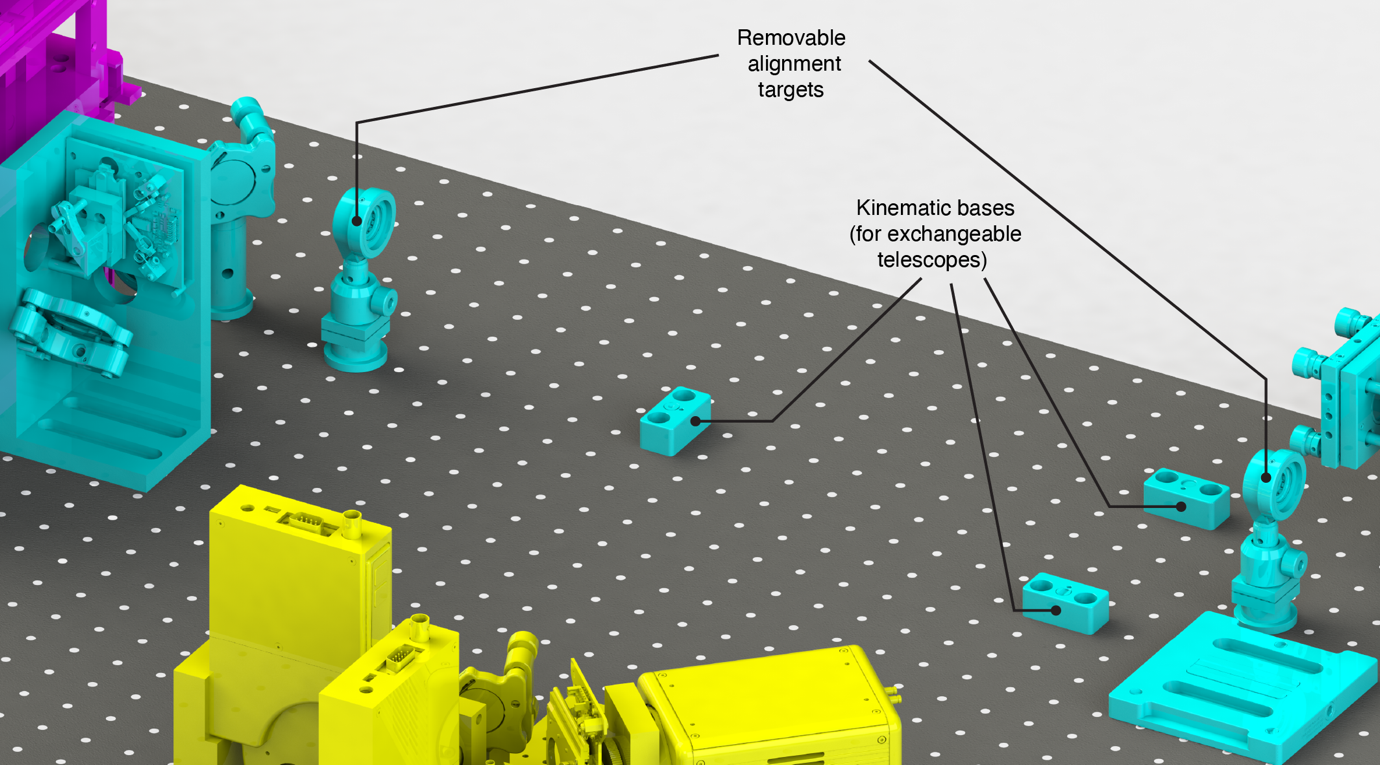

Supplementary Figure 45. Installation of a pair of alignment targets with the upper portion of the refractive beam shaper mount and the platform-mounted telescope removed. The targets aid future realignment of the system by indicating the beam position to which the beam shaper and telescopes have been aligned.

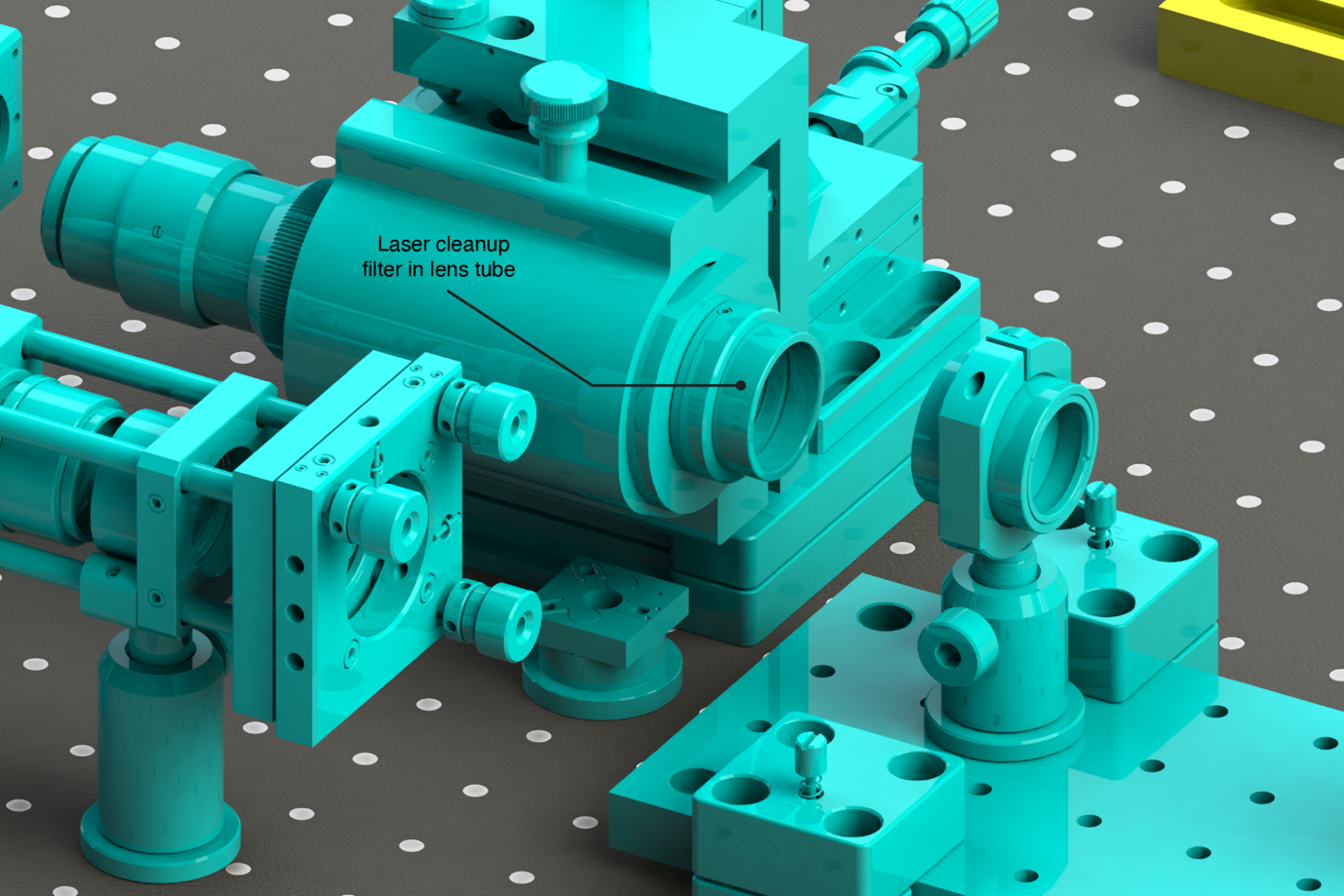

Supplementary Figure 46. Installation of the laser cleanup filter. The cleanup filter is housed in a lens tube, which is threaded into the iris of the refractive beam shaper assembly.

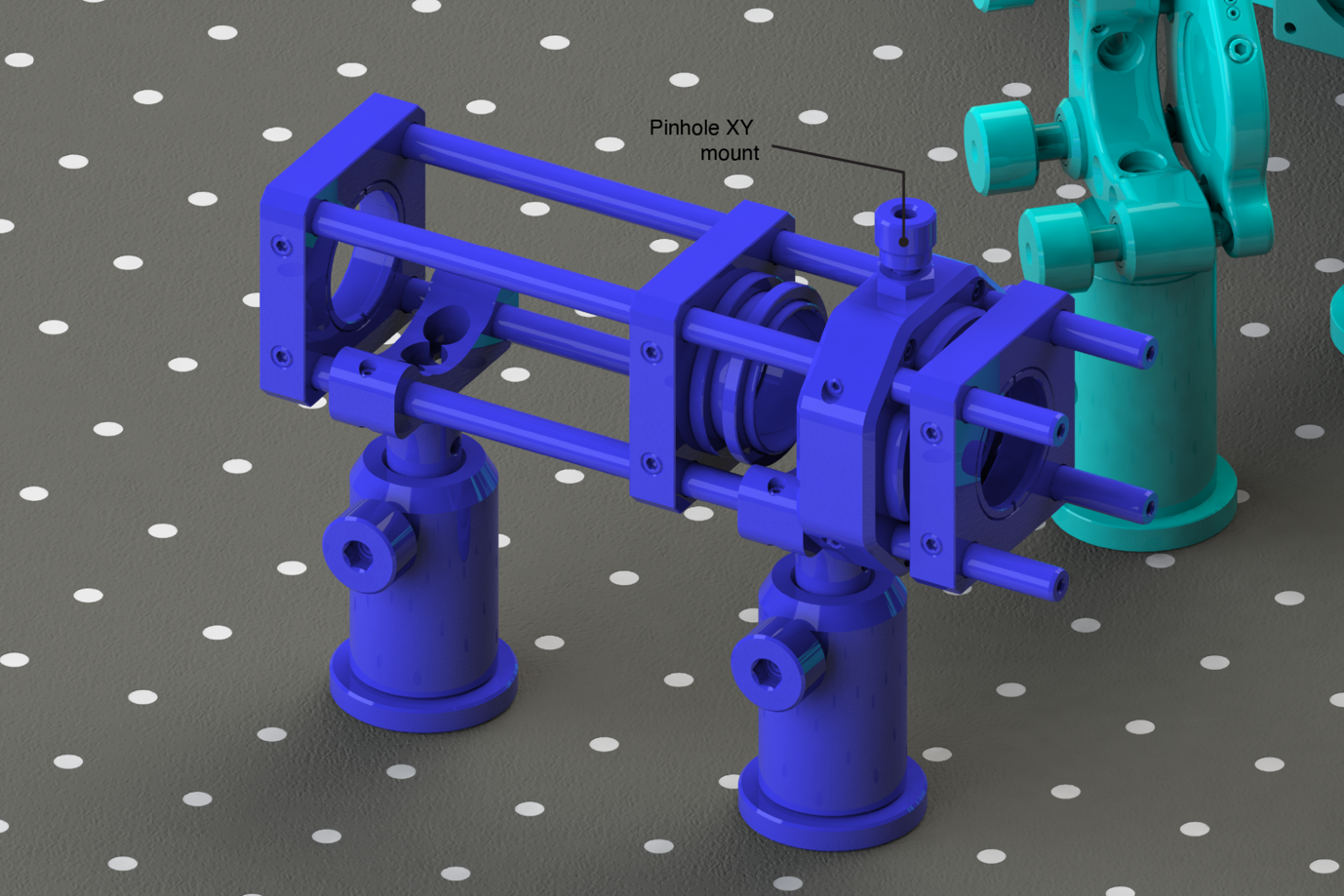

Supplementary Figure 47. The single–mode booster laser spatial filter with the pinhole and lenses omitted.

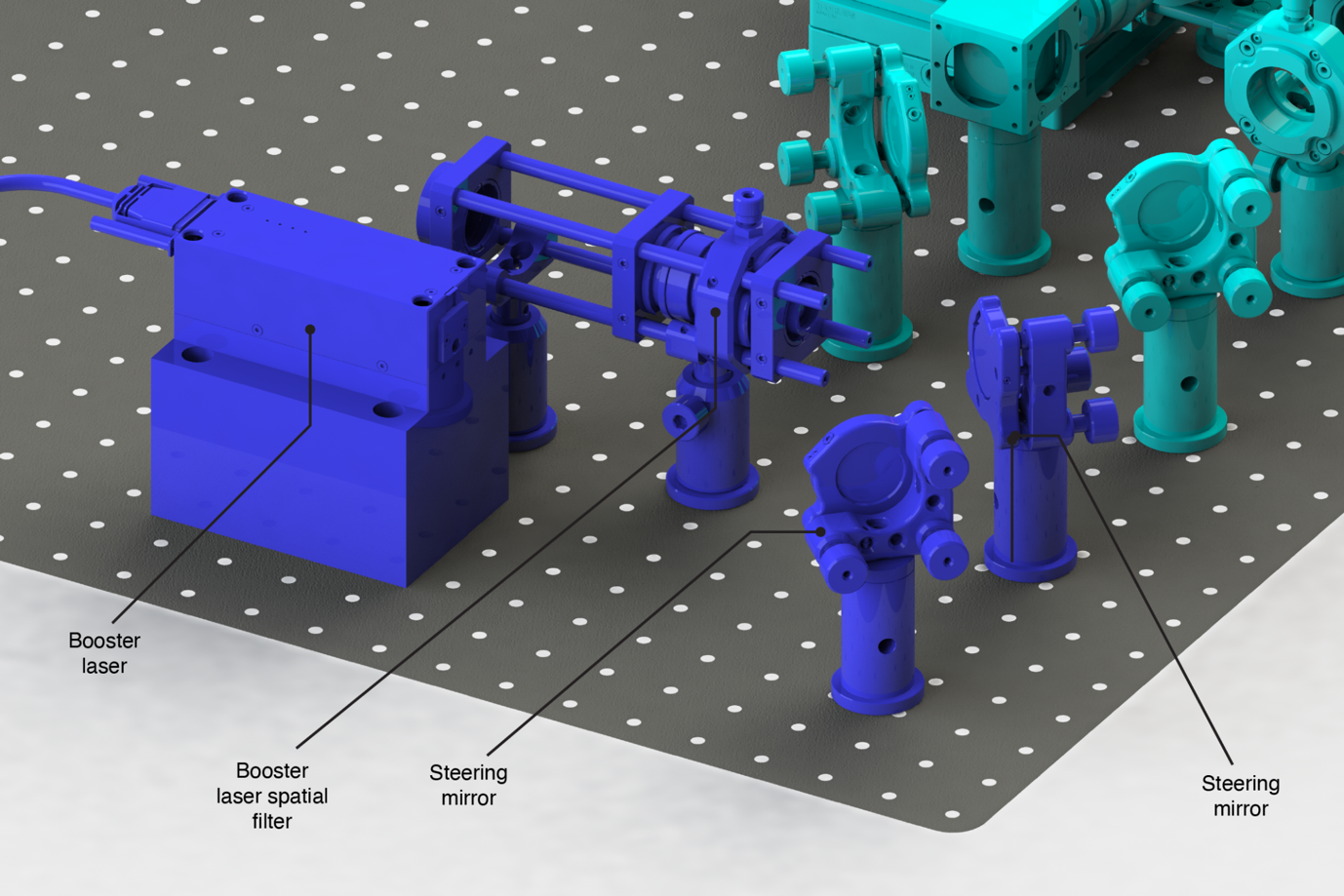

Supplementary Figure 48. The single–mode booster laser and first pair of steering mirrors are introduced and aligned to route the beam through the cage system of the spatial filter on-axis. The lenses and pinhole are subsequently installed in the spatial filter to focus the beam through the pinhole and collimate the filtered output to the required beam diameter.

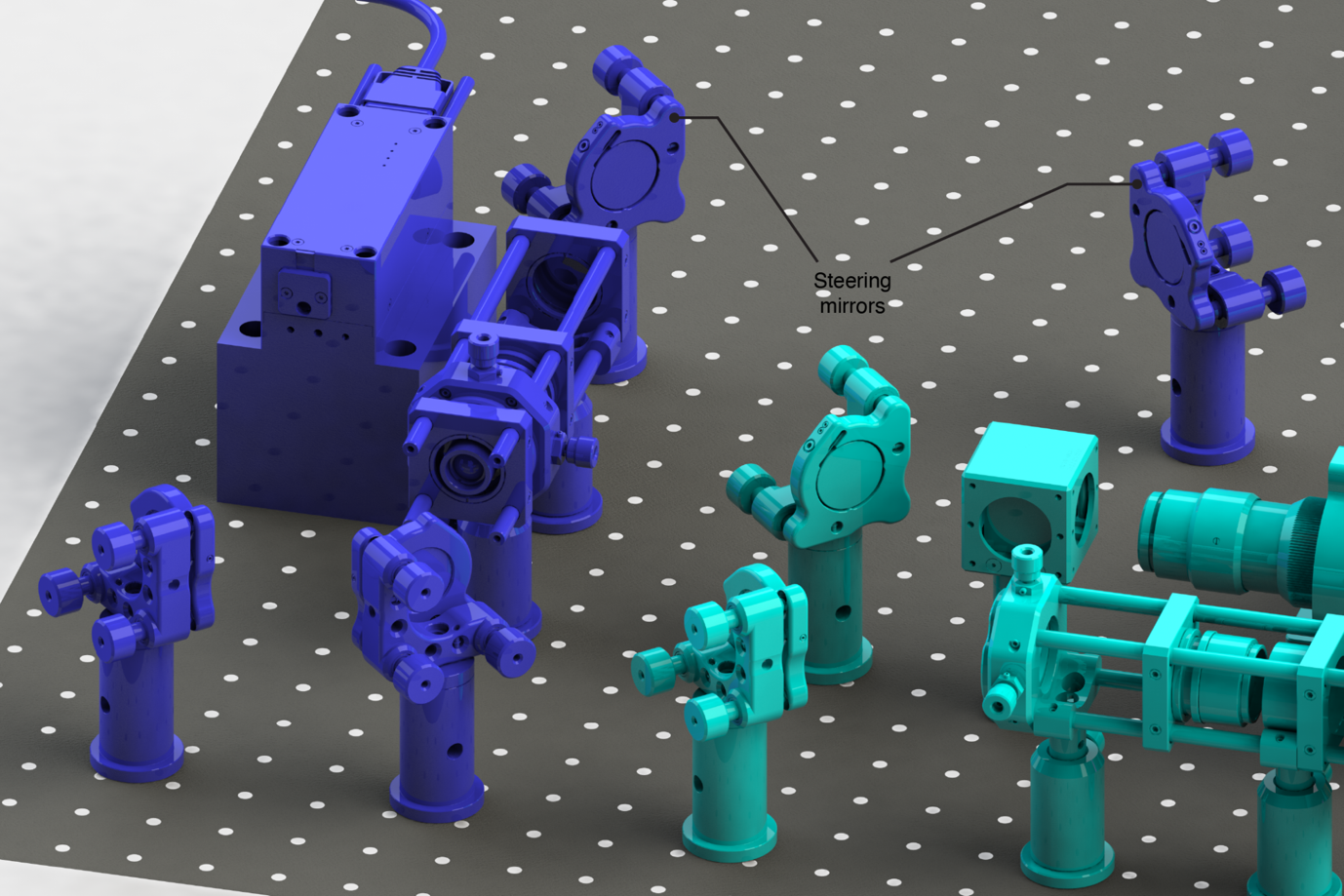

Supplementary Figure 49. The single-mode booster laser is steered via a pair of steering mirrors and combined with the single-mode laser engine at a polarization beam splitter cube (see also Supplementary Figure 36).

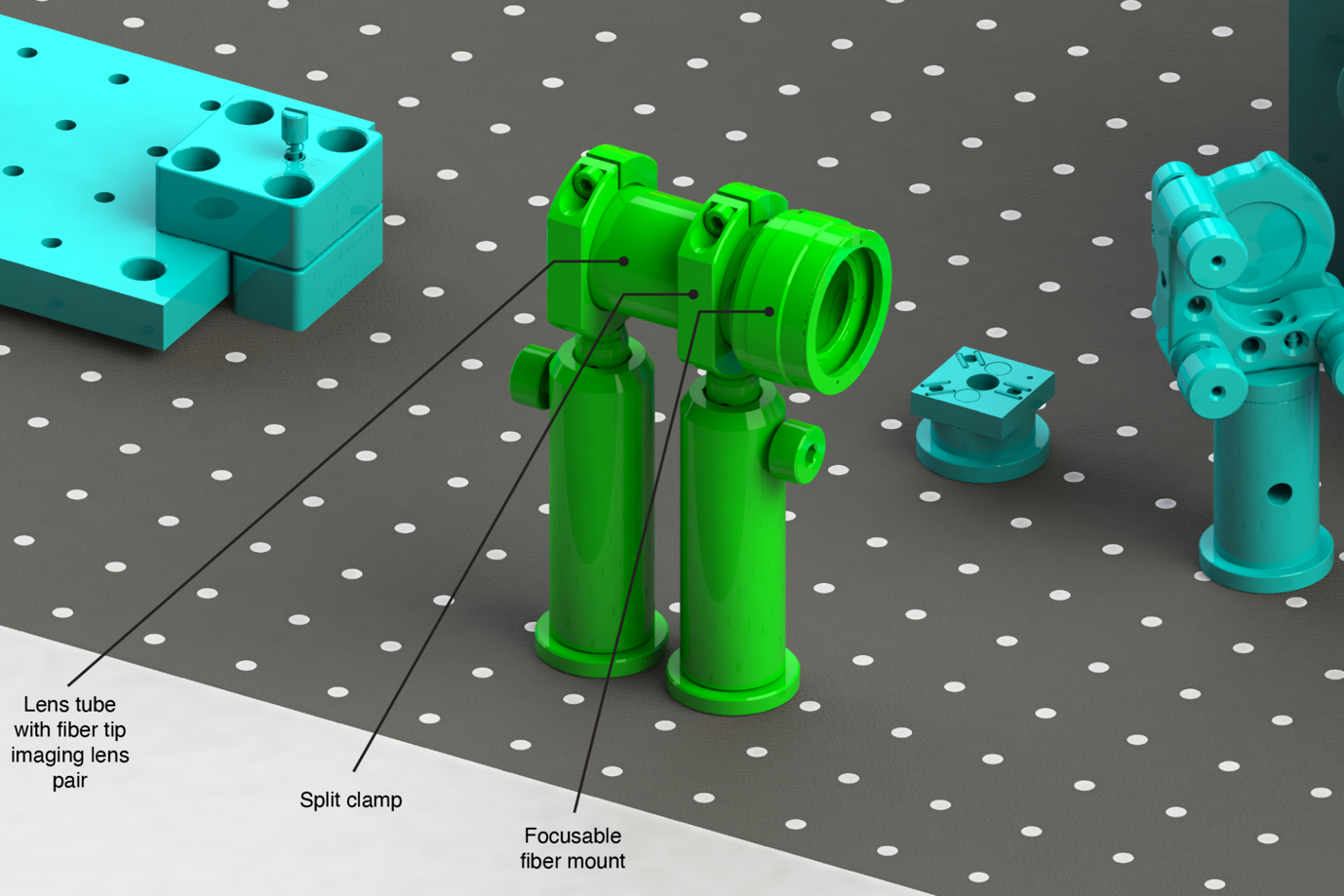

Supplementary Figure 50. The multi-mode laser engine beam launch comprises a focusable fiber mount and pair of achromatic doublets, which allows precise focusing of the fiber tip image to a conjugate object plane later in the protocol. The split clamps allow the lens tube assembly to be rotated to orient the square tip of the fiber correctly with regard to the sCMOS camera chip.

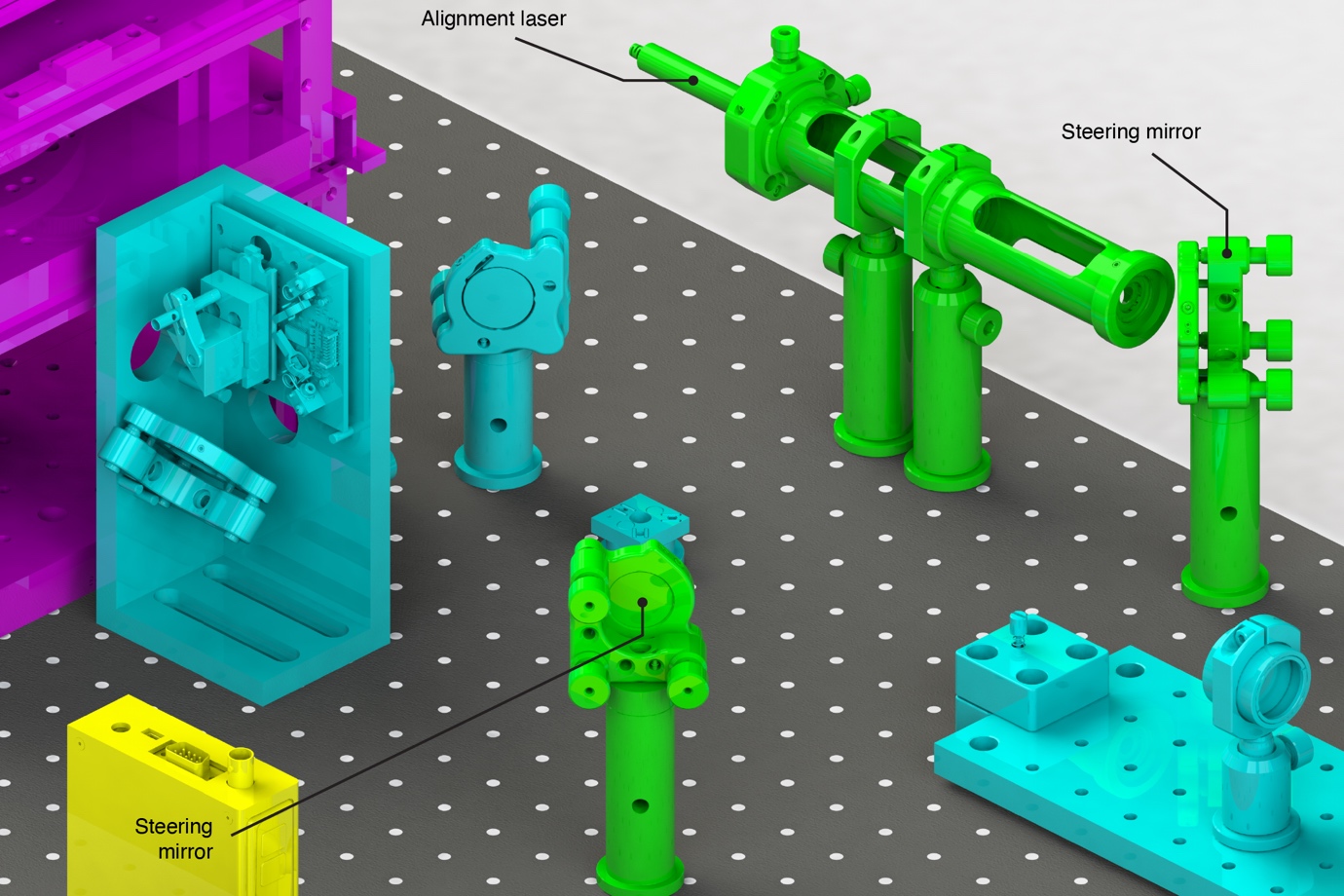

Supplementary Figure 51. The alignment laser is installed through the split clamps of the multi-mode laser engine beam launch and provides a proxy for the multi-mode beam. The two steering mirrors are installed to route the alignment laser on-axis out of the objective port.

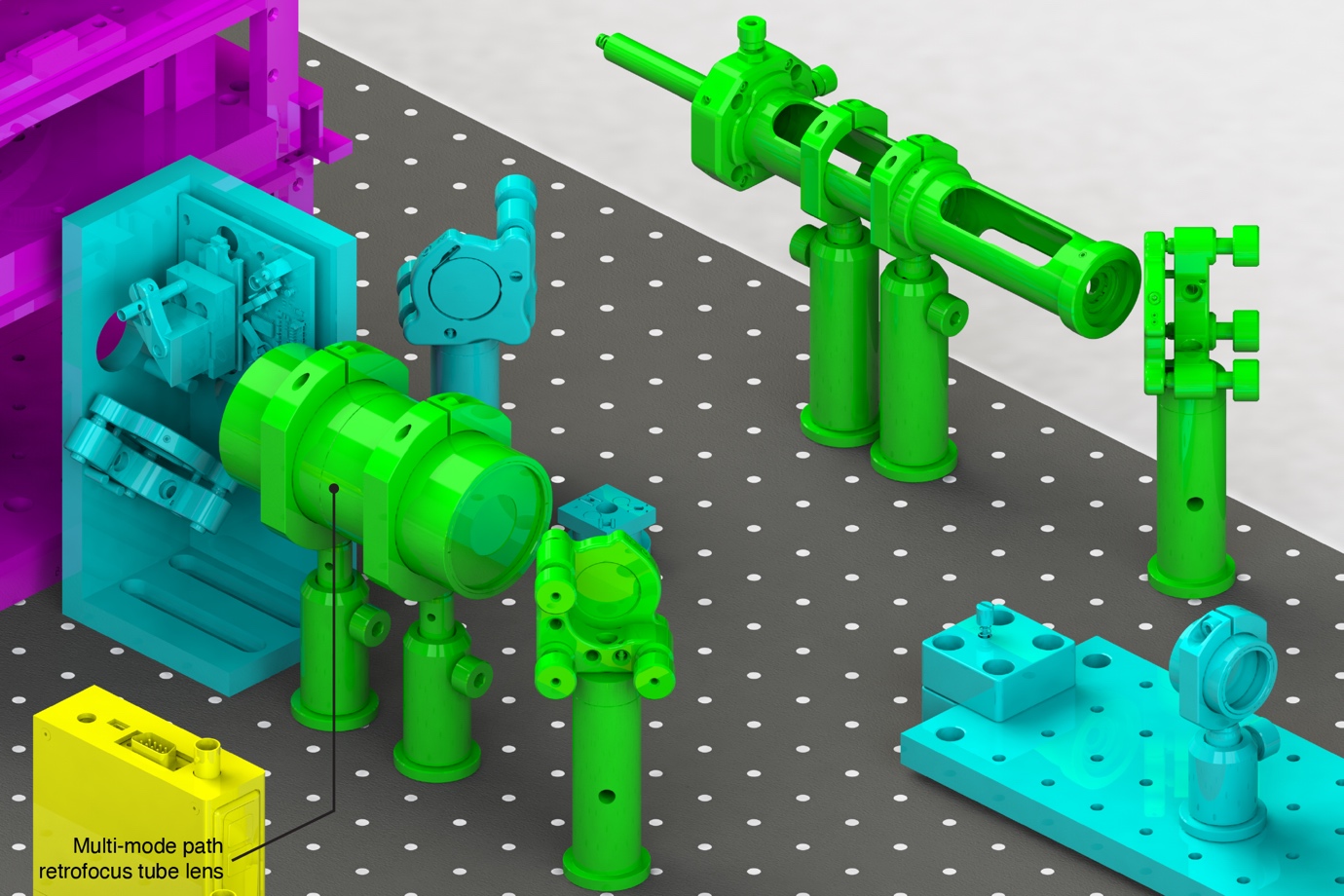

Supplementary Figure 52. Installation and alignment of the multi-mode laser retrofocus tube lens. The lens is focused to produce a collimated output of the alignment laser from the objective port.

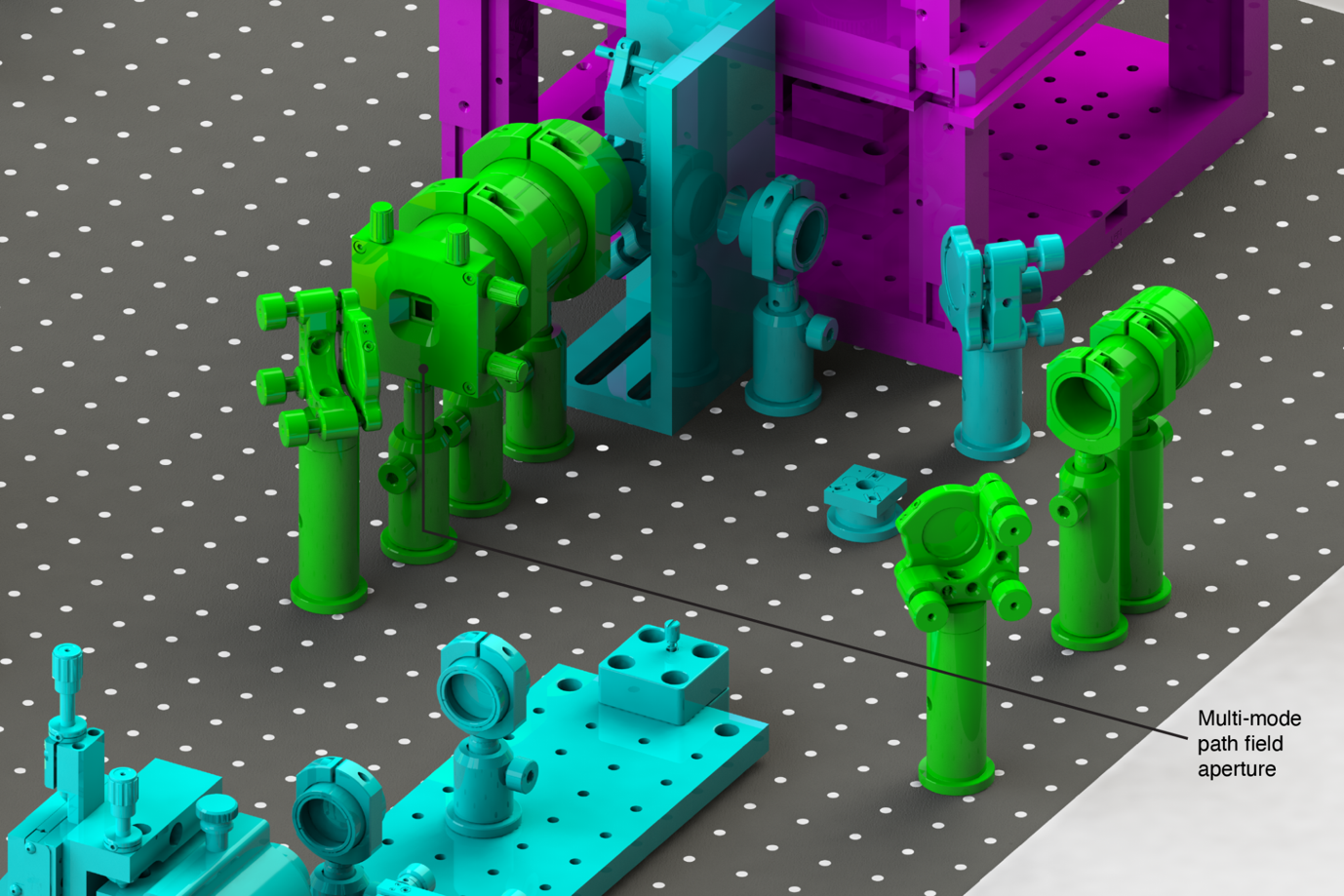

Supplementary Figure 53. Introduction and installation of the multi-mode path field aperture. The aperture is positioned conjugate to the object plane by focusing its image on the infinite conjugate viewing camera installed at the objective port.

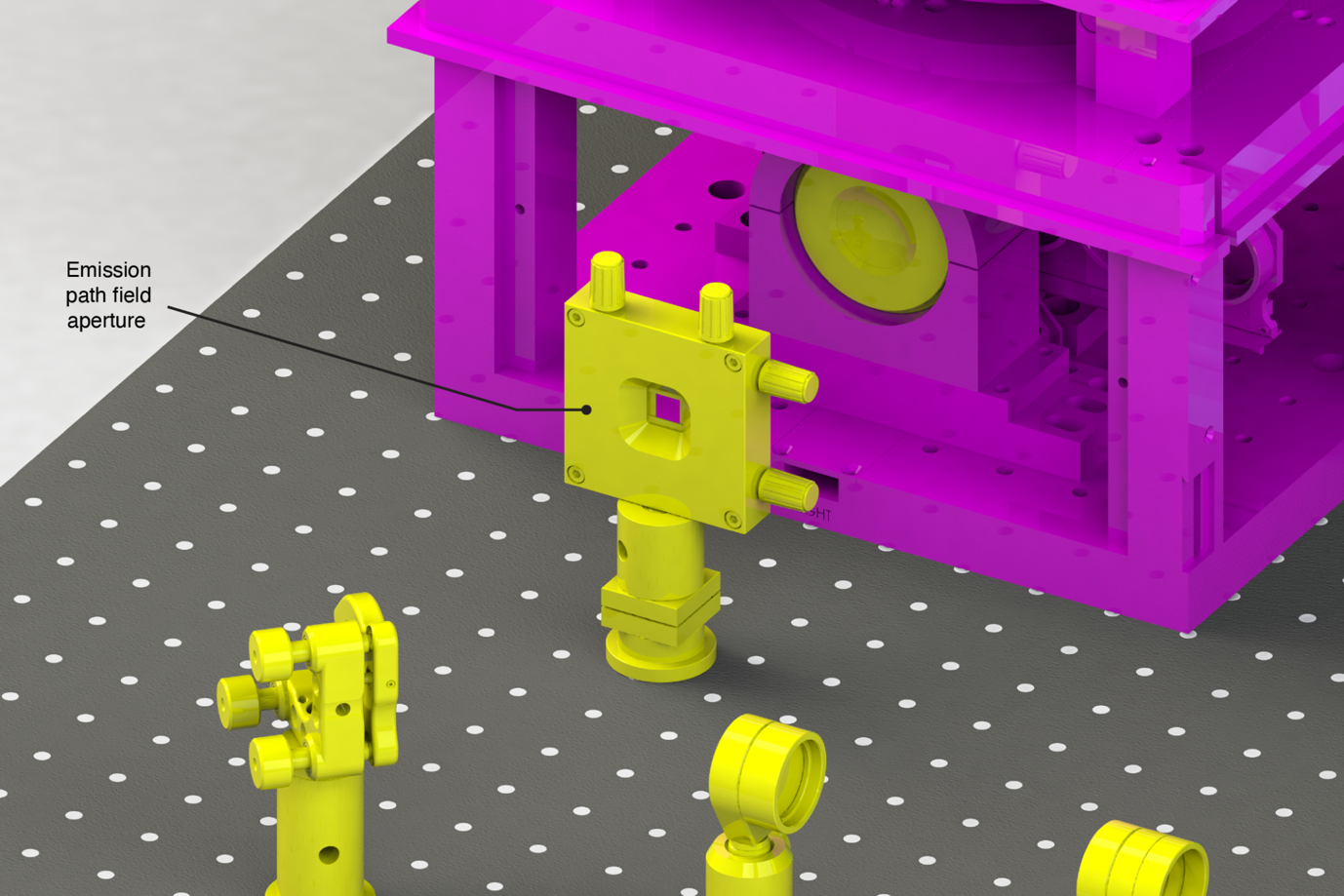

Supplementary Figure 54. Introduction and installation of the emission path field aperture. The aperture is positioned at the primary image plane by focusing its image on the sCMOS camera.

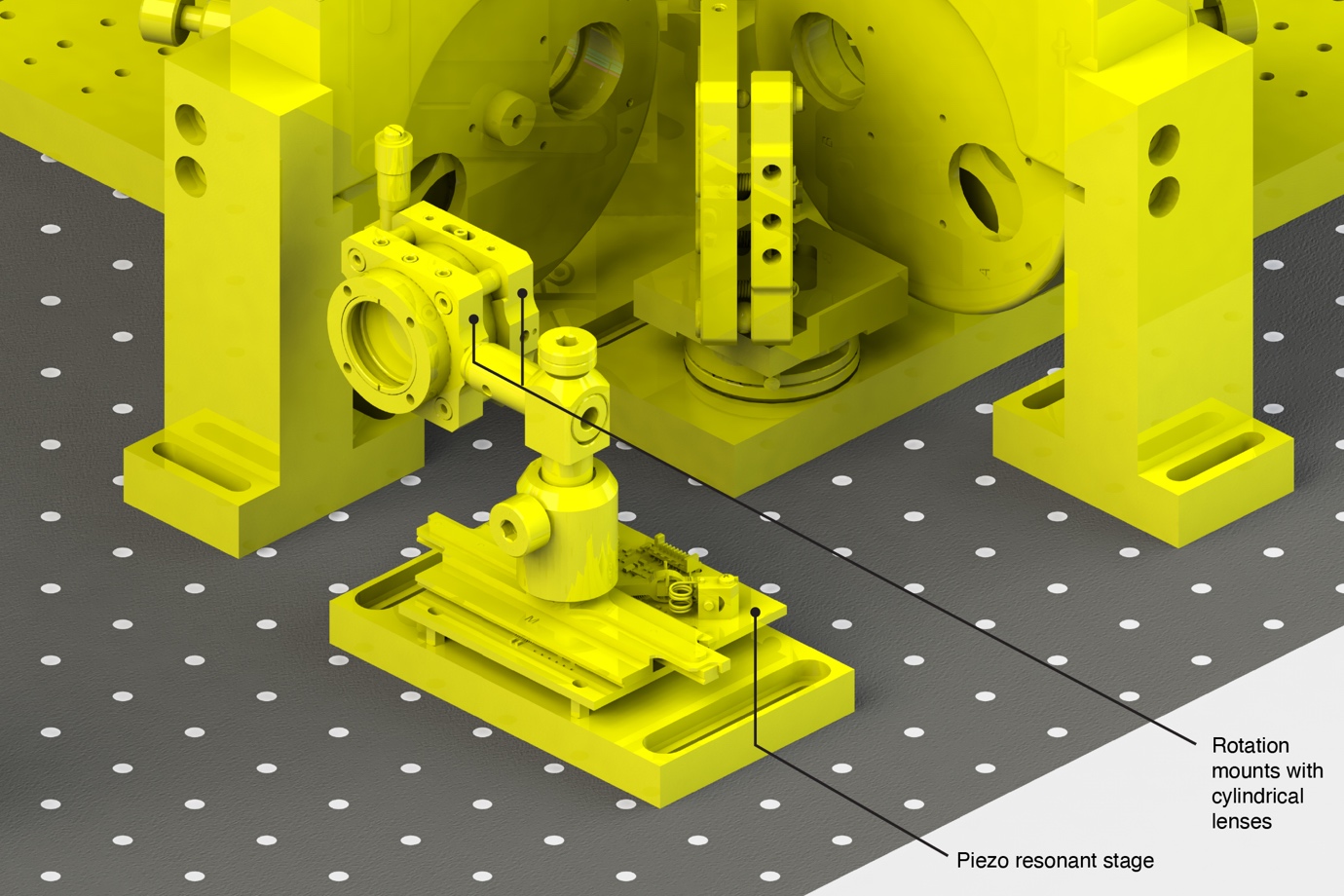

Supplementary Figure 55. Introduction and installation of the astigmatic lens for 3D imaging. The astigmatic lens is longitudinally positioned conjugate to the objective lens back focal plane using the Bertrand lens to image a fixed aperture target placed intermediated between the cylindrical lenses and laterally aligned to the alignment laser installed at the objective port.

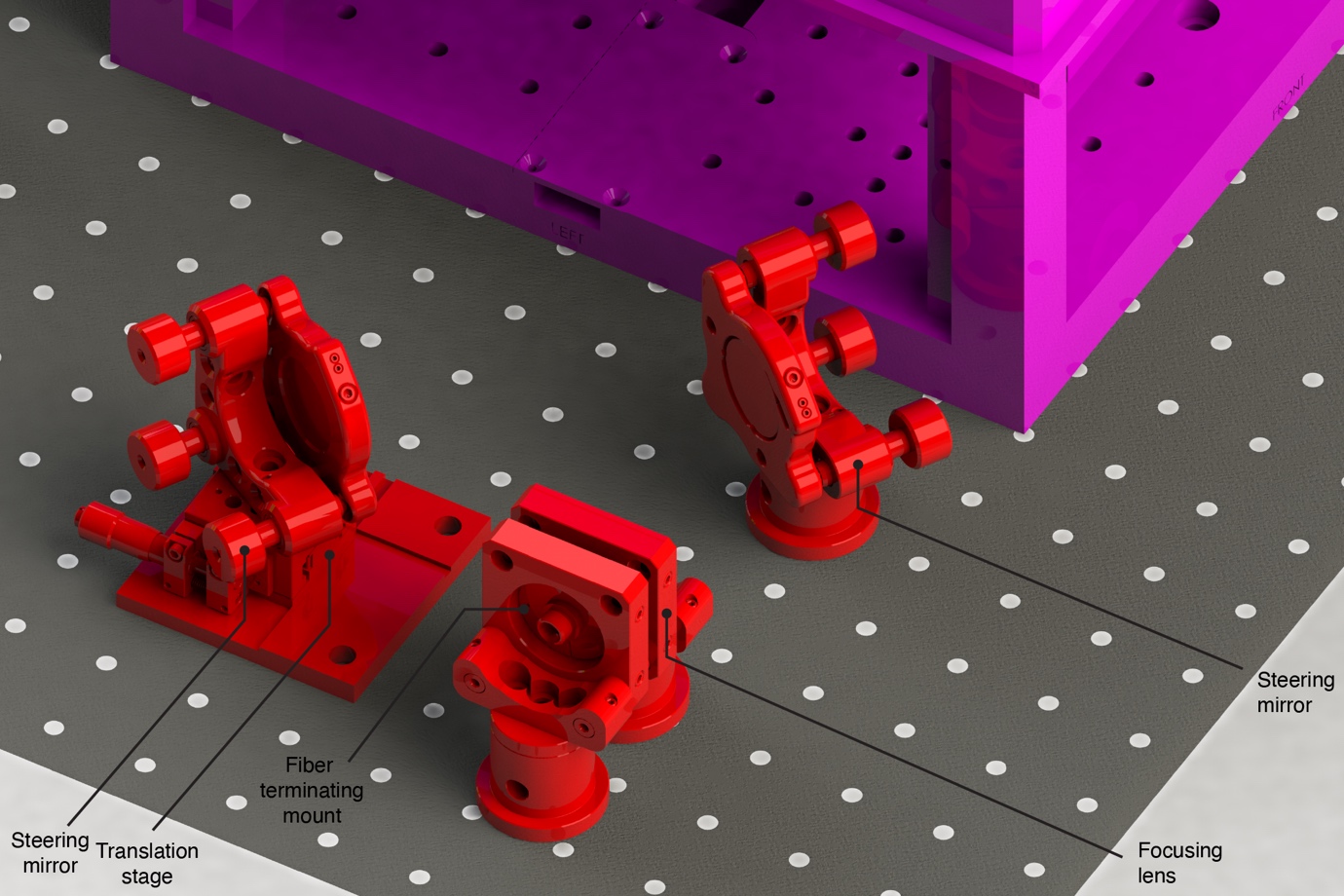

Supplementary Figure 56. Installation of the inward bound portion of the focus lock path. The output of the focus lock laser fiber is steered to the objective port using the two steering mirrors and imaged onto the objective back focal plane by adjusting the position of the focusing lens relative to the fiber terminating mount. The focused beam is offset from the back focal plane center using the translation stage to achieve (total internal) reflection of the beam at the coverslip/sample interface.

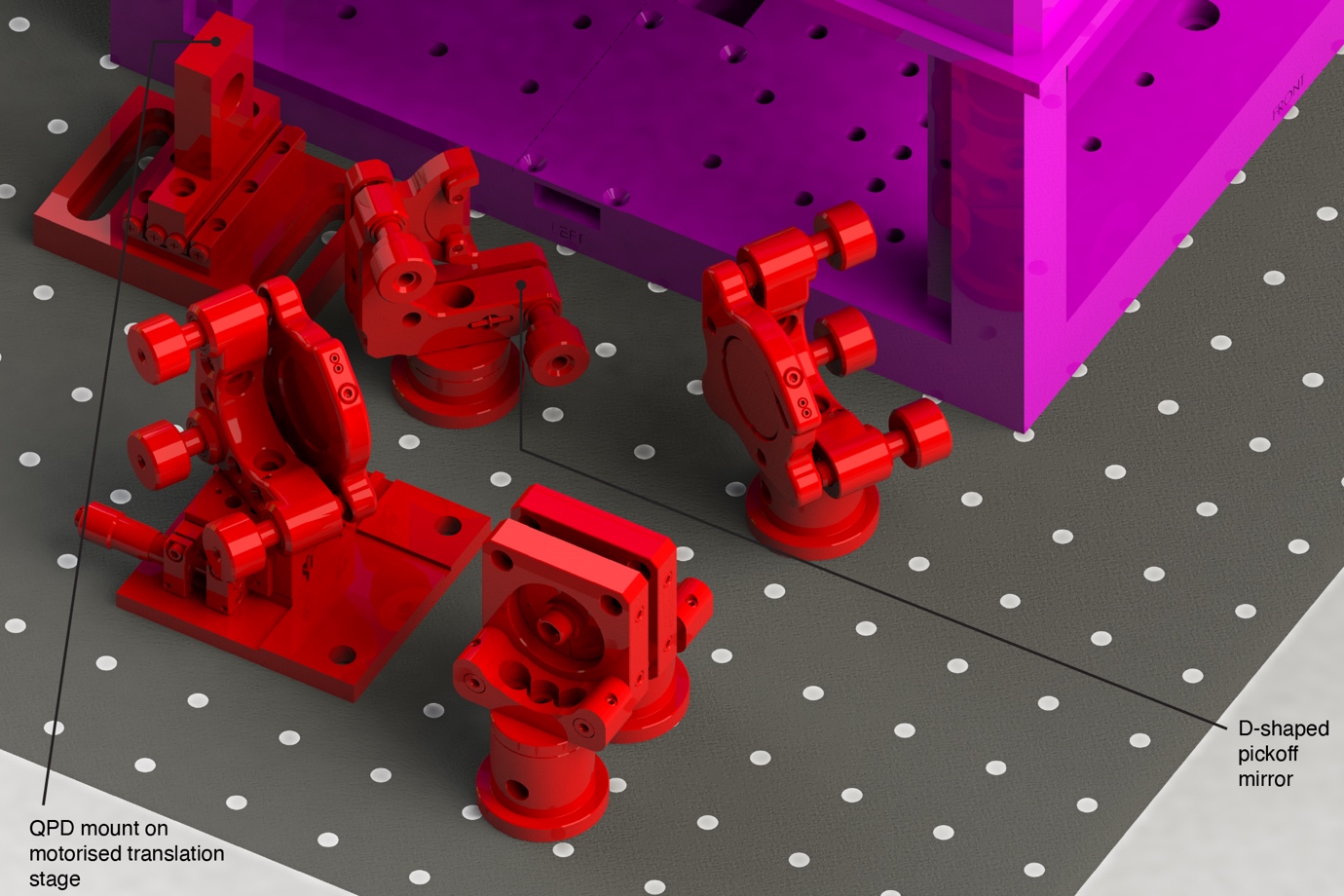

Supplementary Figure 57. Installation of the outward-bound portion of the focus lock path. The returned beam is picked off with a D-shaped mirror onto a quadrant photodiode (QPD).

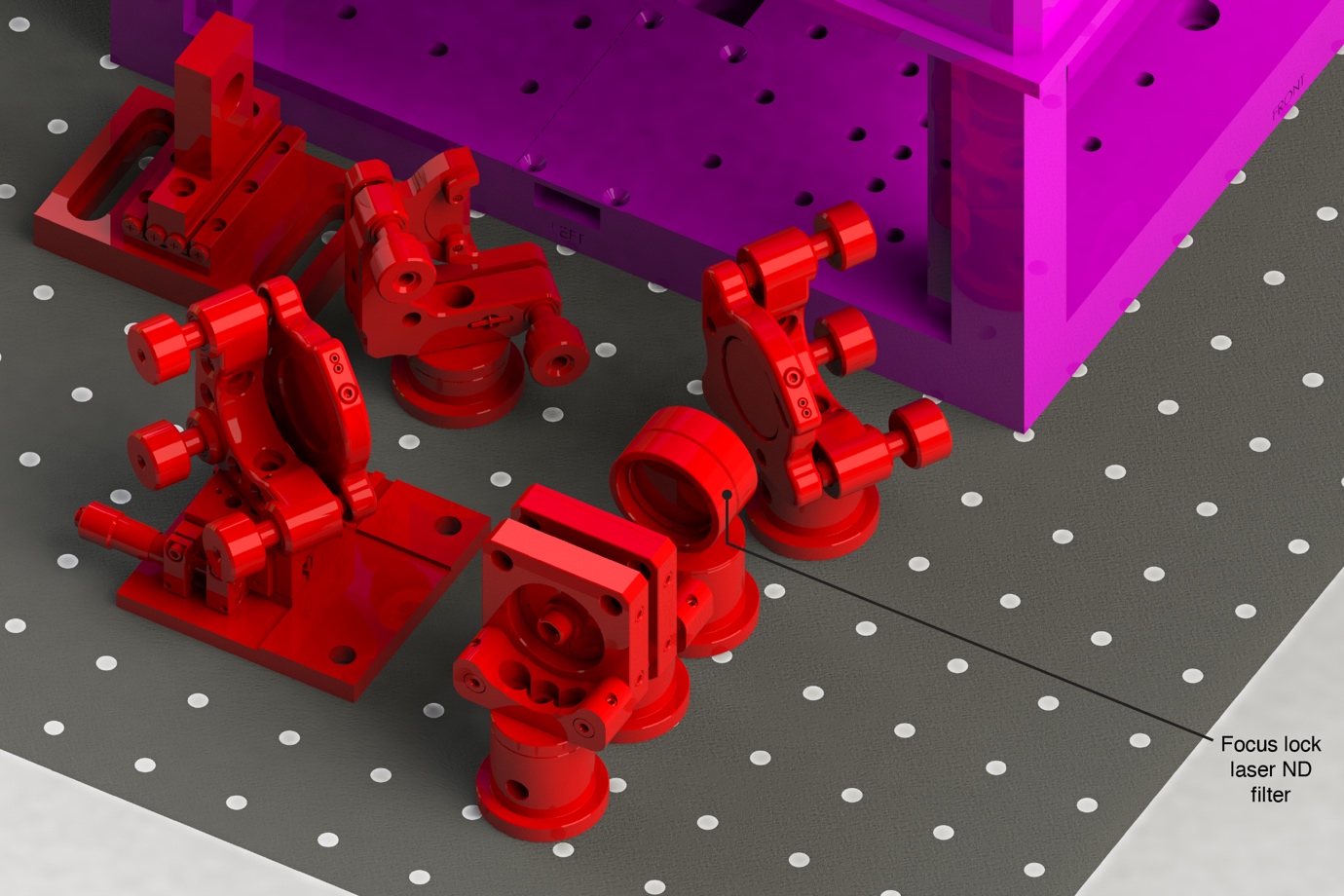

Supplementary Figure 58. Installation of the focus lock laser neutral density filter.

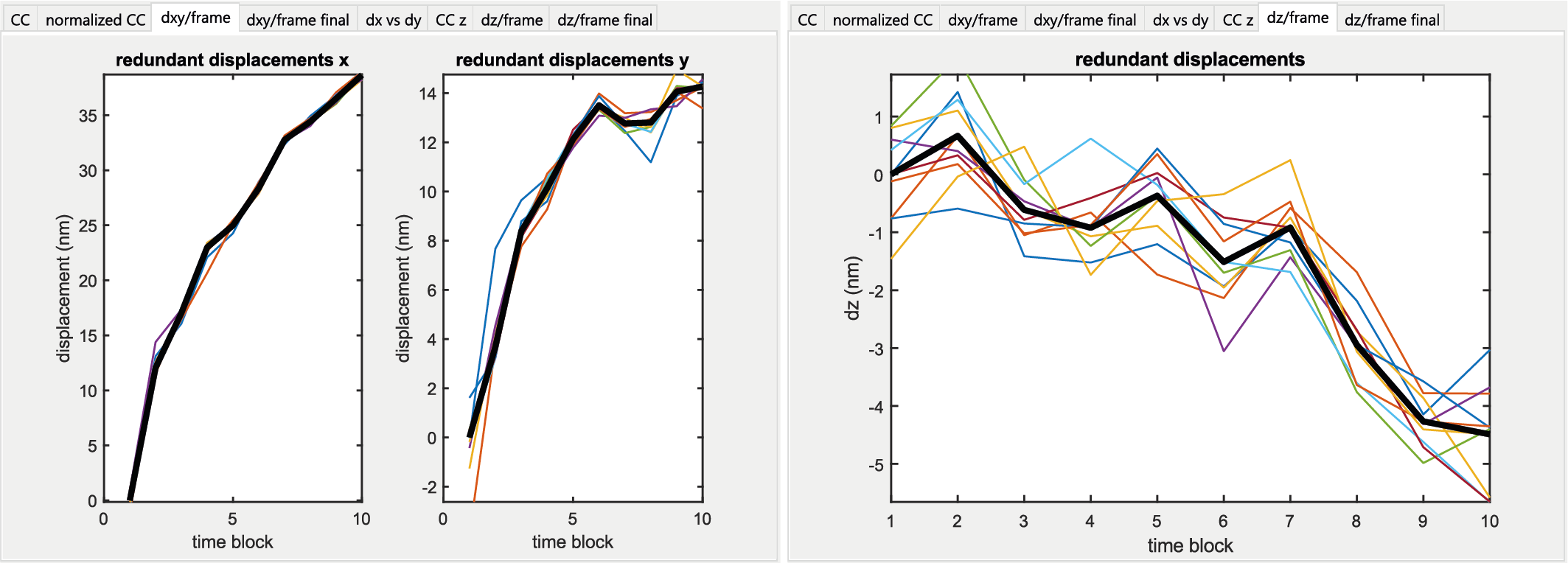

Supplementary Figure 59. An example of drift correction applied to the data shown in Figure 9. Drift correction is performed in SMAP using an algorithm based on redundant cross-correlations. The data are distributed into time bins of equal length (10 in this case), for which super-resolution images were reconstructed. Cross-correlation among all super-resolved images were then calculated to extract the relative displacements in x and y (left). Individual (colored lines) and mean (black lines) displacements between maxima of cross-correlated images are plotted (left) and interpolated by a smoothed spline function (see dz/frame final). This interpolation was subsequently used to correct the xy coordinates of localizations. Displacements in z are calculated analogously using intensity profiles in z as opposed to images for cross-correlation (right).
