## Supplementary Table 1 for "Automated 3D multi-color single-molecule localization microscopy"

| | miCube | K2TIRF | liteTIRF | NanoPro 1.0 | 3D-SMLM |
|---|---|---|---|---|---|
| Lateral spatial res (reported) | < 50 nm (requirement of study but unconfirmed) | Unknown | < 10 nm on origami after drift correct | < 20 nm (20 nm nanoruler resolvable after drift correction) | < 12 nm on nuclear pore samples - resolving distance between Nup96 proteins |
| Axial spatial res (reported) | N/A | N/A | < 86 nm on tetrahedral origamis | | |
| 3D capability | No | No | Yes (non standard piezo free), distortion. Non-trivial adjustment, not automated, no correction collar | | Yes (distortion free, trivial adjustment, automated, integrated analysis) |
| Flat field | No | Yes | No | Yes | Yes |
| TIRF | Yes, manual | Yes, manual | Yes, manual | Not shown | Yes, automated |
| Variable FOV | No | No | No | No | Yes |
| Laser power monitoring | No | No | No | No | Yes |
| Low cost option available | Yes (lasers) | No | Yes (it is low cost) | Hardware noted but not integrated into control scheme | Yes (laser engine/exchangeable hardware) |
| Temperature control | No | Objective heater and enclosure, Omr | No | Not shown (enclosure though) | Body heating possible but not shown |
| Gas control | No | No | No | Not shown (enclosure though) | No |
| Multi-sample height | No | Yes (Smaract 3D stage - 30 mm trave | Yes (13 mm z travel) | Yes (Smaract 3D stage 50 mm travel) | Yes manual 6 mm travel |
| Multi-color imaging (*) | No | Yes | No | Yes | Yes |
| Synchronous multi-color imaging | No | Yes (3 channels) | No | No | Yes (2 channels) |
| Bertrand lens | No | No | No | No | Yes |
| High-throughput multi-FOV | No (no focus lock) | Yes | No | Yes | Yes |
| Smart activation with emitter counting | No | No | No | No | Yes |
| Axial drift (focus) stabilization | No | Yes | No | Yes | Yes |
| Lateral drift stabilization | No | No | No | No | No (handled in integrated analysis pipeline) |
| Multi-laser co-alignment | High-quality laser engine internal | High-quality laser engine internal | N/A | External self-aligned combiner | High-quality laser engine internal, with automated re-optimization |
| Inherent multi-device support | No | No | No | No | Yes (MM2 basis) |

| | | | | | |
|---|---|---|---|---|---|
| Footprint /mm | 600 x 300 | 1800 x 900 | 450 x 300 mm | 1500 x 900 mm | 1500 x 900 |
| Cost /USD equivalent | | | | | |
| Full instructions available | Buildup instructions | No (miCube buildup) | No | Yes | Yes |
| Complexity | Low | Moderate | Low | Moderate | Moderate |
| Project currently maintained | Yes | Yes | Yes | Yes | Yes |
