## Supplementary Table 2 for "Automated 3D multi-color single-molecule localization microscopy"

### EMBL 3D-SMLM Assemblies List

| Part number | Category | Description | Quantity | | | | | | Comments |
|---|---|---|---|---|---|---|---|---|---|
| | | | Base | SM engine | Booster | MM engine | | | |
| 3D-SMLM-EMBL-001 | Assembly | Master assembly | 1 | | | | | | |
| 3D-SMLM-EMBL-001-001 | Assembly | Microscope body | 1 | | | | | | |
| 3D-SMLM-EMBL-001-001-001 | Assembly | Mounted emission tube lens | 1 | | | | | | |
| 3D-SMLM-EMBL-001-002 | Assembly | Emission path | 1 | | | | | | |
| 3D-SMLM-EMBL-001-002-001 | Assembly | Mounted astigmatic lens and piezo stage | 1 | | | | | | |
| 3D-SMLM-EMBL-001-002-002 | Assembly | Mounted image splitter dichroic | 1 | | | | | | |
| 3D-SMLM-EMBL-001-002-003 | Assembly | Mounted knife-edge mirror | 1 | | | | | | |
| 3D-SMLM-EMBL-001-002-004 | Assembly | Camera assembly | 1 | | | | | | |
| 3D-SMLM-EMBL-001-003 | Assembly | Illumination path | 1 | | | | | | |
| 3D-SMLM-EMBL-001-003-001 | Assembly | Single-mode laser engine beam launch | | 1 | | | | | |
| 3D-SMLM-EMBL-001-003-002 | Assembly | Booster laser spatial filter | | | | | | | |
| 3D-SMLM-EMBL-001-003-003 | Assembly | Illumination path polaris mirror left-handed 80.4/110.4 mm beam height | | 2 | 2 | | | | |
| 3D-SMLM-EMBL-001-003-004 | Assembly | Illumination path polaris mirror right-handed 80.4/110.4 mm beam height | | 1 | 2 | | | | |
| 3D-SMLM-EMBL-001-003-005 | Assembly | Mounted refractive beam shaper | | 1 | | | | | |
| 3D-SMLM-EMBL-001-003-006 | Assembly | Kinematic magnetic variable magnification telescope | | 1 | | | | | |
| 3D-SMLM-EMBL-001-003-007 | Assembly | Illumination periscope and inclination control assembly | | 1 | | | | | |
| 3D-SMLM-EMBL-001-003-008 | Assembly | Multi-mode laser engine beam launch | | | | 1 | | | |
| 3D-SMLM-EMBL-001-003-009 | Assembly | Multi-mode laser engine retro focus tube lens and image crop slit | | | | 1 | | | |
| 3D-SMLM-EMBL-001-004 | Assembly | Focus lock path | 1 | | | | | | |
| 3D-SMLM-EMBL-001-004-001 | Assembly | Focus lock laser beam launch | 1 | | | | | | |
| 3D-SMLM-EMBL-001-004-002 | Assembly | Focus lock laser beam translator | | | | | | | |
| 3D-SMLM-EMBL-001-004-003 | Assembly | Quadrant photodiode translation stage assembly | | | | | | | |
| 3D-SMLM-EMBL-002 | Tool Assembly | Tube lens alignment target | 1 | | | | | | |
| 3D-SMLM-EMBL-003 | Tool Assembly | 80.4/80/110.4 mm beam height alignment target | 2 | | | | | | |
| 3D-SMLM-EMBL-004 | Tool Assembly | Infinite conjugate viewing camera and infinite conjugate transmission target | | | | | | | |
| 3D-SMLM-EMBL-005 | Tool Assembly | 80.4/80 mm/objective port configurable shear plate aligner | 1 | | | | | | |
| 3D-SMLM-EMBL-006 | Tool Assembly | Objective port alignment target | 1 | | | | | | |
| 3D-SMLM-EMBL-007 | Tool Assembly | Alignment laser objective port/tube lens port assembly | 1 | | | | | | |
| 3D-SMLM-EMBL-008 | Tool Assembly | 80.4 mm magnetic kinematic alignment target | | 2 | | | | | |
| 3D-SMLM-EMBL-009 | Tool Assembly | Splitter platform alignment targets for out of body alignment | 1 | | | | | | Uses parts from emission path |

### EMBL 3D-SMLM Custom Parts List

| Part number | Category | Description | Material | Quantity | | | | | Comments |
|---|---|---|---|---|---|---|---|---|---|
| 3D-SMLM-FAB-EMBL-000001 | Mechanical | Body bottom plate | Aluminum | 1 | | | | | |
| 3D-SMLM-FAB-EMBL-000002 | Mechanical | Body lower back-right standoff | Aluminum | 1 | | | | | Configurable for heating foil recess |
| 3D-SMLM-FAB-EMBL-000003 | Mechanical | Body lower back-left standoff | Aluminum | 1 | | | | | |
| 3D-SMLM-FAB-EMBL-000004 | Mechanical | Body lower front-right standoff | Aluminum | 1 | | | | | |
| 3D-SMLM-FAB-EMBL-000005 | Mechanical | Body lower front-left standoff | Aluminum | 1 | | | | | |
| 3D-SMLM-FAB-EMBL-000006 | Mechanical | Body insulating intermediate plate | Aluminum/Plastic | 1 | | | | | If not including the heating foil (see below part), then aluminum/ otherwise plastic |
| 3D-SMLM-FAB-EMBL-000007 | Mechanical | Body middle plate | Aluminum | 1 | | | | | Configurable for heating foil recess |
| 3D-SMLM-FAB-EMBL-000008 | Mechanical | Body middle back-right-standoff | Aluminum | 1 | | | | | |
| 3D-SMLM-FAB-EMBL-000009 | Mechanical | Body middle back-left standoff | Aluminum | 1 | | | | | |
| 3D-SMLM-FAB-EMBL-000010 | Mechanical | Body middle front-right standoff | Aluminum | 1 | | | | | |
| 3D-SMLM-FAB-EMBL-000011 | Mechanical | Body middle front-left standoff | Aluminum | 1 | | | | | |
| 3D-SMLM-FAB-EMBL-000012 | Mechanical | Body top plate | Aluminum | 1 | | | | | |
| 3D-SMLM-FAB-EMBL-000013 | Mechanical | Body lower back cover | Plastic | 1 | | | | | |
| 3D-SMLM-FAB-EMBL-000014 | Mechanical | Body lower right cover | Plastic | 1 | | | | | |
| 3D-SMLM-FAB-EMBL-000015 | Mechanical | Body lower front cover | Plastic | 1 | | | | | |
| 3D-SMLM-FAB-EMBL-000016 | Mechanical | Body lower left cover | Plastic | 1 | | | | | |
| 3D-SMLM-FAB-EMBL-000017 | Mechanical | Body LED cable strain relief | Plastic | 2 | | | | | Can be 3D printed |
| 3D-SMLM-FAB-EMBL-000018 | Mechanical | Body middle cover | Plastic | 4 | | | | | |
| 3D-SMLM-FAB-EMBL-000019 | Mechanical | Body cover back-front | Plastic | 1 | | | | | |
| 3D-SMLM-FAB-EMBL-000020 | Mechanical | Body cover left-right | Plastic | 1 | | | | | |
| 3D-SMLM-FAB-EMBL-000021 | Mechanical | Body cover top | Aluminum | 1 | | | | | |
| 3D-SMLM-FAB-EMBL-000022 | Mechanical | Body optical pillar base plate | Aluminum | 1 | | | | | Require either this part or mirrored version below |
| 3D-SMLM-FAB-EMBL-000023 | Mechanical | Body optical pillar | Aluminum | 1 | | | | | Require either this part or mirrored version below |
| 3D-SMLM-FAB-EMBL-000024 | Mechanical | Body focus lock mirror mount | Aluminum | 1 | | | | | Require either this part or mirrored version below |
| 3D-SMLM-FAB-EMBL-000025 | Mechanical | Body emission dichroic mount | Aluminum | 1 | | | | | Require either this part or mirrored version below |
| 3D-SMLM-FAB-EMBL-000026 | Mechanical | Body illumination dichroic mount 3 mm thickness | Aluminum | 1 | | | | | Require either this part or mirrored version below |
| 3D-SMLM-FAB-EMBL-000027 | Mechanical | Body shared path piezo filter slider mount | Aluminum | 1 | | | | | Require either this part or mirrored version below |
| 3D-SMLM-FAB-EMBL-000028 | Mechanical | Body optical pillar base plate mirrored | Aluminum | 1 | | | | | |
| 3D-SMLM-FAB-EMBL-000029 | Mechanical | Body optical pillar mirrored | Aluminum | 1 | | | | | |
| 3D-SMLM-FAB-EMBL-000030 | Mechanical | Body focus lock mirror mount mirrored | Aluminum | 1 | | | | | |
| 3D-SMLM-FAB-EMBL-000031 | Mechanical | Body emission dichroic mount mirrored | Aluminum | 1 | | | | | |
| 3D-SMLM-FAB-EMBL-000032 | Mechanical | Body illumination dichroic mount 3 mm thickness mirrored | Aluminum | 1 | | | | | |
| 3D-SMLM-FAB-EMBL-000033 | Mechanical | Body shared path piezo filter slider mount mirrored | Aluminum | 1 | | | | | |
| 3D-SMLM-FAB-EMBL-000034 | Mechanical | Body tube lens referencing base | Aluminum | 1 | | | | | |
| 3D-SMLM-FAB-EMBL-000035 | Mechanical | Body tube lens split clamp lower | Aluminum | 1 | | | | | |
| 3D-SMLM-FAB-EMBL-000036 | Mechanical | Body tube lens split clamp upper | Aluminum | 1 | | | | | |

| | | | | | | | | | |
|---|---|---|---|---|---|---|---|---|---|
| 3D-SMLM-FAB-EMBL-000037 | Electronic | Ring LED | | 1 | | | | | |
| 3D-SMLM-FAB-EMBL-000038 | Mechanical | Body objective piezo translation stage plate | Aluminum | 1 | | | | | |
| 3D-SMLM-FAB-EMBL-000039 | Mechanical | Body objective piezo translation stage actuator M6x0.25 | Stainless steel | 3 | | | | | Require either this part or hex version below |
| 3D-SMLM-FAB-EMBL-000040 | Mechanical | Body objective piezo translation stage actuator hex | Stainless steel | 3 | | | | | |
| 3D-SMLM-FAB-EMBL-000041 | Mechanical | Body heating foil cover | Aluminum | 1 | | | | | Configurable for heating foil recess |
| 3D-SMLM-FAB-EMBL-000042 | Mechanical | Body heating foil | | 1 | | | | | Configurable for heating foil recess |
| 3D-SMLM-FAB-EMBL-000043 | Mechanical | Body objective thread spacer RMS 45 mm parfocal length | Brass | 1 | | | | | |
| 3D-SMLM-FAB-EMBL-000044 | Mechanical | Body XY stage sample holder adapter plate | Aluminum | 1 | | | | | |
| 3D-SMLM-FAB-EMBL-000045 | Mechanical | Body sample holder plate | Magnetic steel | 3 | | | | | |
| 3D-SMLM-FAB-EMBL-000046 | Mechanical | Body sample holder magnet ring cover | Plastic | 3 | | | | | |
| 3D-SMLM-FAB-EMBL-000047 | Mechanical | Body XY stage spacer | Plastic | 4 | | | | | |
| 3D-SMLM-FAB-EMBL-000048 | Mechanical | Body XY stage sample holder adapter plate legacy stages | Aluminum | 1 | | | | | |
| 3D-SMLM-FAB-EMBL-000049 | Mechanical | Body magnetic safety switch mount | Plastic | 2 | | | | | |
| 3D-SMLM-FAB-EMBL-000050 | Mechanical | Body cable cover bottom plate | Aluminum | 1 | | | | | |
| 3D-SMLM-FAB-EMBL-000051 | Mechanical | Body cable cover heating foil and sensor 1 | Aluminum | 1 | | | | | Configurable for heating foil recess |
| 3D-SMLM-FAB-EMBL-000052 | Mechanical | Body cable cover heating foil and sensor 2 | Aluminum | 1 | | | | | Configurable for heating foil recess |
| 3D-SMLM-FAB-EMBL-000053 | Mechanical | Body cable cover heating foil and sensor 3 | Aluminum | 1 | | | | | Configurable for heating foil recess |
| 3D-SMLM-FAB-EMBL-000054 | Mechanical | Body cable cover objective piezo 1 | Aluminum | 1 | | | | | |
| 3D-SMLM-FAB-EMBL-000055 | Mechanical | Body cable cover objective piezo 2 | Aluminum | 1 | | | | | |
| 3D-SMLM-FAB-EMBL-000056 | Mechanical | Body cable cover stage and magnetic switches 1 | Aluminum | 1 | | | | | |
| 3D-SMLM-FAB-EMBL-000057 | Mechanical | Body cable cover stage and magnetic switches 2 | Aluminum | 1 | | | | | |
| 3D-SMLM-FAB-EMBL-000058 | Mechanical | Body cable cover stage and magnetic switches 3 | Aluminum | 1 | | | | | |
| 3D-SMLM-FAB-EMBL-000059 | Mechanical | Body cable channel | Aluminum | 1 | | | | | |
| 3D-SMLM-FAB-EMBL-000060 | Mechanical | Body emission dichroic mount 5 mm thickness | Aluminum | 1 | | | | | Replace 3 mm versions above for thicker substrates |
| 3D-SMLM-FAB-EMBL-000061 | Mechanical | Body illumination dichroic mount 5 mm thickness | Aluminum | 1 | | | | | Replace 3 mm versions above for thicker substrates |
| 3D-SMLM-FAB-EMBL-000062 | Mechanical | Body emission dichroic mount 5 mm thickness mirrored | Aluminum | 1 | | | | | Replace 3 mm versions above for thicker substrates |
| 3D-SMLM-FAB-EMBL-000063 | Mechanical | Body illumination dichroic mount 5 mm thickness mirrored | Aluminum | 1 | | | | | Replace 3 mm versions above for thicker substrates |
| 3D-SMLM-FAB-EMBL-000064 | Mechanical | Body idler pin M4 tap | Steel | 1 | | | | | |
| 3D-SMLM-FAB-EMBL-000065 | Mechanical | Body idler channel cover | Plastic | 1 | | | | | |
| 3D-SMLM-FAB-EMBL-000066 | Mechanical | Body cover clearance lip | Plastic | 1 | | | | | |
| 3D-SMLM-FAB-EMBL-000067 | Mechanical | Astigmatic lens stage base plate | Aluminum | 1 | | | | | |
| 3D-SMLM-FAB-EMBL-000068 | Mechanical | Splitter dichroic kinematic to magnetic mount plate | Aluminum | 1 | | | | | |
| 3D-SMLM-FAB-EMBL-000069 | Mechanical | Splitter dichroic mount 3 mm thickness | Aluminum | 1 | | | | | |
| 3D-SMLM-FAB-EMBL-000070 | Mechanical | Splitter dichroic mount 5 mm thickness | Aluminum | 1 | | | | | Replace 3 mm versions above for thicker substrates |
| 3D-SMLM-FAB-EMBL-000071 | Mechanical | Camera base plate | Aluminum | 1 | | | | | |
| 3D-SMLM-FAB-EMBL-000072 | Mechanical | Camera mounting plate | Aluminum | 1 | | | | | |
| 3D-SMLM-FAB-EMBL-000073 | Mechanical | Camera c-mount fastener | Stainless steel | 1 | | | | | |
| 3D-SMLM-FAB-EMBL-000074 | Mechanical | Camera Bertrand lens piezo stage mount | Aluminum | 1 | | | | | |
| 3D-SMLM-FAB-EMBL-000075 | Mechanical | Knife-edge mirror pedestal | Aluminum | 2 | | | | | |
| 3D-SMLM-FAB-EMBL-000076 | Mechanical | Filter wheel mount | Aluminum | 2 | | | | | |
| 3D-SMLM-FAB-EMBL-000077 | Mechanical | Splitter base plate | Aluminum | 1 | | | | | |
| 3D-SMLM-FAB-EMBL-000078 | Mechanical | Splitter relay lens split clamp | Aluminum | 2 | | | | | |
| 3D-SMLM-FAB-EMBL-000079 | Mechanical | Splitter reflected path steering mirror pedestal | Aluminum | 1 | | | | | |
| 3D-SMLM-FAB-EMBL-000080 | Mechanical | Splitter transmitted path steering mirror pedestal | Aluminum | 1 | | | | | |
| 3D-SMLM-FAB-EMBL-000081 | Mechanical | Splitter alignment target | Aluminum | 2 | | | | | |
| 3D-SMLM-FAB-EMBL-000082 | Mechanical | piShaper mount | Aluminum | 1 | | | | | |
| 3D-SMLM-FAB-EMBL-000083 | Mechanical | Illumination control bottom plate | Aluminum | 1 | | | | | |
| 3D-SMLM-FAB-EMBL-000084 | Mechanical | Illumination control entrance plate | Aluminum | 1 | | | | | |
| 3D-SMLM-FAB-EMBL-000085 | Mechanical | Illumination control exit plate | Aluminum | 1 | | | | | |
| 3D-SMLM-FAB-EMBL-000086 | Mechanical | Illumination control prism mirror mount | Aluminum | 1 | | | | | |
| 3D-SMLM-FAB-EMBL-000087 | Mechanical | Toptica iBeam smart free space mounting plate | Aluminum | 1 | | | | | Required for booster laser |
| 3D-SMLM-FAB-EMBL-000088 | Mechanical | Toptica iBeam smart PT mounting plate | Aluminum | 1 | | | | | |
| 3D-SMLM-FAB-EMBL-000089 | Mechanical | QPD stage base | Aluminum | 1 | | | | | |
| 3D-SMLM-FAB-EMBL-000090 | Mechanical | QPD stage mount | Aluminum | 1 | | | | | |
| 3D-SMLM-FAB-EMBL-000091 | Mechanical | QPD mount | Aluminum | 1 | | | | | |
| 3D-SMLM-FAB-EMBL-000092 | Mechanical | Focus lock BFP offset stage mount | Aluminum | 1 | | | | | |
| 3D-SMLM-FAB-EMBL-000093 | Mechanical | Camera image alignment target | Aluminum | 1 | | | | | |
| 3D-SMLM-FAB-EMBL-000094 | Mechanical | Multimode laser engine | Aluminum | 1 | | | | | Required for multimode laser engine |
| 3D-SMLM-FAB-EMBL-000095 | Mechanical | Elliptec control board base | 3D print | 1 | | | | | |
| 3D-SMLM-FAB-EMBL-000096 | Mechanical | Elliptec control board cover | 3D print | 1 | | | | | |
| 3D-SMLM-FAB-EMBL-000097 | Mechanical | Body illumination dichroic mount 1 mm thickness | Aluminum | 1 | | | | | Replace 3 mm versions above for thinner substrates |
| 3D-SMLM-FAB-EMBL-000098 | Mechanical | Body illumination dichroic mount 1 mm thickness mirrored | Aluminum | 1 | | | | | Replace 3 mm versions above for thinner substrates |
| 3D-SMLM-FAB-EMBL-000099 | Electronic | microFPGA front panel 1 | Various | 1 | | | | | |
| 3D-SMLM-FAB-EMBL-000100 | Electronic | microFPGA front panel 2 | Various | 1 | | | | | |
| 3D-SMLM-FAB-EMBL-000101 | Electronic | microFPGA front panel 3 | Various | 1 | | | | | |
| 3D-SMLM-FAB-EMBL-000102 | Electronic | microFPGA custom shield | Various | 1 | | | | | |
| 3D-SMLM-FAB-EMBL-000103 | Electronic | microFPGA analog conversion board | Various | 1 | | | | | |
| 3D-SMLM-FAB-EMBL-000104 | Electronic | microFPGA signal conversion board | Various | 1 | | | | | |
| 3D-SMLM-FAB-EMBL-000105 | Electronic | microFPGA enclosure lid | Aluminum | 1 | | | | | modified from 3D-SMLM-COTS-FC-102520-GS |
| 3D-SMLM-FAB-EMBL-000106 | Electronic | microFPGA side panel 1 | Aluminum | 1 | | | | | modified from 3D-SMLM-COTS-FC-102520-GS |
| 3D-SMLM-FAB-EMBL-000107 | Electronic | microFPGA side panel 2 | Aluminum | 1 | | | | | modified from 3D-SMLM-COTS-FC-102520-GS |
| 3D-SMLM-FAB-EMBL-000108 | Electronic | microFPGA side panel 3 | Aluminum | 1 | | | | | modified from 3D-SMLM-COTS-FC-102520-GS |
| 3D-SMLM-FAB-EMBL-000109 | Electronic | microFPGA side panel 4 | Aluminum | 1 | | | | | modified from 3D-SMLM-COTS-FC-102520-GS |

| | | | | | | | | | |
|---|---|---|---|---|---|---|---|---|---|
| 3D-SMLM-FAB-EMBL-000110 | Electronic | microFPGA enclosure base | Aluminum | 1 | | | | | modified from 3D-SMLM-COTS-FC-102520-GS |
| 3D-SMLM-FAB-EMBL-000111 | Electronic | Filter wheel mount base | Aluminum | 2 | | | | | |
| 3D-SMLM-FAB-EMBL-000112 | Electronic | LED controller front panel | Aluminum | 1 | | | | | modified from 3D-SMLM-COTS-FC-101010-GS |
| 3D-SMLM-FAB-EMBL-000113 | Electronic | LED controller top panel | Aluminum | 1 | | | | | modified from 3D-SMLM-COTS-FC-101010-GS |
| 3D-SMLM-FAB-EMBL-000114 | Electronic | LED controller rear panel | Aluminum | 1 | | | | | modified from 3D-SMLM-COTS-FC-101010-GS |
| **EMBL 3D-SMLM COTS Parts List** | | | | | | | | | |
| **Part number** | **Category** | **Description** | **Vendor** | **Quantity** | | | | | **Comments** |
| | | | | Base | SM engine | Booster | MM engine | Tools | |
| 3D-SMLM-COTS-XY-STAGE-LEGACY-RIES | Motion control | Custom XY stage assembly (Ries) | Smaract | 1 | | | | | Legacy component |
| 3D-SMLM-COTS-XY-STAGE-LEGACY-IC | Motion control | Custom XY stage assembly (Imaging Centre) | Smaract | 1 | | | | | Legacy component |
| 3D-SMLM-COTS-SOM-12090 | Motion control | XY stage | Smaract | 1 | | | | | |
| 3D-SMLM-COTS-2445-L | Motion control | 1D linear stage 25 mm travel | Smaract | 1 | | | | | |
| 3D-SMLM-COTS-HCU-3D | Motion control | 3-axis stage controller | Smaract | 1 | | | | | |
| 3D-SMLM-COTS-P-726.1CD | Motion control | Objective piezo flexure stage | Physik Instrumente | 1 | | | | | |
| 3D-SMLM-COTS-ELL9 | Motion control | 4 position piezo slider | Thorlabs | 1 | | | | | Optional |
| 3D-SMLM-COTS-ELL20-M | Motion control | 60 mm travel piezo stage | Thorlabs | 2 | | | | | |
| 3D-SMLM-COTS-ELL17-M | Motion control | 28 mm travel piezo stage | Thorlabs | | 1 | | | | |
| 3D-SMLM-COTS-ELL6 | Motion control | 2 position piezo slider | Thorlabs | 1 | | | | | |
| 3D-SMLM-COTS-ELLC2 | Motion control | piezo stage control board and power supply | Thorlabs | 1 | | | | | |
| 3D-SMLM-COTS-ELLB | Motion control | piezo stage bus distributor | Thorlabs | 1 | | | | | |
| 3D-SMLM-COTS-FW102C | Motion control | 6 position motorised filter wheel | Thorlabs | 2 | | | | | |
| 3D-SMLM-MS1S-M | Motion control | 6.5 mm manual linear stage | Thorlabs | 1 | | | | | |
| 3D-SMLM-COTS-SFL625ZZ | Mechanical | Bearing | Misumi | 3 | | | | | |
| 3D-SMLM-COTS-HTPT48S2M060_A_P6_35 | Mechanical | S2M timing pulley | Misumi | 3 | | | | | |
| 3D-SMLM-COTS-SFDF11 | Mechanical | Idler | Misumi | 1 | | | | | |
| 3D-SMLM-COTS-F6MSSA1 | Mechanical | M6x0.25 threaded bushing | Thorlabs | 3 | | | | | |
| 3D-SMLM-COTS-F25US100 | Mechanical | M6x0.25 hex actuator | Thorlabs | 3 | | | | | If not using the custom actuator |
| 3D-SMLM-COTS-60S2M494 | Mechanical | S2M timing belt 494 mm | Misumi | 1 | | | | | Not in model |
| 3D-SMLM-COTS-91585A437 | Mechanical | 4x10 mm dowel pin | McMaster | 6 | | | | | |
| 3D-SMLM-COTS-91585A371 | Mechanical | 3x12 mm dowel pin | McMaster | 12 | | | | | |
| 3D-SMLM-COTS-24540.00021 | Mechanical | Panel knob M5 threaded | Misumi | 12 | | | | | |
| 3D-SMLM-COTS-98952A104 | Mechanical | 6 mm M3 thread/tap hex standoff | McMaster | 4 | | | | | |
| 3D-SMLM-COTS-93655A099 | Mechanical | 12 mm M3 thread/tap hex standoff | McMaster | 8 | | | | | |
| 3D-SMLM-COTS-91585A668 | Mechanical | 6x30 mm dowel pin | McMaster | 1 | | | | | |
| 3D-SMLM-COTS-91585A454 | Mechanical | 4x16 mm dowel pin | McMaster | 5 | | | | | |
| 3D-SMLM-COTS-94868A002 | Mechanical | 4 mm M3 tap/tap hex standoff | McMaser | 4 | | | | | |
| 3D-SMLM-COTS-91075A873 | Mechanical | 1/8" 4-40 thread/tap hex standoff | McMaser | 4 | | | | | Only needed for ELL9 tertiary filter wheel |
| 3D-SMLM-COTS-RZ099L | Mechanical | 6 mm x 25.5 mm hooked spring | Gutekunst Federn | 6 | | | | | |
| 3D-SMLM-COTS-S-08-05-N | Mechanical | 8 mm diameter, 5 mm height magnet | Supermagnete | 16 |  |  |  |  |  |
| 3D-SMLM-COTS-AP6M4M | Mechanical | M6 to M4 external to external thread adapter | Thorlabs | 2 | | | | | |
| 3D-SMLM-COTS-94459140 | Mechanical | Heat installed thread insert M3 | McMaster | 4 | | | | | Adjust bore diameter in 3D-SMLM-FAB-EMBL-000095 for thread insert |
| 3D-SMLM-COTS-9292K37 | Mechanical | 4 mm alloy steel balls | McMaster | 3 | | | | | If using the custom actuator |
| 3D-SMLM-COTS-UPLAPO100XOHR | Optics | Oil immersion TIRF objective | Olympus | 1 | | | | | As specified by the builder |
| 3D-SMLM-COTS-UPLXAPO100XO | Optics | Oil immersion objective | Olympus | 1 | | | | | As specified by the builder |
| 3D-SMLM-COTS-UPLSAPO100XS | Optics | Silicone oil immersion objective | Olympus | 1 | | | | | As specified by the builder |
| 3D-SMLM-COTS-25.5x36x3MM-DICHROIC | Optics | Dichroic mirror | Semrock/Chroma | 3 | | | | | As specified by the builder (note dimensions), dichroics called out in protocol with suffix -ILL/-EM/-SP |
| 3D-SMLM-COTS-25.5x36x5MM-DICHROIC | Optics | Dichroic mirror | Semrock/Chroma | 3 | | | | | As specified by the builder (note dimensions), alternative to above |
| 3D-SMLM-COTS-25MM-HOUSED-FILTER | Optics | Laser cleanup/emission/notch filter | Semrock/Chroma | >6 | | | | | As specified by the builder |
| 3D-SMLM-COTS-MRA20-E03 | Optics | 90 degree NIR prism mirror | Thorlabs | 1 | | | | | |
| 3D-SMLM-COTS-LJ1516RM | Optics | effl = 1000 mm Plano-convex cylindrical lens mounted | Thorlabs | 1 | | | | | |
| 3D-SMLM-COTS-LK1002RM | Optics | effl = -1000 mm Plano-convex cylindrical lens mounted | Thorlabs | 1 | | | | | |
| 3D-SMLM-COTS-#32-963 | Optics | effl = 60 mm Plano-convex lens (20 mm diameter) | Edmund Optics | 1 | | | | | |
| 3D-SMLM-COTS-MRAK25-P01 | Optics | 25 mm knife-edge mirror | Thorlabs | 1 | | | | | |
| 3D-SMLM-COTS-TTL165-A | Optics | effl = 165 mm widefield tube lens | Thorlabs | 1 | | | | | |
| 3D-SMLM-COTS-BB1-E02 | Optics | 1" diameter A-coated broadband mirror | Thorlabs | 3 | 4 | 4 | 2 | | |
| 3D-SMLM-COTS-AC254-300-A-ML | Optics | effl = 300 mm achromatic doublet, mounted with SM1 thread | Thorlabs | 1 | | | | | |
| 3D-SMLM-COTS-AC254-200-A | Optics | effl = 200 mm achromatic doublet | Thorlabs | 2 | | | | | |
| 3D-SMLM-COTS-PISHAPER-6_6_VIS | Optics | piShaper 6 mm beam in/out, visible | AdlOptica | | 1 | | | | |
| 3D-SMLM-COTS-AC254-075-A-ML | Optics | effl = 75 mm achromatic doublet, mounted with SM1 thread | Thorlabs | | 2 | | | | |
| 3D-SMLM-COTS-AC254-060-A | Optics | effl = 60 mm achromatic doublet | Thorlabs | | 2 | | | | |
| 3D-SMLM-COTS-AC254-040-A | Optics | effl = 40 mm achromatic doublet | Thorlabs | | 2 | | 2 | | |
| 3D-SMLM-COTS-AC254-080-A | Optics | effl = 80 mm achromatic doublet | Thorlabs | | 2 | | | | |
| 3D-SMLM-COTS-AC254-075-A | Optics | effl = 75 mm achromatic doublet | Thorlabs | | 1 | | | | 1x Only needed for 2.85x telescope |
| 3D-SMLM-COTS-AC254-050-A | Optics | effl = 50 mm achromatic doublet | Thorlabs | | 1 | | | | 1x Only needed for 2.85x telescope |
| 3D-SMLM-COTS-#49-769 | Optics | effl = 88.9 mm achromatic doublet | Edmund Optics | | 1 | | | | 1x Only needed for 2.85x telescope |
| 3D-SMLM-COTS-MRA20-E02 | Optics | 20 mm A-coated broadband prism mirror | Thorlabs | | 1 | | | | |

| | | | | | | | | | |
|---|---|---|---|---|---|---|---|---|---|
| 3D-SMLM-COTS-PBS251 | Optics | 1" polarisation beam splitter cube | Thorlabs | | | 1 | | | |
| 3D-SMLM-COTS-LA1986-A-ML | Optics | effl = 125 mm Plano-convex lens, mounted with SM1 thread | Thorlabs | | | 2 | | | |
| 3D-SMLM-COTS-LA1540-A-ML | Optics | effl = 15 mm Plano-convex lens, mounted with SM05 thread | Thorlabs | | | 1 | | | |
| 3D-SMLM-COTS-AC127-019-B-ML | Optics | effl = 19 mm achromatic doublet, mounted with SM05 thread, B-coated | Thorlabs | 1 | | | | | |
| 3D-SMLM-COTS-BB1-E03 | Optics | 1" diameter B-coated broadband mirror | Thorlabs | 2 | | | | | |
| 3D-SMLM-COTS-NE10A-B | Optics | B-coated absorptive neutral density filter, ND = 1 | Thorlabs | 1 | | | | | |
| 3D-SMLM-COTS-NF808-34 | Optics | 808 nm notch filter | Thorlabs | 1 | | | | | |
| 3D-SMLM-COTS-FBH810-10 | Optics | 808 nm cleanup filter | Thorlabs | 1 | | | | | |
| 3D-SMLM-COTS-PFD10-03-P01 | Optics | D-shaped mirror protected silver | Thorlabs | 1 | | | | | |
| 3D-SMLM-COTS-ACN254-050-A | Optics | effl = -50 mm achromatic doublet | Thorlabs | | | | 1 | | |
| 3D-SMLM-COTS-AC508-100-A | Optics | effl = 100 mm achromatic doublet (2") | Thorlabs | | | | 1 | | |
| 3D-SMLM-COTS-DG10-1500-H1-MD | Optics | 1" ground glass diffuser alignment target mounted | Thorlabs | | | | | 1 | |
| 3D-SMLM-COTS-SI100 | Optics | Shear interferometer with 5 - 10 mm beam plate | Thorlabs | | | | | 1 | |
| 3D-SMLM-COTS-SI050P | Optics | Shear plate for 2.5 - 5 mm beam | Thorlabs | | | | | 1 | |
| 3D-SMLM-COTS-SI254P | Optics | Shear plate for 10 - 25.4 mm beam | Thorlabs | | | | | 1 | |
| 3D-SMLM-COTS-NE10A-A | Optics | A-coated absorptive neutral density filter, ND = 1 | Thorlabs | | | | | 1 | |
| 3D-SMLM-COTS-NE20A-A | Optics | A-coated absorptive neutral density filter, ND = 2 | Thorlabs | | | | | 1 | |
| 3D-SMLM-COTS-NE40A-A | Optics | A-coated absorptive neutral density filter, ND = 4 | Thorlabs | | | | | 1 | |
| 3D-SMLM-COTS-AC254-200-A-ML | Optics | effl = 200 mm achromatic doublet, mounted with SM1 thread | Thorlabs | | | | | 1 | |
| 3D-SMLM-COTS-AC254-100-A-ML | Optics | effl = 100 mm achromatic doublet, mounted with SM1 thread | Thorlabs | | | | | 1 | |
| 3D-SMLM-COTS-M-VIS3660-PG4-325A | Optomechanics | 900 x 1500 mm optical table | Newport | 1 | | | | | |
| 3D-SMLM-COTS-KB25-M | Optomechanics | 25 mm kinematic magnetic mount | Thorlabs | 2 | 2 | | | | KB25/M |
| 3D-SMLM-COTS-SM1L03 | Optomechanics | 0.3" length lens tube with SM1 internal and external threads | Thorlabs | 9 | 1 | | | | 4 are for the ELL9 filter slider. Omit if not using |
| 3D-SMLM-COTS-PH30-M | Optomechanics | 30 mm height 1/2" post holder M6 tap | Thorlabs | 1 | | | | | |
| 3D-SMLM-COTS-TRA20-M | Optomechanics | 20 mm length 1/2" post | Thorlabs | 2 | 8 | | | | |
| 3D-SMLM-COTS-RA180-M | Optomechanics | 90 degree M6 to 1/2" post adapter | Thorlabs | 1 | | | | | |
| 3D-SMLM-COTS-TRA40-M | Optomechanics | 40 mm length 1/2" post | Thorlabs | 1 | | | 3 | | |
| 3D-SMLM-COTS-CRM1PT-M | Optomechanics | 30 mm cage rotational mount with micrometer | Thorlabs | 1 | | | | | |
| 3D-SMLM-COTS-CRM1T-M | Optomechanics | 30 mm cage rotational mount | Thorlabs | 1 | | | | | |
| 3D-SMLM-COTS-KM200S | Optomechanics | kinematic mount for rectangular optics | Thorlabs | 1 | | | | | |
| 3D-SMLM-COTS-SB1-M | Optomechanics | round kinematic magnetic mount with release lever | Thorlabs | 1 | | | | | |
| 3D-SMLM-COTS-F25SS050 | Optomechanics | 1/4"-80 fine hex adjuster, 1/2" long | Thorlabs | 2 | | | | | |
| 3D-SMLM-COTS-SM1NT | Optomechanics | Slotted SM1 lock ring | Thorlabs | 1 | | | | 1 | |
| 3D-SMLM-COTS-SM1AD20 | Optomechanics | SM1 externally threaded mounting tube for 20 mm optics | Thorlabs | 1 | | | | | |
| 3D-SMLM-COTS-PM3-M | Optomechanics | M4 threaded clamping mounting post | Thorlabs | 1 | 1 | | | | |
| 3D-SMLM-COTS-SP60 | Optomechanics | 4 bladed rectangular slit aperture | Owis | 1 | | | 1 | | |
| 3D-SMLM-COTS-RS1P4M | Optomechanics | 25 mm height , 25 mm diameter post with M4 taps | Thorlabs | 1 | | | | | |
| 3D-SMLM-COTS-RS05P4M | Optomechanics | 12.5mm height , 25 mm diameter post with M4 taps | Thorlabs | 2 | 2 | | | | |
| 3D-SMLM-COTS-POLARIS-K1 | Optomechanics | Polaris 1" mirror mount with 3 adjusters | Thorlabs | 3 | 3 | 4 | 2 | | |
| 3D-SMLM-COTS-RS5M | Optomechanics | 5 mm height, 25 mm diameter post spacer | Thorlabs | 1 | 3 | 4 | | | |
| 3D-SMLM-COTS-RS2P4M | Optomechanics | 50 mm height, 25 mm diameter pedestal post with M4 taps | Thorlabs | 1 | 3 | 5 | | | |
| 3D-SMLM-COTS-PH50E-M | Optomechanics | 50 mm height, 1/2" post holder pedestal base | Thorlabs | 2 | | | 3 | 2 | |
| 3D-SMLM-COTS-LMR1-M | Optomechanics | 1" lens holder, M4 tap | Thorlabs | 4 | 2 | | | 2 | |
| 3D-SMLM-COTS-SM1M10 | Optomechanics | 1" length lens tube with SM1 internal threads | Thorlabs | 2 | 9 | | | | |
| 3D-SMLM-COTS-ER1 | Optomechanics | 1" cage rod | Thorlabs | 8 | | | | | |
| 3D-SMLM-COTS-PH30-M | Optomechanics | 30 mm height 1/2" post holder M6 tap | Thorlabs | 1 | | | | | |
| 3D-SMLM-COTS-RA180-M | Optomechanics | 90 degree M6 to 1/2" post adapter | Thorlabs | 1 | | | | | |
| 3D-SMLM-COTS-TRA40-M | Optomechanics | 40 mm length 1/2" post | Thorlabs | 1 | | | | | |
| 3D-SMLM-COTS-CRM1PT-M | Optomechanics | 30 mm cage rotational mount with micrometer | Thorlabs | 1 | | | | | |
| 3D-SMLM-COTS-CRM1T-M | Optomechanics | 30 mm cage rotational mount | Thorlabs | 1 | | | | | |
| 3D-SMLM-COTS-KL02L | Optomechanics | Ball tipped kinematic adjuster | Thorlabs | 1 | | | | | |
| 3D-SMLM-COTS-KC1-T-M | Optomechanics | 30 mm cage kinematic tip/tilt/translate mount | Thorlabs | | 1 | | | | |
| 3D-SMLM-COTS-LN2580 | Optomechanics | bronze locking nut for 1/4"-80 adjusters | Thorlabs | | 3 | | | | |
| 3D-SMLM-COTS-S1FCA | Optomechanics | FC/APC terminated fiber adapter | Thorlabs | | 1 | | | | |
| 3D-SMLM-COTS-ER6 | Optomechanics | 6" cage rod | Thorlabs | | 4 | 4 | | | |
| 3D-SMLM-COTS-CP33-M | Optomechanics | 30 mm cage plate SM1 tap | Thorlabs | 1 | 2 | 3 | | | |
| 3D-SMLM-COTS-CP02B | Optomechanics | 30 mm cage mounting bracket | Thorlabs | 2 | 2 | 2 | | | |
| 3D-SMLM-COTS-SM1T2 | Optomechanics | SM1 lens tube coupler | Thorlabs | | 1 | 1 | | | |
| 3D-SMLM-COTS-CXY1 | Optomechanics | 30 mm cage XY mount | Thorlabs | | 1 | 1 | | 1 | |
| 3D-SMLM-COTS-SM1D12D | Optomechanics | SM1 ring actuated iris | Thorlabs | | 4 | | | 4 | |
| 3D-SMLM-COTS-TRA30-M | Optomechanics | 30 mm length 1/2" post | Thorlabs | 1 | 5 | 2 | 2 | | |
| 3D-SMLM-COTS-PH40E-M | Optomechanics | 40 mm height, 1/2" post holder pedestal base | Thorlabs | 1 | 3 | 2 | | 1 | |
| 3D-SMLM-COTS-KBM1-M | Optomechanics | Kinematic magnetic base | Thorlabs | | 1 | | | | |
| 3D-SMLM-COTS-ULM-TILT | Optomechanics | Tip/tilt mount for cylindrical optics | Newport | | 1 | | | | |
| 3D-SMLM-COTS-M-462-XZ-M | Optomechanics | XZ manual stage 25 mm travel | Newport | | 1 | | | | |
| 3D-SMLM-COTS-AJS100 | Optomechanics | Actuator screw | Newport | | 2 | | | | |
| 3D-SMLM-COTS-M-SA2-04X12 | Optomechanics | 100 x 300 mm breadboard | Newport | | 4 | | | | |
| 3D-SMLM-COTS-AKP-BF | Optomechanics | Breadboard mounting kinematic base (lower) | Newport | | 1 | | | | |
| 3D-SMLM-COTS-AKP-BV | Optomechanics | Breadboard mounting kinematic base (lower) | Newport | | 1 | | | | |
| 3D-SMLM-COTS-AKP-BC | Optomechanics | Breadboard mounting kinematic base (lower) | Newport | | 1 | | | | |
| 3D-SMLM-COTS-AKP-TF | Optomechanics | Breadboard mounting kinematic base (upper) | Newport | | 12 | | | | |

| | | | | | | | | | |
|---|---|---|---|---|---|---|---|---|---|
| 3D-SMLM-COTS-PH30E-M | Optomechanics | 30 mm height, 1/2" post holder pedestal base | Thorlabs | | 8 | | | | |
| 3D-SMLM-COTS-SM1RC-M | Optomechanics | Post-mountable split clamp for SM1 lens tubes | Thorlabs | | 9 | | 2 | | |
| 3D-SMLM-COTS-PH20-M | Optomechanics | 20 mm height 1/2" post holder M6 tap | Thorlabs | | 2 | | | | |
| 3D-SMLM-COTS-POLARIS-K1E3 | Optomechanics | Polaris 1" mirror mount with 3 adjusters | Thorlabs | | 1 | | | | |
| 3D-SMLM-COTS-RS10M | Optomechanics | 10 mm height, 25 mm diameter post spacer | Thorlabs | | | 1 | 2 | | |
| 3D-SMLM-COTS-CCM1-4ER-M | Optomechanics | 30 mm cage cube for cube beamsplitters | Thorlabs | | | 1 | | | |
| 3D-SMLM-COTS-SM1A6 | Optomechanics | SM1 external to SM05 thread adapter | Thorlabs | | | 1 | | | |
| 3D-SMLM-COTS-P15K | Optomechanics | 1" 15 µm pinhole | Thorlabs | | | 1 | | | |
| 3D-SMLM-COTS-CP32-M | Optomechanics | 30 mm cage plate SM05 tap | Thorlabs | 1 | | | | | |
| 3D-SMLM-COTS-ER2 | Optomechanics | 2" cage rod | Thorlabs | 2 | | | | | |
| 3D-SMLM-COTS-RS1P4M | Optomechanics | 25 mm height , 25 mm diameter pedestal post with M4 taps | Thorlabs | 4 | | | | | |
| 3D-SMLM-COTS-RS4M | Optomechanics | 4 mm height, 25 mm diameter post spacer | Thorlabs | 3 | | | | | |
| 3D-SMLM-COTS-RS1M | Optomechanics | 1 mm height, 25 mm diameter post spacer | Thorlabs | 1 | | | | | |
| 3D-SMLM-COTS-KM100DL | Optomechanics | D-shaped mirror kinematic mount | Thorlabs | 1 | | | | | |
| 3D-SMLM-COTS-SM1FC | Optomechanics | FC/PC terminated fiber adapter | Thorlabs | | | | 1 | | |
| 3D-SMLM-COTS-SM1ZM | Optomechanics | SM1 threaded focusing mount | Thorlabs | | | | 1 | | |
| 3D-SMLM-COTS-SM1M20 | Optomechanics | 2" length lens tube with SM1 internal threads | Thorlabs | | | | 1 | | |
| 3D-SMLM-COTS-PH75E-M | Optomechanics | 75 mm height, 1/2" post holder pedestal base | Thorlabs | | | | 2 | | |
| 3D-SMLM-COTS-RS3P4M | Optomechanics | 75 mm height, 25 mm diameter pedestal post with M4 taps | Thorlabs | | | | 2 | | |
| 3D-SMLM-COTS-SM2M25 | Optomechanics | 2.5" length lens tube with SM2 internal threads | Thorlabs | | | | 1 | | |
| 3D-SMLM-COTS-SM2RC-M | Optomechanics | Post-mountable split clamp for SM2 lens tubes | Thorlabs | | | | 2 | | |
| 3D-SMLM-COTS-AD2 | Optomechanics | 1" to 2" optics adapter | Thorlabs | | | | 1 | | |
| 3D-SMLM-COTS-SM2L20 | Optomechanics | 2" length lens tube with SM2 internal and external threads | Thorlabs | | | | 1 | | |
| 3D-SMLM-COTS-SM1L30C | Optomechanics | 3" length slotted lens tube with SM1 internal and external threads | Thorlabs | | | | | 3 | |
| 3D-SMLM-COTS-SM1A2 | Optomechanics | SM1 internal to SM2 internal thread adapter | Thorlabs | | | | | 1 | |
| 3D-SMLM-COTS-TRA50-M | Optomechanics | 50 mm length 1/2" post | Thorlabs | | | | | 2 | |
| 3D-SMLM-COTS-SICP | Optomechanics | 30 mm cage adapter for shearing interferometer | Thorlabs | | | | | 1 | |
| 3D-SMLM-COTS-SM1A4 | Optomechanics | external RMS to internal SM1 thread adapter | Thorlabs | | | | | 1 | |
| 3D-SMLM-COTS-SM1A39 | Optomechanics | external SM1 to external c-mount thread adapter | Thorlabs | | | | | 1 | |
| 3D-SMLM-COTS-KAD11F | Optomechanics | SM1 external threaded pitch/yaw adapter for 11 mm diameter components | Thorlabs | | | | | 1 | |
| 3D-SMLM-COTS-SM1S10 | Optomechanics | 1" length lens tube with SM1 external threads | Thorlabs | | | | | 1 | |
| 3D-SMLM-COTS-SM1L10 | Optomechanics | 1" length lens tube with SM1 internal and external threads | Thorlabs | | | | | 1 | |
| 3D-SMLM-COTS-PS-F | Optomechanics | Pedestal post clamping fork | Newport | 13 | 10 | 7 | 7 | 3 | |
| 3D-SMLM-COTS-PS-0.125 | Optomechanics | 1/8" post height spacer | Newport | | | | | 1 | |
| 3D-SMLM-COTS-SM1A9 | Optomechanics | SM1 internal to c-mount external thread adapter | Thorlabs | | | | | 1 | |
| 3D-SMLM-COTS-SM1L30 | Optomechanics | 3.0" length lens tube with SM1 internal and external threads | Thorlabs | | | | | 1 | |
| 3D-SMLM-COTS-SM1L15 | Optomechanics | 1.5" length lens tube with SM1 internal and external threads | Thorlabs | | | | | 1 | |
| 3D-SMLM-COTS-SM1M30 | Optomechanics | 3.0" length lens tube with SM1 internal threads | Thorlabs | | | | | 1 | |
| 3D-SMLM-COTS-SM1M15 | Optomechanics | 1.5" length lens tube with SM1 internal threads | Thorlabs | | | | | 1 | |
| 3D-SMLM-COTS-SM1T10 | Optomechanics | SM1 lens tube coupler | Thorlabs | | | | | 1 | |
| 3D-SMLM-COTS-SM1RR | Optomechanics | SM1 retaining ring | Thorlabs | | | | | 2 | More will be supplied with associated parts (e.g. lens tubes/cage plates) than required. |
| 3D-SMLM-COTS-R-27-16-05-N | Consumables | Ring magnet 16/26.75 mm ID/OD, 5 mm height | Supermagnete | | | | | | As many as required |
| 3D-SMLM-COTS-#1.5-PRECISION-COVERSLIP | Consumables | 24 mm diameter, #1.5 precision coverslip +/- 5 µm thickness | Various | | | | | | As many as required, high precision e.g. Marienfeld Superior 0117640 |
| 3D-SMLM-COTS-12X12MM-COVERSLIP | Consumables | 12 x 12 mm coverslip | Various | | | | | | As many as required, e.g. Marienfeld Superior 0101000 (lower precision options are sufficient here) |
| 3D-SMLM-COTS-IMMOIL-F30CC | Consumables | Immersion oil (n = 1.51) | Olympus | 1 | | | | | As appropriate for objective lens |
| 3D-SMLM-COTS-SIL300CS-30CC | Consumables | Silicone immersion oil (n = 1.41) | Olympus | 1 | | | | | As appropriate for objective lens |
| 3D-SMLM-COTS-VACUUM-GREASE | Consumables | Vacuum grease/lubricant | Various | 1 | | | | | |
| 3D-SMLM-COTS-NOA65 | Consumables | UV cure epoxy | Thorlabs | 1 | | | | | |
| 3D-SMLM-COTS-JB-WELD | Consumables | Two part epoxy | Various | 1 | | | | | Many other two part epoxy adhesives are suitable |
| 3D-SMLM-COTS-C15440-20UP | Radiometry | Orca Fusion BT sCMOS camera (USB3.0) | Hamamatsu | 1 | | | | | |
| 3D-SMLM-COTS-SM1PD1A | Radiometry | SM1 threaded photodiode | Thorlabs | 1 | | | | | Only required if using the laser power monitoring photodiode |
| 3D-SMLM-COTS-SD197-23-21-041 | Radiometry | Si quad cell photodiode (5 mm) TO-8 | LaserComponents | 1 | | | | | |
| 3D-SMLM-COTS-LC301DQD-PV | Radiometry | Amplifier for quad cell photodiodes | LaserComponents | 1 | | | | | |
| 3D-SMLM-COTS-ICHROME-MLE | Lasers | Single-mode fiber-coupled laser engine (405/488/561/640 nm typical) | Toptica | | 1 | | | | |
| 3D-SMLM-COTS-IBEAM-SMART | Lasers | Single-mode free-space laser head (640 nm typical) | Toptica | | | 1 | | | |
| 3D-SMLM-COTS-IBEAM-SMART-PT | Lasers | Single-mode fiber-pigtailed laser engine (808 nm) | Toptica | 1 | | | | | |
| 3D-SMLM-COTS-CPS532 | Lasers | Single-mode free-space compact laser (532 nm) | Thorlabs | | | | | | |
| 3D-SMLM-COTS-M04 2500000004 | Electronics | Meder magnetic door switch magnet | Buerklin | 2 | | | | | Only used if employing laser safety interlock with cover |
| 3D-SMLM-COTS-MK04-1A66B-2000W 224271 | Electronics | Meder magnetic door switch wired switch | Buerklin | 2 | | | | | Only used if employing laser safety interlock with cover |
| 3D-SMLM-COTS-90327-0308 | Electronics | 8-pin connector | Mouser | 8 | | | | | Manufacturer: Molex, 2 used for ELL9 shared path filter slider |
| 3D-SMLM-COTS-AWG 28-08G 3M | Electronics | 8 x 1.27 mm pitch ribbon cable (3m) | Reichelt | 1 | | | | | |
| 3D-SMLM-COTS-TBS1052C | Electronics | 2 channel 50 MHz oscilloscope | Tektronix | | | | | 1 | Only required if self-building/testing microFPGA. Otherwise source microFPGA directly from EMBL/EMBLEM |
| 3D-SMLM-COTS-PEAKTECH-6227 | Electronics | Variable lab power supply | Reichelt | | | | | 1 | Only required if self-building/testing microFPGA. Otherwise source microFPGA directly from EMBL/EMBLEM |
| 3D-SMLM-COTS-FC-102520-GS | Electronics | microFPGA enclosure | Daub CNC Technik ( | 1 | | | | | Only required if self-building/testing microFPGA. Otherwise source microFPGA directly from EMBL/EMBLEM |
| 3D-SMLM-COTS-681183 | Electronics | USB 2.0 Type-C 22 mm diameter, 15 cm cable length connector | Ribu Elektronik | 1 | | | | | Only required if self-building/testing microFPGA. Otherwise source microFPGA directly from EMBL/EMBLEM |

| | | | | | | | | | |
|---|---|---|---|---|---|---|---|---|---|
| 3D-SMLM-COTS-AU-FPGA | Electronics | Alchitry Au FPGA development board (DEV-16527) | Sparkfun | 1 | | | | | Only required if self-building/testing microFPGA. Otherwise source microFPGA directly from EMBL/EMBLEM |
| 3D-SMLM-COTS-BR-SHIELD | Electronics | Alchitry Br prototype shield (DEV-16254) | Sparkfun | 1 | | | | | Only required if self-building/testing microFPGA. Otherwise source microFPGA directly from EMBL/EMBLEM |
| 3D-SMLM-COTS-GSM18E12-P1J | Electronics | Mean Well 12V, 1.5 A power supply | RS | 1 | | | | | Only required if self-building/testing microFPGA. Otherwise source microFPGA directly from EMBL/EMBLEM |
| 3D-SMLM-COTS-99-0402-00-02 | Electronics | microFPGA power supply connector | Farnell | 1 | | | | | Only required if self-building/testing microFPGA. Otherwise source microFPGA directly from EMBL/EMBLEM |
| 3D-SMLM-COTS-09-0403-00-02 | Electronics | microFPGA enclosure power connector | Farnell | 1 | | | | | Only required if self-building/testing microFPGA. Otherwise source microFPGA directly from EMBL/EMBLEM |
| 3D-SMLM-COTS-661002113322 | Electronics | 2 pin cable connector | Würth Elektronik | 55 | | | | | Only required if self-building/testing microFPGA. Otherwise source microFPGA directly from EMBL/EMBLEM |
| 3D-SMLM-COTS-661161122030 | Electronics | cable 30 cm both ends crimped | Würth Elektronik | 55 | | | | | Only required if self-building/testing microFPGA. Otherwise source microFPGA directly from EMBL/EMBLEM |
| 3D-SMLM-COTS-INT-714020 | Electronics | Power strip (EU) 4 sockets | Reichelt | 3 | | | | | Example. Manufacturer: Intellinet |
| 3D-SMLM-COTS-UA0148 | Electronics | 7 port industrial USB hub | Reichelt | 1 | | | | | Example. Manufacturer: Logilink |
| 3D-SMLM-COTS-61860 | Electronics | USB 2.0 to 8x RS232 serial hub | Reichelt | 1 | | | | | Example. Manufacturer: Delock |
| 3D-SMLM-COTS-3616111 | Electronics | Heating foil 92 mm outer diameter | Conrad | 1 | | | | | Manufacturer: Thermo Technologies, Only required if using the heating foil |
| 3D-SMLM-COTS-GST18E28-P1J | Electronics | Mean Well 18W, 0.64 A power supply | Reichelt | 1 | | | | | Only required if self-building/testing LED and controller. Otherwise source directly from EMBL/EMBLEM |
| 3D-SMLM-COTS-FC-101010-GS | Electronics | LED controller enclosure | Daub CNC Technik | 1 | | | | | Only required if self-building/testing LED and controller. Otherwise source directly from EMBL/EMBLEM |
| 3D-SMLM-COTS-9904020002 | Electronics | 2 pin cable connector | Farnell | 1 | | | | | Only required if self-building/testing LED and controller. Otherwise source directly from EMBL/EMBLEM |
| 3D-SMLM-COTS-9904010002 | Electronics | 2 pin cable connector | Farnell | 1 | | | | | Only required if self-building/testing LED and controller. Otherwise source directly from EMBL/EMBLEM |
| 3D-SMLM-COTS-0904030002 | Electronics | 2 pin flange socket | Farnell | 1 | | | | | Only required if self-building/testing LED and controller. Otherwise source directly from EMBL/EMBLEM |
| 3D-SMLM-COTS-0904040002 | Electronics | 2 pin flange socket | Farnell | 1 | | | | | Only required if self-building/testing LED and controller. Otherwise source directly from EMBL/EMBLEM |
| 3D-SMLM-COTS-J01160A0021 | Electronics | Coaxial connector | Buerklin | 4 | | | | | Only required if self-building/testing LED and controller. Otherwise source directly from EMBL/EMBLEM |
| 3D-SMLM-COTS-534-102-1K0 | Electronics | 1 kOhm potentiometer | Buerklin | 1 | | | | | Only required if self-building/testing LED and controller. Otherwise source directly from EMBL/EMBLEM |
| 3D-SMLM-COTS-2606 | Electronics | Potentiometer knob | Buerklin | 1 | | | | | Only required if self-building/testing LED and controller. Otherwise source directly from EMBL/EMBLEM |
| 3D-SMLM-COTS-SPW602 | Tools | SM1 spanner wrench | Thorlabs | | | | | 1 | |
| 3D-SMLM-COTS-SPW801 | Tools | Adjustable spanner wrench | Thorlabs | | | | | 1 | |
| 3D-SMLM-COTS-7426A48 | Tools | Spring pulling hooks | McMaster | | | | | 1 | |
| 3D-SMLM-COTS-RS7M | Tools | 7 mm height, 25 mm diameter post spacer | Thorlabs | | | | | 3 | |
| 3D-SMLM-COTS-CPA1 | Tools | Cage alignment target (0.9 mm hole) | Thorlabs | | | | | 1 | |
| 3D-SMLM-COTS-CPA2 | Tools | Cage alignment target (5.0 mm hole) | Thorlabs | | | | | 1 | |
| 3D-SMLM-COTS-PM100D | Tools | Digital laser power meter | Thorlabs | | | | | 1 | |
| 3D-SMLM-COTS-S121C | Tools | Photodiode power sensor (up to 500 mW) | Thorlabs | | | | | 1 | |
| 3D-SMLM-COTS-LASERCAM-HRII | Tools | Beam profiling camera with laser flat ND filter | Coherent | | | | | 1 | |
| 3D-SMLM-COTS-VRC2 | Tools | NIR beam viewer card | Thorlabs | | | | | 1 | |
| 3D-SMLM-COTS-LVL01 | Tools | Bubble level | Thorlabs | | | | | 1 | |
| 3D-SMLM-COTS-LAB16/LAB16-EU | Tools | Compressed air optics grade | Newport | | | | | 1 | |
| 3D-SMLM-COTS-TGP1 | Tools | Non-marring pliers | Thorlabs | | | | | 1 | |
| 3D-SMLM-COTS-DIGC6 | Tools | Digital calliper | Thorlabs | | | | | 1 | Manufacturer: Mitutoyo |
| 3D-SMLM-COTS-LG12 | Tools | Laser safety glasses | Thorlabs | | | | | 1 | For use at 405, 488, 640, 808 nm |
| 3D-SMLM-COTS-NIR-ZS2-900 | Tools | Laser safety glasses | Laser2000 | | | | | 1 | For use at 561 nm |
| 3D-SMLM-COTS-TPSM1-M | Tools | Magnetic laser safety screen | Thorlabs | | | | | 1 | |
