## Supplementary Table 3 for "Automated 3D multi-color single-molecule localization microscopy"

| Module | Budgetary item | Price /EUR * |
|---|---|---|
| Core | PC | 2000 |
| | microFPGA (self-sourced) | 1000 |
| | Cables and connectors | 250 |
| | Optical table | 5500 |
| | Custom mechanical parts (body) | 7500 |
| | Smaract SOM-12090 XY stage | 11000 |
| | PI P-726.1CD PIFOC | 7500 |
| | Dichroic mirrors and notch filter | 2000 |
| | Basic optics and optomechanics and mechanical | 1500 |
| | **TOTAL** | 38250 |
| Objective lenses | Olympus 100x/1.5 oil objective | 13500 |
| | Olympus 100x/1.45 oil objective | 6000 |
| | Olympus 100x/1.35 silicone objective | 9000 |
| Emission | Filters and dichroic mirror | 3500 |
| | Filter wheels | 2500 |
| | Custom mechanical parts (splitter) | 2000 |
| | Thorlabs ELL20/M piezo stage | 400 |
| | Thorlabs ELL6 slider | 400 |
| | Thorlabs ELL9 slider | 400 |
| | Thorlabs widefield tube lens | 750 |
| | Basic optics and optomechanics | 1750 |
| | **TOTAL** | 11700 |
| Camera | Hamamatsu Fusion BT sCMOS camera | 20000 |
| | Hamamatsu Fusion sCMOS camera | 14500 |
| SM engine illumination | Thorlabs ELL17/M piezo stage | 400 |
| | Toptica iChrome MLE laser engine | 44000 |
| | Basic optics and optomechanics | 2250 |
| | ADL optica piShaper refractive beam shaper | 4000 |
| | X,Z, pitch/yaw stage assembly | 2000 |
| | Swappable telescope (cost for two) | 1000 |
| | Filters | 500 |
| | **TOTAL** | 54150 |
| SM booster laser | Booster laser | 3500 |
| | Basic optics and optomechanics | 2000 |
| | Custom mechanical parts (laser mount) | 100 |
| | **TOTAL** | 5600 |
| MM laser | Multi-mode laser engine | 3000 |
| | Basic optics and optomechanics | 1500 |
| | **TOTAL** | 4500 |
| Focus lock | 808 nm laser | 5500 |
| | Blocking filters | 500 |
| | Basic optics and optomechanics | 1000 |
| | 1D stage** | 1200 |
| | QPD and amplifier | 750 |
| | **TOTAL** | 8950 |

| Configuration | Total cost /EUR * | Equivalent /US dollar *** |
|---|---|---|
| Core, Emission, Orca Fusion, MM laser engine, Focus Lock, 100x/1.45 (low cost) | 83900 | 95126 |
| Core, Emission, Orca Fusion BT, SM laser engine, Focus Lock, 100x/1.5 (base configuration) | 146550 | 166158 |
| Core, Emission, Orca Fusion BT, SM laser engine, Booster Focus Lock, 100x/1.5, 100x/1.35 (high-end) | 161150 | 182712 |

| |
|---|
| * All prices exclude sales taxes/VAT. Prices are subject to change. |
| ** Must be combined with Smaract SOM-12090 XY stage since controllers are shared. If another XY stage is to be used, other options should be sought for the 1D focus lock stage. |
| *** US dollar equivalent based on 5 year average exchange rate. This figure is provided for guidance only. Certain hardware may be more or less expensive in certain jurisdictions due to import duties and other factors. |
