## Supplementary Table 4 for "Automated 3D multi-color single-molecule localization microscopy"

| Hardware configuration and communications settings | | | | | | | | |
|---|---|---|---|---|---|---|---|---|
| **Manufacturer** | **Hardware** | **uManager Device Module/Adapter** | **Baud** | **Bits** | **Handshake** | **Parity** | **StopBits** | **Other settings and notes** |
| PI | E-709.CRG | PI_FocusLock/PIZStage | 57600 | 8 | Off | None | 1 | Axis = Z, Limit-um = 100 |
| Toptica | iChrome MLE | Toptica_iChrome_MLE/iChromeMLE | 115200 | 8 | Off | None | 1 | |
| Toptica | iBeam 640 nm | Toptica_iBeamSmartCW/iBeamSmartCW | 115200 | 8 | Off | None | 1 | Note use Normal Mode adapter to restore iBeam functionality |
| Toptica | iBeam 808 nm | Toptica_iBeamSmartCW/iBeamSmartCW | 115200 | 8 | Off | None | 1 | Note use Normal Mode adapter to restore iBeam functionality |
| Thorlabs | FW102C | Thorlabs Filter Wheel/Thorlabs Filter Wheel | 115200 | 8 | Off | None | 1 | |
| Thorlabs | FW102C | Thorlabs Filter Wheel/Thorlabs Filter Wheel | 115200 | 8 | Off | None | 1 | |
| Thorlabs | ELL6 | Thorlabs ElliptecSlider/Thorlabs ELL6 | 9600 | 8 | Off | None | 1 | Bertrand lens, Channel = 2 |
| Thorlabs | ELL9 | Thorlabs ElliptecSlider/Thorlabs ELL9 | 9600 | 8 | Off | None | 1 | Shared path filter slider, Channel = 3 (optional) |
| Thorlabs | ELL17/M | Thorlabs ElliptecSlider/Thorlabs ELL17 | 9600 | 8 | Off | None | 1 | Illumination inclination control stage (HILO/TIRF), Channel = 1 |
| Thorlabs | ELL20/M | Thorlabs ElliptecSlider/Thorlabs ELL17 | 9600 | 8 | Off | None | 1 | 3D astigmatic lens stage, Channel = 0 |
| Smaract | XY stage | SmarActHCU-3D/SmarAct 2D | 9600 | 8 | Off | None | 1 | X channel = 0, X direction = 1, Y channel = 1, Y direction = 1 |
| Smaract | QPD stage | SmarActHCU-3D/SmarAct 1D | 9600 | 8 | Off | None | 1 | Z channel = 2, Z direction = 1 |
| EMBL | microFPGA-Hub | MicroFPGA/MicroFPGA-Hub | 57600 | 8 | Off | None | 1 | |
| EMBL | microFPGA-LaserTrig | | | | | | | Number of lasers = 4 |
| EMBL | microFPGA-AnalogInput | | | | | | | Number of channels = 4 |
| EMBL | microFPGA-Servos | | | | | | | Number of servos = 4 |
| EMBL | microFPGA-PWM | | | | | | | Number of PWM = 4 |
| EMBL | microFPGA-TTL | | | | | | | Number of channels = 4 |
| Hamamatsu | Fusion BT | HamamatsuHam_DCAM | | | | | | |

| Other settings | | |
|---|---|---|
| Default Camera | Hamamatsu_DCAM | |
| Default Shutter | | |
| Default Focus Stage | PIZStage | |
| Stage focus directions | All 'Unknown' | |
| Delay | iChromeMLE | Delay[ms] = 500 ms |
| | Thorlabs Filter Wheels | Delay[ms] = 2000 ms |
| iChrome MLE State 0 | State-0 | |
| iChrome MLE State 1 | State-1 | |
| iChrome MLE State 2 | State-2 | |
| Thorlabs Filter Wheel State 0 | Filter-1 | |
| Thorlabs Filter Wheel State 1 | Filter-2 | |
| Thorlabs Filter Wheel State 2 | Filter-3 | |
| Thorlabs Filter Wheel State 3 | Filter-4 | |
| Thorlabs Filter Wheel State 4 | Filter-5 | |
| Thorlabs Filter Wheel State 5 | Filter-6 | |
| Thorlabs ELL6 State 0 | Position 0 | |
| Thorlabs ELL6 State 1 | Position 1 | |
| Thorlabs ELL9 State 0 | Position 0 | optional |
| Thorlabs ELL9 State 1 | Position 1 | optional |
| Thorlabs ELL9 State 2 | Position 2 | optional |
| Thorlabs ELL9 State 3 | Position 3 | optional |
