## Supplementary Table 5 for "Automated 3D multi-color single-molecule localization microscopy"

| Hardware | microFPGA I/O | Intended functionality | Associated designation on custom shield |
|---|---|---|---|
| Camera | Digital input | Camera exposure for active camera synchronisation | P_Cam |
| Camera | Digital output | Camera trigger for passive camera synchronisation | P_TTL4 |
| Single-mode laser engine (L0 - L3) | Digital output | Laser enable | P_L0 - P_L3 |
| Booster laser (L0/1/2/3) | Digital output | Laser enable | P_L0/1/2/3 |
| Multi-mode laser engine (L0 - L4) | Digital output | Laser enable | P_L0 - P_L4 |
| Multi-mode laser engine (L0 - L3) | Pulse-width modulation | Laser power modulation | P_P0 - P_P3 |
| Quadrant photodiode | Analog input (0 - 10 V converted to 0 - 1 V) | X signal from QPD | P_AI0 |
| Quadrant photodiode | Analog input (0 - 10 V converted to 0 - 1 V) | Y signal from QPD | P_AI1 |
| Quadrant photodiode | Analog input (0 - 10 V converted to 0 - 1 V) | SUM signal from QPD | P_AI2 |
| Photodiode | Analog input (voltage across shunt resistor from photodiode current truncated to 0 - 1 V) | Laser power measurement | P_AI3 |
