## Supplementary Table 7 for "Automated 3D multi-color single-molecule localization microscopy"

| **Possible hardware** | **microFPGA I/O** | **Intended functionality** | **Associated designation on custom shield** |
|---|---|---|---|
| Flip mirror | Digital output x4 | TTL level control of binary state hardware | P_TTl0 - P_TTL3 |
| Servo motors | PWM x 7 | Control of servo motor position | P_S0 - P_S6 |
| Acousto-optical modulators/tunable filters | Analog output (via low pass filter of PWM on signal conversion board) | Control of continuously modulatable hardware | P_S0 - P_S6 |
| Additional lasers | Digital output | Laser enable | P_L4 - P_L7 |
