## Supplementary Table 8 for "Automated 3D multi-color single-molecule localization microscopy"

**Normal Mode**

| Property | Channel | Setting | Units |
|---|---|---|---|
| Hamamatsu_DCAM, DEFECT CORRECT MODE | | OFF (0) | |
| Hamamatsu_DCAM, Exposure | | 100 | ms |
| Hamamatsu_DCAM, HOT PIXEL CORRECT LEVEL | | MINIMUM | |
| Hamamatsu_DCAM, OUTPUT TRIGGER KIND | 0 | EXPOSURE | |
| Hamamatsu_DCAM, OUTPUT TRIGGER POLARITY | 0 | Positive | |
| Hamamatsu_DCAM, ScanMode | | 2 | |
| iChrome-MLE, Laser 1: 4. Use TTL | | 1 | |
| iChrome-MLE, Laser 2: 4. Use TTL | | 1 | |
| iChrome-MLE, Laser 3: 4. Use TTL | | 1 | |
| iChrome-MLE, Laser 4: 4. Use TTL | | 1 | |

**Fast Mode**

| Property | Channel | Setting | Units |
|---|---|---|---|
| Hamamatsu_DCAM, DEFECT CORRECT MODE | | OFF (0) | |
| Hamamatsu_DCAM, Exposure | | 25 | |
| Hamamatsu_DCAM, HOT PIXEL CORRECT LEVEL | | MINIMUM | |
| Hamamatsu_DCAM, OUTPUT TRIGGER KIND | 0 | EXPOSURE | |
| Hamamatsu_DCAM, OUTPUT TRIGGER POLARITY | 0 | Positive | |
| Hamamatsu_DCAM, ScanMode | | 3 | |
| iChrome-MLE, Laser 1: 4. Use TTL | | 1 | |
| iChrome-MLE, Laser 2: 4. Use TTL | | 1 | |
| iChrome-MLE, Laser 3: 4. Use TTL | | 1 | |
| iChrome-MLE, Laser 4: 4. Use TTL | | 1 | |

**Slow Mode**

| Property | Channel | Setting | Units |
|---|---|---|---|
| Hamamatsu_DCAM, DEFECT CORRECT MODE | | OFF (0) | |
| Hamamatsu_DCAM, Exposure | | 250 | |
| Hamamatsu_DCAM, HOT PIXEL CORRECT LEVEL | | MINIMUM | |
| Hamamatsu_DCAM, OUTPUT TRIGGER KIND | 0 | EXPOSURE | |
| Hamamatsu_DCAM, OUTPUT TRIGGER POLARITY | 0 | Positive | |
| Hamamatsu_DCAM, ScanMode | | 1 | |
| iChrome-MLE, Laser 1: 4. Use TTL | | 1 | |
| iChrome-MLE, Laser 2: 4. Use TTL | | 1 | |
| iChrome-MLE, Laser 3: 4. Use TTL | | 1 | |
| iChrome-MLE, Laser 4: 4. Use TTL | | 1 | |

**Startup**

| Property | Channel | Setting | Units |
|---|---|---|---|
| Hamamatsu_DCAM, DEFECT CORRECT MODE | | OFF (0) | |
| Hamamatsu_DCAM, Exposure | | 100 | |
| Hamamatsu_DCAM, HOT PIXEL CORRECT LEVEL | | MINIMUM | |
| Hamamatsu_DCAM, OUTPUT TRIGGER KIND | 0 | EXPOSURE | |
| Hamamatsu_DCAM, OUTPUT TRIGGER POLARITY | 0 | Positive | |
| Hamamatsu_DCAM, ScanMode | | 2 | |
| iChrome-MLE, Laser 1: 4. Use TTL | | 1 | |
| iChrome-MLE, Laser 2: 4. Use TTL | | 1 | |
| iChrome-MLE, Laser 3: 4. Use TTL | | 1 | |
| iChrome-MLE, Laser 4: 4. Use TTL | | 1 | |
