## Supplementary Table 9 for "Automated 3D multi-color single-molecule localization microscopy"

**Plugin settings**

| | |
|---|---|
| Additional FW tab | Yes if using ELL9 |
| Additional FW tab title | Shared filters |
| Powermeter tab | Yes |
| QPD tab | Yes |
| Single FW panel | No |
| Trigger tab | Yes |
| iBeamSmart #1 | Yes if using booster |
| iBeamSmart #1 name | XXX nm Booster |
| iBeamSmart #2 | Yes |
| iBeamSmart #2 name | Focus Lock Laser |

**Properties**

| **UI parameter** | **Value** | |
|---|---|---|
| XXX nm Booster - external trigger available | Yes | |
| XXX nm Booster - fine available | Yes | |
| XXX nm Booster - max power | XXX | mW |
| Acquisitions - BFP lens | Two-state device 4 | |
| Acquisitions - Bright field | Two-state device 5 | |
| Acquisitions - focus stabilization | Z stage focus locking | |
| Activation - Default feedback | 0.01 | Suited to dSTORM imaging |
| Activation - Default sd coeff | 4 | Suited to dSTORM imaging |
| Activation - Idle time (ms) | 100 | Suited to dSTORM imaging |
| Activation - Number of points | 30 | Suited to dSTORM imaging |
| Controls - Enable two-state device 1 | Yes | |
| Controls - Enable two-state device 2 | Yes | |
| Controls - Enable two-state device 3 | Yes | |
| Controls - Enable two-state device 4 | Yes | |
| Controls - Enable two-state device 5 | Yes | |

| | | |
|---|---|---|
| Controls - Enable two-state device 6 | Yes | |
| Controls - Title | Controls | |
| Controls - Enable two-state device 1 name | 3D | |
| Controls - Enable two-state device 2 name | TIRF | |
| Controls - Enable two-state device 3 name | HILO | |
| Controls - Enable two-state device 4 name | BFP | |
| Controls - Enable two-state device 5 name | Brightfield | |
| Controls - Enable two-state device 6 name | Multi-Mode | |
| Filters - Filter colors | blue, green, red, grey, grey, grey | Example configuration |
| Filters - Filter colors 2 | red, grey, grey, grey, grey, grey | Example configuration |
| Filters - Filter names | 525/50, 600/52, 676/37, None, None, None | Example configuration |
| Filters - Filter names 2 | 685/70, None, None, None, None, None | Example configuration |
| Filters - Panel title | Filters | |
| Focus - Idle time (ms) | 100 | |
| Focus - Large step | 2 | microns |
| Focus - Number of points | 30 | |
| Focus - Small step | 0.2 | microns |
| Focus Lock Laser - external trigger available | No | |
| Focus Lock Laser - fine available | Yes | |
| Focus Lock laser - max power | 75 | mW |

**Single-mode Toptica MLE laser engine**

| **UI parameter** | **Value** | |
|---|---|---|
| Laser 0 - Color | violet | Example configuration, Laser 0 is always activation laser |
| Laser 0 - Default max pulse | 10000 | Example configuration, Laser 0 is always activation laser |
| Laser 0 - Name | 405 | Example configuration, Laser 0 is always activation laser |
| Laser 0 - Use on/off | Yes | Example configuration, Laser 0 is always activation laser |
| Laser 0 - Use slider | Yes | Example configuration, Laser 0 is always activation laser |
| Laser 0 trigger - Color | violet | Example configuration, Laser 0 is always activation laser |
| Laser 0 trigger - Name | 405 | Example configuration |
| Laser 1 - Color | blue | Example configuration |

| | | |
|---|---|---|
| Laser 1 - Name | 488 | Example configuration |
| Laser 1 - Use on/off | yes | Example configuration |
| Laser 1 - Use slider | yes | Example configuration |
| Laser 1 trigger - Color | blue | Example configuration |
| Laser 1 trigger - Name | 488 | Example configuration |
| Laser 2 - Color | green | Example configuration |
| Laser 2 - Name | 561 | Example configuration |
| Laser 2 - Use on/off | yes | Example configuration |
| Laser 2 - Use slider | yes | Example configuration |
| Laser 2 trigger - Color | green | Example configuration |
| Laser 2 trigger - Name | 561 | Example configuration |
| Laser 3 - Color | red | Example configuration |
| Laser 3 - Name | 640 | Example configuration |
| Laser 3 - Use on/off | yes | Example configuration |
| Laser 3 - Use slider | yes | Example configuration |
| Laser 3 trigger - Color | red | Example configuration |
| Laser 3 trigger - Name | 640 | Example configuration |

**Multi-mode laser engine**

| **UI parameter** | **Value** | |
|---|---|---|
| Laser 0 - Color | violet | Example configuration, Laser 0 is always activation laser |
| Laser 0 - Default max pulse | 10000 | Example configuration, Laser 0 is always activation laser |
| Laser 0 - Name | 405 | Example configuration, Laser 0 is always activation laser |
| Laser 0 - Use on/off | No | Example configuration, Laser 0 is always activation laser |
| Laser 0 - Use slider | Yes | Example configuration, Laser 0 is always activation laser |
| Laser 0 trigger - Color | violet | Example configuration, Laser 0 is always activation laser |
| Laser 0 trigger - Name | 405 | Example configuration |
| Laser 1 - Color | blue | Example configuration |
| Laser 1 - Name | 488 | Example configuration |
| Laser 1 - Use on/off | No | Example configuration |
| Laser 1 - Use slider | yes | Example configuration |

| | | |
|---|---|---|
| Laser 1 trigger - Color | blue | Example configuration |
| Laser 1 trigger - Name | 488 | Example configuration |
| Laser 2 - Color | red | Example configuration |
| Laser 2 - Name | 640(1) | Example configuration |
| Laser 2 - Use on/off | No | Example configuration |
| Laser 2 - Use slider | yes | Example configuration |
| Laser 2 trigger - Color | | laser typically in the range 488 < lambda0 < 640, (e.g. 561 nm) which can then be triggered appropriately and either controlled external to uManager or with |
| Laser 2 trigger - Name | | |
| Laser 3 - Color | red | Example configuration |
| Laser 3 - Name | 640(2) | Example configuration |
| Laser 3 - Use on/off | No | Example configuration |
| Laser 3 - Use slider | yes | Example configuration |
| Laser 3 trigger - Color | red | Example configuration |
| Laser 3 trigger - Name | 640 (1,2) | Example configuration |

| | | |
|---|---|---|
| Powermeter - idle time (ms) | 1000 | Example configuration |
| Powermeter - number of points | 100 | Example configuration |
| Powermeter - offsets | 0,0,0,0 | Requires calibration |
| Powermeter - slopes | 1,1,1,1 | Requires calibration |
| Powermeter - wavelengths | 405,488, 561, 640 | Example configuration |
| QPD idle time (ms) | 100 | |
| QPD - XY max | 65535 | |
| QPD - Z max | 65535 | |

**Global Settings**

| **Setting** | **Value** |
|---|---|
| Enable unallocated warnings | No |
