## Supplementary Table 10 for "Automated 3D multi-color single-molecule localization microscopy"

**Properties**

| UI property | Device | Property |
|---|---|---|
| XXX nm Booster enable fine | iBeam-XXXnm | Enable Fine |
| XXX nm Booster enable fine - On | | On |
| XXX nm Booster enable fine - Off | | Off |
| XXX nm Booster ext trigger | iBeam-XXXnm | Enable ext trigger |
| XXX nm Booster ext trigger - On | | On |
| XXX nm Booster ext trigger - Off | | Off |
| XXX nm Booster fine a (%) | iBeam-XXXnm | Fine A (%) |
| XXX nm Booster fine b (%) | iBeam-XXXnm | Fine B (%) |
| XXX nm Booster laser power | iBeam-XXXnm | Power (mW) |
| XXX nm Booster operation - On value | | On |
| XXX nm Booster laser power - Off value | | Off |
| Camera exposure | HamamatsuHam_DCAM | Exposure |
| Filter wheel 2 position | Thorlabs Filter Wheel-1 | State |
| Filter wheel 2 position state 0 | | 0 |
| Filter wheel 2 position state 1 | | 1 |
| Filter wheel 2 position state 2 | | 2 |
| Filter wheel 2 position state 3 | | 3 |
| Filter wheel 2 position state 4 | | 4 |
| Filter wheel 2 position state 5 | | 5 |
| Filter wheel position | Thorlabs Filter Wheel | State |
| Filter wheel position state 0 | | 0 |
| Filter wheel position state 1 | | 1 |
| Filter wheel position state 2 | | 2 |
| Filter wheel position state 3 | | 3 |
| Filter wheel position state 4 | | 4 |
| Filter wheel position state 5 | | 5 |
| Focus Lock Laser enable fine | iBeam-808nm | Enable Fine |
| Focus Lock Laser enable fine - On | | On |
| Focus Lock Laser enable fine - Off | | Off |

| | | |
|---|---|---|
| Focus Lock Laser ext trigger | iBeam-808nm | Enable ext trigger |
| Focus Lock Laser ext trigger - On | | On |
| Focus Lock Laser ext trigger - Off | | Off |
| Focus Lock Laser fine a (%) | iBeam-808nm | Fine A (%) |
| Focus Lock Laser fine b (%) | iBeam-808nm | Fine B (%) |
| Focus Lock Laser laser power | iBeam-808nm | Power (mW) |
| Focus Lock Laser operation - On value | | On |
| Focus Lock Laser laser power - Off value | | Off |

**Single-mode Toptica MLE laser engine**

| UI property | Device | Property |
|---|---|---|
| Laser 0 enable | iChrome-MLE | Laser 4: 1. Enable |
| Laser 0 enable - on value | | 1 |
| Laser 0 enable - off value | | 0 |
| Laser 0 power percentage | iChrome-MLE | Laser 4: 3. Level % |
| Laser 0 power percentage slope | | 1 |
| Laser 0 power percentage offset | | 0 |
| Laser 0 trigger mode | LaserTrig | Mode0 |
| Laser 0 trigger mode state 0 | | 0 |
| Laser 0 trigger mode state 1 | | 1 |
| Laser 0 trigger mode state 2 | | 2 |
| Laser 0 trigger mode state 3 | | 3 |
| Laser 0 trigger mode state 4 | | 4 |
| Laser 0 trigger pulse duration | LaserTrig | Duration0 |
| Laser 0 trigger pulse sequence | LaserTrig | Sequence0 |
| Laser 1 enable | iChrome-MLE | Laser 3: 1. Enable |
| Laser 1 enable - on value | | 1 |
| Laser 1 enable - off value | | 0 |
| Laser 1 power percentage | iChrome-MLE | Laser 3: 3. Level % |
| Laser 1 power percentage slope | | 1 |
| Laser 1 power percentage offset | | 0 |

| | | |
|---|---|---|
| Laser 1 trigger mode | LaserTrig | Mode1 |
| Laser 1 trigger mode state 0 | | 0 |
| Laser 1 trigger mode state 1 | | 1 |
| Laser 1 trigger mode state 2 | | 2 |
| Laser 1 trigger mode state 3 | | 3 |
| Laser 1 trigger mode state 4 | | 4 |
| Laser 1 trigger pulse duration | LaserTrig | Duration1 |
| Laser 1 trigger pulse sequence | LaserTrig | Sequence1 |
| Laser 2 enable | iChrome-MLE | Laser 2: 1. Enable |
| Laser 2 enable - on value | | 1 |
| Laser 2 enable - off value | | 0 |
| Laser 2 power percentage | iChrome-MLE | Laser 2: 3. Level % |
| Laser 2 power percentage slope | | 1 |
| Laser 2 power percentage offset | | 0 |
| Laser 2 trigger mode | LaserTrig | Mode2 |
| Laser 2 trigger mode state 0 | | 0 |
| Laser 2 trigger mode state 1 | | 1 |
| Laser 2 trigger mode state 2 | | 2 |
| Laser 2 trigger mode state 3 | | 3 |
| Laser 2 trigger mode state 4 | | 4 |
| Laser 2 trigger pulse duration | LaserTrig | Duration2 |
| Laser 2 trigger pulse sequence | LaserTrig | Sequence2 |
| Laser 3 enable | iChrome-MLE | Laser 1: 1. Enable |
| Laser 3 enable - on value | | 1 |
| Laser 3 enable - off value | | 0 |
| Laser 3 power percentage | iChrome-MLE | Laser 1: 3. Level % |
| Laser 3 power percentage slope | | 1 |
| Laser 3 power percentage offset | | 0 |
| Laser 3 trigger mode | LaserTrig | Mode3 |
| Laser 3 trigger mode state 0 | | 0 |
| Laser 3 trigger mode state 1 | | 1 |
| Laser 3 trigger mode state 2 | | 2 |

| | | |
|---|---|---|
| Laser 3 trigger mode state 3 | | 3 |
| Laser 3 trigger mode state 4 | | 4 |
| Laser 3 trigger pulse duration | LaserTrig | Duration3 |
| Laser 3 trigger pulse sequence | LaserTrig | Sequence3 |

**Multi-mode laser engine**

| **UI property** | **Device** | **Property** |
|---|---|---|
| Laser 0 enable | | |
| Laser 0 enable - on value | | |
| Laser 0 enable - off value | | |
| Laser 0 power percentage | PWM | Position0 |
| Laser 0 power percentage slope | | 0 |
| Laser 0 power percentage offset | | 2.55 |
| Laser 0 trigger mode | LaserTrig | Mode0 |
| Laser 0 trigger mode state 0 | | 0 |
| Laser 0 trigger mode state 1 | | 1 |
| Laser 0 trigger mode state 2 | | 2 |
| Laser 0 trigger mode state 3 | | 3 |
| Laser 0 trigger mode state 4 | | 4 |
| Laser 0 trigger pulse duration | LaserTrig | Duration0 |
| Laser 0 trigger pulse sequence | LaserTrig | Sequence0 |
| Laser 1 enable | | |
| Laser 1 enable - on value | | |
| Laser 1 enable - off value | | |
| Laser 1 power percentage | PWM | Position1 |
| Laser 1 power percentage slope | | 0 |
| Laser 1 power percentage offset | | 2.55 |
| Laser 1 trigger mode | LaserTrig | Mode1 |
| Laser 1 trigger mode state 0 | | 0 |
| Laser 1 trigger mode state 1 | | 1 |
| Laser 1 trigger mode state 2 | | 2 |

| | | |
|---|---|---|
| Laser 1 trigger mode state 3 | | 3 |
| Laser 1 trigger mode state 4 | | 4 |
| Laser 1 trigger pulse duration | LaserTrig | Duration1 |
| Laser 1 trigger pulse sequence | LaserTrig | Sequence1 |
| Laser 2 enable | | |
| Laser 2 enable - on value | | |
| Laser 2 enable - off value | | |
| Laser 2 power percentage | PWM | Position2 |
| Laser 2 power percentage slope | | 0 |
| Laser 2 power percentage offset | | 2.55 |
| Laser 2 trigger mode | LaserTrig | Mode2 |
| Laser 2 trigger mode state 0 | | 0 |
| Laser 2 trigger mode state 1 | | 1 |
| Laser 2 trigger mode state 2 | | 2 |
| Laser 2 trigger mode state 3 | | 3 |
| Laser 2 trigger mode state 4 | | 4 |
| Laser 2 trigger pulse duration | LaserTrig | Duration2 |
| Laser 2 trigger pulse sequence | LaserTrig | Sequence2 |
| Laser 3 enable | | |
| Laser 3 enable - on value | | |
| Laser 3 enable - off value | | |
| Laser 3 power percentage | PWM | Position3 |
| Laser 3 power percentage slope | | 0 |
| Laser 3 power percentage offset | | 2.55 |
| Laser 3 trigger mode | LaserTrig | Mode3 |
| Laser 3 trigger mode state 0 | | 0 |
| Laser 3 trigger mode state 1 | | 1 |
| Laser 3 trigger mode state 2 | | 2 |
| Laser 3 trigger mode state 3 | | 3 |
| Laser 3 trigger mode state 4 | | 4 |
| Laser 3 trigger pulse duration | LaserTrig | Duration3 |
| Laser 3 trigger pulse sequence | LaserTrig | Sequence3 |

| UI property | Device | Property |
|---|---|---|
| Laser powermeter | AnalogInput | AnalogInput3 |
| QPD X | AnalogInput | AnalogInput0 |
| QPD Y | AnalogInput | AnalogInput1 |
| QPD Z | AnalogInput | AnalogInput2 |
| Two-state device 1 | Thorlabs ELL17/ELL20 | Position (µm) |
| Two-state device 1 - On value | | [INSERT STAGE POSITION IN µm FOR 3D LENS IN EMISSION PATH] |
| Two-state device 1 - Off value | | [INSERT STAGE POSITION IN µm FOR 3D LENS NOT IN EMISSION PATH] |
| Two-state device 2 | Thorlabs ELL17/ELL20-1 | Position (µm) |
| Two-state device 2 - On value | | [INSERT STAGE POSITION IN µm FOR TIRF] |
| Two-state device 2 - Off value | | [INSERT STAGE POSITION IN µm FOR EPI-] |
| Two-state device 2 | Thorlabs ELL17/ELL20-1 | Position (µm) |
| Two-state device 2 - On value | | [INSERT STAGE POSITION IN µm FOR HILO] |
| Two-state device 2 - Off value | | [INSERT STAGE POSITION IN µm FOR EPI-] |
| Two-state device 4 | Thorlabs ELL6 | Position (µm) |
| Two-state device 4 - On value | | 1 |
| Two-state device 4 - Off value | | 0 |
| Two-state device 5 | TTL | State0 |
| Two-state device 5 - On value | | 0 |
| Two-state device 5 - Off value | | 1 |
| Two-state device 6 | Thorlabs ELL17/ELL20-1 | Position (µm) |
| Two-state device 6 - On value | | [INSERT STAGE POSITION IN µm FOR MULTI-MODE] |
| Two-state device 6 - Off value | | [INSERT STAGE POSITION IN µm FOR SINGLE-MODE] |
| UV pulse duration (activation) | LaserTrig | Duration0 |
| UV pulse duration (main frame) | LaserTrig | Duration0 |
| Z stage focus locking | PIZStage | External sensor |
| Z stage focus locking - On value | | 1 |
| Z stage focus locking - Off value | | 0 |
| Z stage position | PIZStage | Position |
