## Supplementary Table 11 for "Automated 3D multi-color single-molecule localization microscopy"

| | | | | | |
|---|---|---|---|---|---|
| | Maximum magnification of telescope | 0.5 | 1 | 2 | 2.85 |
| | Illumination field diameter at object /µm | 14 | 27 | 54 | 77 |
| | Illumination field diameter at camera /px | 127 | 254 | 508 | 723 |
| | Required offset of FOV from centre of chip /px | 63 | 127 | 254 | 362 |
| 2304 x 2304 px | reflected FOV centre position x /px | 1088 | 1025 | 898 | 790 |
| | reflected FOV centre position y /px | 1152 | 1152 | 1152 | 1152 |
| | transmitted FOV centre position x /px | 1216 | 1279 | 1406 | 1514 |
| | transmitted FOV centre position y /px | 1152 | 1152 | 1152 | 1152 |
| 2048 x 2048 px | reflected FOV centre position x /px | 960 | 897 | 770 | 662 |
| | reflected FOV centre position y /px | 1024 | 1024 | 1024 | 1024 |
| | transmitted FOV centre position x /px | 1088 | 1151 | 1278 | 1386 |
| | transmitted FOV centre position y /px | 1024 | 1024 | 1024 | 1024 |
