## Supplementary Table 12 for "Automated 3D multi-color single-molecule localization microscopy"

| Part number(s) lens 1 | Part number(s) lens 2 | Effective focal length lens 1 /mm | Effective focal length lens 2 /mm | Effective magnification | Approximate 4f track length /mm |
|---|---|---|---|---|---|
| 3D-SMLM-COTS-AC254-080-A | 3D-SMLM-COTS-AC254-040-A | 80 | 40 | 0.50 | 240 |
| 3D-SMLM-COTS-AC254-060-A | 3D-SMLM-COTS-AC254-060-A | 60 | 60 | 1.00 | 240 |
| 3D-SMLM-COTS-AC254-040-A | 3D-SMLM-COTS-AC254-080-A | 40 | 80 | 2.00 | 240 |
| 3D-SMLM-COTS-AC254-075-A, 3D-SMLM-COTS-AC254-050-A | 3D-SMLM-COTS-#49-769 | 31.2 | 88.9 | 2.85 | 240.2 |
